# An explicit birth-death-reticulation model for studying the diversification of phylogenetic networks

**DOI:** 10.64898/2026.09.11.750987

**Authors:** Michael R. May, Carl J. Rothfels

## Abstract

Much of the history of life is reticulate and better represented by phylogenetic networks than by strictly bifurcating trees. Understanding the processes that generated that history thus requires models of diversification (speciation and extinction) that incorporate reticulation. However, we currently lack tractable reticulate diversification models. Here we develop a simple birth-death-reticulation model that includes unidirectional and bidirectional gene flow, homoploid hybrid speciation, and allopolyploidization, and derive a practical probability density function for networks under this model. We demonstrate that the model can extract information about reticulation processes from known phylogenetic networks. We also explore the empirical utility of the model using an allopolyploid network of ferns of the family Cystopteridaceae, revealing evidence in favor of the controversial hypothesis that polyploids have lower diversification rates than their diploid relatives. While the model represents an advance in our ability to learn about the diversification of reticulate lineages, we also identify significant statistical, computational, and empirical challenges that face this nascent framework.

---

Reticulate evolutionary processes—for example, introgressive hybridization, hybrid speciation, and allopolyploidization—play a significant role in producing and shaping diversity in many groups. These events have significant implications for our understanding of relationships among taxa, requiring us to represent lines of descent as reticulate networks rather than bifurcating trees. In addition to complicating our view of phylogenetic relationships, these processes are also of direct scientific interest, raising important questions about the evolutionary dynamics of reticulation and of reticulate lineages.

Phylogenetic networks can be either implicit or explicit (Huson and Bryant 2006). Implicit phylogenetic networks depict conflict/discordance (*e*.*g*., `splitsTree`; Huson and Bryant 2006), but do not directly model the processes that produced the conflict. By contrast, explicit phylogenetic networks treat reticulations as specific historical events where contemporaneous lineages exchanged or combined genetic material. Methods for estimating explicit phylogenetic networks from sequence data are an active area of research and development. Differences in the genomic consequences of different types of reticulation events necessitate different approaches, but tools for estimating hybrid networks (*e*.*g*., Solís-Lemus and Ané 2016; Solís-Lemus et al. 2017; Wen et al. 2018; Zhang et al. 2018) and allopolyploid networks (*e*.*g*., Huson et al. 2012; Jones et al. 2013; Jones 2017; Yan et al. 2022) have begun to emerge.

Methods for estimating explicit phylogenetic networks have largely focused on modeling the details of genomic inheritance, gene genealogies, and sequence evolution: What reticulate history makes a given sequence dataset or a set of genealogies most probable? However, this treatment ignores the fact that different species histories will themselves have different probabilities given a process of diversification. For example, a reticulation event may imply the existence of an extinct ancestral lineage that is highly implausible for a given set of speciation and extinction rates. Phrased differently, the underlying species network is itself the product of evolution and different networks will have different probabilities, independent of any sequence data that evolves along them.

Birth-death processes are commonly used to model the processes of speciation and extinction that produce bifurcating phylogenetic trees. These birth-death models are applied as tools to study diversification, typically on a pre-estimated topology (Morlon 2014), and as priors in Bayesian phylogenetic analysis (Yang and Rannala 1997). Although birth-death models have been expanded to include reticulation events (Morin and Moret 2006; Woodhams et al. 2016; Justison and Heath 2024), these extensions primarily serve as simulation tools for understanding genetic or diversification patterns, rather than as a basis for explicit probability densities of phylogenetic networks. This impairs our ability to study the diversification processes that produce phylogenetic networks.

The lack of tractable birth-death-reticulation processes may also impair our ability to estimate phylogenetic networks. For example, Bayesian phylogenetic network inference methods do exist and require specifying a prior probability distribution for the network. However, in these cases, the prior is either an arbitrary, abiological probability distribution (Jones 2017), or makes convenient but unrealistic simplifying assumptions about the network-generating process (*e*.*g*., no extinction and complete species sampling, and merging of lineages rather than sharing of genes between lineages; Zhang et al. 2018). The impact of these priors on the inferred network is unclear.

In this paper, we develop a simple birth-death-reticulation (BDR) model that accommodates a variety of reticulation events, including introgressive hybridization, homoploid hybrid speciation, and allopolyploid speciation. Our goal is to establish a practical model that is suitable both for inferring diversification (and reticulation) parameters in a traditional two-step macroevolutionary study and as a prior distribution in a Bayesian phylogenetic network inference model. To that end, we first articulate a general but abstract view of the phylogenetic reticulation process, from the processes that forms the network down to the sequences. We then specify the stochastic BDR process and derive (approximate) probability density functions for networks under this general model. We validate the probability density function (confirming that both the theory and implementation are correct), and perform a simulation study to assess how well the parameters can be estimated from networks. Finally, we apply the model to an existing phylogenetic allopolyploid network of ferns from the Cystopteridaceae to evaluate the plausibility of competing allopolyploid ancestry scenarios and to understand whether allopolyploids might diversify at different rates than diploids.

## Reticulation processes from genes to species networks

Broadly speaking, we conceive of reticulation events as episodes of genetic exchange or combination between otherwise genetically independent species (lineages). These episodes may alter the genetic composition of one or both species, or may even produce new species that are reproductively isolated from their parents. In addition to having direct consequences on the genomes of the involved species, these events will also shape the relationships among species, reflected as species-level phylogenetic networks. Ultimately, our goal is to develop a BDR model that describes the distribution of species networks. However, because gene genealogies and species networks are inextricable, we begin by framing our BDR model in the context of a full reticulation model that explicitly connects gene-level and network-level processes.

We view this full reticulation model as an extension of the multispecies coalescent (MSC) model (Maddison 1997; Rannala and Yang 2003). Under the MSC, genetic lineages trace their histories back from the present through branches of the species tree, potentially coalescing with other genetic lineages in the same species (branch of the species tree). This conceptualization leads to a two-level hierarchical model, with one level (the coalescent level) governing the probabilities of gene trees (which record which genetic lineages coalesce and when) inside of the species tree, and the other level (the mutation level) governing the probabilities of the observed sequences evolving on the gene trees. In many applications, the MSC model considers just these two levels. However, the MSC can include an additional level that models the species tree itself. In fact, this third level is necessary if we are estimating the species tree in a Bayesian framework, where we are required to specify a prior on the species tree. This prior may be abiological (using arbitrary densities on branch lengths; Rannala and Yang 2003) or biological (for example a birth-death process; Heled and Drummond 2010). Using a birth-death process in this role constrains the species tree based on a biological interpretable model, and allows us to simultaneously estimate speciation and extinction rates as either a byproduct of tree inference or a focal inference itself.

By analogy, we imagine a full reticulation model with three levels (Fig. 1): a level describing the evolution of the species network, a level describing the gene-tree coalescence within the species network, and a level describing the evolution of sequences on the gene trees. For simplicity, we assume that sequence evolution on gene trees operates the same way(s) for a species network as for a species tree. However, modeling the coalescent and network-generating processes requires careful consideration of the specific reticulation events at work.

**Figure 1:**
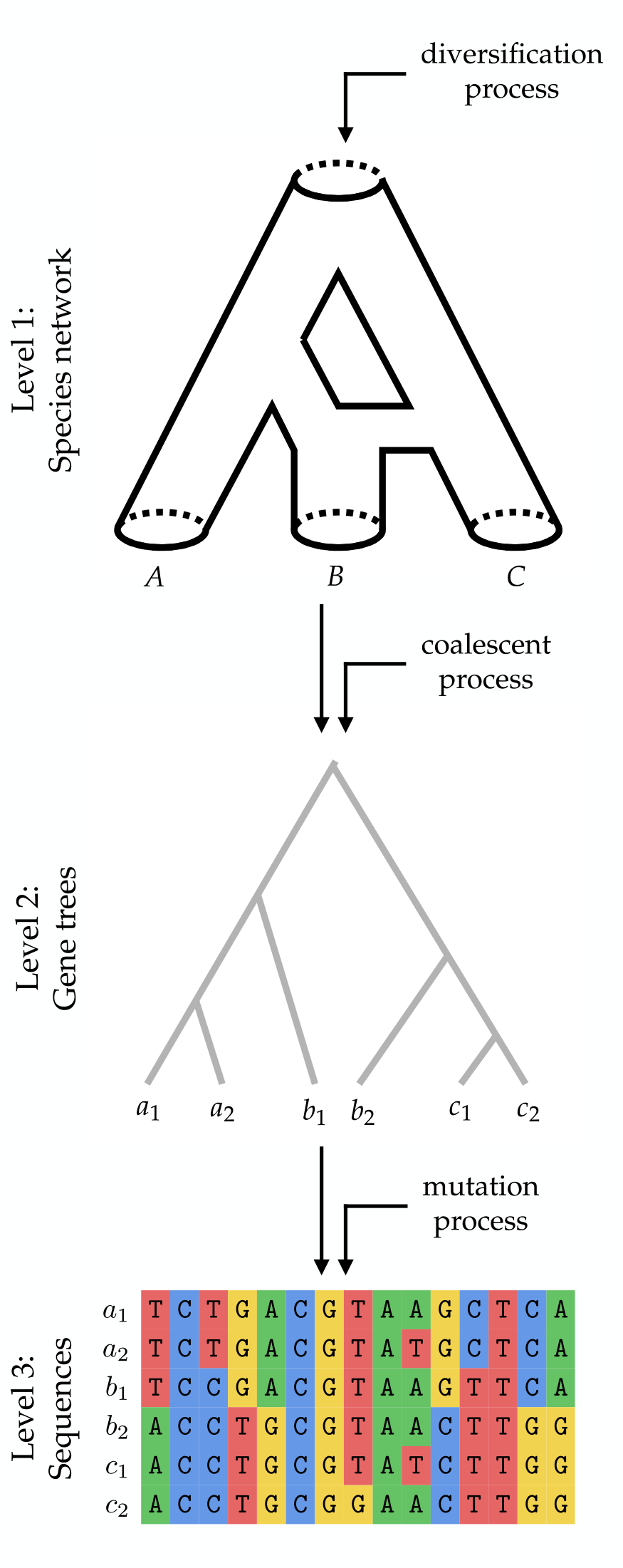
The three-level multispecies-network-coalescent model. We conceive of a full network process that operates at three levels. At the first level, a diversification process with parameters produces a species network, in this case for species *A, B*, and *C*. At the second level, gene trees evolve within the species network according to a coalescent process, producing samples *a*_1_, *a*_2_, *b*_1_, *b*_2_, *c*_1_ and *c*_2_. At the third level, sequence evolve along the gene trees according to a mutation process, producing a sequence alignment for the samples.

### Types of reticulation and their genetic consequences

We consider three broad types of reticulation: “introgressive hybridization”, where two species exchange genetic material without producing a new species; “homoploid hybrid speciation”, where genetic material from two species merges to produce a new hybrid species with the same number of genomes as each of its parents; and “allopolyploid speciation”, where two species produce a new species with the full genomes of each parent. Within introgressive hybridization, we further distinguish between unidirectional and bidirectional hybridization.

At the genetic level, hybridization causes introgression, producing lineages with mixed genetic ancestry. In unidirectional hybridization, one lineage donates genes to another, producing a single hybrid lineage and leaving the donor unchanged (Fig. 2, first row, first column). In bidirectional hybridization, each lineage donates genes to the other lineage, producing two hybrid lineages (Fig. 2, first row, second column). In homoploid hybrid speciation, both lineages (the parents) donate genes to a new hybrid lineage; the parents themselves are unchanged (Fig. 2, first row, third column). Multi-species network coalescent models have been developed for these hybridization events (Kubatko 2009; Yu et al. 2014; Flouri et al. 2020). These models are variously referred to as multispecies network coalescent models (MSNC, which we consider ambiguous because not all network models are hybrid models; Wen et al. 2016), or multispecies coalescent with introgression models (MSCi; Flouri et al. 2020).

**Figure 2:**
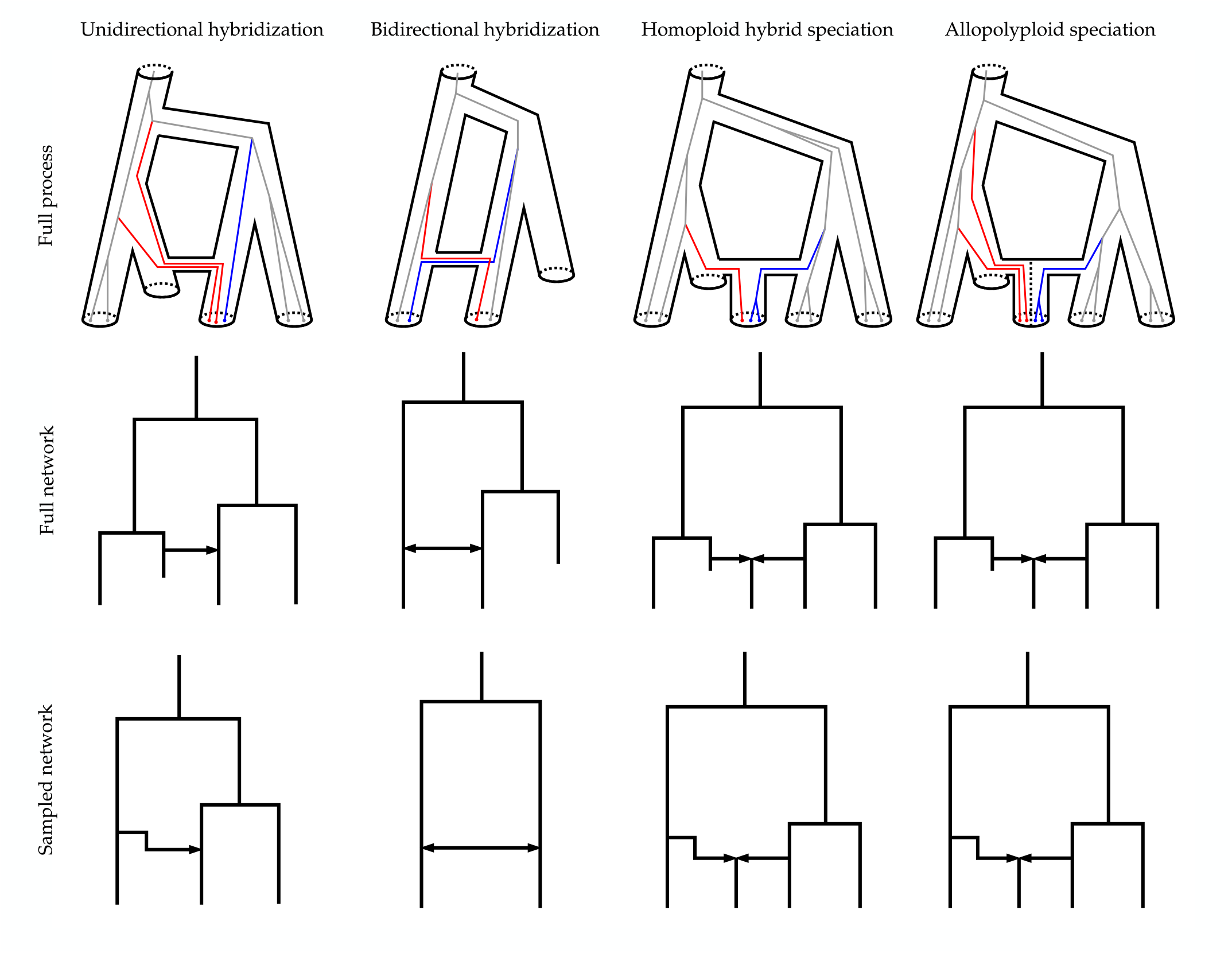
Four types of reticulation events, and their consequences on gene genealogies and network shapes. First column: unidirectional hybridization, where one species donates genes to another. Second column: bidirectional hybridization, where two lineages exchange genes. Third column: homoploid hybrid speciation, where two parents produce a new hybrid with mixed ancestry. Fourth column: allopolypoid speciation, where two parents produce a new species with both parental genomes (the hybrid species branch is subdivided into the two respective subgenomes by a dashed line). First row: an example full process showing coalescence within a species network that includes the focal reticulation type as well as speciation and extinction. In the hybrid networks, introgressed genes are depicted in blue and red. In the allopolyploid network, gene copies are colored by the subgenome of origin. Reticulation events are assumed to be instananeous, but they are depicted with some vertical extent for clarity. Second row: the species networks including all species, with arrows depicting the direction of reticulation. Third row: the species networks including only surviving species (the “sampled” or “reconstructed” network).

In each case, genetic lineages tracing their ancestry back to a hybridization event are inherited from one parent (with probability typically denoted *γ*) or the other parent (with probability 1 − *γ*).

By contrast, the genetic consequence of allopolyplodization is whole-genome duplication: the new allopolyploid species has multiple subgenomes (at least soon after the event), one set inherited from each parent. Existing multispecies allopolyploid network models (Jones et al. 2013; Jones 2017) treat the subgenomes of allopolyploids as independent, so that genetic lineages only coalesce with other genetic lineages in the same subgenome (homeologous recombination is not allowed). When genetic lineages in one subgenome trace back to an allopolyploidization event, they are necessarily inherited from the corresponding parent (Fig. 2, first row, fourth column).

### Reticulation, extinction, and species networks

In addition to their direct genetic consequences, reticulation events affect the shapes of species networks by adding reticulation edges and by operating as a potential source of new species. The patterns of reticulation in a species network are also influenced by extinction and incomplete sampling.

In principle, reticulation events should always occur between contemporaneous lineages: if we knew the complete history of the network including all extinct species, all reticulation edges would be horizontal (Fig. 2, second row). However, if one of the parents leaves no extant sampled descendants, reticulation edges have kinks or angles (Fig. 2, third row).

Extinction can also confuse both gene-level and network-level signatures of reticulation. For example, unidirectional and bidirectional hybridization will be indistinguishable if one of the parents goes extinct, taking with it the record of genes that might have traced their ancestry back to the surviving parent. Likewise, homoploid hybrid speciation with both parents going extinct can look like other unidirectional or bidirectional hybridization with one extinct parent.

A coherent, mechanistic network-generating process suitable for the top level of our full reticulation model must be able to explain both horizontal and angled edges in a coherent way; in particular, the presence of angled reticulation edges implies the extinction of (or failure to sample) one or both of the parents of the reticulate lineage (so-called “ghost lineages”). The model must also be able to account for the different types of reticulation events that could lead to the same genetic signals because of extinction of parental lineages. In the section that follows, we derive such a model in the form of an explicit stochastic BDR model that describes the formation of phylogenetic networks by speciation, extinction, and reticulation events.

## A stochastic birth-death-reticulation process

We develop a stochastic birth-death-reticulation process that produces species networks. Our stochastic birth-death-reticulation model shares many features of the models described in Justison and Heath (2024), but is simplified for tractability and explicitly distinguishes between hybridization and allopolyploidization events. We refer to the number of species alive at time *t* as *N*(*t*). The process begins at time *t* = *T* with one lineage, *N*(*T*) = 1, or at a speciation event producing two lineages (*i*.*e*., the root, *N*(*t*) = 2), and evolves forward in time by speciation, extinction, and reticulation events.

For simplicity, we assume that speciation and extinction events occur at a constant rate across species and over time (similar to a standard constant-rate birth-death process). As with a standard BD process, speciation events occur at rate *λ* per species, splitting an existing species into two daughter species. Likewise, extinction events occur at rate *µ* per species, terminating the affected species. Because these events occur independently for each species, the total rate of speciation (and extinction) at time *t* increases linearly with *N*(*t*).

In contrast, we assume that each contemporaneous pair of species represents an independent opportunity for reticulation to occur, so that the rate of reticulation at time *t* is proportional to 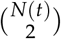 rather than *N*(*t*) (Justison and Heath 2024); the possibility for reticulation increase much more quickly with the number of species than does speciation or extinction. Reticulation events either keep the number of species the same (“lineage-neutral” events, *sensu* Justison and Heath 2024) or increase the number of species (“lineage-generative” events). We do not consider reticulation events that decrease the number of species (“lineage degenerative” events). We model four distinct types of reticulation events: 1) unidirectional hybridization; 2) bidirectional hybridization; 3) homoploid hybrid speciation, and; 4) allopolyploid speciation.

Unidirectional hybridization is a lineage-neutral event that occurs at rate *η* per species pair, and involves one species donating genes to another (Fig. 3, event marked U). During such an event, each possible direction is equally likely, so a unidirectional hybridization event from species *x* to species *y* occurs at rate *η*/2. Bidirectional hybridization is also a lineage-neutral event, occuring at rate *ζ* per species pair (Fig. 3, event marked B). During a bidirectional hybridization, each species donates genes to the other, creating two hybrid species. Homoploid hybrid speciation is a lineage-generative event, occuring at rate *ν* per species pair (Fig. 3, event marked H). During such an event, a new hybrid species is produced, and both parents remain the same. Allopolyploidization is a lineage-generative event that occurs at rate *Ψ* (Fig. 3, event marked P). Like homoploid hybrid speciation, a new species is produced and both parents remain the same. However, the new species inherits the genomes of each parent. (Note that the network-generating process is agnostic about the ploidy of the parents, so the new lineage may be tetraploid, hexaploid, octoploid, etc., depending on the ploidy of the parents.)

**Figure 3:**
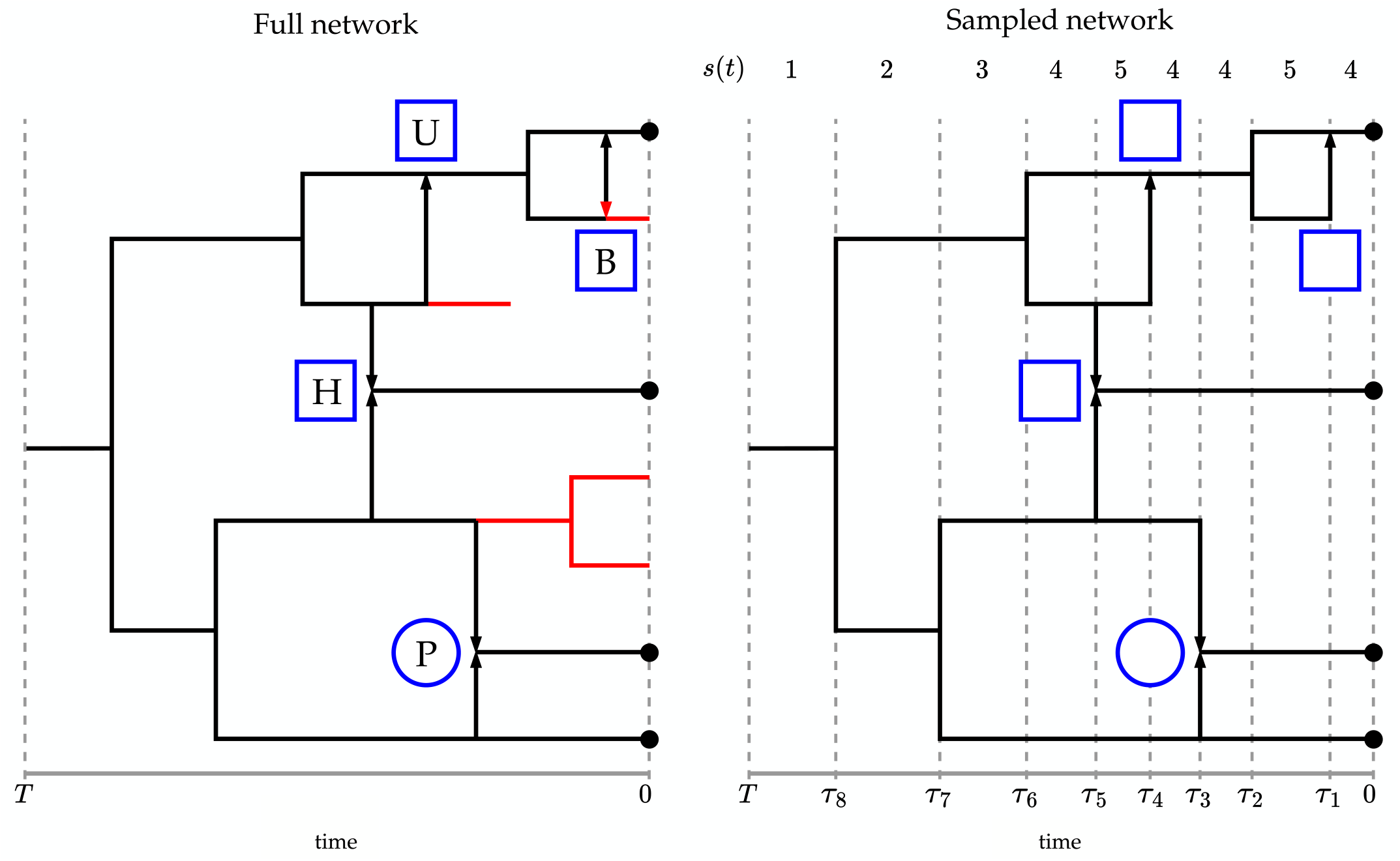
An example phylogenetic network produced by the birth-death-reticulation process. Left) A full phylogenetic network produced by the birth-death-reticulation process, in this case beginning with a single lineage at time *T* in the past. Black lines represent sampled lineages, while red lines represent extinct or unsampled lineages. Reticulation events are indicated by blue shapes; hybridization events are indicated by squares and allopolyploidization events are indicated by circles. A unidirectional hybridization event is marked U; a bidirectional hybridization event is marked B; a homoploid hybrid speciation event is marked H; and an allopolyploid speciation event is marked P. Otherwise, nodes in the tree are produced by speciation events, and extinction events terminate lineages before they can reach the present. Species sampled at the present are marked with black dots. Arrows indicate the direction of the reticulation events. Note that reticulation events pointing into extinct/unsampled lineages are themselves red, since their signature is erased. Right) The sampled phylogenetic network, produced by pruning extinct and unsampled lineages from the full network. The times of each node in the network are indicated by *τ*_*i*_, and the number of sampled lineages, *s*, alive during each interval is marked above the network. Note that reticulation events in the sampled network retain information about whether they involved hybridization or allopolyploidization, because we assume the resulting lineages contain genetic information about the type of event that produced them.

The process continues evolving according to these events until all species die out, or until the present, *t* = 0, at which point each species is sampled with probability *ρ*. We then prune out all the extinct and unsampled lineages, as well as reticulation arrows (indicating their direction) if the recipient lineage went extinct or unsampled; for example, a bidirectional hybridization event where one of the descendant lineages went extinct would not retain the signature of hybridization leading into the extinct lineage. We call the network with all lineages (even those that are extinct or unsampled) the full network, and the network with extinct and unsampled lineages pruned out the sampled (or reconstructed) network, denoted ℵ (Fig. 3, left and right, respectively). The sampled network retains information about the genetic category of each reticulation event, *i*.*e*., either hybridization or polyploidization, because the descendant lineages will retain genetic information about the nature of the event from which they arose (indicated by squares and circles in Fig. 3).

The birth-death-reticulation process we introduce, with all four type of reticulation events, is meant to be general, covering most of the types of reticulation that are thought to occur in nature. However, in many cases it may make sense to use only one or a few of the reticulation events. We will sometimes refer to the special case-models with just one parameter according to that event, *e*.*g*., the birth-death-allopolyploidization model includes *Ψ >* 0, with the other reticulation rates set to zero.

### The probability density of a network

Computing the probability of a sampled network under this model requires overcoming some special challenges. First, because lineages interact during reticulation events, we cannot make the assumption common to standard birth-death models that each lineage evolves independently. Second, the probability of the sampled network depends on the probability that—apart from the reticulation events recorded in the network—sampled lineages *did not* draw ancestry from lineages that ultimately went extinct (or survived but were unsampled); this probability depends on how many unobserved lineages there are at any given time in the past, which is unknown.

These issues lead us to model the joint evolution of the number of sampled and unobserved lineages. (An unobserved lineage is one that went extinct before the present or survived but was not sampled.) We denote the number of sampled and unobserved lineages at time *t* as *s*(*t*) and *u*(*t*), respectively, and the joint state as {*s, u*} (*t*). As we describe later, the probability of the network relies on the probability that {*s, u*} (*t*_0_) at time *t*_0_ evolved from state {*s, u*} (*t*_1_) at some time *t*_1_ in the past. The number of sampled lineages *s* is naturally constrained by the number observed in the network at time *t*. However, the number of unobserved lineages *u* can range from zero to infinity, which means that the state space of {*s, u*} is infinitely large, complicating practical calculations.

We derive a practical approximation to the network probability density by truncating the maximum number of unobserved lineages to a fixed constant *u*_max_; events that would increase the number of unobserved lineages beyond this number are allowed to occur, but flow into an implicit absorbing state that represents “more than the maximum number of represented unobserved lineages”. As *u*_max_→∞, this approximate probability density converges to the true, untruncated probability density. This approach is conceptually similar to the solutions adopted for stochastic models of chromosome number evolution (Mayrose et al. 2010; Freyman and Höhna 2018), and for density-dependent diversification models (Etienne et al. 2012).

Our probability density calculation begins at the present (*t* = 0) and works backward (*t >* 0) toward time *T* (either the root of the network or the origin of the stem of the network, depending on the network in question). We denote the probability density of the network downstream from time *t* (toward the present) as *p*_*u*_(*t*). We denote the vector of probabilities for all values of *u* as 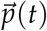. Computing the probability density of the entire network involves four types of operations: (1) defining the time intervals between node events; (2) specifying the probability at the present, *p*_*u*_ (*i*.*e*., specifying the initial conditions); (3) computing how *p*_*u*_(*t*) changes as we move backward, between node/reticulation events in the sampled network, and; (4) computing how *p*_*u*_(*t*) changes at speciation and reticulation events (which we refer to generically as “node” events), accommodating all possible node events that could have produced the observed configuration of branches at time *t*. These operations consider all possible values of *u* (up to *u*_max_), effectively integrating over all possible histories of unobserved lineages that could have produced the sampled network. Repeating these operations from the tips to the root (or stem), we end up with the quantity *p*_1_(*T*), which is the probability density of the entire network. We now describe how we perform each of these operations.

#### Time intervals

We denote the age of the *i*^th^ node event in the network as *τ*_*i*_. We divide the time from 0 to *T* into a finite set of time intervals based on the ages of the node events. For a network with *k* node events, there will be *k* intervals if the network begins at the root, and *k* + 1 intervals if it begins at the stem. We order the intervals from the present to the past. The first interval spans (*τ*_1_, 0), while subsequent intervals span (*τ*_*i*_, *τ*_*i*−1_) for interval *i*. This scheme is depicted in Fig. 3 (right). We denote the number of sampled lineages in the *i*^th^ interval as *s*_*i*_. Note that we treat these intervals as closed (not including the exact time point on either end of the interval) because calculations that take place over the intervals (transition probabilities) are different from calculations that take place exactly at the breakpoints between intervals (node-event densities).

#### Initial conditions

We assume for simplicity that extant species are sampled uniformly and independently at random with probability *ρ*. Because the sampled network is fixed, we know there are exactly *s*_1_ sampled species at the present, so the initial probability is just the probability that those *s*_1_ lineages were each sampled times the probability that each of the *u* unsampled lineages were not. Accordingly, we set

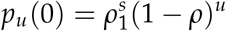

for each value of *u*.

#### Transition probabilities between nodes

Our approach for computing how 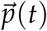 changes over time intervals closely follows Maddison et al. (2007). Specifically, we consider the finite set of events that could happen over a small interval of time from *t* to *t* + Δ(backward in time) to produce the sampled network descendant from time *t*. We then use this finite set of events (and their associated probabilities) to derive a set of ordinary differential equations (ODEs) that describe how each *p*_*u*_(*t*) changes over time:

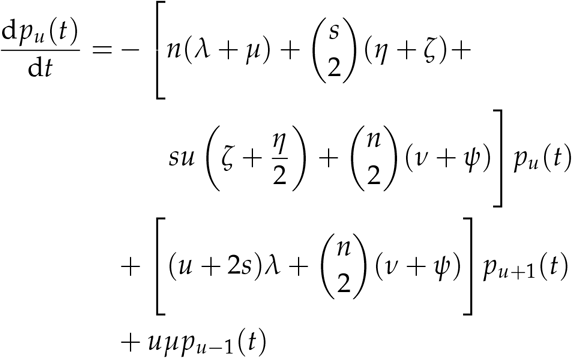

with *n* = *s* + *u*, and *s* set to *s*_*i*_ for the corresponding interval. Note that this equation is for a typical value of *u*, with some modifications necessary near boundaries. (See Supplemental Material for derivation.)

To compute the probability 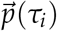 given the probability 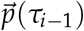, we solve this set of ODEs from *τ*_*i*−1_ to *τ*_*i*_. In contrast to the ODEs derived for BiSSE (Maddison et al. 2007) and related state-dependent models, these ODEs can be solved using matrix exponentation:

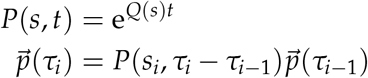

where *Q*(*s*) is a sparse matrix of coefficients defined for a given number of sampled lineages *s* (described in Supplemental Material).

#### Transition densities at nodes

We now turn to the effect of observed nodes on the probability density of the sampled network. In bifurcating birth-death models, nodes only represent splits, and generally involve multiplying the probability density by the speciation rate. In contrast, nodes in a reticulate network can also involve the merging of ancestral lineages, with different possible configurations depending on the reticulation events involved and the subsequent history of extinction and sampling. These additional possibilities necessitate some additional bookkeeping when we encounter reticulate nodes in the sampled network.

We enumerate all the node-generating events that could occur to produce observed nodes in the sampled network. Each node event is characterized by the stochastic event involved (either speciation or a particular reticulation event), and the history of extinction and sampling that occurs subsequent (forward in time) to the event. The events, their descriptions, and their corresponding probability densities are depicted in Fig. 4.

**Figure 4:**
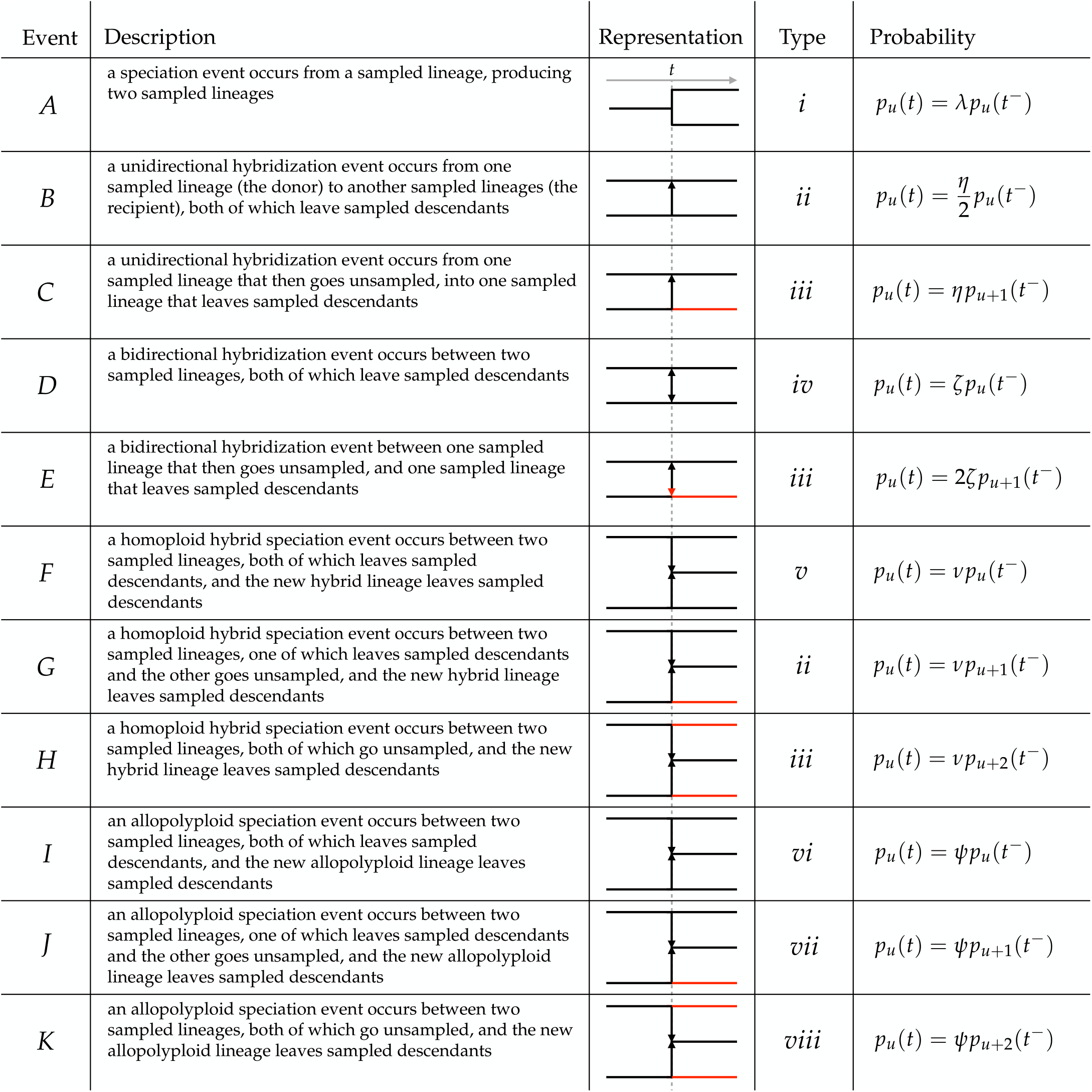
Node events in sampled networks. Node events (in rows, lettered *A* through *K*) are defined as a combination of a particular reticulation event and a particular history of extinction (or incomplete sampling) that could produce an observed node in the sampled network. The representation of each event depicts the reticulation event, with extinct (or unsampled) lineages depicted in red, and sampled lineages depicted in black, and time decreasing to the right (toward the present). Reticulation events are depicted as horizontal arrows pointing in the direction of the reticulation (the recipient or new lineage); note that extinction can also remove information about the directionality, resulting in a red arrowhead (event E). There are eight types of node events, numbered *i* through *viii*. The type of a node event is determined by the pattern of the sampled lineages that are produced; a specific node event cannot be distinguished from other node events of the same type. In particular, node events *B* and *G* are both of type *ii*, while events *C, E*, and *H* are all of type *iii*. The probabilities of the downstream network immediately after the event at time *t, p*_*u*_ (*t*), depends on the rate of the corresponding reticulation event, and the appropriate probability at the time point immediately before the event, which we denote time *t*^−^.

Some node events generate the same pattern of sampled lineages, leading to ambiguous patterns in the sampled network. When we encounter a node in the sampled network, we must therefore consider all possible node events that are consistent with that configuration of sampled lineages. To do this, we categorize the node events into distinct types, which we enumerate *i* through *viii*. Node events of the same type cannot be distinguished, but different types of node events are distinguishable (Fig. 4). Most node event types contain just one node event, but there are two types (*ii* and *iii*) that contain multiple node events. Type *ii* is comprised of a unidirectional hybridization event where both donors are sampled (a type *B* event) and a homoploid hybrid speciation event where one of the parent lineages goes unobserved (a type *G* event). In either case, there are two sampled lineages prior to the event and two sampled lineages after the event. Type *iii* comprises a unidirectional hybridization event where the donor goes unobserved (a type *C* event), a bidirectional hybridization event where one of the lineages goes unobserved (a type *E* event), and a homoploid hybrid speciation event where both parents go unobserved (a type *H* event). In these cases, there are two sampled lineages prior to the event and one sampled lineage after the event.

When we encounter a node of a particular type as we move back through the sampled network, we update the probabilities *p*_*u*_(*t*) according to the probability rules in Fig. 4. For example, when we encounter a type *i* node (a speciation event), we set *p*_*u*_(*t*) = *λp*_*u*_(*t*^−^), where *t* is the time of the node event and *t*^−^ is the moment immediately before the speciation event. For ambiguous node types, we have to sum the probabilities of events with the same index of *u*. For example, for a type *ii* node, we set

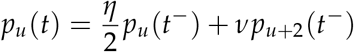

for all values of *u*.

For each type of node event, we can define a transition probability matrix that encodes the effect of that type of event on the entire probability vector 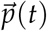. Specifically, 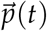can be computed as:

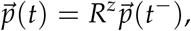

where *R*^*z*^ is a transition probability matrix for type *z*∈ {*i, ii*, … *viii*}.

We derive the probability densities of each type of node event, and the corresponding transition probability matrices, in the Supplemental Material.

#### Probability density of the sampled network

The probability density of the sampled network depends on whether it began with a stem (a single lineage) or the root (two lineages). If the process began with the root, then we first compute 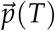 according to a speciation event, as described above. If the process began with the stem, we do not change 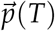, as there was no event at time *T*. In either case, the probability density of the network is

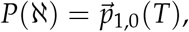

because the process began with one sampled lineage (that immediately split into two sampled lineages in the root case) and no unsampled lineages, by assumption.

We note that this probability density is *unconditional*, specifically we do not condition on either survival (as is typical for birth-death models) or, perhaps more importantly, on there being at least one reticulation event in the sampled network. We currently lack a straightforward approach for calculating the marginal probabilty of either of these cases, which prohibits us from presenting the conditional probability.

#### Matrix representation

The probability density of the network can be represented compactly as a set of matrix operations acting on the initial probability 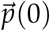. Recalling that time intervals are sorted from youngest to oldest, the probability density for a network starting at the root is

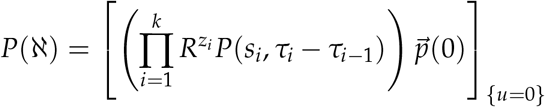

where 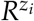is the event matrix of the *i*^th^ event and *k* is the number of node events and intervals. (The product is ordered from right to left, so that factors with smaller *i* operate first.) Starting with the stem, the probability density is

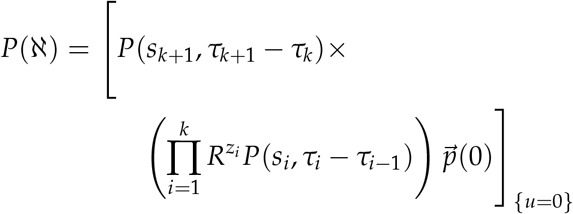

## Implementation

### C++ library

We implemented the probability density calculation for our BDR model in a `C++` library we call `Diversinet`. The library accepts a network (in a modestly extended Extended Newick Format, described below) as well as model parameters and other numerical settings (*e*.*g*., *u*_max_), and returns the probability density of the network given those parameters.

The runtime is dominated by matrix operations, particularly the matrix exponential, because non-trivial values of *u*_max_ lead to large matrices. The library uses `Eigen` for sparse matrix types and operations (Guennebaud et al. 2010), as well as `Boost` ODE solvers to compute the matrix exponential (*e*.*g*., Euler, adaptive Fehlberg and Dormand-Prince Runge-Kutta schemes; Fehlberg 1969; Dormand and Prince 1980; Boost.org contributors 2026). However, we found that these ODE solvers were unstable for large values of *u*_max_, and so implemented a more stable matrix exponential solution using uniformization (Grassmann 1977) with adaptive truncation (Reibman and Trivedi 1988). We describe this uniformization approach in the Supplemental Material.

### Network input format

Our library accepts input networks in an extended version of the Extended Newick Format (Cardona et al. 2008). In the Extended Newick Format, hybridization events are represented by duplicate node labels annotated as either `LGT#` or `H#`, where # is an index. We extended this format to include the annotation `P#` to represent polyploid reticulation events. The library interprets reticulation events annotated with either LGT or H as hybridization events and those annotated with P as allopolyploidization events.

### Julia package

We implemented a lightweight Julia interface to our `C++` library called `Diversinet.jl`. The package exposes the probability density function (pdf) for a network, allowing users to employ the pdf as a likelihood function in existing inference machinery available in `Julia` for maximum likelihood inference (*e*.*g*., `Optim`, Mogensen and Riseth 2018) or Bayesian inference (*e*.*g*., `AdaptiveMCMC`, Vihola 2022, 2026). We use `AdaptiveMCMC` for the analyses that follow (except where noted otherwise).

## Validation

We performed validation tests to ensure that both our theory and implementation of the network probability density were correct. Under the BDR model, the number of tips, *n*, and reticulation events, *r*, are both random variables with joint probability mass function *P*(*n, r*| *θ, T*), where *θ* are the parameters of the process, and *T* is the age of the process. This probability mass can be computed as a sum over network topologies with *n* tips and *r* reticulation events, and a multidimensional integral over the ages of nodes in the network:

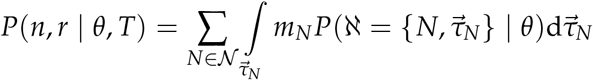

where *N* is the network topology, *N* is the set of all network topologies with *n* tips and *r* reticulation events, 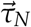is the vector of ages of the events for topology *N*, and *m*_*N*_ is the number of orientations of topology *N*; a particular network is defined by the network topology *N* and the ages 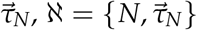. This equation allows us to validate our probability density function, *P*(ℵ|*θ*), by brute force enumeration of topologies and integration over node ages, and comparing against the joint probability mass of *n* and *r*.

We validated five special-case scenarios that together cover every piece of our implementation code: 1) a unidirectional hybridization model; 2) a bidirectional hybridization model; 3) a homoploid hybrid speciation model; 4) a model with all non-allopolyploid hybridization events, and; 5) an allopolyploid speciation model. We chose these scenarios to cover each of the possible event types, and to cover every part of the likelihood code. For each scenario, the included parameter(s) were set to *θ >* 0 and the remaining parameters were excluded (set to zero). In all cases, we assumed networks began with the stem lineage at time *T* = 1, that the relative extinction rate was *€* = *µ*/*λ* = 0.1, and the sampling fraction was *ρ* = 0.2; we then tuned the speciation rate *λ* and the relevant parameter *θ* such that the expected number of tips was relatively small (two for scenarios 1 and 2 and three for scenarios 3 through 5). In scenarios 1 and 2, we validated against the probability mass of *n* = 2, *r* = 1; for the remaining scenarios, we validated against the probability mass of *n* = 3, *r* = 1.

For each scenario, we enumerated all network topologies with the target number of *n* and *r*. We then computed the target probability mass by summing our network probability density over each network topology, and for each network topology, integrating over node ages using multidimensional numerical integration implemented in the `Julia` package `cubature` (Johnson 2013). We refer to this quantity as the marginalized network density. Critically, these scenarios together involve every piece of mathematical machinery implemented in `Diversinet`.

An analytical formula for the joint probability mass *P*(*n, r* |*θ, T*) is unknown. For each scenario, we used Monte Carlo simulation to approximate the joint probability mass against which we compared the marginalized network density; *e*.*g*., for scenario 1, we simulated many networks under the same model parameters and computed the fraction of simulations with *n* = 2 and *r* = 1. Further details of the network density marginalization and Monte Carlo procedures are described in the Supplemental Material.

We validated over a range of values of the relevant *θ* spanning the tuned value. (For scenario 4, which has three parameters (*η, ζ, ν*), we fixed the ratios of these parameters and validated over a range of sums, *θ* = *η* + *ζ* + *ν*.) We also performed the validation over a set of truncation values, *u*_max_ = {4, 8, 16, 32, 64}. We compared against target probabilities approximated from 10 million Monte Carlo replicates. Figure 5 shows that the marginalized network density approaches the Monte Carlo approximation as *u*_max_ grows, demonstrating that the theory and implementation of our density are correct.

**Figure 5:**
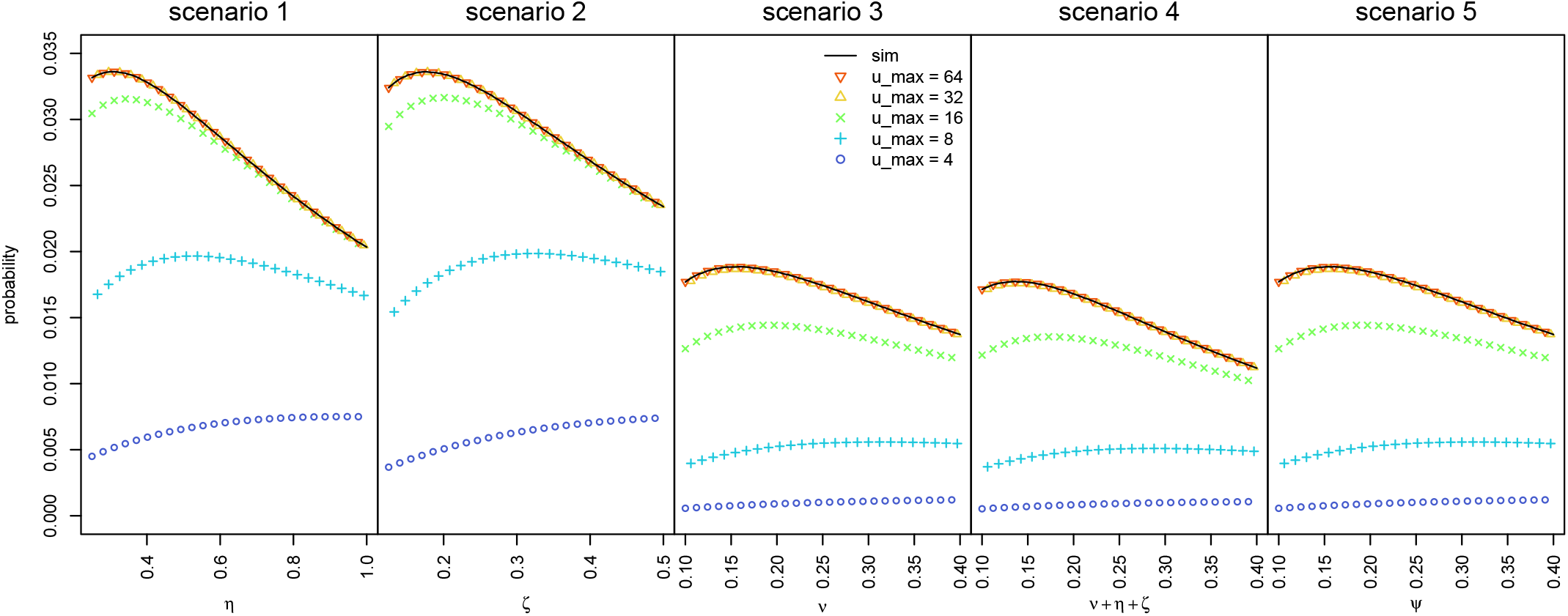
Validating the probability density of the network. We computed the probability of *n* tips and *r* reticulation events by averaging our network density over all topologies and sets of node ages with that *n* and *r* (the marginalized network density). We repeated this calculation for five scenarios (panels, described in the text), over a range of parameter values (x-axis), and for different values of *u*_max_ (colored symbols). We then compared these values against the frequencies of the target *n* and *r* approximated from 10 million Monte Carlo simulations under the same model and parameter values (black lines). As *u*_max_ increases, the marginalized network density approaches the Monte Carlo estimate, demonstrating that the theory and implementation of our network density are correct.

## Simulation Study

We assessed our ability to estimate the parameters of the BDR model using simulated phylogenies. Before describing the simulations in detail, we wish to note some important caveats.

First, because we lack a solid empirical understanding of reasonable reticulation rates, our choice of simulating rates is somewhat arbitrary. Further, the positive density-dependence of the reticulation rate means that even small rates can cause individual simulations to explode in size—producing networks with millions of reticulation events—so the range of values we explored is relatively small. Nonetheless, our simulations cover a broad range in the number of reticulation events (from none to dozens) that we hope includes most reasonable empirical networks. We summarize the distribution of the number of events as a function of simulation parameters in the Supplemental Material, and note that the results in Figure 6 (for the allopolyploid model) is color coded by the number of reticulation events, to give a sense of the range covered.

**Figure 6:**
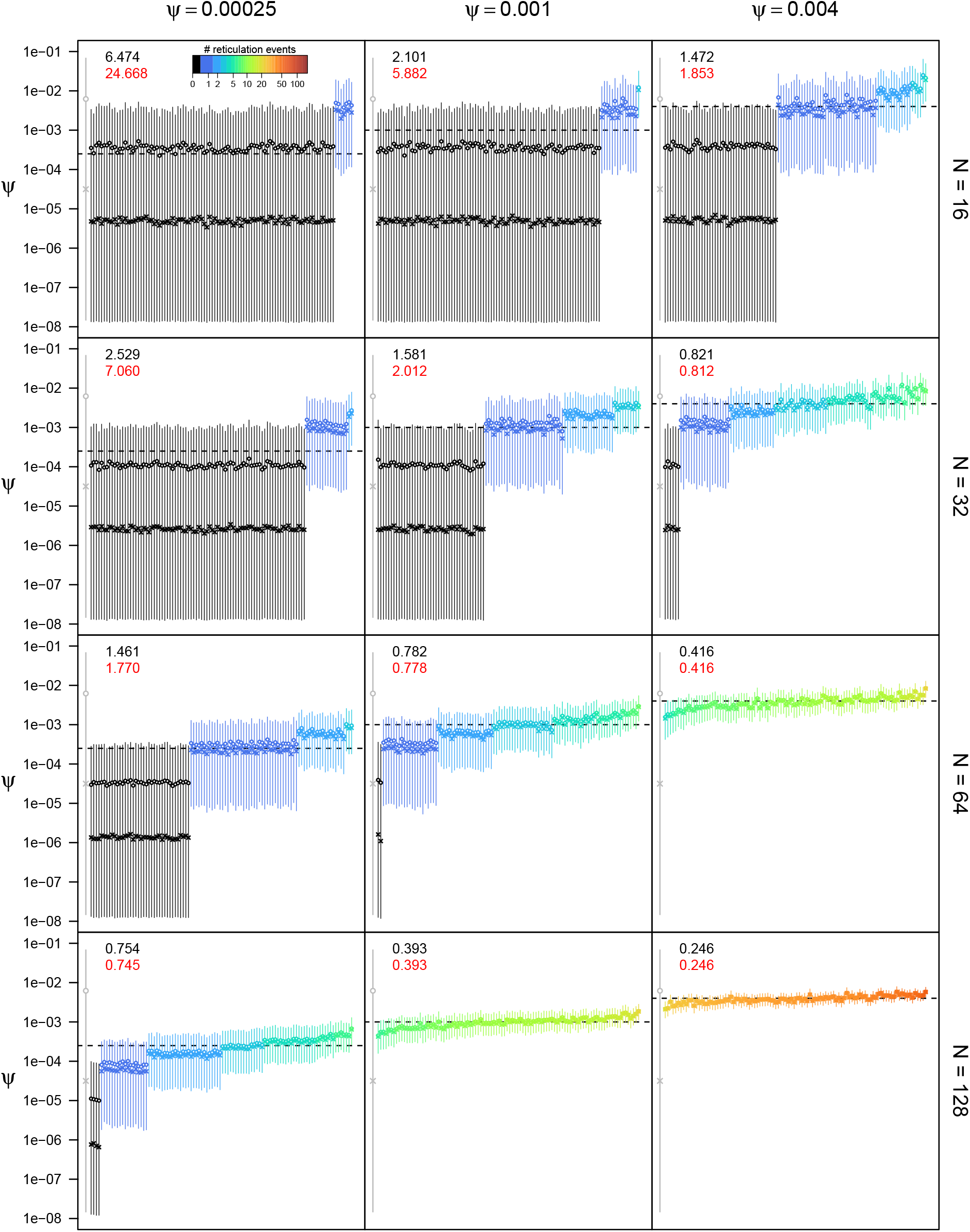
Posterior estimates of the allopolyploidization rate *Ψ* for simulated networks for *€* = 0.3, *ρ* = 0.5, and *u*_max_ = 4*N*. Rows correspond to different network sizes, *N*, and columns correspond to different true rate parameters (also depicted as a dashed horizontal line). Each cell shows the posterior distribution of 100 replicate networks; the vertical bar corresponds to the 95% credible interval, the circle to the posterior mean, and the cross to the posterior median. The grey bar at the far left of each panel is the prior distribution on *Ψ*. Bars are colored by the number of reticulation events (colored legend in first panel). The black number in the top left is the root-mean-squared error (RMSE, described in the text) averaged over all datasets; the red number is the RMSE averaged over datasets with at least one reticulation event.

Second, our simulations include phylogenies that, by random chance, have no reticulation events, *i*.*e*., they are unconditional on reticulation. We retain these phylogenies because our probability density function is likewise unconditional on reticulation, owing to the difficulties of the necessary calculations. Our results therefore reflect how the model should behave were it systematically applied to any phylogeny, whether it is a tree or a network. However, researchers are more likely to use a BDR model on a phylogeny with reticulation events, which in practice may upwardly bias estimates of reticulation rates. We address the potential consequences of this ascertainment bias in the simulation results.

Third, our simulations assume that the true phylogeny is known. Of course, all phylogenies are estimates rather than direct observations, and are therefore subject to random error, systematic error, and uncertainty. While many issues that attend treating phylogenies as direct observations are common to birth-death analysis, our model has the added problem that estimated phylogenetic networks may under- or over-count the number of reticulation events. Because they will necessarily depend on the biology, data, and methods, the risks and consequences of missing or erroneous reticulation events are difficult for us to anticipate. We return to this issue in the Discussion.

### Simulation settings

We performed simulations under three special-case models: 1) a model of unidirectional hybridization; 2) a model of bidirectional hybridization, and; 3) a model of allopolyploid speciation. For each simulation scenario, we simulated with a positive value of the focal parameter, which we denote *θ*, and set the remaining reticulation rates to zero. We did not simulate under a model of homoploid hybrid speciation because it is essentially identical to the allopolyploid speciation model, except that the events are labeled as homoploid hybrid speciation events rather than allopolyploid speciation events (see Fig. 2); therefore, our ability to estimate the homoploid hybrid speciation rate *ν* should be identical to our ability to estimate the allopolyploid speciation rate *Ψ*.

We explored the influence of both parameter values and network size on our ability to estimate parameters. For each simulation scenario, we simulated networks with a fixed speciation rate *λ* = 1, and varied the other parameters as follows:

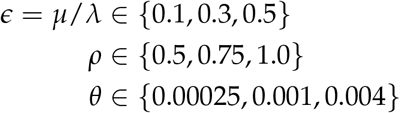

where *θ* is the relevant reticulation parameter.

We varied the network size (number of tips *N*) over the values *N* ∈{16, 32, 64, 128}. We used rejection sampling to simulate trees of a given size *N, i*.*e*., we repeated a given simulation replicate until the network was of the target size. To ensure that the resulting networks were not unusually large or small for a given parameter combination, we used Monte Carlo simulation to tune the root time *T* for that parameter combination so that the expected number of tips was 60. We then used the same value of *T* (for a given parameter combination) for all values of *N*, so that networks spanned the expected number and were not sampled from an usual part of the network outcome space.

We simulated 100 replicate phylogenies for each combination of *€, ρ, θ*, and *N*. This resulted in a total of 3 ×3 ×3 ×4 ×100 = 10, 800 simulated phylogenies for each of the three simulation scenarios. We provide more details of our simulation procedure and the distribution of the number of reticulation events in the Supplemental Material.

### Simulation analyses

We performed Bayesian inference on each simulated phylogeny, estimating the joint posterior distribution of the parameters *λ, €*, and *θ*, with the prior distributions

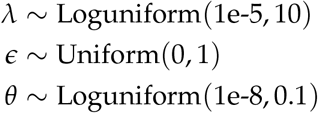

and *ρ* set to the true value. We repeated each analysis for three different values of *u*_max_. Specifically, we set *u*_max_ to *u*_max_ ∈{*N*, 2*N*, 4*N*} where *N* is the number of tips in the analysis phylogeny.

We estimated the posterior distribution for each analysis with the the `Julia` package `AdaptiveMCMC` (Vihola 2026). We ran each chain for 50, 000 iterations, which in the vast majority of cases resulted in ESS values *>*1000 for all model parameters. The rare failures (about 20 in total) were the result of stuck chains and/or badly tuned MCMC proposals; we simply reran them until they worked. In total, we performed 97, 200 MCMC analyses, plus the small number that had to be rerun.

### Simulation results

We present results for the allopolyploid speciation model for paremeter combination *€* = 0.3, *ρ* = 0.5, and *u*_max_ = 4*N* (Fig. 6). Results for other simulation scenarios, and other *u*_max_ and parameter values are qualitatively very similar (see Supplemental Material). Posterior distributions of the allopolyploidization rate *Ψ* (and likewise for the other reticulation parameters) are almost entirely determined by the number of reticulation events in the phylogeny (posteriors are color coded by the number of events in Fig. 6). With zero reticulation events (black bars), the posterior estimates are low but differ from the prior (grey bars), indicating that there is evidence in these cases that the reticulation rate is not very high. For a given network size, the posterior estimates of *Ψ* are essentially identical across true values of *Ψ* for networks with a given number of reticulation events (*e*.*g*., compare dark blue bars for a given row across columns). This is quite sensible, because the information about the reticulation rate primarily derives from the number of reticulation events. However, across values of *Ψ*, average posterior estimates track the true values because a larger proportion of networks have more reticulation events in them as the rate increases. As expected, the widths of posterior distributions decrease with both network size and increasing values of *Ψ*, because there is more information available to estimate the rate. The posterior distributions contain the true value in almost all cases. These results suggest that the reticulation rate can be estimated relatively accurately, though with a fair amount of uncertainty when the rate is low or the network is small.

In addition to visually interpreting posterior distributions of model parameters, we quantified the quality of posterior estimates using the relative root-mean squared error (which we call RMSE). For a given network ℵ with posterior distribution *P*(*θ* | ℵ) for parameter *θ*, the RMSE is

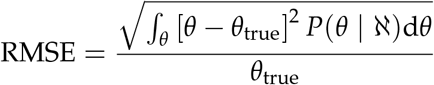

*i*.*e*., it is the square root of the average squared distance between the true parameter *θ*_true_ and an estimate *θ*, weighted by the posterior probability of that estimate and normalized by the true value. We chose this quantity for a number of reasons. First, the squared distance ensures that positive and negative errors do not cancel out. Second, taking the square root ensures that the unit of the numerator is the same as the unit of the parameter itself; further normalizing by the true value ensures that the quantity is dimensionless (it has no units), and is therefore comparable across parameters and scales. Finally, the RMSE captures both “bias” (in the sense that the RMSE will increase if the mean of the posterior distribution is farther away from the true value) and uncertainty (it will be large if the posterior distribution is wide).

As expected, the average RMSE (across replicates in a given parameter combination) is large when the reticulation rate is low, but decreases as *N* and *Ψ* each increase (black numbers, top left of panels in Fig. 6). In general, the RMSE for *Ψ* is between the RMSE of *µ* (which has relatively high RMSE) and *λ* (which has relatively low RMSE; see Supplemental Material). The main exception to this pattern is when *N* and *Ψ* are small, owing to very wide posterior distributions for cases where there are no reticulation events. Most notably, the RMSE significantly increases when we consider only replicates with at least one reticulation event (the conditional, RMSE_c_; red numbers, top left of panels in Fig. 6). The effect is large for small phylogenies and low rates— increasing the RMSE by a factor of up to 6—but decreases as the fraction of networks with reticulation events increases, suggesting that posterior estimates will be subject to a potentially strong ascertainment bias—induced by only studying phylogenies with reticulation events—when the true value of *Ψ* is low. Though they are not the focal parameter in our simulation study, posterior estimates of *λ* and *µ* are relatively sensible across simulation settings. Additionally, posterior estimates of all parameters were relatively insensitive to values of *u*_max_, though there was some effect for large values of *µ* and low values of *ρ* (*i*.*e*., cases where the number of extinct/unsampled lineages are more likely to be large). We present figures for all simulation scenarios and parameters, as well as tables summarizing the RMSE, RMSE_c_, and the coefficient of variation (the standard deviation of the posterior divided by the mean, characterizing uncertainty) in the Supplemental Material.

## Allopolyploid Diversification in Cystopteridaceae

A persistent debate in macroevolution regards the long-term evolutionary dynamics of polyploid lineages: are polyploid lineages fated for extinction, or are they instead significant long-term contributors to biodiversity? This debate is especially salient in plants, where more than 30% of recently formed plant species are estimated to be polyploids (Wood et al. 2009). The prevailing view in the 20th century was that polyploid lineages, despite enjoying some short-term ecological advantages, would suffer long-term evolutionary disadvantages, primarily due to their reduced genetic variation and to the selection-buffering effect of additional genomes (Stebbins 1950; Wagner 1970; Stebbins 1971). Consequently, polyploid lineages should struggle to adapt, and experience elevated rates of extinction over long timescales.

This “dead-end hypothesis” remains theoretically and empirically controversial (*e*.*g*., Mayrose et al. 2011; Tank et al. 2015; Vamosi et al. 2018; Rothfels and Otto 2026). In part, the longevity of this controversy is due to the challenges the reticulate nature of allopolyploidy poses to phylogenetic inference: before our BDR model, the available machinery for studying polyploid diversification over evolutionary time did not exist. This deficit led researchers to avoid empirical investigations of the macroevolutionary consequences of polyploidy or to adopt *ad hoc* approaches that ignore reticulation, which has unknown consequences (Mayrose et al. 2015; Rothfels 2021). Ideally, researchers would use network-based methods for addressing these questions.

We applied our BDR model to an allopolyploid phylogenetic network of the fern family Cystopteridaceae (fragile and oak ferns) to understand how allopolyploidization has shaped the diversification of this family. The Cystopteridaceae are noteworthy for having a large fraction of polyploid species, most of which are recognized as allopolyploids (Rothfels et al. 2017). The group is comprised of three genera, *Cystopteris, Acystopteris*, and *Gymnocarpium*, with≈ 40 recognized species distributed across the Northern Hemisphere (Pteridophyte Phylogeny Group II view). Ideally, we would jointly estimate the network and the parameters of the birth-death-reticulation process (Tribble et al. 2026), but implementing such a joint inference method is a significant technical challenge in its own right and the focus of our ongoing work. As a practical compromise, we employ a sequential procedure, where we fit the model to an existing estimate of the species network from Rothfels et al. (2017). Though our model assumes that diploid and polyploids diversify at the same rates, it nonetheless serves as a useful starting point, and allows us understand whether such a model provides an adequate description of the true diversification process.

### Cystopteridaceae species network

We performed our analyses on the phylogenetic network of Cystopteridaceae from Rothfels et al. (2017). This network is derived from a maximum-clade-credibility (MCC) multilabeled (MUL) species tree estimated from four nuclear gene sequences using `AlloppNET` (Jones et al. 2013). We obtained the .ai (Adobe Illustrator) file for Figure 7 of Rothfels et al. (2017), then used `ChatGPT` (OpenAI 2026) to construct the Extended Newick string corresponding to panel 7b, but with the ghost lineages removed, and rescaling the root age to *T* = 1. We discarded× *Cystocarpium*, as it is known to be a relatively recent, sterile allopolyploid, rather than an established allopolyploid population (Rothfels et al. 2015). We visually inspected the network to ensure it matched Rothfels et al. (2017). The resulting network comprises 26 tips, 17 of which are allopolyploids.

**Figure 7:**
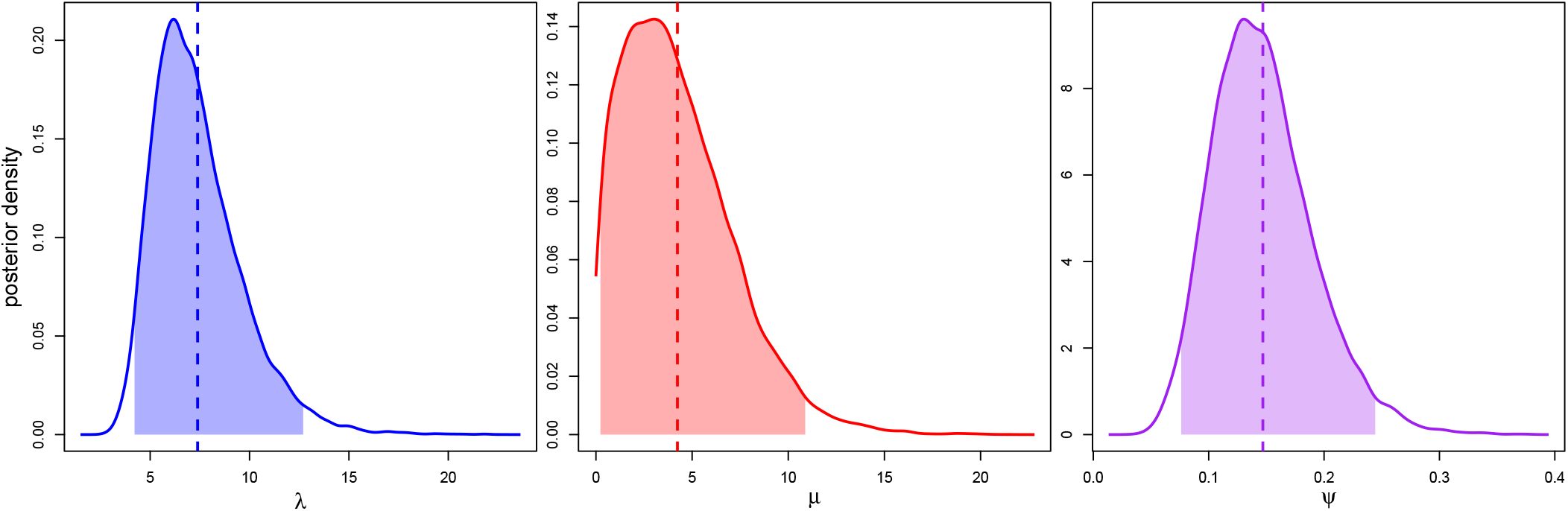
Posterior distributions of the birth-death-allopolyploidization parameters for Cystopteridaceae. We estimated the joint posterior distribution of the speciation rate, *λ*, extinction rate *µ*, and allopolyploidization rate *Ψ*, from the Cystopteridaceae network. The marginal posterior distributions are depicted as curves, with the posterior mean (vertical dashed lines) and the 95% credible intervals (shaded regions).

### Limitations of MUL trees

The derivation of the Rothfels et al. (2017) species network from the MUL tree has significant implications for our diversification analysis. A MUL tree represents each allopolyploid as multiple tips, one per subgenome, in a bifurcating tree. This approach is based on the (usually reasonable) assumption that the two subgenomes of an allopolyploid do not recombine, and therefore evolve effectively independently. However, the MUL tree does not record the age of the actual reticulation events; it shows the time at which the ancestors of each polyploid subgenome diverged from its most recent ancestor with a sampled diploid lineage but not the time that that subgenome lineage entered a polyploid. As a consequence, the MUL tree constrains but does not determine the ages of reticulation events in the corresponding species network. To derive a species network from a MUL tree, one must choose ages for reticulation events which obey the constraints imposed by the MUL tree. Specifically, a reticulation event can be no older than the time at which either of its two subgenomes diverged from their closest sampled relatives: divergences in the MUL tree are upper bounds on the ages of reticulation events. Furthermore, each subgenome implies the existence of a ghost lineage if its divergence time in the MUL tree is older than the reticulation event: if the reticulation event happened after both divergences, then there are two ghost lineages; after one divergence, one ghost lineage. If no divergences are older than the reticulation event, then there are no ghost lineages, but this situation requires that both divergences in the MUL tree are of exactly the same age.

In any case, the determination of a species network from a MUL tree depends on some assumptions that may have consequences on diversification analysis because they imply different numbers of extinction events. In general, Rothfels et al. (2017) chose the ages of allopolyploidization events to be as old as possible, implying the minimal number of ghost lineages. In the analyses that follow, we either assume the network of Rothfels et al. (2017) is the true network (taking the old ages at face value), or consider all possible ages that are consistent with the MUL tree.

### Bayesian analysis

We estimated the posterior distribution of the allopolyploid BDR model from the Cystopteridaceae network using priors similar to those described in our simulation study, with slight modification:

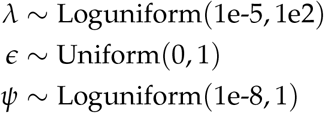

and *ρ* = 0.5, because there are about 50 Cystopteridaceae species, assuming approximately 10 currently undescribed species (see, *e*.*g*., Rothfels et al. 2017, 2014).

To avoid taking the reticulation ages in the Rothfels et al. (2017) network at face value, we considered all possible networks that are consistent with the MUL tree. As described above, the network we extracted from Rothfels et al. (2017) assumes the maximum possible reticulation ages (and minimum number of ghost lineages) that are consistent with the MUL tree. Therefore, to consider all possible ages consistent with the MUL tree, we have to sample ages that are no older than those in the input network ℵ. To do this, we augment our Bayesian model to include the ages of reticulation events, 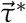, with the corresponding posterior distribution

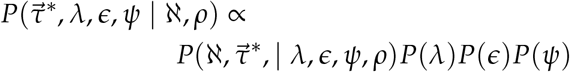

The probability density of the augmented network is

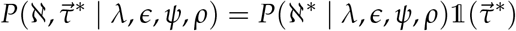

where ℵ^∗^ is the augmented network—*i*.*e*., the network ℵwith the reticulation ages replaced by 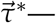and 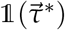 is an indicator function that ensures the augmented ages are consistent with the network ℵ

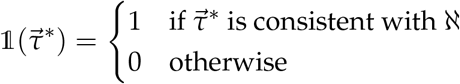

(where consistent means has ages no older than those in ℵfor each event). We provide further theoretical justification for this procedure in the Supplemental Material.

We sampled from the above augmented posterior distribution using MCMC, effectively sampling over reticulation ages that are consistent with the MUL using MCMC proposals. However, the augmented ages are not only bounded between the present and the maximum age of the corresponding reticulation event, they may also be exactly as old as the maximum age; for a particular reticulation event age 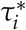 with maximum *m*_*i*_, the chain must explore values such that 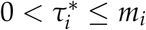. We therefore designed MCMC proposals that proposed continuous changes to *τ*_*i*_, as well as changes between *τ*_*i*_ = *m*_*i*_ and *τ*_*i*_ ≠ *m*_*i*_ (essentially reversible-jump MCMC on an augmented variable; Green 1995). We describe our MCMC sampler in the Supplemental Material. We performed our main MCMC analyses under five different values of *u*_max_, *u*_max_ = {32, 64, 128, 256, 512}. These analyses sampled the model parameters *λ, €*, and *Ψ*, as well as the reticulation event ages 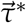 using the custom MCMC sampler described above. As a partial validation of our custom MCMC sampler, we repeated these analyses using our sampler on the fixed network (with proposals on reticulation ages turned off), as well as the same `AdaptiveMCMC` code we used for the simulation study. To assess the impact of the fixed sampling fraction *ρ*, we also performed analyses with *ρ* = 1. In all cases, we ran MCMC analyses for 100,000 iterations, and discarded 10% as burnin. All analyses mixed well, with post-burnin samples satisfying ESS *>* 1000 for all parameters (and allopolyploid ages when those were estimated).

### Assessing model adequacy

To explore potential differences in diversification rates between polyploids and diploids in the Cystopteridaceae, we used Bayesian posterior-predictive simulation (Gelman et al. 1996) to assess whether the birth-death-allopolyploidization model—which assumes that diploids and polyploids diversify at the same rate—provides an adequate description of the actual network-generating process. Posterior-predictive simulation involves simulating datasets from parameters sampled from the posterior distribution, and comparing those simulated datasets to the original empirical dataset; if the simulated datasets resemble the empirical dataset, then the model is considered adequate. This resemblance can be formalized using test statistics computed on the simulated and observed datasets, and measured by a posterior-predictive p-value, which is the fraction of simulated statistics that are more extreme than the observed statistic (either as a one-tailed or two-tailed test). Different posterior-predictive test statistics will capture different features of the data, and may therefore be more or less sensitive to different model violations (for phylogenetic examples see, *e*.*g*., Schwery et al. 2023; Mulvey et al. 2025).

We targeted our posterior-predictive simulations at detecting model violations that would be consistent with the dead-end hypothesis. The dead-end hypothesis predicts that polyploid lineages have elevated extinction rates and/or depressed speciation rates, and so should be relatively young. We therefore used the mean polyploid age as our primary test statistic.

We drew samples from the posterior distribution of our empirical analysis with estimated reticulation ages and *u*_max_ = 512, and for a given parameter sample, simulated a network with 26 total tips and 17 allopolyploid tips using rejection sampling.

We conditioned on the number of tips and allopolyploids because otherwise the variance in network size was extremely large, confusing the interpretation of the distribution of statistics. We repeated this procedure for 1000 posterior samples, generating 1000 simulated test statistics. We then computed the test statistic on the empirical network in two ways: first, we computed the test statistic on 1000 reticulation ages sampled from the posterior; second, we computed the test statistic using the posterior mean age of each reticulation event.

### Results

We report results for the analyses with *u*_max_ = 512, and with estimated reticulation ages. Results for different values of *u*_max_ indicate convergence at around *u*_max_ = 128. Our other analyses indicate that the inference of the reticulation ages has a modest effect on posterior estimates of rate parameters (versus using the fixed ages in the Rothfels et al. (2017) network), and that our custom MCMC machinery gives the same results as AdaptiveMCMC. We present these auxiliary results in the Supplemental Material.

#### Inferred rates of diversification and allopolyploidization

Marginal posterior distributions of the rate parameters are depicted in Figure 7. Per-lineage rates of speciation (posterior mean 7.3, [4.2 −12.7] 95% credible interval [CI]) exceed extinction rates by almost two-fold (posterior mean 4.2, [0.25 −10.9] 95% CI). With a positive net-diversification rate (*λ*−*µ*, posterior mean 3.14, [0.62−5.49] 95% CI), diploid Cystopteridaceae species are diversifying fast enough to avoid clade extinction on their own. The per-lineage-pair allopolyploidization rate is relatively small (posterior mean 0.14, [0.076−0.24] 95% CI), but the fact that it is pairwise makes limits direct comparison to the speciation and extinction rates.

#### Ages of reticulation events, and the plausibility of ghost lineages

We estimated the posterior distribution of the age of each reticulation event in the Cystopteridaceae network. We computed the posterior mean age of each reticulation event, producing a posterior mean summary network (Fig. 8). For allopolyploid nodes whose ages we estimated (that is, whose ages were not uniquely defined by the MUL tree), the mean ages were generally younger than the maximum possible age. The inferred ages of allopolyploid nodes is directly related to the inferred number of ghost lineages, *i*.*e*., parents of allopolyploids that subsequently went extinct (or were unsampled at the present). In fact, our analysis allows us to assign posterior probabilities that particular allopolyploids have ghost parents.

**Figure 8:**
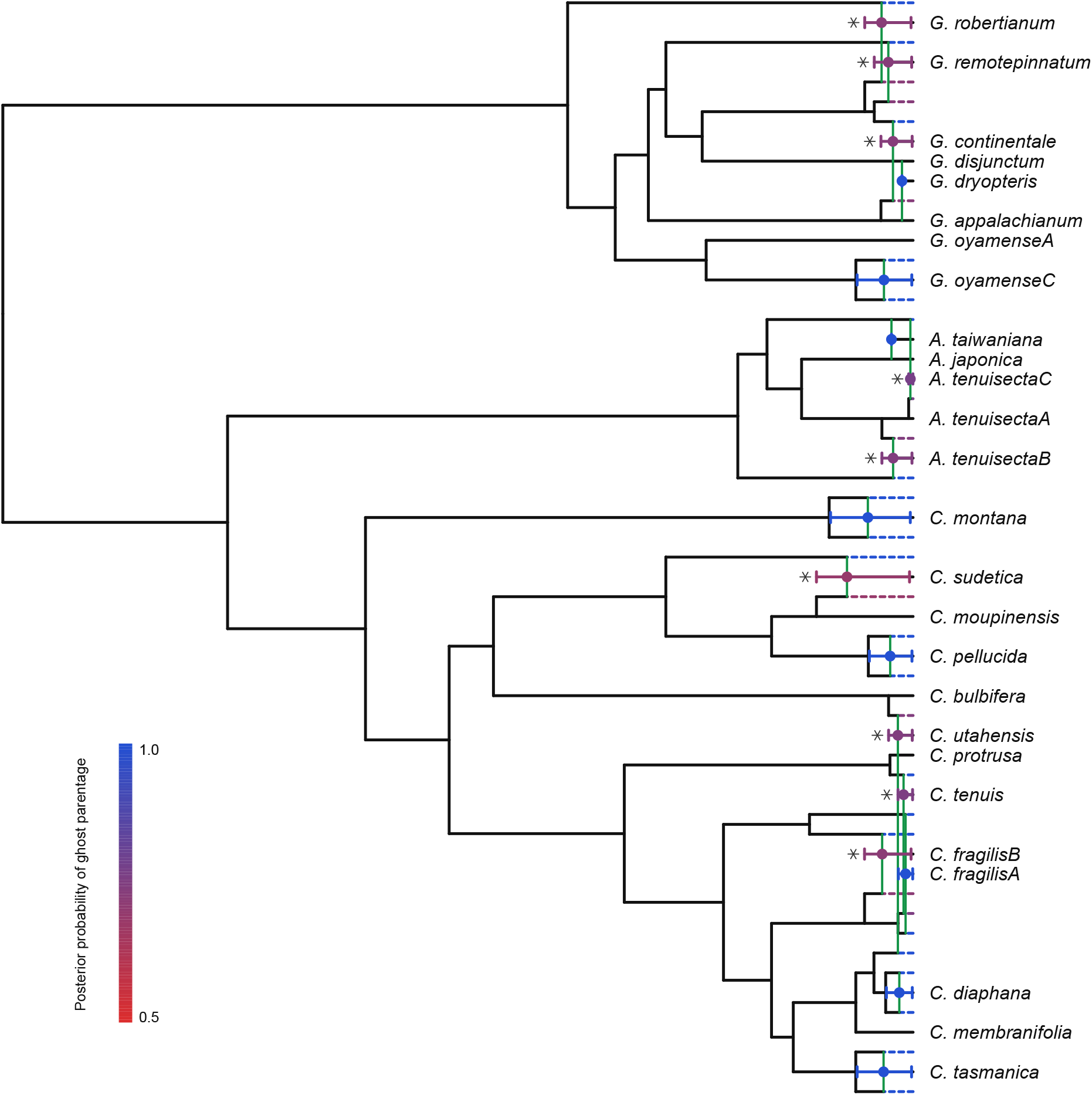
Summary network of Cystopteridaceae estimated using MCMC under the birth-death-allopolyploidization model. We estimated the parameters of the birth-death-allopolyploidization model as well as the reticulation ages of the phylogenetic network from Rothfels et al. (2017). Reticulation event ages were constrained by the maximum age in the Rothfels et al. (2017) network, as described in the text. We computed a summary network based on the posterior estimates of the ages of the reticulation events. Posterior mean ages of events are depicted as dots, with horizontal intervals depicting the 95% credible interval of the ages. Dashed horizontal lines depict ghost lineages. For some reticulation events (marked with asterisks), there is some ambiguity about whether one of the parents is a ghost lineage; in these cases, the nodes (and corresponding ghost lineage) are colored according to the posterior probability of ghost parentage. Other reticulation nodes require the invocation of ghost lineages by the nature of the topology, in which case the nodes and ghost parents have a posterior probability of 1.

For many of the allopolyploids in our network, both subgenomes trace back to a recent common ancestor that left no other descendants (*e*.*g*., *C. tasmanica*); in these cases, the only possible explanation is that both of the parents are extinct or unsampled (two ghost lineages), and so the posterior probability of two ghost parents is 1. Other allopolyploids have identical divergence between their two subgenomes and the corresponding parental lineages (*e*.*g*., *A. taiwaniana*) indicating that both parents survived (again with probability 1). The remaining nodes in the tree have two parents of unequal age, in which case the older parent must be a ghost, but the younger parent may or may not be a ghost. For example, *C. sudetica* has one parent that must be a ghost because the lineage branches off from deeper in the network; however, the question remains whether the other parent was the ancestor of *C. sudentica*, or a ghost side branch. Allopolyploids where the younger parent was inferred to be a ghost are indicated by asterisks in Figure 8. In all cases, the posterior probability of the ghost parent is *>*70%, though no case is decisive, with higher posterior probabilities for older allopolyploids (*e*.*g*., *C. sudetica*) compared to younger ones (*e*.*g*., *A. tenuisectaC*).

The network reconciliation of Rothfels et al. (2017) implies a total of 21 ghost lineages, which is the minimum number required to explain the network. By contrast, our summary network depicts 30 ghost lineages, which is the maximum possible. This is somewhat illusory, as the marginal probabilities of each ghost lineage are ≈70%. In reality, the posterior distribution of networks sampled in our analysis support a range of ghost lineage numbers, with a posterior mean of 27.63 ([25 −30] 95% CI, see Supplemental Material).

#### Model inadequacy

Our posterior-predictive simulations reveal that the (equal-rates) BDR model is insufficient to describe the true diversification process for Cystopteridaceae. In particular, allopolyploids in simulated networks (Fig. 9, red) are much older than those in the empirical network (Fig. 9, blue and black). The posterior-predictive p-values of the empirical mean ages are 0, whether we consider just the network with mean reticulation ages or the range of reticulation ages in the posterior sample. In other words, a model where the diversification rates of diploids and polyploids are the same predicts much older allopolyploidization events than does our empirical data: we can reject the equal-rates model. The significantly young age of the empirical allopolyploidizations lend support to the dead-end hypothesis, which favors young, transient allopolyploid lineages over old, persistent ones. We return to the implications of these results in the discussion.

**Figure 9:**
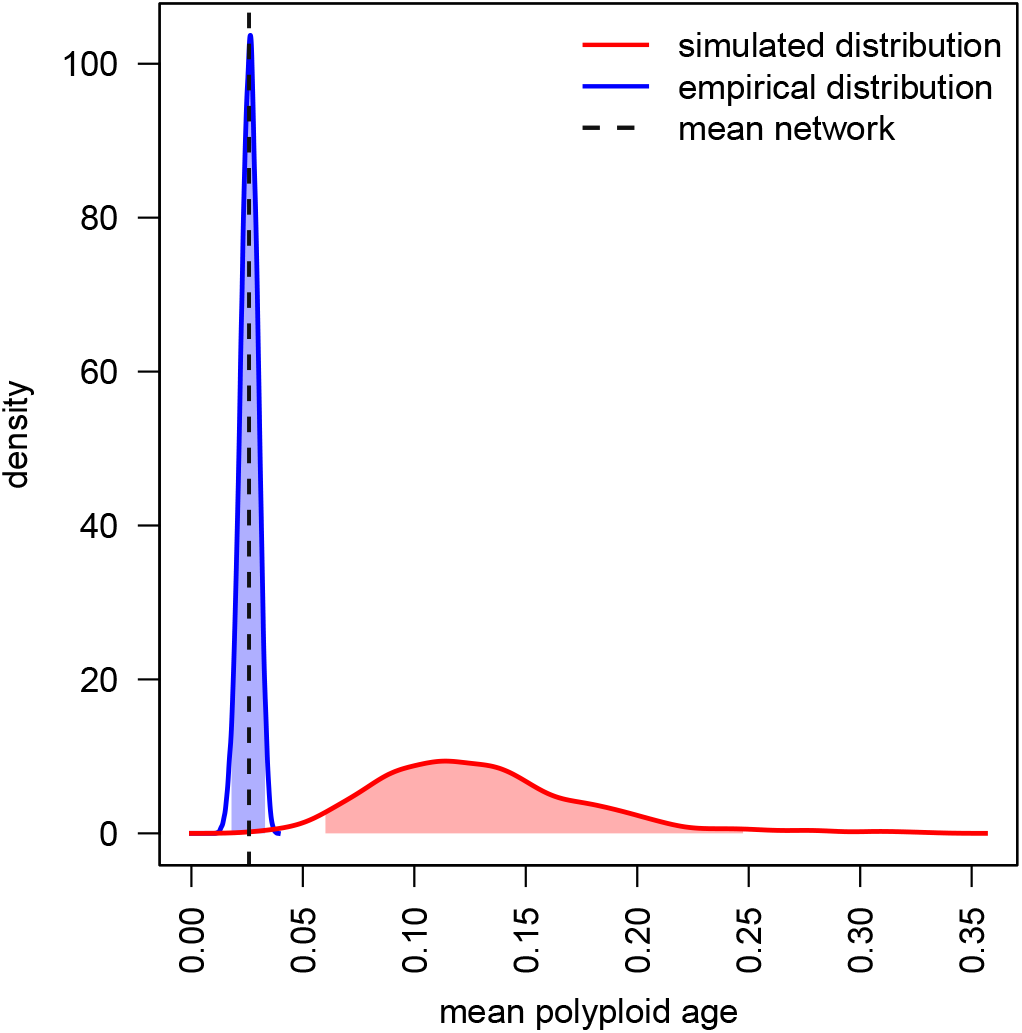
Posterior-predictive adequacy of the birth-death-allopolyploidization model for Cystopteridaceae. We simulated networks using parameters sampled from the posterior distribution of our empirical analysis. We computed the mean age of polyploid lineages in each of those simulated networks (red curve shows distribution of mean ages across simulated networks), and compared against the corresponding mean age of polyploid lineages in the empirical network (using either the mean event ages, vertical dashed line, or a distribution of event ages, blue distribution). Shaded regions of each distribution show the 95% equal-tailed predictive interval (analogous to the 95% credible interval of a parameter).

## Discussion

We present a simple BDR process that generates rooted, ultrametric phylogenetic networks. The model makes many simplifying assumptions, for example, that reticulation events are effectively instantaneous rather than protracted, and that all lineages are equally likely to reticulate with each other. However, this simplicity allows us to derive an explicit (if approximate) probability density function for phylogenetic networks under the model. In turn, this probability density is suitable for application as a likelihood function in a sequential inference framework focused on estimating diversification and reticulation rates (*i*.*e*., on a pre-estimated network, as we do here with Cystopteridaceae), or as a prior distribution in a Bayesian phylogenetic network inference framework. While we believe this model represents a useful step in understanding the processes that shape phylogenetic networks, we now turn to discussing important caveats, implications, and future developments.

### Statistical and numerical properties

Our simulations demonstrate that reticulation rates can be estimated from known phylogenetic networks. Naturally, the uncertainty in reticulation rate estimates is high for networks with no reticulation events (*i*.*e*., phylogenetic *trees*), but diminishes with even a modest number of reticulation events (*e*.*g*., Fig. 6). However, we must temper these promising results with some comments about statistical behavior and the numerical properties of our approximation (namely *u*_max_).

#### Acquisition bias

Researchers may be unlikely to apply a BDR process to phylogenetic trees, where standard birth-death models are available. This acquisition bias has the potential to lead to systematic bias, because the BDR process will tend to be applied only to networks with at least one reticulation, despite the fact that it will produce strictly bifurcating trees with some frequency, particularly when reticulation rates are low. This problem is similar to that in discrete morphological phylogenetics: in practice, we only analyze (discrete) morphological traits that vary among our taxa, even though our models will predict that some traits are invariant, resulting in upwardly biased rate estimates (Lewis 2001).

Indeed, our simulations indicate that only considering networks with at least one reticulation event dramatically inflates estimates of the reticulation rate when the true rate is low (see Fig. 6). Unfortunately, we were unable to derive a conditional probability density to correct for this bias. An analytical solution seems rather hopeless, but we continue to work on practical numerical solutions. In the meantime, researchers should interpret rate estimates with caution if they are studying networks with one or a few reticulation events.

#### Identifiability

The approximate nature of our probability density function makes it difficult to derive clean analytical statements about parameter identifiability. However, we should expect the BDR model to inherit most of the problems of birth-death models. We suspect, for example, that *λ, µ*, and *ρ* are difficult to estimate simultaneously (*i*.*e*., one must be fixed to do reliable inference; Stadler 2009), and that the extinction rate may only be weakly identifiable.

Currently, we can only speculate about the identifiability of reticulation-rate parameters. Because reticulation events leave an unmistakable signature that cannot be mimicked by speciation or extinction, we are optimistic that reticulation rates are unlikely to be confounded with *λ, µ*, or *ρ* (though it is possible in principle). However, we note that different types of hybridization events can leave similar histories of reticulation in sampled phylogenetic networks; for example, unidirectional and bidirectional hybridization look the same if one of the parental lineages goes extinct. As a result, it may be difficult to separately estimate *η, ζ*, and *ν*, particularly when extinction rates are high or sampling probabilities are low.

#### The importance of u_max_

Our probability density function depends on a truncation parameter, *u*_max_, which represents the maximum number of lineages that are allowed to exist over the history of the process. If *u*_max_ is sufficiently large, the approximate probability density should approach the true, untruncated probability density. However, a large *u*_max_ necessarily increases the time it takes to compute the probability density; runtime should scale approximately with 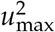. Researchers may therefore be motivated to choose relatively small values of *u*_max_, but a value that is too small could compromise parameter estimates.

In our simulations, we found that setting *u*_max_ to two to four times the number of sampled taxa was generally sufficient to achieve an accurate approximation, but naturally this will depend on features of the dataset. Our empirical analyses tended to stabilize by *u*_max_ = 128, with runtimes on the order of an hour (to achieve very large ESS) on a modern laptop, suggesting that good approximations are not out of practical reach. While it may be possible to develop a theoretically grounded, dynamic method for determining *u*_max_ based on parameter values, for now we consider it good practice to increment *u*_max_ to the point that parameter estimates are stable.

### The practicality of sequential inference

The common practice in phylogenetic diversification analysis is to estimate a phylogenetic tree from sequence data, then fit a diversification model to a point estimate of the tree (or, in some cases, a sample of trees; Morlon 2014). We refer to this as sequential inference. A similar approach may be taken with our probability density, essentially using it as a likelihood function on a pre-estimated phylogenetic network. Such an approach has all the usual issues of sequential inference for diversification modeling, for example the assumptions made in the first step may bias downstream inferences in one direction or another (Tribble et al. 2026). However, sequential analysis of networks may have special hazards.

The primary concern is that different network inference programs will be able to infer (or have an implicit preference for) different reticulation histories, which could bias downstream sequential inferences. The most popular quartet-based method for inferring hybrid networks, SNaQ (Solís-Lemus and Ané 2016), is limited in the number and configuration of permitted reticulation events, focusing on identifiable level-1 phylogenetic networks (networks where reticulate histories do not overlap; Solís-Lemus and Ané 2016; Kong et al. 2022; Kolbow et al. 2026). Recent work demonstrates that some level-2 networks are identifiable (under the Jukes-Cantor model, Englander et al. 2025, with quartets, Holtgrefe et al. 2025), but methods for inferring such networks do not yet exist. While model-based network inference methods that do not constrain the number and configurations of reticulation events do exist (*e*.*g*., `PhyloNet` for hybrid networks, Wen et al. 2018; `AlloppNET` for allopolyploid networks, Jones 2017), the identifiability of reticulation events under these models is an open question.

Importantly, our BDR model can generate a wide range of phylogenetic networks without special regard to the number and configuration of the reticulation events. For example, it is not a probability distribution on just level-1 networks. Because the limits and biases of network inference methods are themselves not well understood, it is difficult to predict how a sequential procedure will behave when the assumptions of the birth-death-reticulation process are violated in the first, network-inference, step.

### Implications for network inference

Network-inference methods often only consider the probability of a set of sequences (or gene trees relating those sequences) when estimating the network, without regard to the plausibility of the implied diversification history. Methods like `SNaQ` (Solís-Lemus and Ané 2016) and `PhyloNet` (Wen et al. 2018) estimate networks that are not constrained by time, so that reticulation events may connect lineages that are in reality temporally distant. The timing of these events implies something about the existence of ghost lineages, the plausibility of which will necessarily depend on the ages of the participating lineages and on rates of extinction and sampling.

Other network inference methods, notably Bayesian multispecies-coalescent methods like `AlloppNET` (Jones et al. 2013; Jones 2017) and `SpeciesNetwork` (Zhang et al. 2018), require a prior density on the network. The network densities used by these approaches are arbitrary or have questionable empirical justification. For `AlloppNET,` the network density takes the form of 1) a Yule (pure birth) prior in the case of a MUL tree; 2) separate birth-death priors on diploid and polyploid subtrees in the case of a network with one allopolyploidization, and; 3) or an arbitrary density in the case of a network with an estimated number of allopolyploidization events. For `SpeciesNetwork`, the prior density on the species network is a Yule-hybridization model, which ignores both extinction and incomplete sampling, and models the hybridization process as the merging of two lineages rather than the exchange of genetic material (*i*.*e*., it is a lineage-degenerative process *sensu* Justison and Heath 2024).

We might hope that genetic data are sufficiently informative that either ignoring the plausibility of reticulation events (*e*.*g*., `SNaQ` or `PhyloNet`) or using an arbitrary density on reticulate networks (*e*.*g*., `AlloppNET` or `SpeciesNetwork`) has little impact on the estimates we care about. It seems likely that the number and placement of reticulation events is primarily informed by genetic data. For example, in the context of the multispecies network coalescent, gene trees may imply coalescent events in ancestral populations that require (or strongly favor) the invocation of reticulation events. If our primary goal is to infer the number of reticulation events or which lineages have hybrid ancestry, then ignoring the diversification dynamics or using an arbitrary network density may be perfectly acceptable.

However, inferring the ages of reticulation events from genetic data alone may be more challenging. In the case of allopolyploids, where the subgenomes do not recombine (at least by assumption), coalescent events between each subgenome and their respective closest relatives only provide an upper bound on the age of the allopolyploidization event. Therefore, estimates of exact ages of allopolyploidization events should be nonidentifiable from genetic data alone. The situation is more complicated with hybridization, because coalescent histories may be informative about whether introgressed genes trace their ancestry through ghost lineages (Ottenburghs 2020), thus providing some information about the age of the hybridization event. However, this information may be limited when the number of genetic lineages with the potential to coalesce is small (*i*.*e*., when population-level sampling is sparse or hybridization events are very old), and is likely to depend on assumptions about population sizes (coalescent rates) that may be difficult to justify. For these reasons, we should expect the chosen network density—or the lack of any explicit network density—to have potentially significant influence on estimates of the ages of reticulation lineages. Explicit BDR models—either the simple one we describe here or a more complex variant—provide coherent and biologically interpretable prior densities for a joint network inference framework, and a formal way to evaluate the plausibility of the timing of reticulation events. We believe such models are critical for making statements about the geological or temporal context in which reticulate lineages arose, and for macroevolutionary inferences that depend on the ages of reticulate lineages, for example Stebbins’ (1971) dead-end hypothesis.

### Cystopteridaceae, ghost lineages, and the dead-end hypothesis

Our analysis of an allopolyploid phylogenetic network of Cystopteridaceae ferns highlights some of the opportunities and associated with network diversification analysis. We performed these analyses using an existing network derived from a MUL tree estimated with `AlloppNET` (Jones et al. 2013); consequently, this network is only informative about the maximum ages of the allopolyploid lineages.

In order to accommodate this uncertainty in a sequential framework, we sampled over all possible ages of the allopolyploidization events, which required careful consideration of the network topology and the design of special MCMC algorithms. Our analyses demonstrate that fixing the ages to those in the original network (*i*.*e*., taking the reconciliation of Rothfels et al. (2017) at face value) had a modest but non-zero effect on inferred diversification rates, compared to our age-sampling approach. Sampling the ages of allopolyploid lineages also provided us with the opportunity to evaluate the plausibility of ghost lineages. The network presented by Rothfels et al. (2017) invokes the minimum number of ghost lineages (21) necessary to reconcile the network with the MUL tree. In contrast, we estimate that there were probably 27–28 ghost lineages (posterior mean 27.63). This inference may seem unparsimonious, particularly because many of the allopolyploids are quite recent and therefore their parents should not have had much time to have gone extinct. However, our sampling fraction is only *ρ* = 0.5, which allows even recent parents to be unsampled. Subsequent analyses using *ρ* = 1 indicate that higher sampling fractions disfavor recent ghost lineages (see Supplemental Material), indicating that these inferences should be considered fairly sensitive to assumptions about extinction and sampling rates.

While our model does not accommodate different diversification rates for allopolyploids, we were nonetheless able to use Bayesian posterior-predictive simulation to uncover evidence in favor of the dead-end hypothesis in the Cystopteridaceae. Stebbins’ dead-end hypothesis (Stebbins 1950, 1971) posits that polyploid lineages will exhibit elevated extinction rates (compared to diploid lineages) because of genetic masking, *i*.*e*., the effect of advantageous mutations will be reduced because polyploids have more copies of each gene, limiting adaptability. To address this question, we simulated networks using parameters estimated from the Cystopteridaceae network, then compared the ages of simulated polyploids to the ages of polyploids in the Cystopteridaceae network.

Our posterior-predictive simulations revealed that simulated polyploid ages were always significant older than observed polyploid ages. This result is consistent with the dead-end hypothesis, which predicts that polyploid lineages should be relatively short-lived. Of course, other explanations of our results are possible. For example, an erroneously large *ρ* could push simulated ages forward (make reticulations tend to be younger) and could explain the discrepancy between the empirical and simulated ages in our study. However, we assumed a total of ≈50 Cystopteridaceae species, which is probably a considerable underestimate given the number of inferred cryptic species in the family (see, *e*.*g*., Rothfels et al. 2017; Ekrt et al. 2022), and thus our estimate of *ρ* is probably an overestimate. Alternatively, if extinction rates are decreasing over time and/or speciation rates are increasing, that would push nodes toward the present compared to the expectations of our model. Our model also ignores the role of genetic divergence and geography on the propensity of parents to produce allopolyploid daughters; however, these factors should be operating over the entire history of the group, and it is not obvious that they would drive reticulation events in one direction over another. Although these results are quite consistent with the predictions of the deadend hypothesis, more work is needed to develop models that explicitly incorporate differential diversification for polyploids, and to understand the factors that might confound such inferences.

### Moving forward with birth-death-reticulation models

The BDR process we present is about the simplest process one can imagine for modeling the generation of phylogenetic networks. (The Yule-hybridization process of Zhang et al. (2018) is simpler but ignores key components like extinction and incomplete sampling.) Despite its simplicity, the probability density of a network under this model can only be computed approximately, and even so is quite computationally demanding compared to standard birth-death models. Naturally, this situation raises questions about how far our simple model can be extended before it breaks, either theoretically or practically.

Our birth-death-reticulation process leaves out many key features that could affect the production, establishment, and diversification of reticulate lineages. For example, we ignore the influence of genetic distance (Woodhams et al. 2016; Justison and Heath 2024), geography, ecology, phenology, and other factors on the propensity for lineages to reticulate. Likewise, our model lacks the parameters allowing reticulate lineages to diversify at different rates (*i*.*e*., state dependence), denying us the ability to make direct macroevolutionary inferences about differential diversification rates.

We view the simple BDR process as analogous to the constant-rate birth-death process (Nee et al. 1994), which likewise makes many simplifying assumptions. Of course, the birth-death process has been extended to accommodate time dependence (Stadler 2011; Höhna 2015), state dependence (Maddison et al. 2007), lineage dependence (Maliet and Morlon 2021; Barido-Sottani et al. 2020; Höhna et al. 2026), and fossilization (Heath et al. 2014), among many other phenomena. In principle, our model can similarly be extended to include these phenomena. However, our approach for approximating the probability density of a network—using a truncated set of ODEs to average over all possible histories of unobserved lineages—may not be practical for more complex models, necessitating the development of alternative approximation strategies.

We believe a joint Bayesian multispecies network coalescent framework is the most promising avenue for the study of reticulate diversification. In this framework, we simultaneously estimate the gene trees, the network, and the parameters of the network-generating process directly from genetic sequence data. Practically, this framework entails using the BDR process (or some variant of it) as a prior on the network, and estimating the hyperparameters of the process in a hierarchical model.

Of course, the joint Bayesian framework combines two already difficult problems—network inference and network-diversification modeling—and we should anticipate many statistical and computational challenges. Nonetheless, this framework may have some significant advantages. First, it avoids the significant conceptual and technical issues related to sequential inference discussed above (and in Tribble et al. 2026), and allows the diversification process to inform inferences about the plausibility of inferred reticulation histories. Second, Markov chain Monte Carlo may support approximation machinery for complex network-diversification models—for example, data augmentation (see, *e*.*g*., Maliet and Morlon 2021; Ronquist et al. 2021, for similar approaches for complex birth-death models)—that are unavailable in a sequential framework. We are optimistic that continued development of Bayesian network inference methods and BDR models will allow evolutionary biologists to address critical questions about the diversification of reticulation lineages.

## Software and data availability

The `C++` library `Diversinet` and the `julia` package Diversinet.jl are both released under the open source `GNU GPL3` license, and are actively maintained at https://github.com/mikeryanmay/Diversinet/ and https://github.com/mikeryanmay/Diversinet.jl/, respectively. Versions `v0.1.0` of both softwares were used for this study (tagged at their respective repositories). Users can install `Diversinet.jl v0.1.0` and all of its dependencies (including the C++ library) by following instructions at https://github.com/mikeryanmay/DiversinetRegistry; a Docker image is also available (see https://github.com/mikeryanmay/DiversinetDocker for instructions).

All of the data used in this study are available in the Data Dryad repository at https://doi.org/10.5061/dryad.k6djh9wpf. Code for recreating all of the analyses in this study are available at 10.5281/zenodo.22693703.

## Declaration of AI usage

We used AI (Codex) at various stages of this project. In all cases, these applications reflect our own design decisions with pipelining, debugging, and modest implementation facilitated by AI. We wrote the `C++` library `Diversinet` and the `julia` package `Diversinet.jl` ourselves, but Codex assisted in developing the code for building and distributing the Diversinet.jl package (and associated Docker container). We used Codex to extract the empirical Cystopteridaceae network from Fig. 7 in Rothfels et al. (2017), and to implement the complex MCMC proposals on reticulation-event ages (based on our descriptions of the proposals; see Supplemental Material). We also used Codex to automate the tedious steps of the validation analyses, in particular the categorization of Monte Carlo simulations into target topologies. Finally, we used Codex to generate pipelines for the validation analyses, simulation study, and empirical analyses.

## Acknowledgements

We are grateful to X anonymous reviewers, as well as the members of the Rothfels lab, for their feedback on this study. This study was supported by NSF CAREER grant #2144011 to CJR.

