## Supplemental Material for "An explicit birth-death-reticulation model for studying the diversification of phylogenetic networks"

### Electronic Supplemental Material For: A simple birth-death-reticulation model for phylogenetic networks

#### Contents


**S1 The probability density of a network under a simple birth-death-reticulation process** **S2**
S1.1 Transition probabilities between nodes . . . S2
S1.2 Node-event densities . . . S7

**S2 Validation** **S10**
S2.1 Parameter tuning . . . S10
S2.2 Enumerating the network topologies . . . S11
S2.3 Monte Carlo simulation . . . S11
S2.4 Numerical integration . . . S11

**S3 Simulation Study** **S17**
S3.1 Simulation procedure . . . S17
S3.2 MCMC analyses . . . S17
S3.3 Extended results . . . S18

**S4 Empirical analyses** **S22**
S4.1 Bayesian augmented network analysis . . . S22
S4.1.1 Theory . . . S22
S4.1.2 Implementation . . . S23
S4.2 Ghost lineages . . . S26
S4.3 Sensitivity of $u_{\text{max}}$ . . . S27
S4.4 The effect of estimating allopolyploidization ages . . . S28
S4.5 Consistency between custom sampler and `AdaptiveMCMC` . . . S29
S4.6 Sensitivity of ghost lineages to the sampling fraction $\rho$ . . . S30

### S1 The probability density of a network under a simple birth-death-reticulation process

#### S1.1 Transition probabilities between nodes

Our goal is to derive ordinary differential equations that allow us to compute how $p_u$ changes over the interval $(\tau_i, \tau_j)$. Our approach closely follows Maddison et al. (2007), beginning with finite difference equations to derive a set of ordinary differential equations (ODEs) that describe how $p_u(t)$ changes over time, assuming $s$ is fixed (as it is over a time interval). These differential equations will be used to solve for $p_u(\tau_i)$ (at the left of the interval) given $p_u(\tau_j)$ (at the right of the interval), allowing us to move the probability density down the network toward the next node event. In the next sections, we assume the number of unobserved lineages is not at a boundary (0 or $u_{\max}$) for simplicity; the boundary cases require only modest modifications, which we describe later.

*Finite difference equations.*—To derive finite difference equations, we assume that $p_u(t)$ is known for all $u$ and attempt to compute the probability some small interval of time $\Delta$ later, $p_u(t+\Delta)$. We assume $\Delta$ is small enough that at most one event occurs. We then enumerate all possible scenarios that could occur over time $\Delta$ that could lead to the network descending from time $t+\Delta$:

(a) nothing happens
(b) a speciation event occurs from a sampled lineage, producing an unobserved lineage
(c) a speciation event occurs from an unsampled lineage, producing an unobserved lineage
(d) an extinction event occurs to an unobserved lineage
(e) a unidirectional hybridization event occurs from a sampled lineage to an unoberved lineage
(f) a unidirectional hybridization event occurs between two unobserved lineages
(g) a bidirectional hybridization event occurs between two unoberved lineages
(h) a hybrid speciation event occurs between two sampled lineages, but the hybrid daughter is unobserved
(i) a hybrid speciation event occurs between two unobserved lineages, and the hybrid daughter is unobserved
(j) a hybrid speciation event occurs between a sampled lineage and an unoberved lineage, and the hybrid daughter is unobserved
(k) a polyploid speciation event occurs between two sampled lineages, but the polyploid daughter is unobserved
(l) a polyploid speciation event occurs between two unobserved lineages, and the polyploid daughter is unobserved, and
(m) a polyploid speciation event occurs between a sampled lineage and an unoberved lineage, and the polyploid daughter is unobserved

These scenarios are depicted in Fig. S1. Note that we are excluding events that could have occurred in the interval $(t+\Delta, t)$ but that could not have produced the observed network. For example, a speciation event producing a new sampled lineage is impossible because it could not result in the observed network (*i.e.*, there would be a node event at that time).

The probability that an event with rate $\theta$ occurs once over an interval of size $\Delta$ is approximately $\theta\Delta$. We can use this to compute the probability of the data at time $t+\Delta$ given that an event of type $x$, which we denote $p_u^x(t+\Delta)$. This probability for an event of type $a$ is (with $n = s+u$, and $s$ the

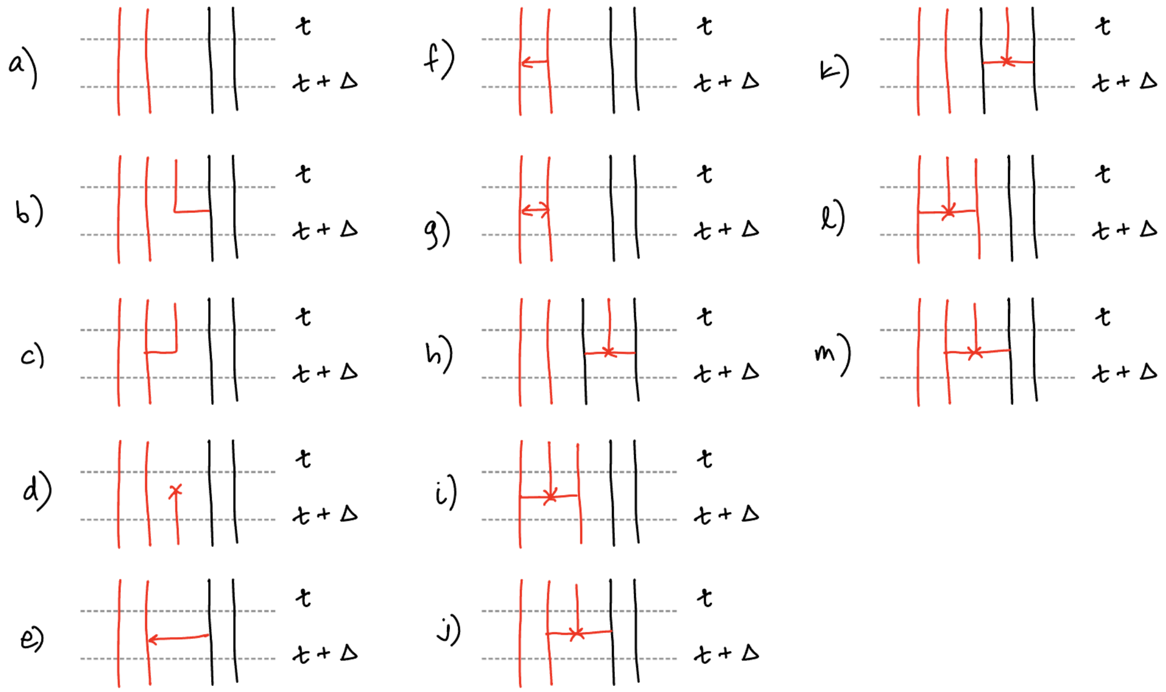


**Figure S1: Transition events leading to observed networks.** Sampled lineages are black and unsampled lineages are red. Unsampled lineages exiting time $t$ (the top) must subsequently go extinct, but are able to participate in diversification and reticulation events inside the interval between $t$ and $t + \Delta$.

number of sampled lineages at a given time $t$):

$$p_u^a(t+\Delta) = \left[ \overbrace{(1 - n\lambda\Delta)}^{\text{no speciation}} \overbrace{(1 - n\mu\Delta)}^{\text{no extinction}} \overbrace{\left(1 - \binom{n}{2}\eta\Delta\right)}^{\text{no unidirectional hybridization}} \overbrace{\left(1 - \binom{n}{2}\zeta\Delta\right)}^{\text{no bidirectional hybridization}} \overbrace{\left(1 - \binom{n}{2}\nu\Delta\right)}^{\text{no hybrid speciation}} \overbrace{\left(1 - \binom{n}{2}\psi\Delta\right)}^{\text{no allopolyploid speciation}} \right] p_u(t)$$

The probability of no events is the product of the probabilities of each individual event because they are independent. We multiply the probability of the scenario in the interval (inside the braces) by the probability of the data at time $t$, which gives us the probability of the data at time $t + \Delta$ given that event $a$ occurred. We apply this logic to the subsequent events, though the relevant probabilities at time $t$ are different, depending on the implied number of unobserved lineages after the event occurs.

The probability for an event of type $b$ is:

$$p_u^b(t+\Delta) = 2s\lambda\Delta p_{u+1}(t)$$

because each of the $s$ sampled lineages could produce the unobserved lineage. The factor of two appears because either descendant of the split could be sampled.

The probability for an event of type $c$ is:

$$p_u^c(t+\Delta) = u\lambda\Delta p_{u+1}(t)$$

because each of the unsampled lineages at time $t + \Delta$ could produce a new lineage that is unsampled.

The probability for an event of type $d$ is:

$$p_u^d(t+\Delta) = u\mu\Delta p_{u-1}(t)$$

because each of the unsampled lineages at time $t + \Delta$ could go extinct in the interval.

The probability for an event of type $e$ is:

$$p_u^e(t + \Delta) = \frac{1}{2} su\eta \Delta p_u(t).$$

Since the unidirectional hybridization occurs between one sampled and one unsampled lineage, the rate is proporitonal to the number of such pairs, $su$. The factor $1/2$ appears because half of the events point into the unobserved lineage.

The probability for an event of type $f$ is:

$$p_u^f(t + \Delta) = \binom{u}{2} \eta \Delta p_u(t).$$

In this case, there are $\binom{u}{2}$ unobserved lineage pairs, each of which may undergo hybridization. The direction of the hybridization is irrelevant, so no factor of $1/2$ appears.

The probability for an event of type $g$ is:

$$p_u^g(t + \Delta) = \binom{u}{2} \zeta \Delta p_u(t),$$

which is the same as event $f$ except that it involves bidirectional hybridization.

The probability for an event of type $h$ is:

$$p_u^h(t + \Delta) = \binom{s}{2} \nu \Delta p_{u+1}(t)$$

because each pair of sampled lineages could give rise to the unsampled homoploid hybrid.

The probability for an event of type $i$ is:

$$p_u^i(t + \Delta) = \binom{u}{2} \nu \Delta p_{u+1}(t)$$

because each pair of unsampled lineages could give rise to the unsampled homoploid hybrid.

The probability for an event of type $j$ is:

$$p_u^j(t + \Delta) = su\nu \Delta p_{u+1}(t).$$

As with event $e$, the rate is proportional to the number of sampled and unobserved lineage pairs, $su$, that could give rise to the unsampled homoploid hybrid.

The events $k$, $l$, and $m$ are the same as $i$, $j$, and $k$, respectively, except that they involve allopolyloid events that occur at rate $\psi$.

Assuming that $\Delta \approx 0$ such that no more than one event can occur, the above events are mutually exclusive. Therefore, the probability of the data at time $t + \Delta$ is the sum of the above terms:

$$\begin{aligned} p_u(t + \Delta) = {} & p_u^a(t + \Delta) + p_u^b(t + \Delta) + p_u^c(t + \Delta) + p_u^d(t + \Delta) + p_u^e(t + \Delta) + \\ & p_u^f(t + \Delta) + p_u^g(t + \Delta) + p_u^h(t + \Delta) + p_u^i(t + \Delta) + p_u^j(t + \Delta) + \\ & p_u^k(t + \Delta) + p_u^l(t + \Delta) + p_u^m(t + \Delta) \end{aligned}$$

Substituting terms and simplifying,

$$p_u(t+\Delta) = \Bigg[(1-n\lambda\Delta)(1-n\mu\Delta)(1-\binom{n}{2}\eta\Delta)(1-\binom{n}{2}\zeta\Delta)(1-\binom{n}{2}\nu\Delta)(1-\binom{n}{2}\psi\Delta) + \frac{1}{2}su\eta\Delta + \binom{u}{2}\eta\Delta + \binom{u}{2}\zeta\Delta\Bigg]p_u(t) + \left[(u+2s)\lambda\Delta + \binom{n}{2}(\nu+\psi)\Delta\right]p_{u+1}(t) + u\mu\Delta p_{u-1}(t)$$

The corresponding finite difference equations are of the form

$$\frac{p_u(t+\Delta)-p_u(t)}{\Delta}$$

*Ordinary differential equations.*—We derive ODEs by taking the limit of the difference equations as $\Delta \to 0$:

$$\lim_{\Delta\to 0}\frac{p_u(t+\Delta)-p_u(t)}{\Delta} = \lim_{\Delta\to 0}\Bigg[(1-n\lambda\Delta)(1-n\mu\Delta)(1-\binom{n}{2}\eta\Delta)(1-\binom{n}{2}\zeta\Delta)(1-\binom{n}{2}\nu\Delta)(1-\binom{n}{2}\psi\Delta) + \frac{1}{2}su\eta\Delta + \binom{u}{2}\eta\Delta + \binom{u}{2}\zeta\Delta - 1\Bigg]\frac{p_u(t)}{\Delta} + \lim_{\Delta\to 0}\left[(u+2s)\lambda\Delta + \binom{n}{2}(\nu+\psi)\Delta\right]\frac{p_{u+1}(t)}{\Delta} + \lim_{\Delta\to 0}u\mu\Delta\frac{p_{u-1}(t)}{\Delta}$$

After some algebra, we arrive at the system of ODEs (over values of $u$) of the form:

$$\frac{\mathrm{d}p_u(t)}{\mathrm{d}t} = -\left[n(\lambda+\mu) + \frac{s(s-1)}{2}(\eta+\zeta) + su\left(\zeta+\frac{\eta}{2}\right) + \frac{n(n-1)}{2}(\nu+\psi)\right]p_u(t) + \left[(u+2s)\lambda + \frac{n(n-1)}{2}(\nu+\psi)\right]p_{u+1}(t) + u\mu p_{u-1}(t) \quad \text{(S.1)}$$

*Boundary states.*—The above equations consider the case where the number of unobserved lineages $u$ is not at or near a boundary (0 or $u_{\max}$). When it is at or near a boundary, some events are not possible and the ODEs must be slightly modified. For example, when $u = 0$, extinction events are impossible. Consequently,

$$\frac{\mathrm{d}p_0(t)}{\mathrm{d}t} = -\left[s(\lambda+\mu) + \frac{s(s-1)}{2}(\eta+\zeta) + \frac{s(s-1)}{2}(\nu+\psi)\right]p_0(t) + \left[2s\lambda + \frac{s(s-1)}{2}(\nu+\psi)\right]p_1(t)$$

At the upper boundary, transitions from $u_{\max}+1$ are omitted because this state is not represented. Events that would move probability from $u_{\max}$ to $u_{\max}+1$ are absorbed into the implicit overflow state and therefore appear only through the negative diagonal term. Consequently, the ODE for $u = u_{\max}$ has the same diagonal and lower-neighbor terms as above, but no contribution from $p_{u_{\max}+1}(t)$.

*Matrix representation.*—The system of ODEs can be represented more compactly by a matrix of coefficients, $Q(s)$, with elements $q_{i,j}(s)$:

$$
\begin{aligned}
q_{u,u-1}(s) &= u\mu && \text{for } u > 0 \\
q_{u,u}(s) &= -\left[ n(\lambda+\mu) + \binom{s}{2}(\eta+\zeta) + su\left(\zeta + \frac{\eta}{2}\right) + \binom{n}{2}(\nu+\psi) \right] && \text{for all } u \\
q_{u,u+1}(s) &= (u+2s)\lambda + \binom{n}{2}(\nu+\psi) && \text{for } u < u_{\max}
\end{aligned}
$$

In this representation, $j$ is the source state and $i$ is the destination state, so $q_{i,j}(s)$ is the coefficient multiplying $p_j(t)$ in the differential equation for $p_i(t)$ (*i.e.*, the matrix is tranposed compared to standard phylogenetic rate matrices). Additionally, the columns of this matrix do not sum to zero. This allows probability to flow out of the vector $p_u(t)$, owing to events that increase $u$ beyond $u_{\max}$ and events that cannot occur without producing observed events in the sampled network (*e.g.*, reticulation events between sampled lineages).

*Computing the transition probability between node events.*—The transition probability matrix that describes the probability of transition from state $\{s, i\}$ to state $\{s, j\}$ over an interval of length $t$ is

$$P(s,t) = \mathrm{e}^{Q(s)t}$$

Recall that the (column) vector of probabilities for all values of $u$ is $\vec{p}(t)$. Given the probability vector at time $\tau_0$, $\vec{p}(\tau_0)$, the probability vector at time $\tau_1$ (further in the past) is:

$$\vec{p}(\tau_1) = P(s, \tau_1 - \tau_0)\vec{p}(\tau_0)$$

The matrix exponential averages over all possible trajectories of $u$ over the interval.

To solve this exponential, we implemented numerical ODE solvers as well as matrix exponentiation using uniformization. Uniformization computes the matrix exponential as:

$$P(t) = \mathrm{e}^{Qt} = \sum_{k=0}^{\infty} \frac{(\gamma t)^k \mathrm{e}^{-\gamma t}}{k!} A^k$$

where $k$ is a Poisson-distributed random variable with rate $\gamma$, $\gamma$ is no smaller in magnitude than any entry of $Q$ (in practice we set $\gamma$ to the largest value of $-Q_{ii}$), and

$$A = I + \frac{Q}{\gamma}$$

is a single-step transition probability matrix for the uniformized process ($I$ is the identity matrix). In practice, we enumerate this sum by brute force, but truncate the largest value of $k$ to a finite number $K$, such that the probability mass for a number of events larger than $K$ is negligible ($\approx 1e - 15$ by default). In experiments (not shown), uniformization proved to be more stable and significantly faster than the adaptive ODE solvers available in `Boost`.

#### S1.2 Node-event densities

*Transition densities for node events.*—Here we describe each possible node event and the corresponding transition density. We denote the time of the event as $t$, and the time immediately before the event as $t^-$. Each of the events occurs with some probability density, and affects the number of unsampled lineages in the process.

A type $A$ event is a speciation event: one ancestral lineage splits into two descendant lineages, neither of which goes extinct or unsampled. The probability density for an event of type $A$ is:

$$p_u^A(t) = \lambda p_u(t^-)$$

A type $B$ event is a undirectional hybridization event where neither parent goes extinct or unsampled. The probability density after an event of type $B$ is:

$$p_u^B(t) = \frac{\eta}{2} p_u(t^-)$$

The factor 1/2 occurs because half of the unidirectional hybridization events point in a particular direction.

A type $C$ event is a unidirectional hybridization event where one of the parents goes extinct or unsampled. The probability density after an event of type $C$ is:

$$p_u^C(t) = \eta p_{u+1}(t^-)$$

This is twice the rate of a type $B$ event because either donor could have gone extinct or unsampled.

A type $D$ event is a bidirectional hybridization event where neither lineage goes extinct or unsampled. The probability density after an event of type $D$ is:

$$p_u^D(t) = \zeta p_u(t^-)$$

A type $D$ event is a bidirectional hybridization event where one of the lineages goes extinct or unsampled. The probability density after an event of type $E$ is:

$$p_u^E(t) = 2\zeta p_{u+1}(t^-)$$

The factor two arises because either parental lineage could go extinct or unsampled to produce the observed event.

A type $F$ event is a homoploid hybrid speciation event where neither parent goes extinct or unsampled. The probability density after an event of type $F$ is:

$$p_u^F(t) = \nu p_u(t^-)$$

A type $G$ event is a homoploid hybrid speciation event where one parent goes extinct or unsampled. The probability density after an event of type $G$ is:

$$p_u^G(t) = \nu p_{u+1}(t^-)$$

Note that, even though one of the parents is extinct or unsampled, we do not multiply by two because the direction of the reticulation implies which parent went extinct (if the other parent went extinct, the arrow would point in the opposite direction).

A type $H$ event is a homoploid hybrid speciation event where both parents go extinct or unsampled. The probability density after an event of type $H$ is:

$$p_u^H(t) = \nu p_{u+2}(t^-)$$

A type $I$ event is a homoploid hybrid speciation event where neither parent goes extinct or unsampled. The probability density after an event of type $I$ is:

$$p_u^I(t) = \psi p_u(t^-)$$

A type $J$ event is a homoploid hybrid speciation event where one parent goes extinct or unsampled. The probability density after an event of type $J$ is:

$$p_u^J(t) = \psi p_{u+1}(t^-)$$

We do not multiply by two for the same reason given for type $G$ events.

A type $K$ event is a homoploid hybrid speciation event where both parents go extinct or unsampled. The probability density after an event of type $K$ is:

$$p_u^K(t) = \psi p_{u+2}(t^-)$$

*Transition densities for node event types.*—Some node events are ambiguous in the sampled network (they involve the same number of ancestors and descendants), so we must sum over all types of events that are consistent with a given node. We categories each node event into one of eight types, *i* through *vii*.

The probability density after a type *i* configuration is:

$$p_u^i(t) = p_u^A(t)$$

The probability density after a type *ii* configuration is:

$$p_u^{ii}(t) = p_u^B(t) + p_u^G(t)$$

because it could be either a type $B$ event or a type $G$ event. Likewise, the probability density after a type *iii* configuration is

$$p_u^{iii}(t) = p_u^C(t) + p_u^E(t) + p_u^H(t).$$

The probability density after a type *iv* configuration is:

$$p_u^{iv}(t) = p_u^D(t)$$

The probability density after a type *v* configuration is:

$$p_u^v(t) = p_u^F(t)$$

The probability density after a type *vi* configuration is:

$$p_u^{vi}(t) = p_u^I(t)$$

The probability density after a type *vii* configuration is:

$$p_u^{vii}(t) = p_u^J(t)$$

The probability density after a type *viii* configuration is:

$$p_u^{viii}(t) = p_u^K(t)$$

*Matrix representation.*—These transition densities can be expressed as a matrix that operates on $p_u(t^-)$ to compute $p_u(t)$. This requires care near the boundary $u_{\max}$, which prohibits some kinds of transition events. The matrices for each type of event are:

$$R^{i} = \begin{cases} r^{i}_{u,u} = \lambda & \text{for } u \leq u_{\max} \\ 0 & \text{otherwise} \end{cases}$$

$$R^{ii} = \begin{cases} r^{ii}_{u,u} = \frac{\eta}{2} & \text{for } u \leq u_{\max} \\ r^{ii}_{u,u+1} = \nu & \text{for } u < u_{\max} \\ 0 & \text{otherwise} \end{cases}$$

$$R^{iii} = \begin{cases} r^{iii}_{u,u+1} = \eta + 2\zeta & \text{for } u < u_{\max} \\ r^{iii}_{u,u+2} = \nu & \text{for } u < u_{\max} - 1 \\ 0 & \text{otherwise} \end{cases}$$

$$R^{iv} = \begin{cases} r^{iv}_{u,u} = \zeta & \text{for } u \leq u_{\max} \\ 0 & \text{otherwise} \end{cases}$$

$$R^{v} = \begin{cases} r^{v}_{u,u} = \nu & \text{for } u \leq u_{\max} \\ 0 & \text{otherwise} \end{cases}$$

$$R^{vi} = \begin{cases} r^{vi}_{u,u} = \psi & \text{for } u \leq u_{\max} \\ 0 & \text{otherwise} \end{cases}$$

$$R^{vii} = \begin{cases} r^{vii}_{u,u+1} = \psi & \text{for } u < u_{\max} \\ 0 & \text{otherwise} \end{cases}$$

$$R^{viii} = \begin{cases} r^{viii}_{u,u+2} = \psi & \text{for } u < u_{\max} - 1 \\ 0 & \text{otherwise} \end{cases}$$

### S2 Validation

As described in the main text, we validated our network probability density by summing over network topologies and integrating over node ages for small networks, and comparing these values against theoretically matching Monte Carlo frequencies. Here, we provide further numerical details of this procedure, including how parameters were selected, how topologies were enumerated, and how we performed high dimensional numerical integration over the node ages.

#### S2.1 Parameter tuning

For each scenario described in the main text, we tuned parameters so the expected number of tips and reticulation events was close to the target number. We note that the exact details of this tuning are somewhat irrelevant, since for given parameters the marginalized network density and the corresponding Monte Carlo estimate should agree regardless of how the parameters are chosen. However, Monte Carlo estimates are very unstable when the quantity being approximated is very small, so we developed this tuning procedure to increase the efficiency of the Monte Carlo approximation.

For these simulations, we set the age of the process to $T = 1$, the relative extinction rate to $\epsilon = \mu/\lambda = 0.1$, and the sampling fraction to $\rho = 0.2$. We then specify a Markovian rate matrix defined by the model parameters, $\lambda$, $\mu$, $\rho$, and $\theta$. The state of this model is the joint number of non-reticulate and reticulate lineages, $\{d, h\}$. In this model, we define a reticulate lineage as one that has any reticulation event in its ancestry. For $h = 1$, this should be fairly similar to the number of reticulation events $r$, though there is some chance that a reticulate lineage has more than one reticulation event in its history. For the allopolyploidy model (with reticulation parameter $\psi$), the instantaneous rate matrix has elements:

$$
\begin{aligned}
q_{\{i,j\}\to\{i-1,j\}} &= \mu i && \text{death of a non-reticulate lineage} \\
q_{\{i,j\}\to\{i+1,j\}} &= \lambda i && \text{birth of a non-reticulate lineage} \\
q_{\{i,j\}\to\{i,j-1\}} &= \mu j && \text{death of a reticulate lineage} \\
q_{\{i,j\}\to\{i,j+1\}} &= \lambda j + \binom{i+j}{2}\psi && \text{birth of a reticulate lineage}
\end{aligned}
$$

with the size of the matrix set to a large truncation value $K$. Using this finite matrix, the distribution of the number of tips at the present (before sampling) is approximately:

$$\pi(0) = \mathrm{e}^{QT}\pi(T),$$

where $\pi(T)$ is a vector of initial probabilities (*i.e.*, $\pi_{\{1,0\}}(T) = 1$ and 0 otherwise, because we began with one non-reticulate and zero reticulate lineages). The marginal probability of $\{d, h\}$ among the

**Table S1**

| Scenario | $\{n, h\}$ | $\lambda$ | $\theta$ | $P(n, h)$ | $\lvert\mathcal{N}\rvert$ |
|---|---|---|---|---|---|
| 1 | $\{2, 1\}$ | 2.4305 | $\eta = 0.5000$ | 0.0524 | 4 |
| 2 | $\{2, 1\}$ | 2.4305 | $\zeta = 0.2500$ | 0.0409 | 4 |
| 3 | $\{3, 1\}$ | 2.4305 | $\nu = 0.2000$ | 0.0242 | 10 |
| 4 | $\{3, 1\}$ | 2.7081 | $\eta = 0.1000$ | 0.0281 | 9 |
| | | | $\zeta = 0.2000$ | | |
| | | | $\nu = 0.0333$ | | |
| 5 | $\{3, 1\}$ | 2.4305 | $\psi = 0.2000$ | 0.0241 | 10 |

sampled lineages is then a sum:

$$P(\{d,h\}) = \sum_{\{x,y\}} \left[\rho^d (1-\rho)^{x-d} \rho^h (1-\rho)^{y-h}\right] \pi_{\{x,y\}}(0)$$

For each scenario with target numbers $n$ and $r$, we set $d = n - r$ and $h = r$, and then performed a grid search over values of $\lambda$ and $\theta$ to find the values that maximized the probability of $d$, $r$. (For scenario 4, we fixed the ratios of the three hybridization parameters and searched over values of their sum.) The resulting tuned parameters for each model are presented in Table S1.

##### S2.2 Enumerating the network topologies

We manually enumerated each network topology with the target number of tips and reticulation events. For a given scenario, we denote the set of such topologies $\mathcal{N}$, and the number of such topologies $|\mathcal{N}|$. The target numbers and total number of topologies for each scenario are listed in Table S1. These topologies and the number of orientations of each are depicted in Figures S2 through S6. Note that we ignored the single topology from scenario 4 that could only be produced by homoploid hybrid speciation (the first panel from scenario 3).

##### S2.3 Monte Carlo simulation

We developed a Monte Carlo simulator for each validation scenario. We forward simulated under the corresponding model and parameters, starting with a single lineage at time $t = T$ until time $t = 0$. We then applied sampling according to the assumed parameter $\rho$, and pruned out extinct and unsampled lineages. We repeat this 10 million times, and classified each replicate as either "off-target" (*i.e.*, it had the wrong number of tips and reticulation events) or as belonging to one of the enumerated topologies for the corresponding scenario. A replicate was treated "on-target" as long as it was classified as one of the enumerated topologies. We then computed the frequency of on-target simulations, which is a Monte Carlo estimate of $P(n, r \mid \theta)$.

##### S2.4 Numerical integration

For each topology, we integrated over all possible sets of node ages for that topology using multidimensional numerical integration available the `Julia` package `cubature` (Johnson 2013). Beyond the relatively high dimensionality (up to four dimensions/ages in some cases), the main technical challenge is that node ages are constrained by each other. We used constrained numerical integration to ensure that the domains of integration for each $\tau$ correctly obeyed the nested relationships of each node during the integration (essentially ensuring that all branch lengths were positive, and all hybridization edges were non-negative).

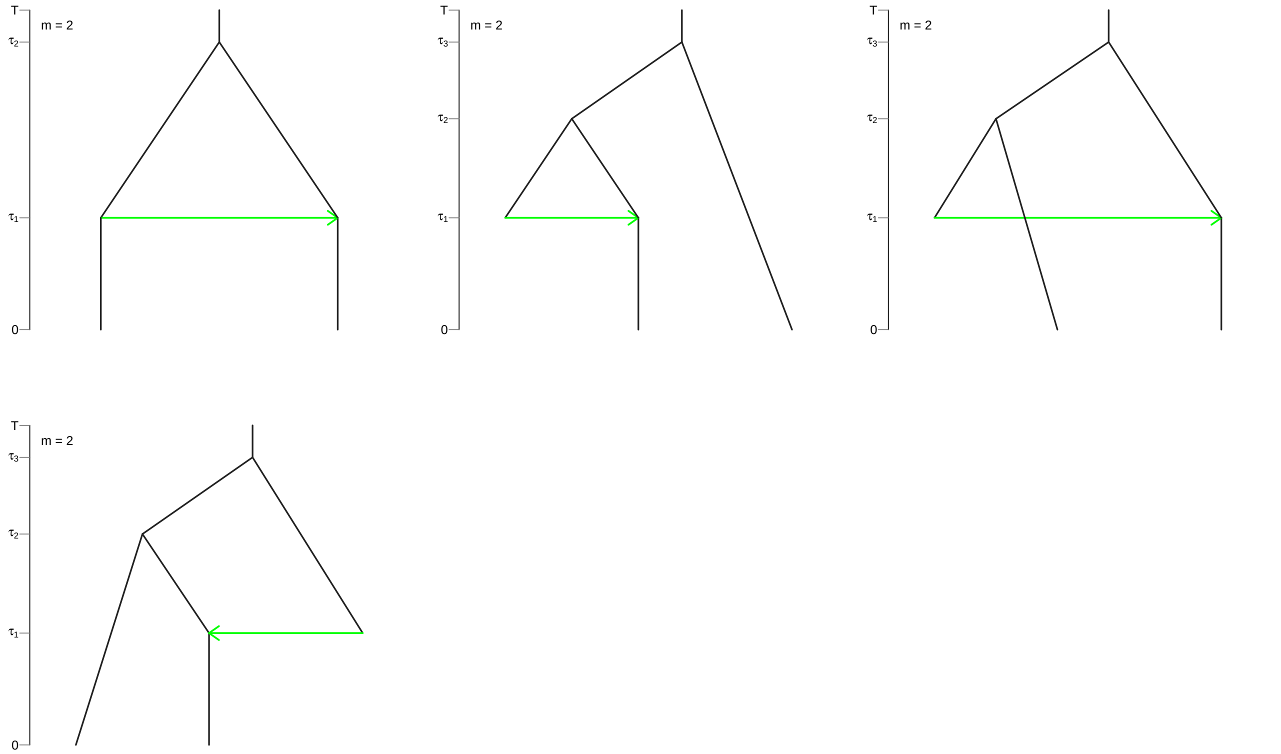


**Figure S2: Network topologies for scenario 1.** Reticulation events are depicted as green horizontal lines. $m$ is the number of orientations of the topology.

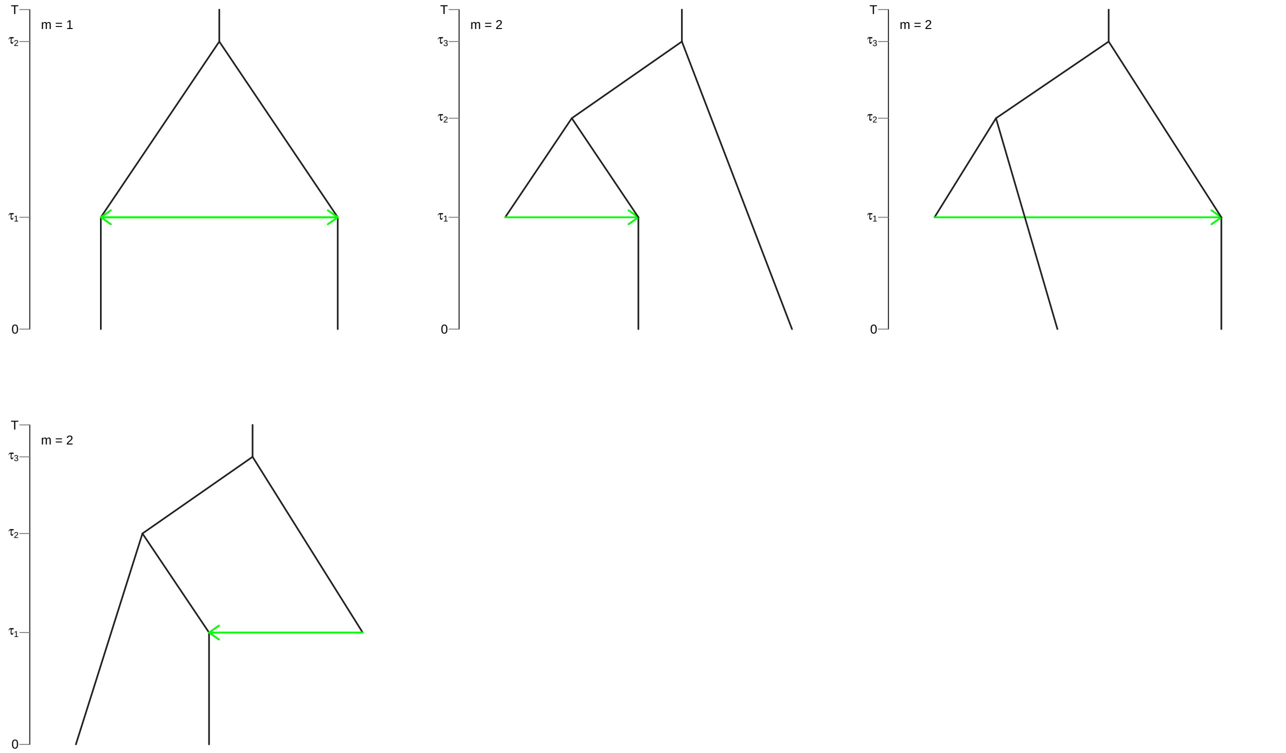


**Figure S3: Network topologies for scenario 2.** Reticulation events are depicted as green horizontal lines. $m$ is the number of orientations of the topology.

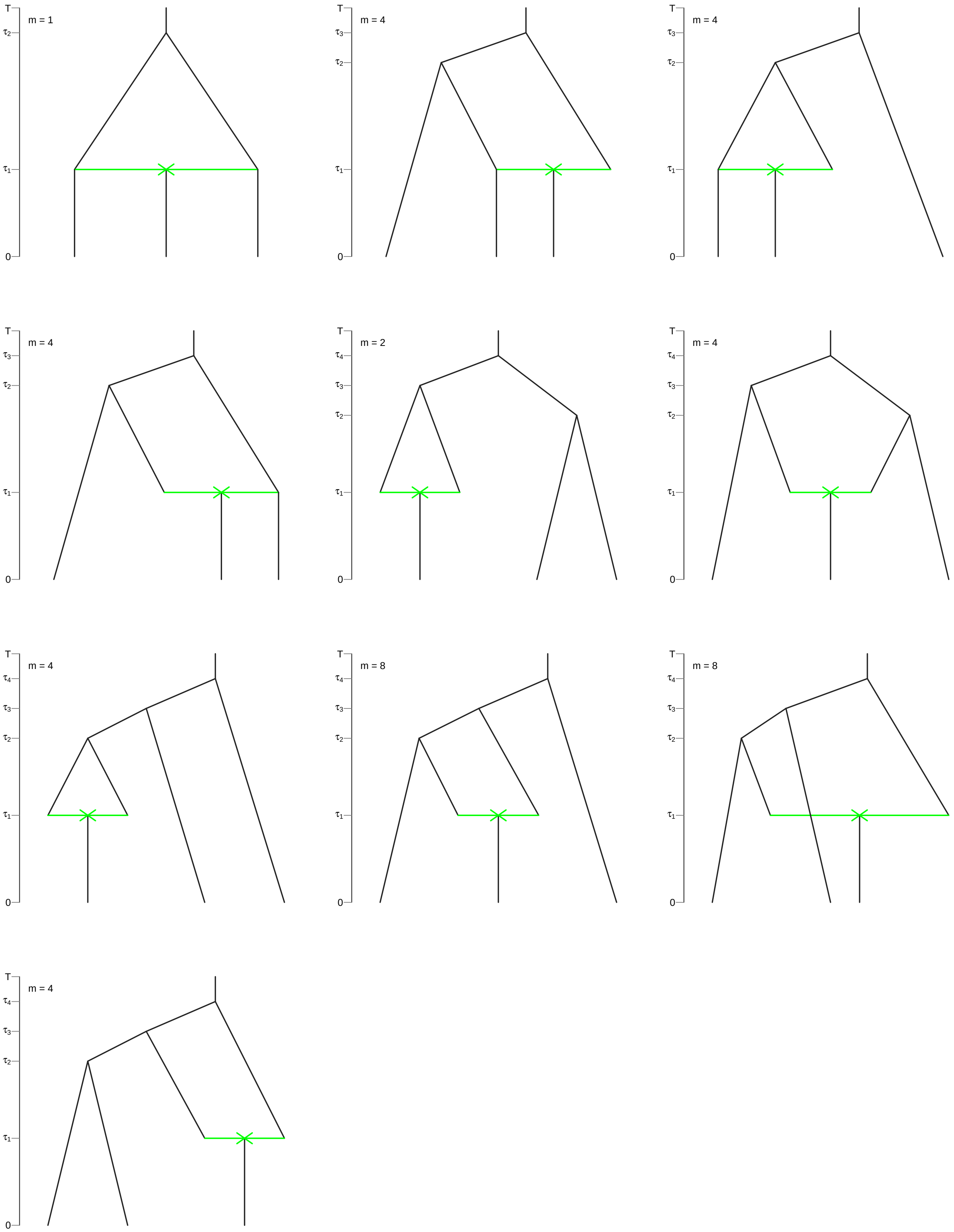


**Figure S4: Network topologies for scenario 3.** Reticulation events are depicted as green horizontal lines. $m$ is the number of orientations of the topology.

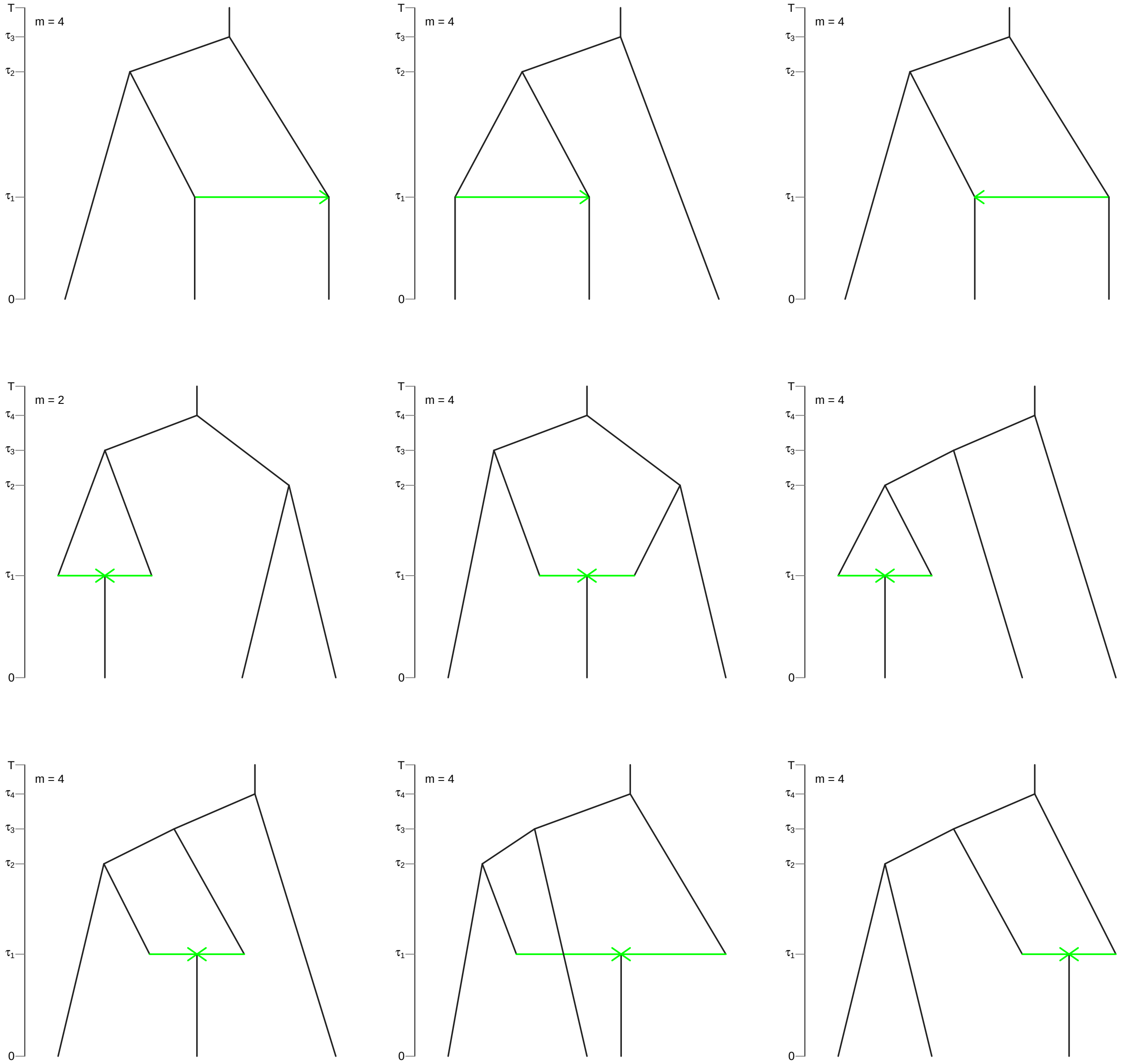


**Figure S5: Network topologies for scenario 4.** Reticulation events are depicted as green horizontal lines. $m$ is the number of orientations of the topology.

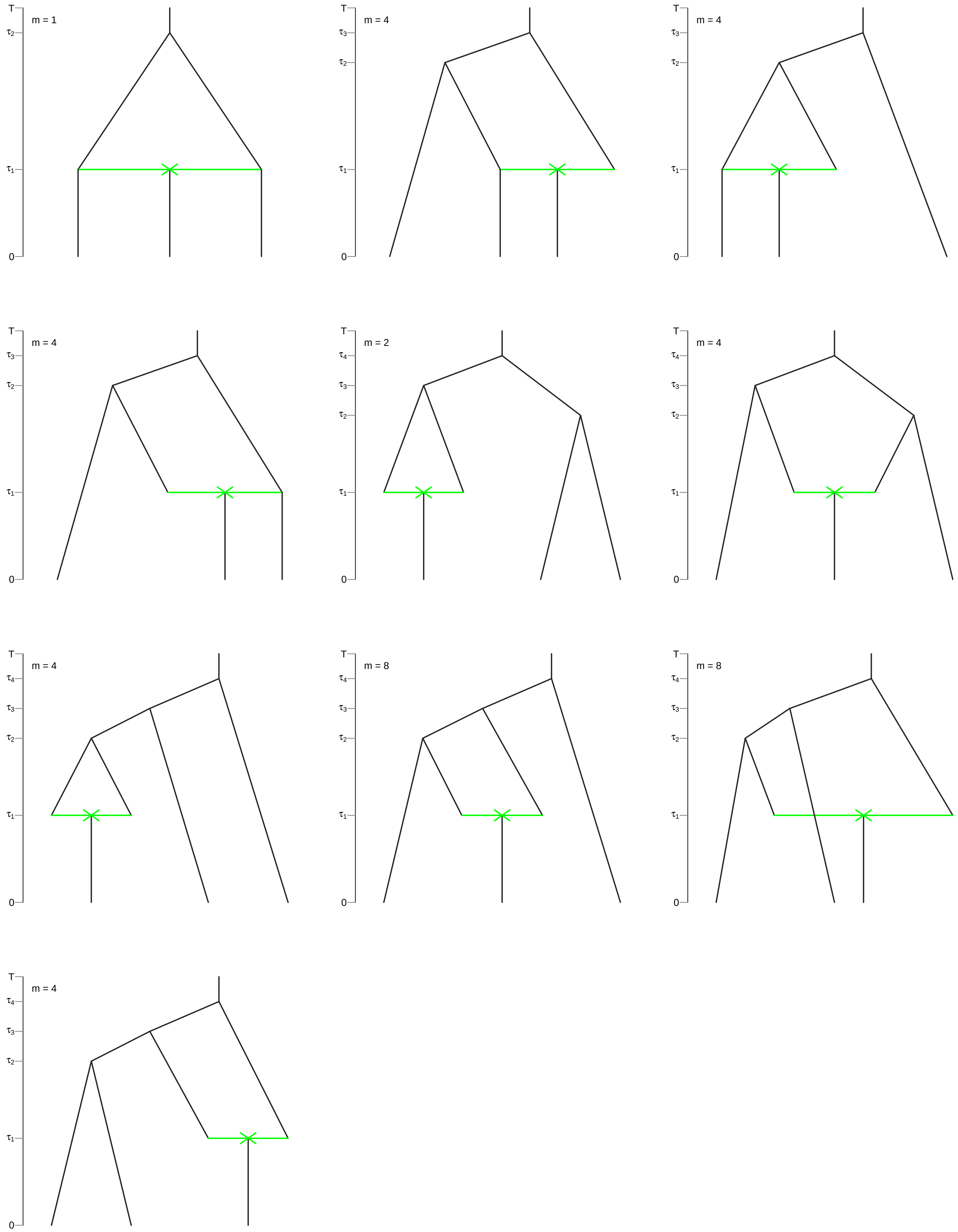


**Figure S6: Network topologies for scenario 5.** Reticulation events are depicted as green horizontal lines. *m* is the number of orientations of the topology.

### S3 Simulation Study

#### S3.1 Simulation procedure

We simulated phylogenetic networks over the scenario and parameter set described in the main text. For each parameter and scenario combination, we tuned the root time $T$ of the network such that the expected number of tips was 60. To achieve this, we chose a particular value of $T$, then simulated many networks under the given parameters, and computed the average number of simulated tips. We increased or decreased $T$ until the expected value converged to $\approx$60, within some tolerance. We chose the initial value of $T$ using the Markov model framework described in the validation section, then used a hill climber to refine $T$ until the expected number of tips (based on 20,000 Monte Carlo replicates) was $60 \pm 2$.

Given the chosen value of $T$, we simulated networks of fixed size $N \in \{16, 32, 64, 128\}$ using rejection sampling. Specifically, we simulated networks, rejecting those that had tip numbers that did not match the target. We repeated this until we collected 100 replicates with the target $N$. Across parameter settings and sizes, the resulting networks covered a broad range of number of reticulation events, shown in Figures S7, S8, and S9.

#### S3.2 MCMC analyses

We used the `Julia` package `AdaptiveMCMC` (Vihola 2026) to estimate the joint posterior distribution of model parameters, as described in the main text. Here, we note some additional details of those analyses. First, we used the adaptive scaling within adaptive Metropolis (ASWAM) algorithm implemented in `AdaptiveMCMC`, which adaptively learns a covariance matrix for a multivariate normal proposal on all parameters. We allowed the proposal to adapt only during the burnin phase of the chain (5,000 iterations, or 10% of the entire run), and used a target acceptance rate of 23%.

*Parameter transformations.*—In preliminary analyses, it proved difficult to achieve good mixing on untransformed parameters, so we transformed the parameter space while keeping the priors the same. Specifically, for the relatively extinction $\epsilon$ with lower bound $l = 0$ and upper bound $u = 1$, we applied a logit transform (and its inverse):

$$x = \text{logit}\left(\frac{\epsilon - l}{u - l}\right) = \text{logit}(\epsilon) = \log\left(\frac{\epsilon}{1-\epsilon}\right)$$
$$\epsilon = l + \frac{u - l}{1 + e^{-z}} = \frac{1}{1 + e^{-z}}$$

The corresponding Jacobian ratio (applied to the Metropolis-Hastings acceptance ratio) is

$$\frac{J(\epsilon')}{J(\epsilon)} = \frac{\epsilon'(1-\epsilon')}{\epsilon(1-\epsilon)}$$

For the speciation and reticulation rates (which had loguniform priors), we first log transformed, then applied the same logit transform, so that (for generic parameter $\theta$), the relevant quantities are:

$$x = \operatorname{logit}\left(\frac{\log\theta - \log l}{\log u - \log l}\right)$$
$$z = \exp\left[\log l + \frac{\log u - \log l}{1 + e^{-z}}\right]$$
$$\frac{J(\theta')}{J(\theta)} = \frac{\theta'}{\theta}\frac{\frac{e^{-z'}}{(1+e^{-z'})^2}}{\frac{e^{-z}}{(1+e^{-z})^2}}$$

The `AdaptiveMCMC` package does not natively support constrained transformations. Therefore, we designed the sampler to sample transformed parameters, and applied the necessary transformations inside our likelihood and prior functions. To apply the Jacobian terms, we added the (log) numerator of the Jacobian ratios to the proposed log prior density and the (log) denominator to the current log prior density to satisfy the Metropolis-Hastings acceptance rate:

$$A = \min\left[1, \frac{P(\aleph \mid \theta')}{P(\aleph \mid \theta)}\frac{P(\theta')}{P(\theta)}\frac{J(\theta')}{J(\theta)}\right]$$

We then transformed the samples post hoc to the original parameter space.

#### S3.3 Extended results

We report the posterior estimates of all model parameters across simulation settings for $u_{\max} = 4N$. These figures follow the same format as the main simulation results figure in the main text. See Figures S17–S97 at the end of this document. Additionally, we report tables of RMSE (relative posterior root-mean squared error) averaged across datasets, and $\text{RMSE}_c$ (averaged over datasets with at least one reticulation event), and coefficients of variation, for all simulation scenarios, parameter combinations, and $u_{\max}$ values in Tables S2–S10.

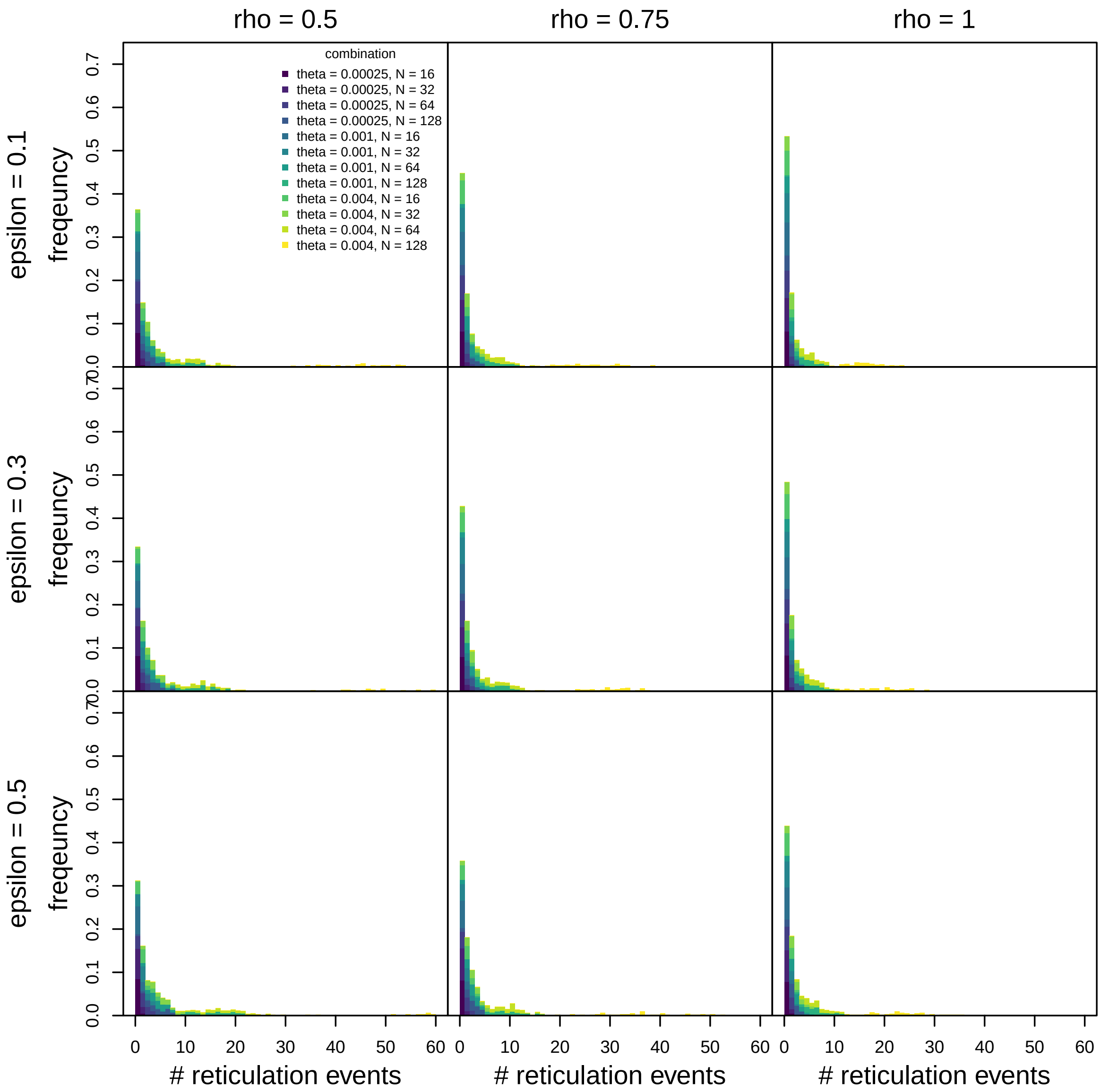


**Figure S7: Distribution of the number of reticulation events in simulated datasets for the unidirectional hybridization simulations.** Values of $\epsilon$ are in rows and values of $\rho$ are in columns. Within panels, each bar represents the frequency of simulations (y axis) with the given number of reticulation events (x axis). Each bar is color-coded according to the frequency of those simulations that come from each parameter setting, as described in the legend.

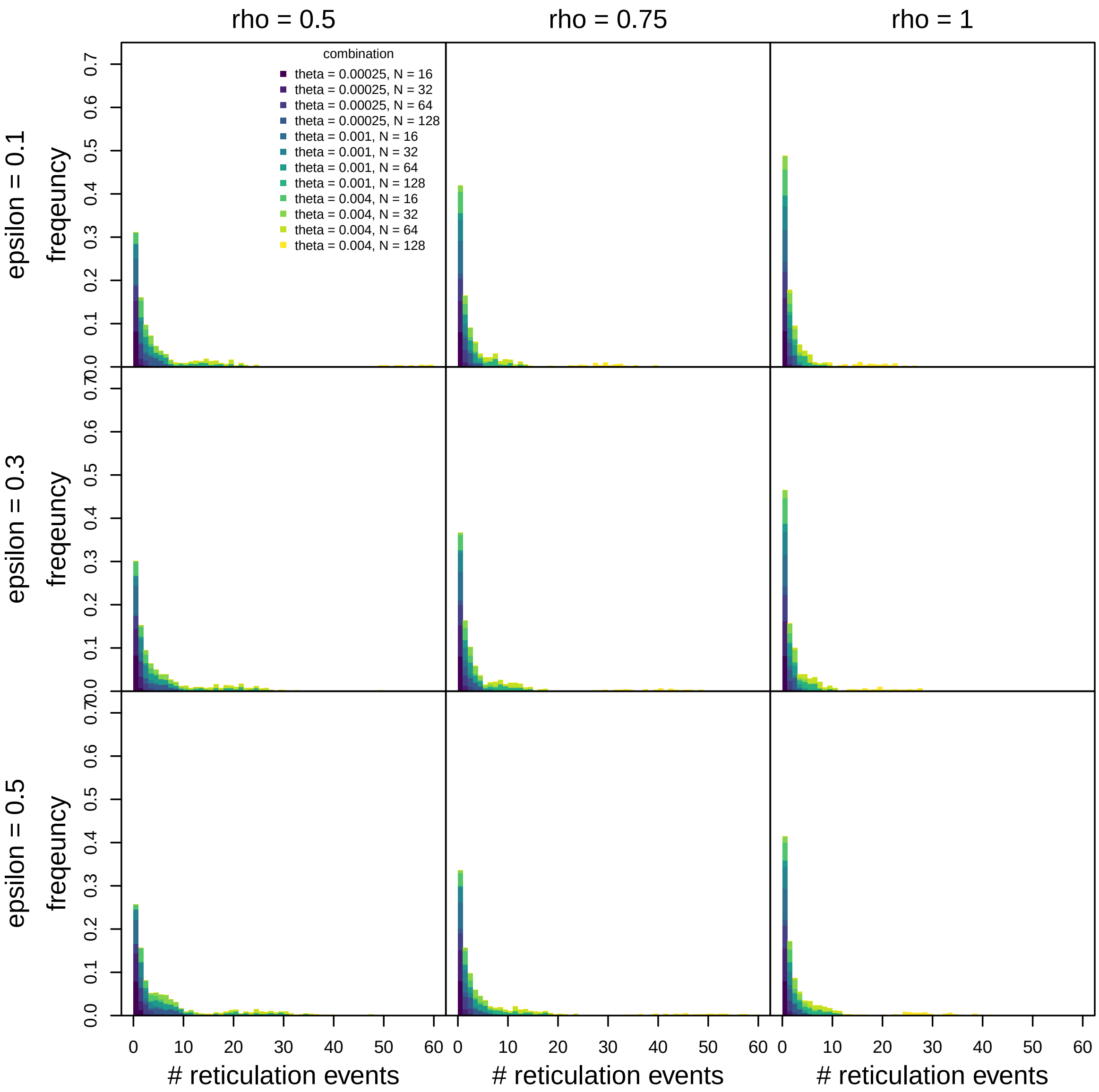


**Figure S8: Distribution of the number of reticulation events in simulated datasets for the bidirectional hybridization simulations.** Values of $\epsilon$ are in rows and values of $\rho$ are in columns. Within panels, each bar represents the frequency of simulations (y axis) with the given number of reticulation events (x axis). Each bar is color-coded according to the frequency of those simulations that come from each parameter setting, as described in the legend.

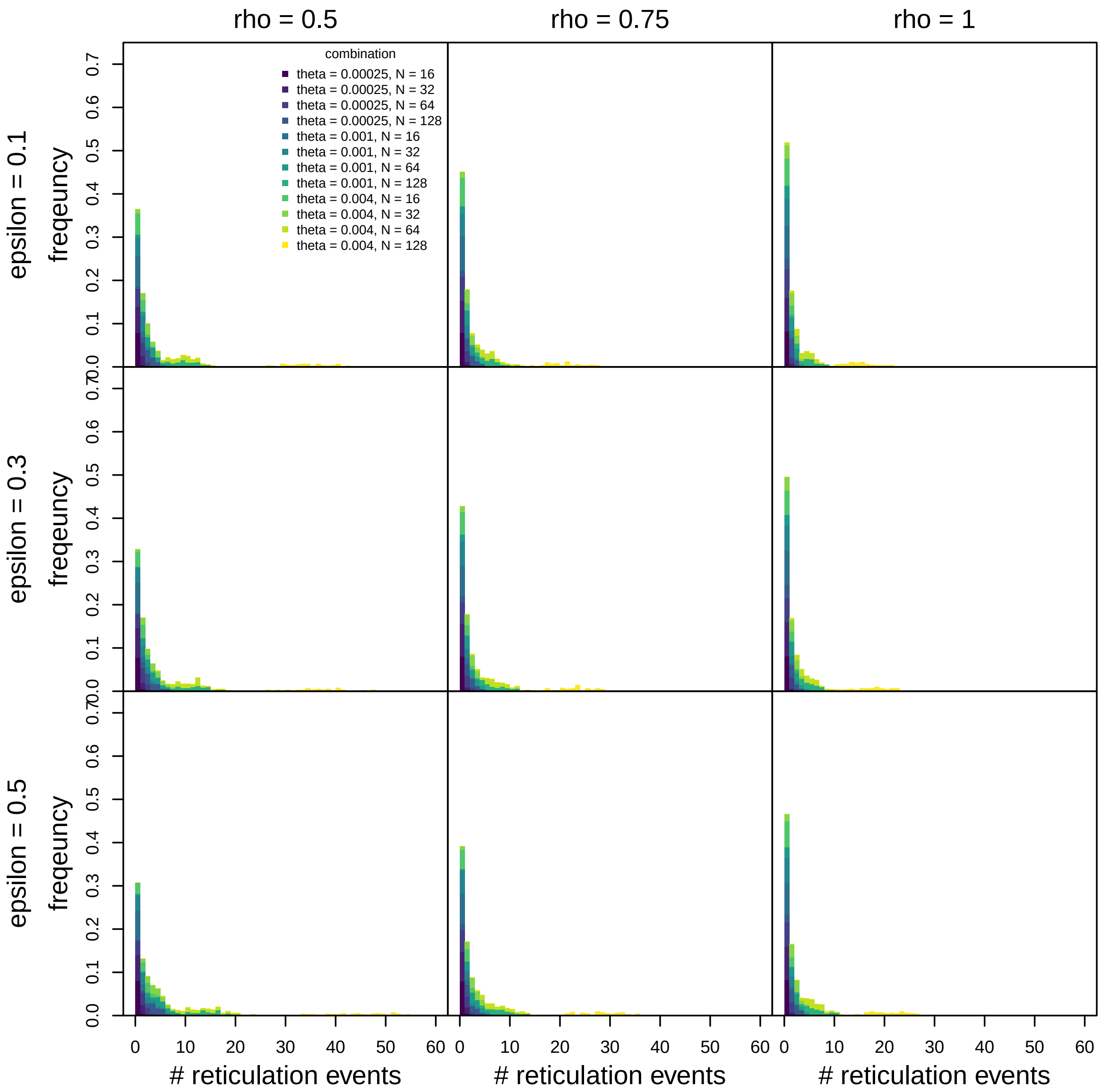


**Figure S9: Distribution of the number of reticulation events in simulated datasets for the allopolyploid speciation simulations.** Values of $\epsilon$ are in rows and values of $\rho$ are in columns. Within panels, each bar represents the frequency of simulations (y axis) with the given number of reticulation events (x axis). Each bar is color-coded according to the frequency of those simulations that come from each parameter setting, as described in the legend.

### S4 Empirical analyses

#### S4.1 Bayesian augmented network analysis

##### *S4.1.1 Theory*

We performed Bayesian analysis under the augmented network model described in the main text. This model has posterior distribution

$$P(\vec{\tau}^*, \lambda, \epsilon, \psi \mid \aleph, \rho) \propto P(\aleph^*, \vec{\tau}^* \mid \lambda, \epsilon, \psi, \rho) P(\lambda) P(\epsilon) P(\psi)$$

where $\vec{\tau}^*$ is a set of augmented reticulation ages, $\aleph^*$ is the augmented network (the network $\aleph$ with the reticulation ages replaced by $\vec{\tau}^*$), and $P(\aleph^*, \vec{\tau}^* \mid \lambda, \epsilon, \psi, \rho)$ is

$$P(\aleph^*, \vec{\tau}^* \mid \lambda, \epsilon, \psi, \rho) = P(\aleph^* \mid \lambda, \epsilon, \psi, \rho) \mathbb{1}(\vec{\tau}^*)$$

where $\mathbb{1}(\vec{\tau}^*)$ is an indicator function that checks whether the augmented ages are consistent with the network $\aleph$

$$\mathbb{1}(\vec{\tau}^*) = \begin{cases} 1 & \text{if } \vec{\tau}^* \text{ is consistent with } \aleph \\ 0 & \text{otherwise} \end{cases}$$

By consistent we mean they are no older than the ages of those events in the network $\aleph$.

The justification for this approach is as follows. While we can compute the probability density of a fully specified network (*i.e.*, with exact node ages) under our birth-death-reticulation model, we don't actually know the exact ages. We consider the ages of reticulation events in the empirical network $\aleph$ to be partial information: the MUL tree from which they are derived provides only an upper bound on the ages of reticulation events. In reality, a given reticulation event could be anywhere between the age in $\aleph$ and their oldest immediate descendant (or the present, in the case they have no descendants, which is true for all allopolyploids in our network). In other words, the exact ages are missing data. The probability density of the network (with partial infromation) should therefore integrate over all possible sets of reticulation ages. Denoting the set of all possible ages $\mathcal{T}$, this marginal density is

$$P(\aleph) = \int\limits_{\vec{\tau}^* \in \mathcal{T}} P(\aleph^* \mid \lambda, \epsilon, \psi, \rho) \mathrm{d}\vec{\tau}^*$$

where again $\aleph^*$ is just the observed network $\aleph$ with the ages $\vec{\tau}^*$ substituted into the reticulation ages, and the integral represents a multidimensional integral over the set of valid ages $\mathcal{T}$.

Rather than doing this integral calculation during MCMC (which would be extremely onerous), we simply sample the values of $\vec{\tau}^*$ subject to the constraint that they must obey the observed network $\aleph$. This should be equivalent to the above integration because

$$P(\lambda, \epsilon, \psi \mid \aleph, \rho) = \int_{\vec{\tau}^*} P(\vec{\tau}^*, \lambda, \epsilon, \psi \mid \aleph, \rho) \mathrm{d}\vec{\tau}^* \propto \left[ \int\limits_{\vec{\tau}^*} P(\aleph^* \mid \lambda, \epsilon, \psi, \rho) \mathrm{d}\vec{\tau}^* \right] P(\lambda) P(\epsilon) P(\psi)$$

*i.e.*, the marginalized posterior (left) is the same as the augmented posterior with $\vec{\tau}^*$ marginalized after the fact.

##### S4.1.2 Implementation

We designed a custom Metropolis-Hastings MCMC sampler to sample the joint posterior distribution of the model parameters and reticulation ages. For the model parameters, we applied the same parameter transformations described in the simulation study, above. We then used the ASWAM machinery from the `AdaptiveMCMC` package to make adaptive multivariate proposals to the transformed parameters. We also implemented reticulation-age proposals (described below) to sample the ages of each reticulation event. For each MCMC generation, our sampler makes one proposal to the continuous parameters (using ASWAM) and one proposal to one reticulation age in sequence, each of which is accepted/rejected independently according to the MH algorithm.

We also designed Metropolis-Hastings proposals for the ages of the reticulation events. These proposals are a mix of continuous-variable proposals and discrete proposals that propose to change a reticulation age to become exactly equal to (or unequal to) its maximum age. The type of proposals that get applied to a particular event depend on the configurations of allopolyploidization events that are possible for that node. There are three types of reticulation events, classified by the lengths of the reticulation edges leading into the event. The classification of reticulation edges by type for the Cystopteridaceae network from Rothfels et al. (2017) is depicted in Figure S10.

Type I nodes have two reticulation edges that are of length zero in the original network. This means that both parents are of identical age, $t$. If the reticulation event occured sometime after $t$, then implicitly there are two ghost lineages which diverged from their respective ancestors at exactly the same moment. This implies that two speciation events occurred at exactly time $t$, which has probability density zero under our Markovian birth-death-reticulation model, which allows at most one event to occur in an infinitesimal amount of time. Therefore, Type I nodes *must* have two surviving parental lineages, and therefore the age of the reticulation event has age $t$ with posterior probability 1. Accordingly, we do not apply any proposals to the ages of Type I nodes.

On the other end of the spectrum, Type III nodes have two reticulation edges that are of positive length in the original network. In our network, Type III nodes are almost all allopolyploid "bubbles", where both reticulation edges descend directly from a common ancestor at time $t$. The maximum age of the reticulation event is necessarily $t$. However, the reticulation event can not have occurred exactly at time $t$, since $t$ is when the two lineages that gave rise to the polyploid diverged. In other words, we know a speciation event occured at time $t$, therefore an allopolyploidization event could not also have occurred at time $t$, under the assumptions of the model. Accordingly we apply only continuous proposals to the ages of Type III nodes, *i.e.*, we never consider the possibility that they have age exactly $t$.

Type II nodes are more complicated. They have parent nodes $A$ and $B$ with ages $t_A$ and $t_B$, respectively. We order the parents so that $t_B > t_A$. The reticulation event can be no older than either of its parents. However, two configurations are possible: either the reticulation event occurred at exactly time $t_A$, in which case the reticulation event has one ghost parent leading to ancestor $B$, or the reticulation event occured after $t_A$, in which case it has two ghost parents leading to $A$ and $B$. In contrast to Type I nodes, the invocation of ghost lineages here does not imply the existence of simultaneous events under the model, and so we must consider both cases. The possible ages for the reticulation event, $\tau_i$, are therefore $0 < \tau_i \leq t_A$. We therefore use continuous proposals on the age of the node (when $\tau_i \neq t_A$), as well as proposals that change states between $\tau_i = t_A$ and $\tau_i \neq t_A$. If the state is currently $\tau_i = t_A$, we apply a discrete age proposal to $\tau_i \neq t_A$ with probability 1; if $\tau_i \neq t_A$ then we apply a continuous proposal with probability 0.5, otherwise we apply a discrete proposal to state $\tau_i = t_A$.

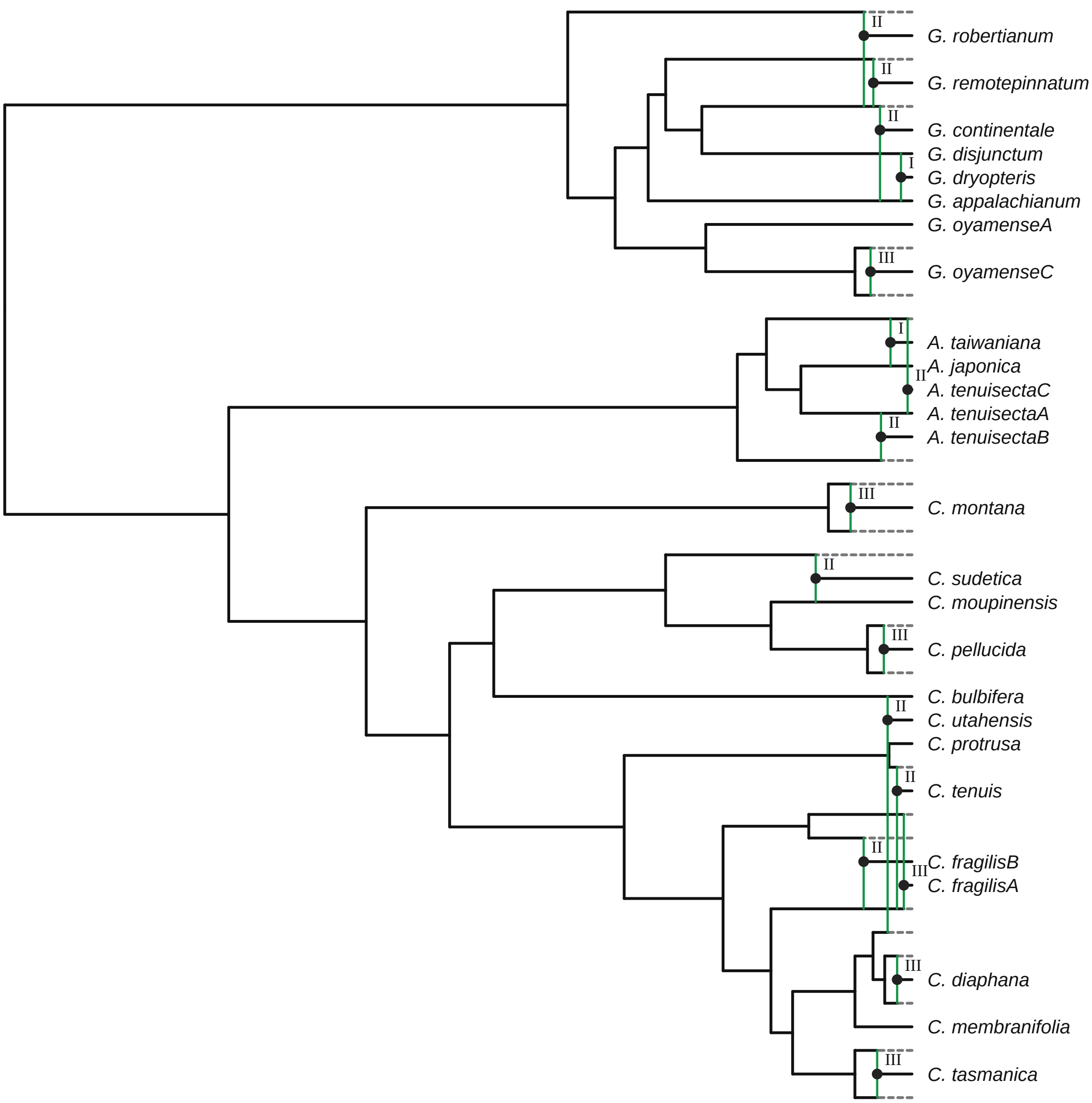


**Figure S10: Classification of reticulation events by type.** We depict the input phylogenetic network of Cystopteridaceae from Rothfels et al. (2017). Type I nodes have two zero-length reticulation edges. Type II nodes have one positive-length edge and one zero-length edge. Type III nodes have two positive-length edges. These types determine which reticulation ages are estimated, and what MCMC proposals we use on them.

*Proposal on continuous ages.*—In cases where the reticulation age $\tau_i$ is different from the maximum age, specifically for Type II and Type III nodes, we use a sliding proposal on the age. We propose a new age from the distribution

$$\tau_i' \sim \text{Uniform}(\tau_i - w, \tau_i + w)$$

reflecting the proposed age at boundaries so it remains between 0 and the maximum age. The proposal is symmetric, with a Hastings ratio of 1. We use the same value $w$ for all nodes, but tune it to have a global acceptance rate of 40% during the burnin phase.

*Proposal on discrete ages.*—Type II nodes can have ages exactly equal to their youngest ancestor. Denote the age of a Type II node $\tau_i$, and the age of the ancestor $t$. If the current state is $\tau_i = t$, the proposal is:

$$\tau_i' \sim \text{Uniform}(0, t)$$

*i.e.*, the new age is drawn uniformly between 0 and the maximum age (which will never propose an age of exactly $t$). If the current state is $\tau_i < t$, the discrete proposal is selected with probability 0.5 and sets the proposed age $\tau_i' = t$. This is essentially reversible jump MCMC, since the dimensionality changes between the two states. The Hastings ratio for the $\tau_i = t$ to $\tau_i' \neq t$ proposal is:

$$H(\tau_i' \neq t) = \frac{q(\tau_i \mid \tau_i')}{q(\tau_i' \mid \tau_i)} = \frac{t}{2},$$

because the forward proposal occurs drawns a new random variable with density $1/t$ and the reverse proposal occurs with probability 0.5. The Hastings ratio in the other direction is just the inverse:

$$H(\tau_i' = t) = \frac{q(\tau_i \mid \tau_i')}{q(\tau_i' \mid \tau_i)} = \frac{2}{t},$$

#### S4.2 Ghost lineages

We compute the posterior number of ghost lineages as 21 (the minimum number) plus one per additional ghost lineage in the network, as determined by the number of reticulation events that are not exactly as old as their divergence from their sister lineage. The posterior distribution for the analysis with $u_{\max} = 512$ is depicited in Fig. S11.

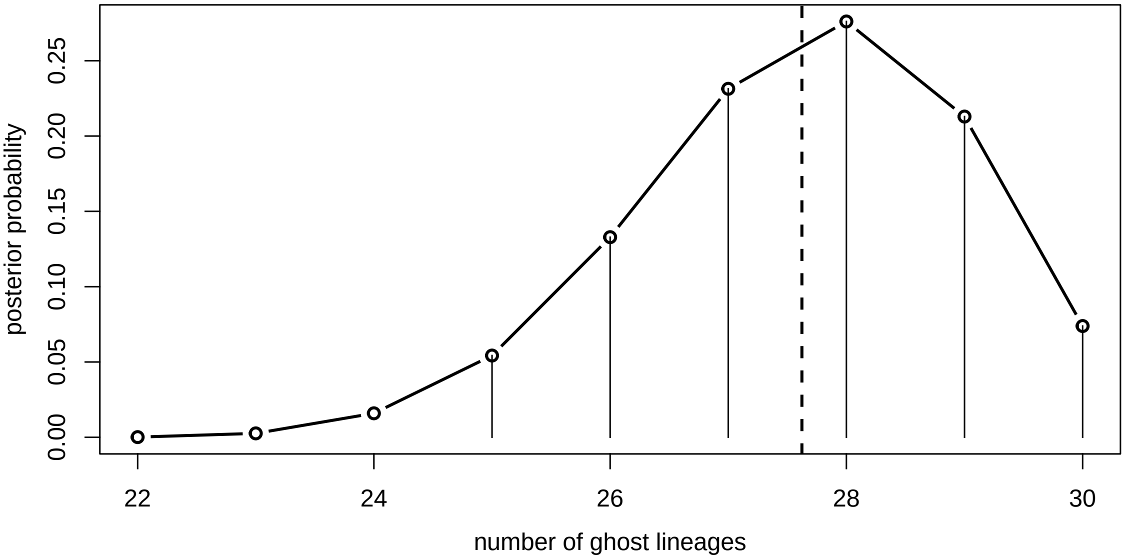


**Figure S11: Posterior distributions of ghost lineages for $u_{\mathbf{max}} = 512$.** Dots represent the full posterior probability distribution of the number of ghost lineages. Dashed vertical line is the posterior mean. Solid vertical lines are drawn for ghost lineage numbers that are contained by the 95% credible interval, which ranges from 25 to 30.

#### S4.3 Sensitivity of $u_{\mathbf{max}}$

We explored the impact of posterior estimates to the truncation parameter $u_{\text{max}}$. As $u_{\text{max}}$ increases, the posterior distribution should converge to the posterior distribution we would estimate under an untruncated (unapproximated) model. We varied $u_{\text{max}} = \{32, 64, 128, 256, 512\}$ (successive doublings), while keeping the remaining settings (including prior distributions) the same. These analyses indicate that the posterior distributions stabilize with by the time $u_{\text{max}} = 128$ (Fig. S12).

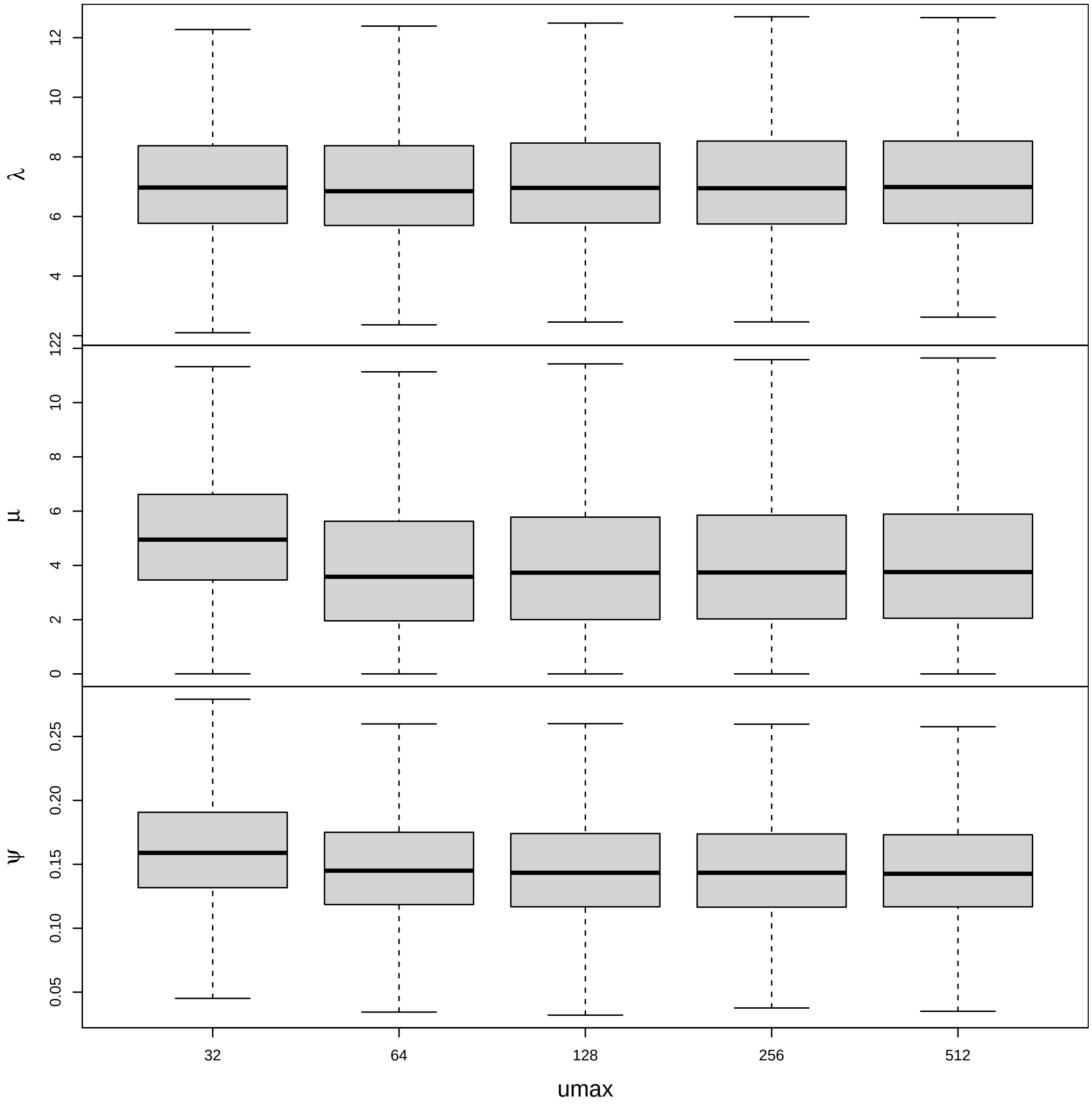


**Figure S12: Posterior distributions of model parameters as a function of $u_{\mathbf{max}}$.** Each row corresponds to a model parameter, the speciation rate $\lambda$, extinction rate $\mu$, and allopolyploidization rate $\psi$. Boxes correspond to the posterior distributions under the $u_{\text{max}}$ value on the x-axis.

#### S4.4 The effect of estimating allopolyploidization ages

In our main analyses, we infer the age of each reticulation event, between 0 (the present) and its maximum age, as determined by the youngest divergence from either sister lineage. Whether a reticulation event occurs at exactly the maximum age, or sometime after the maximum age, has a significant influence in the interpretation of ancestral nodes in the network: in the former case, the ancestral node must be a reticulation event, in the latter case, it must be a speciation event. Whether a particular ancestral node is considered a speciation event or an allopolyploid parent should affect estimates of the corresponding rate parameters.

Indeed, when we estimate parameters using the fixed ages from the Rothfels et al. (2017) network, they are different from those when we estimate the ages (Fig. S13). In particular, speciation and extinction rates are lower and the allopolyploidization rate is higher, when using the fixed network. The decrease in the speciation rate likely reflects the fact that there are fewer observed speciation events in the fixed network. Likewise, the decrease in the extinction rate is likely due to the fact that the fixed network implies fewer ghost lineages, therefore fewer lineages that are required to go extinct (or be unsampled).The modest increase in the allopolyploidization rate is more difficult to understand, since the number of allopolyploidization events in the network is the same regardless of their ages; however, it is possible the increase is because older allopolyploidization events have fewer contemporary lineages to arise from, therefore implying an increased allopolyploidization rate.

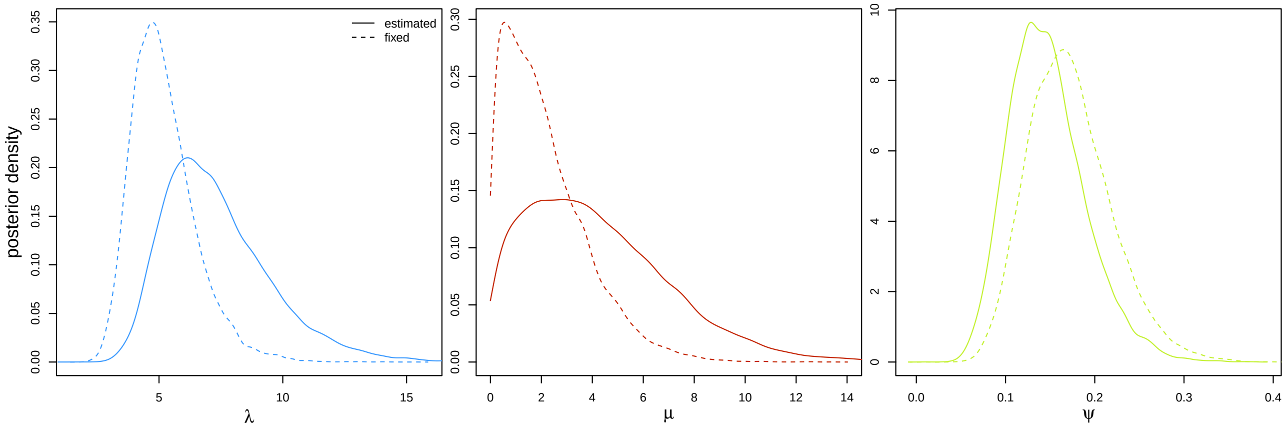


**Figure S13: Posterior distributions of the birth-death-allopolyploidization parameters for Cystopteridaceae, with and without estimated reticulation ages ($u_{\mathbf{max}} = 512$).** Curves correspond to the approximate posterior distribution of the model parameters with estimated ages (solid lines) or with fixed ages (dashed lines).

#### S4.5 Consistency between custom sampler and `AdaptiveMCMC`

To validate our MCMC sampler, we compared posterior estimates between our custom sampler and the equivalent sampler implemented in `AdaptiveMCMC` (Vihola 2026). Figure S14 demonstrates that the posterior estimates under these two implementations are essentially identical.

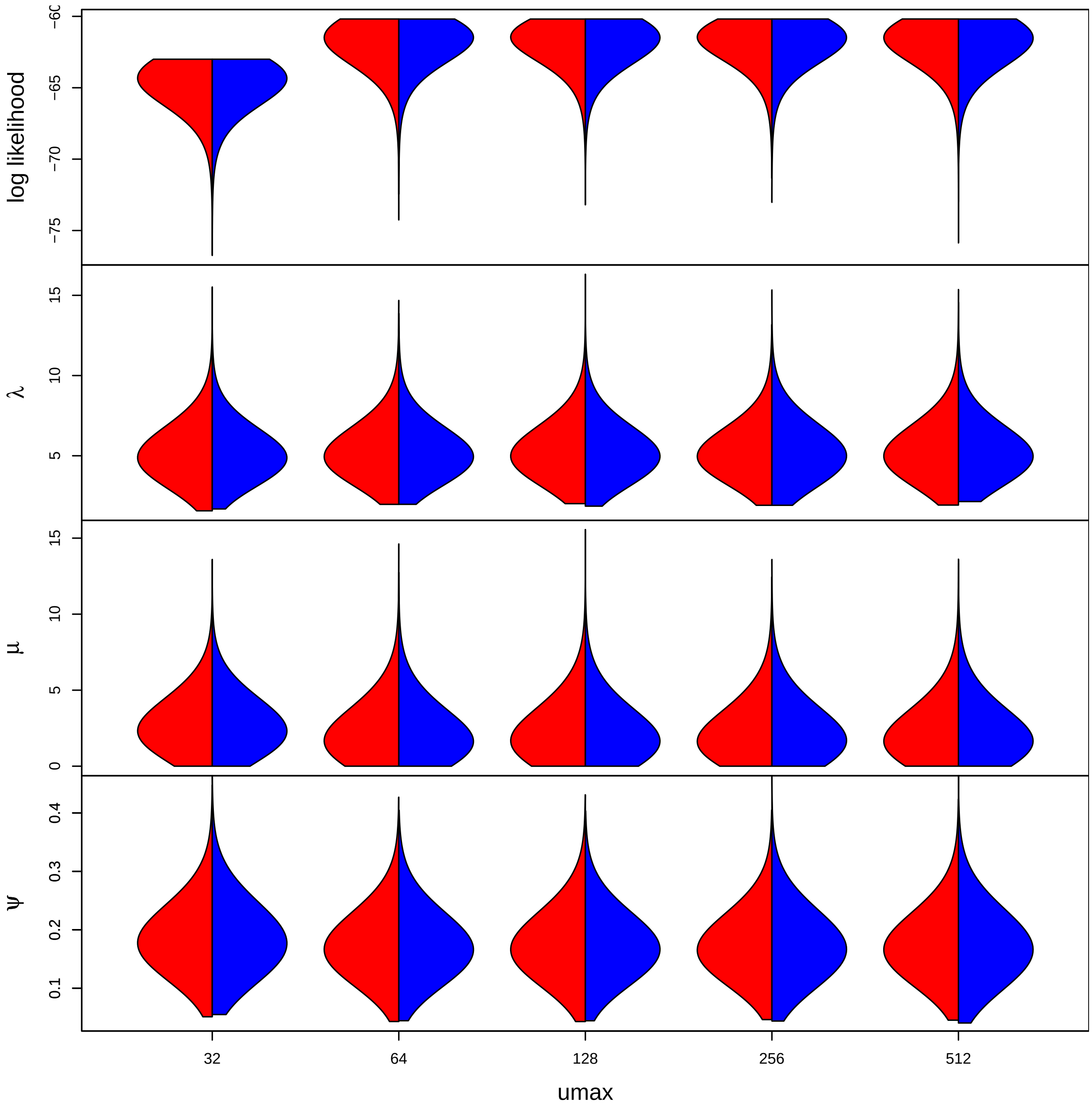


**Figure S14: Posterior distributions using our custom sampler compared to `AdaptiveMCMC`.** We estimated posterior distributions for each value of $u_{max}$ using the fixed network from Rothfels et al. (2017). We did this using our custom sampler (red) and the off-the-shelf sampler implemented in `AdaptiveMCMC`. We show the posterior distributions as a function of $u_{max}$ (x axis) and for each model parameter (and the likelihood) in rows.

#### S4.6 Sensitivity of ghost lineages to the sampling fraction $\rho$

To understand the influence of the assumed sampling fraction on inferences about ghost lineages, we conducted additional analyses with $u_{\text{max}} = 512$ and $\rho = 1$. The posterior-mean-age summary network demonstrates that younger allopolyploid lineages are less likely to have ghost parents (Fig. S16), and the posterior distribution of the number of ghost lineages is shifted down by approximately 1.5, on average (Fig. S15).

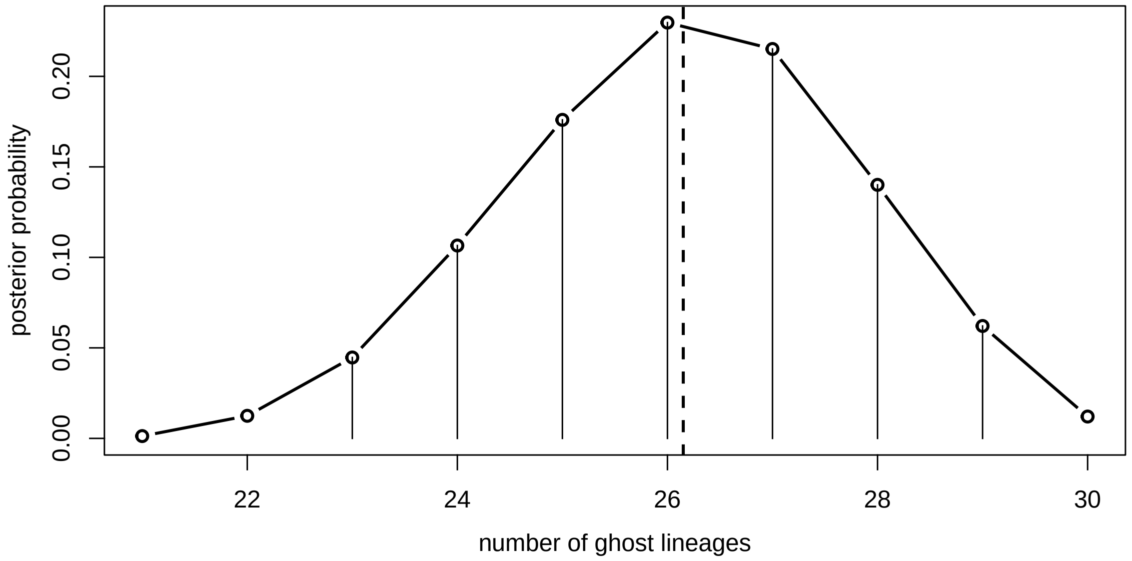


**Figure S15: Posterior distributions of ghost lineages for $u_{\text{max}} = 512$ with $\rho = 1$.** Assuming a larger sampling fraction, $\rho = 1$, decreases the posterior distribution of the number of ghost lineages (compare against Fig. S11).

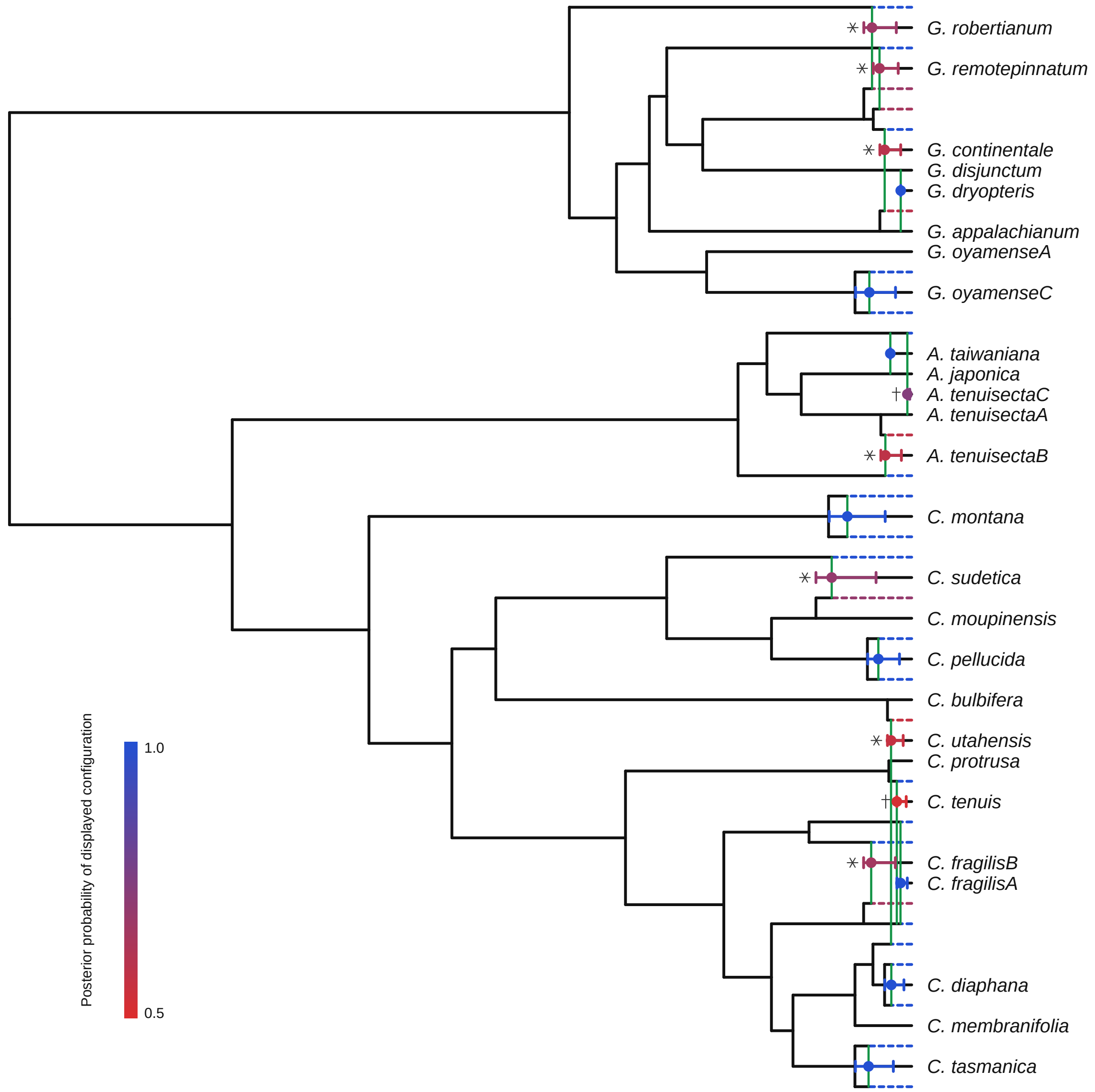


**Figure S16: Summary network of Cystopteridaceae estimated using $\rho = 1$.** Graphical conventions follow Fig. X in the main text.

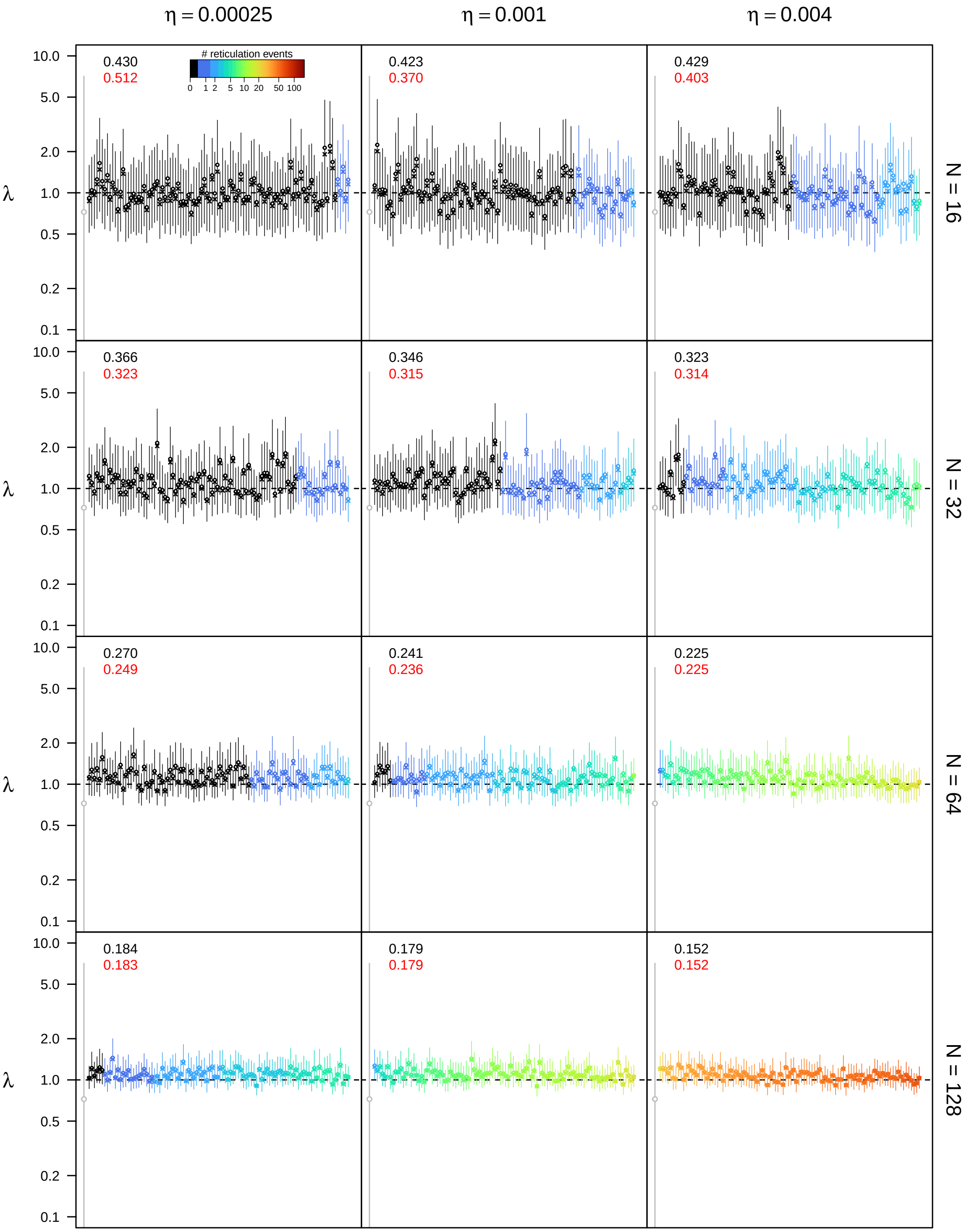


**Figure S17: Posterior estimates of the speciation rate $\lambda$ under the unidirectional hybridization model for $\epsilon = 0.1$, $\rho = 0.5$, and $u_{\mathbf{max}} = 4N$.** See main text for interpretation.

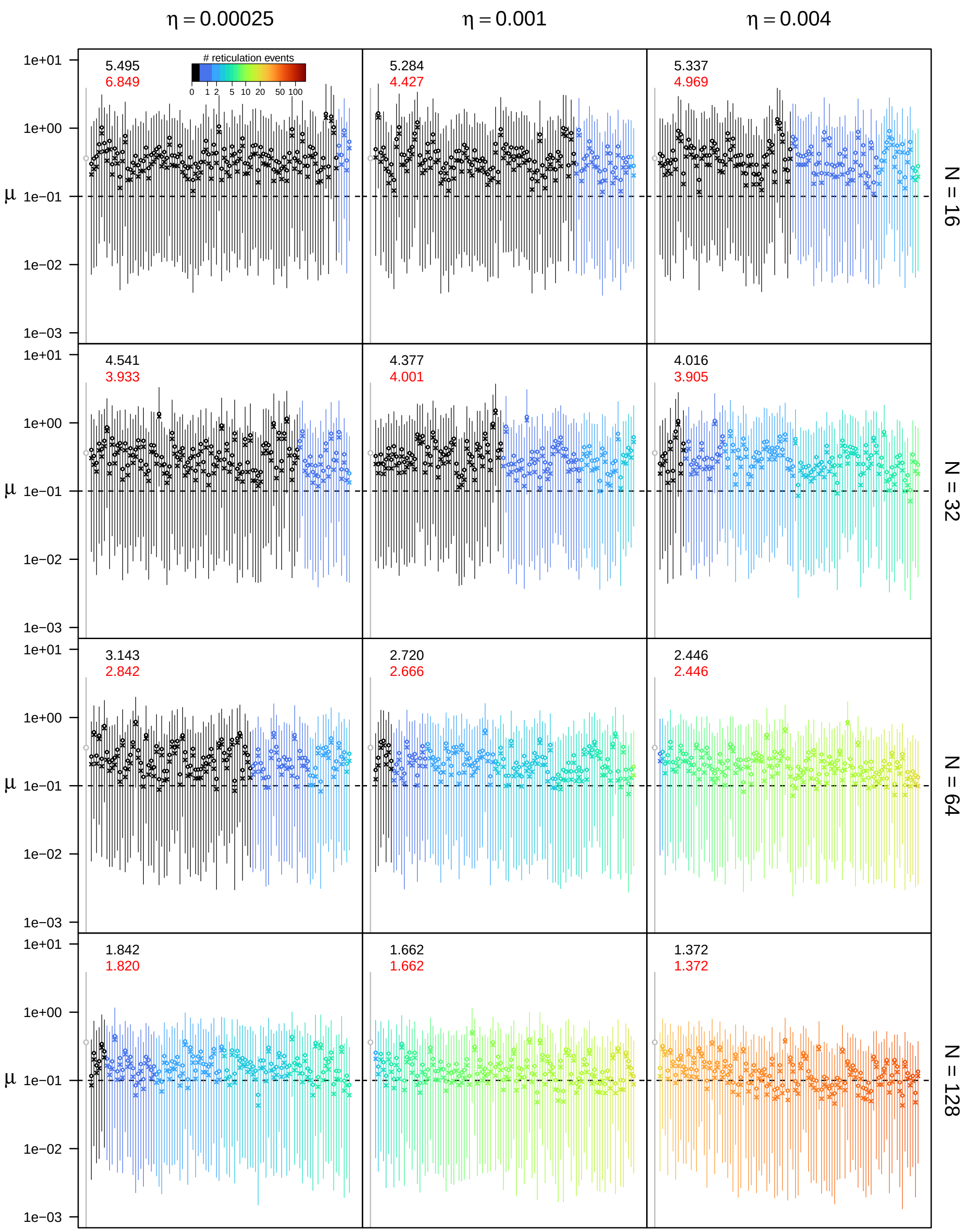


**Figure S18: Posterior estimates of the extinction rate $\mu$ under the unidirectional hybridization model for $\epsilon = 0.1$, $\rho = 0.5$, and $u_{\mathbf{max}} = 4N$.** See main text for interpretation.

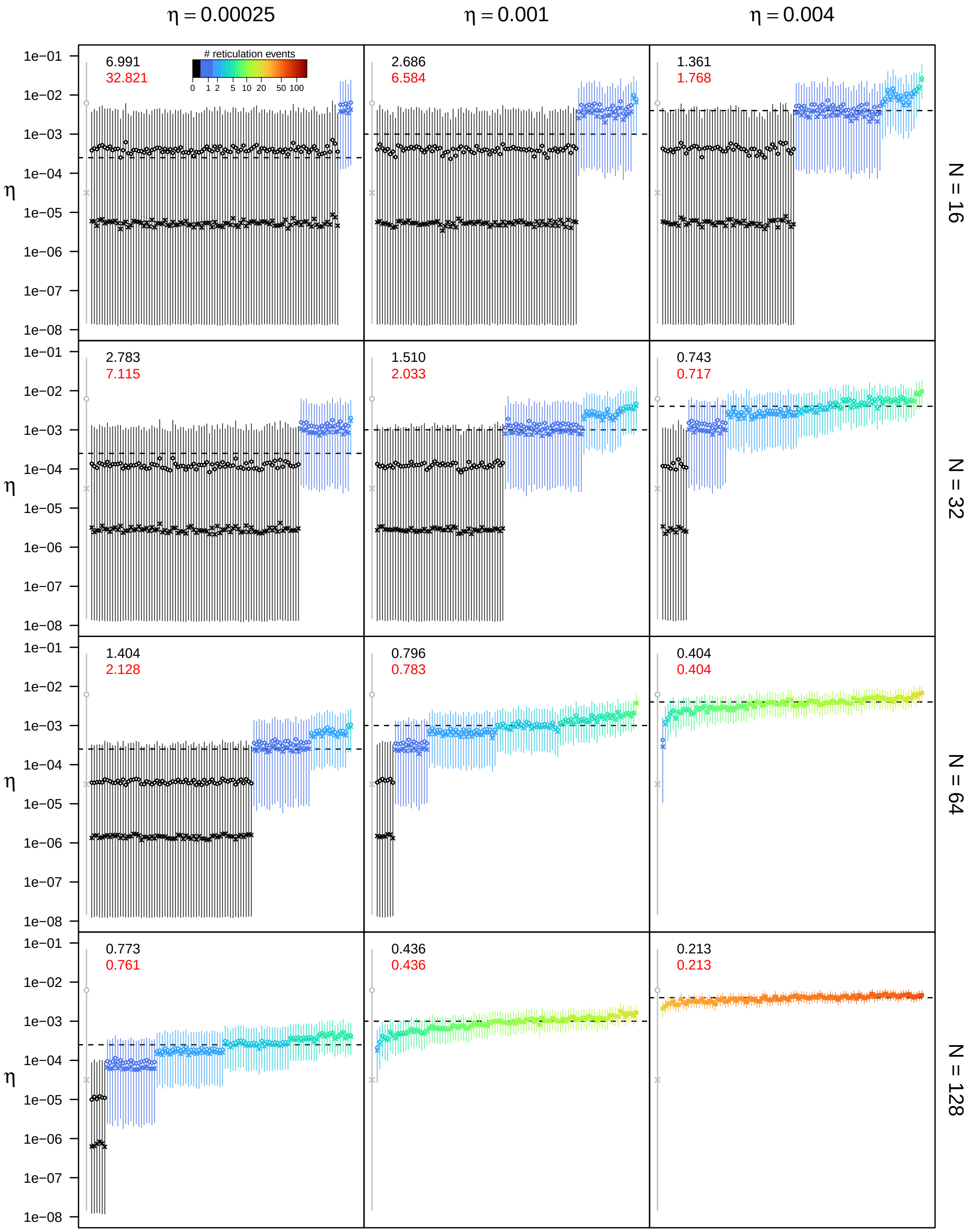


**Figure S19: Posterior estimates of the unidirectional hybridization rate $\eta$ under the unidirectional hybridization model for $\epsilon = 0.1$, $\rho = 0.5$, and $u_{\mathbf{max}} = 4N$.** See main text for interpretation.

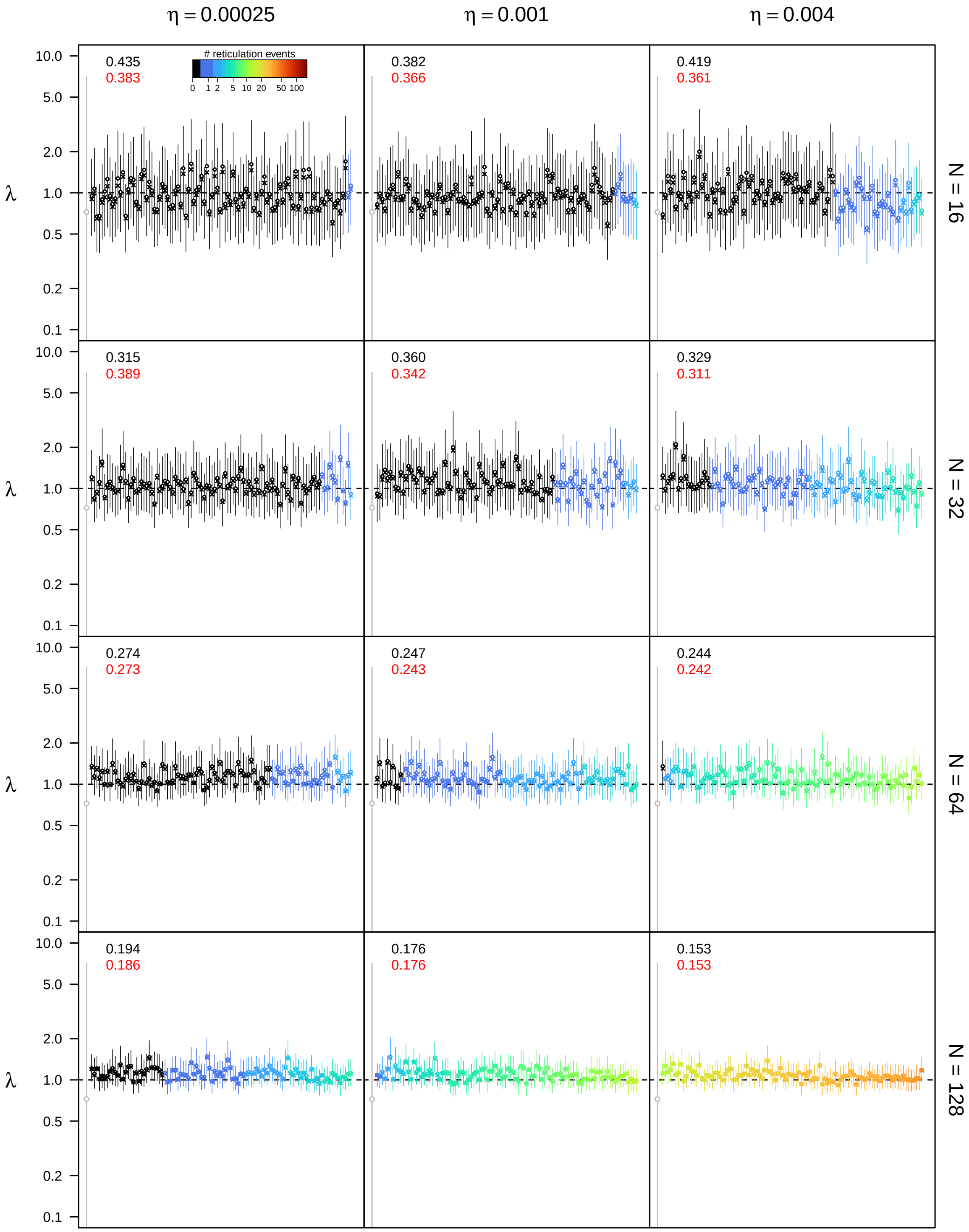


**Figure S20: Posterior estimates of the speciation rate $\lambda$ under the unidirectional hybridization model for $\epsilon = 0.1$, $\rho = 0.75$, and $u_{\mathbf{max}} = 4N$.** See main text for interpretation.

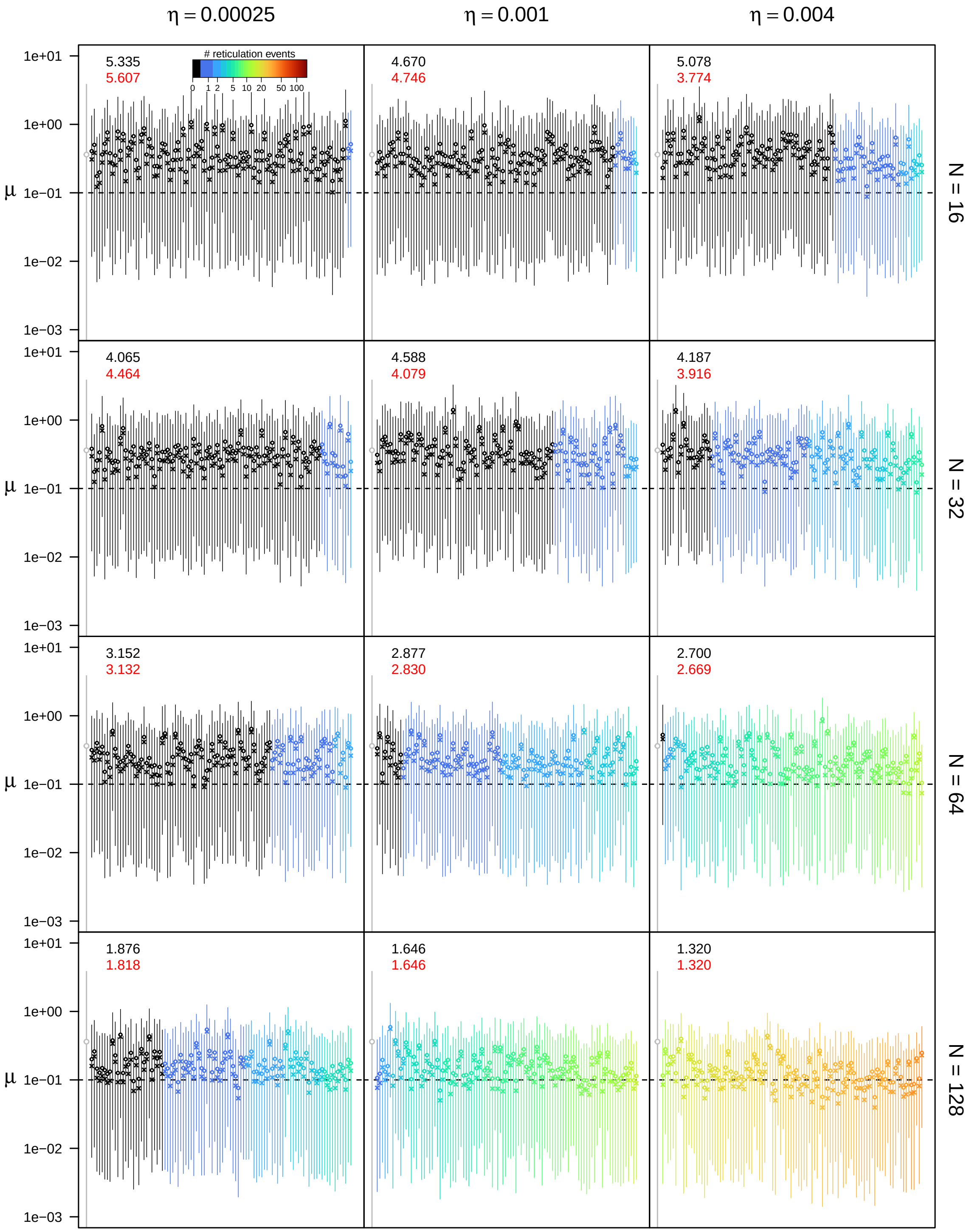


**Figure S21: Posterior estimates of the extinction rate $\mu$ under the unidirectional hybridization model for $\epsilon = 0.1$, $\rho = 0.75$, and $u_{\mathbf{max}} = 4N$.** See main text for interpretation.

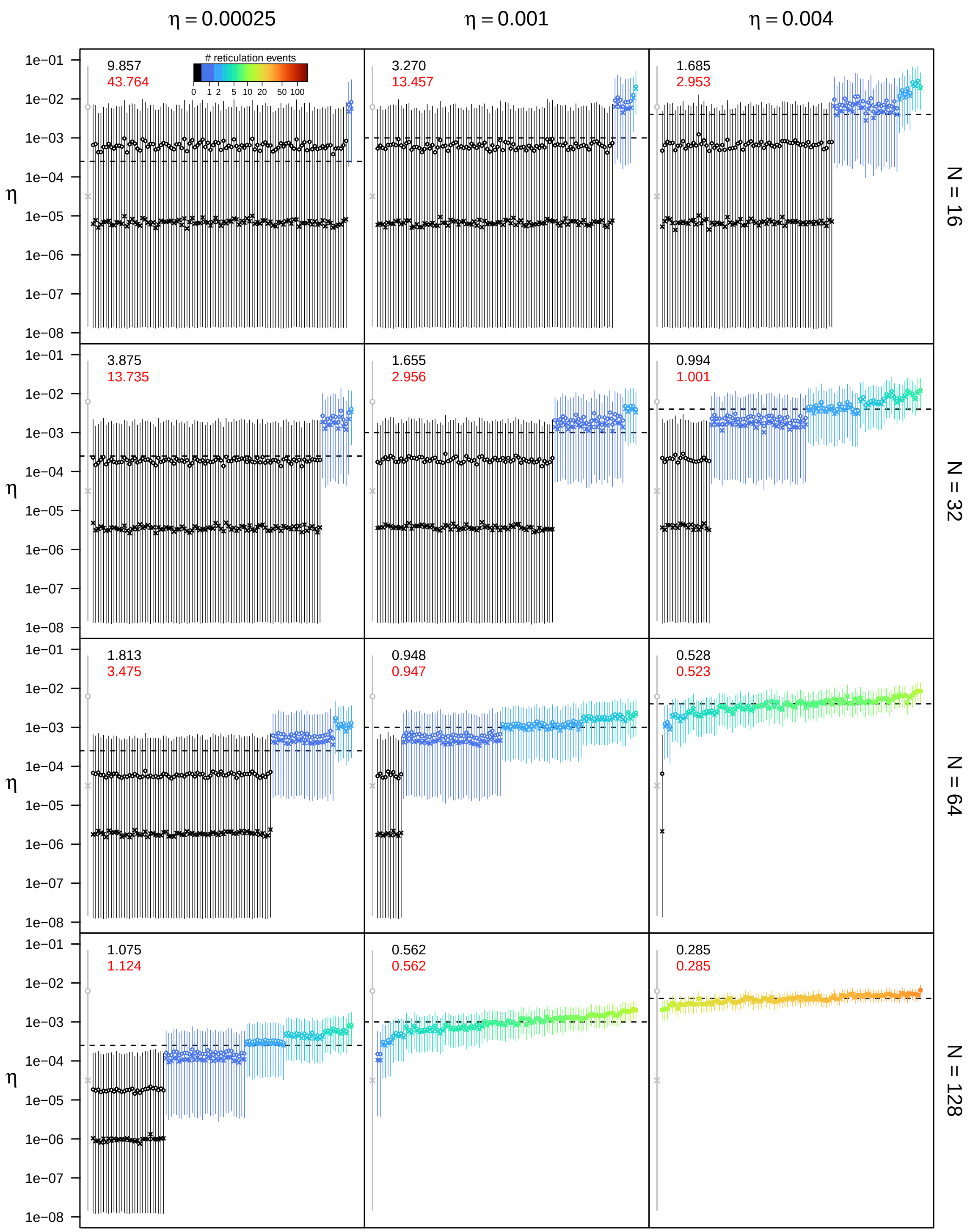


**Figure S22: Posterior estimates of the unidirectional hybridization rate $\eta$ under the unidirectional hybridization model for $\epsilon = 0.1$, $\rho = 0.75$, and $u_{\mathbf{max}} = 4N$.** See main text for interpretation.

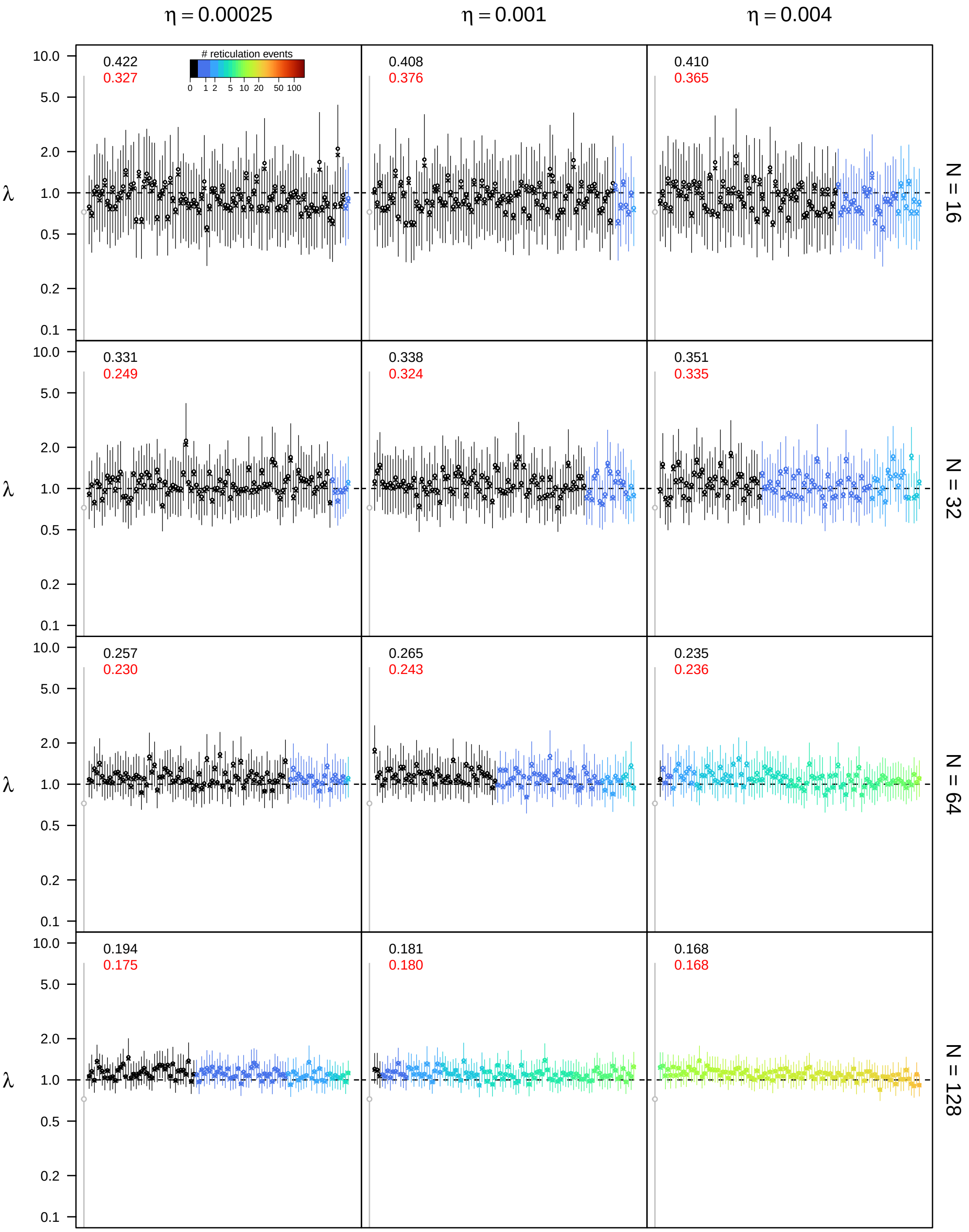


**Figure S23: Posterior estimates of the speciation rate $\lambda$ under the unidirectional hybridization model for $\epsilon = 0.1$, $\rho = 1.0$, and $u_{\max} = 4N$.** See main text for interpretation.

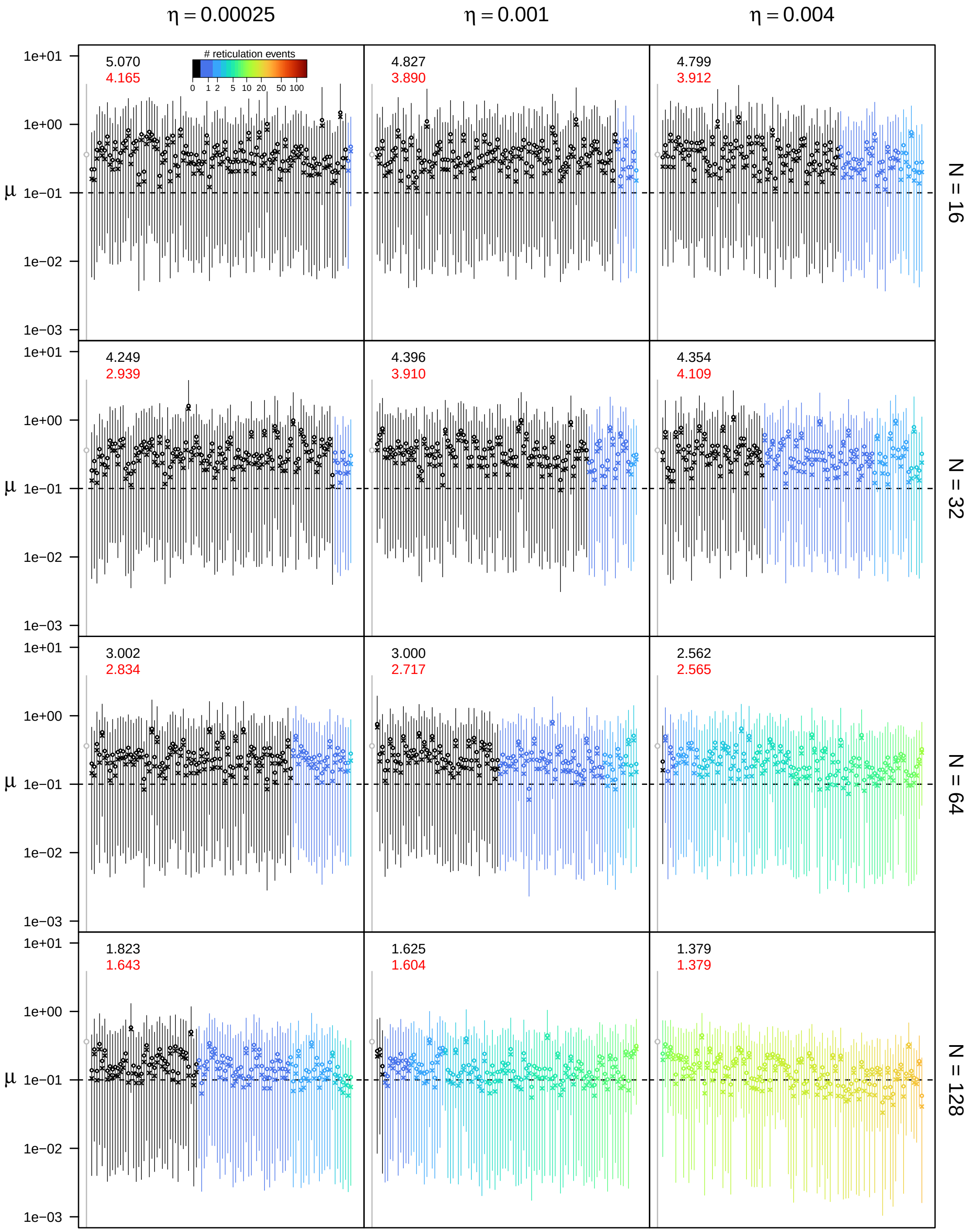


**Figure S24: Posterior estimates of the extinction rate $\mu$ under the unidirectional hybridization model for $\epsilon = 0.1$, $\rho = 1.0$, and $u_{\mathbf{max}} = 4N$.** See main text for interpretation.

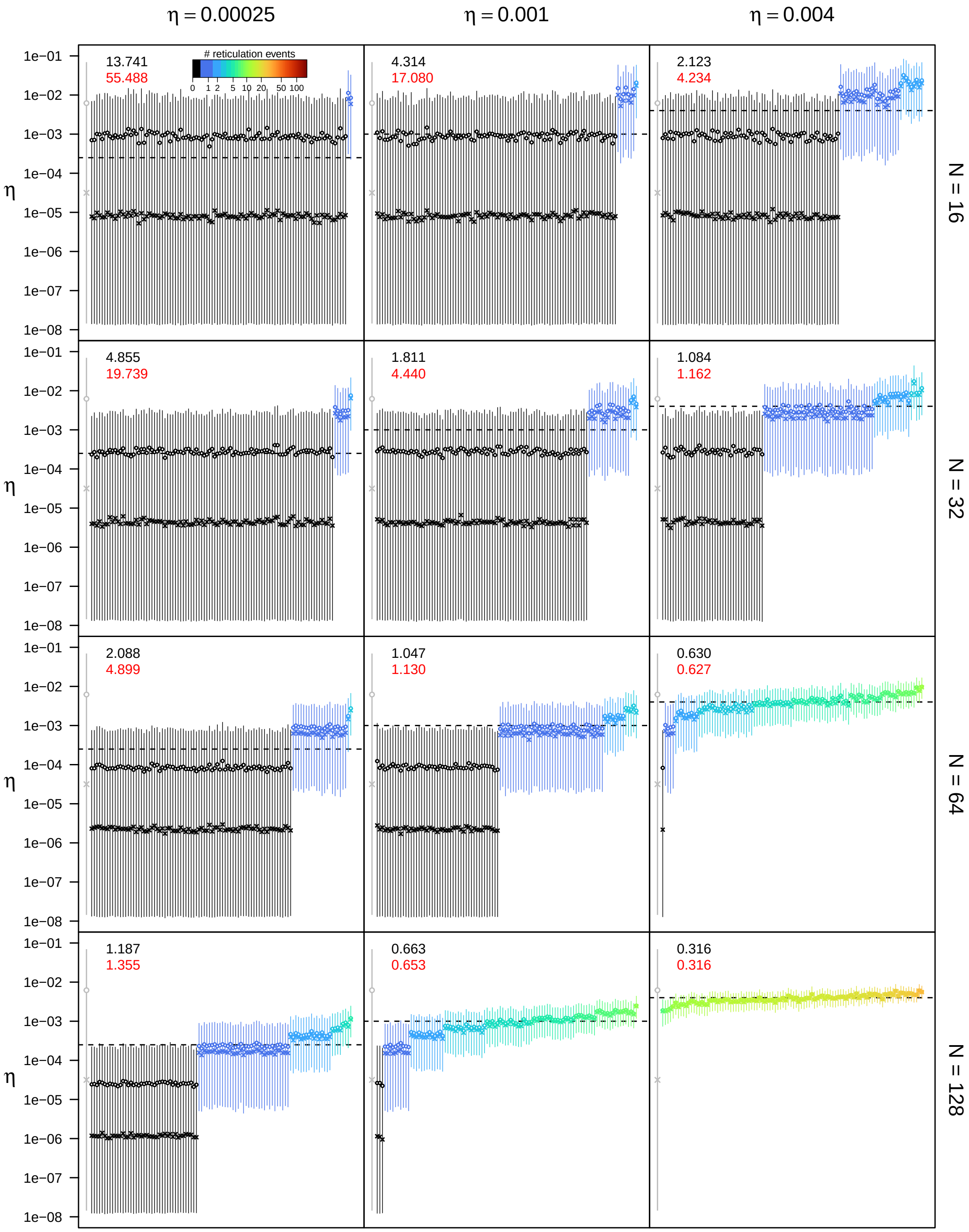


**Figure S25: Posterior estimates of the unidirectional hybridization rate $\eta$ under the unidirectional hybridization model for $\epsilon = 0.1$, $\rho = 1.0$, and $u_{\max} = 4N$.** See main text for interpretation.

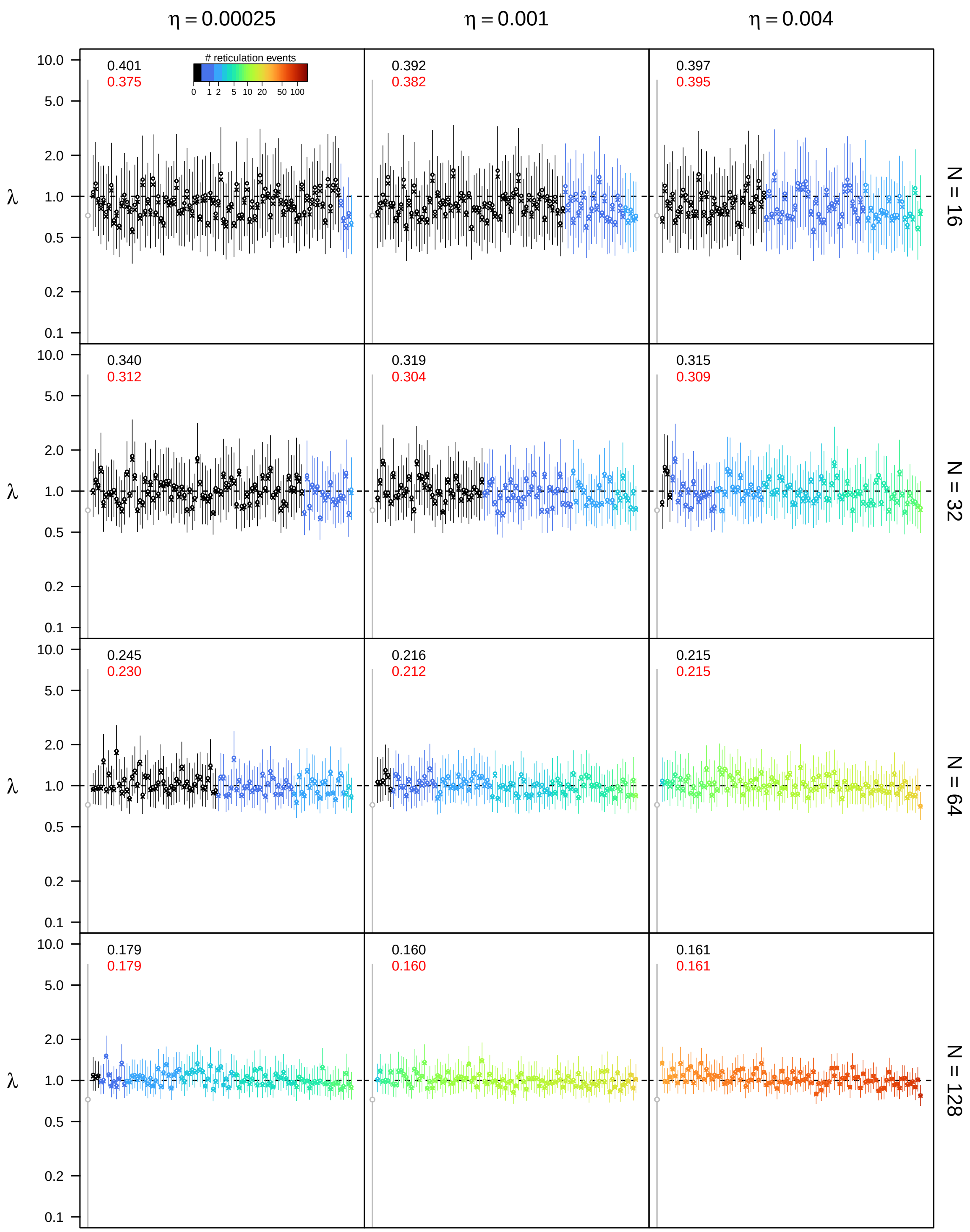


**Figure S26: Posterior estimates of the speciation rate $\lambda$ under the unidirectional hybridization model for $\epsilon = 0.3$, $\rho = 0.5$, and $u_{\mathbf{max}} = 4N$.** See main text for interpretation.

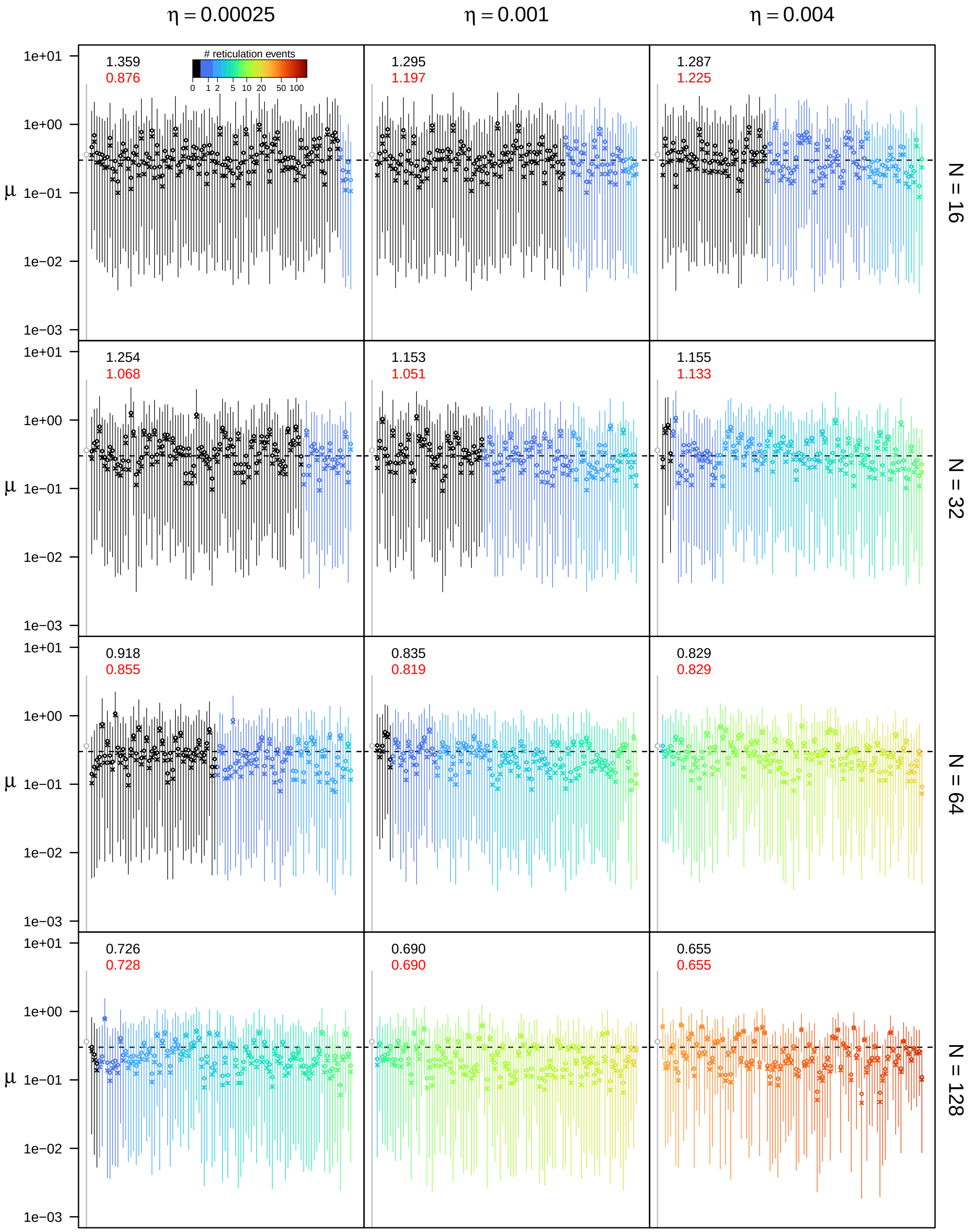


**Figure S27: Posterior estimates of the extinction rate $\mu$ under the unidirectional hybridization model for $\epsilon = 0.3$, $\rho = 0.5$, and $u_{\mathbf{max}} = 4N$.** See main text for interpretation.

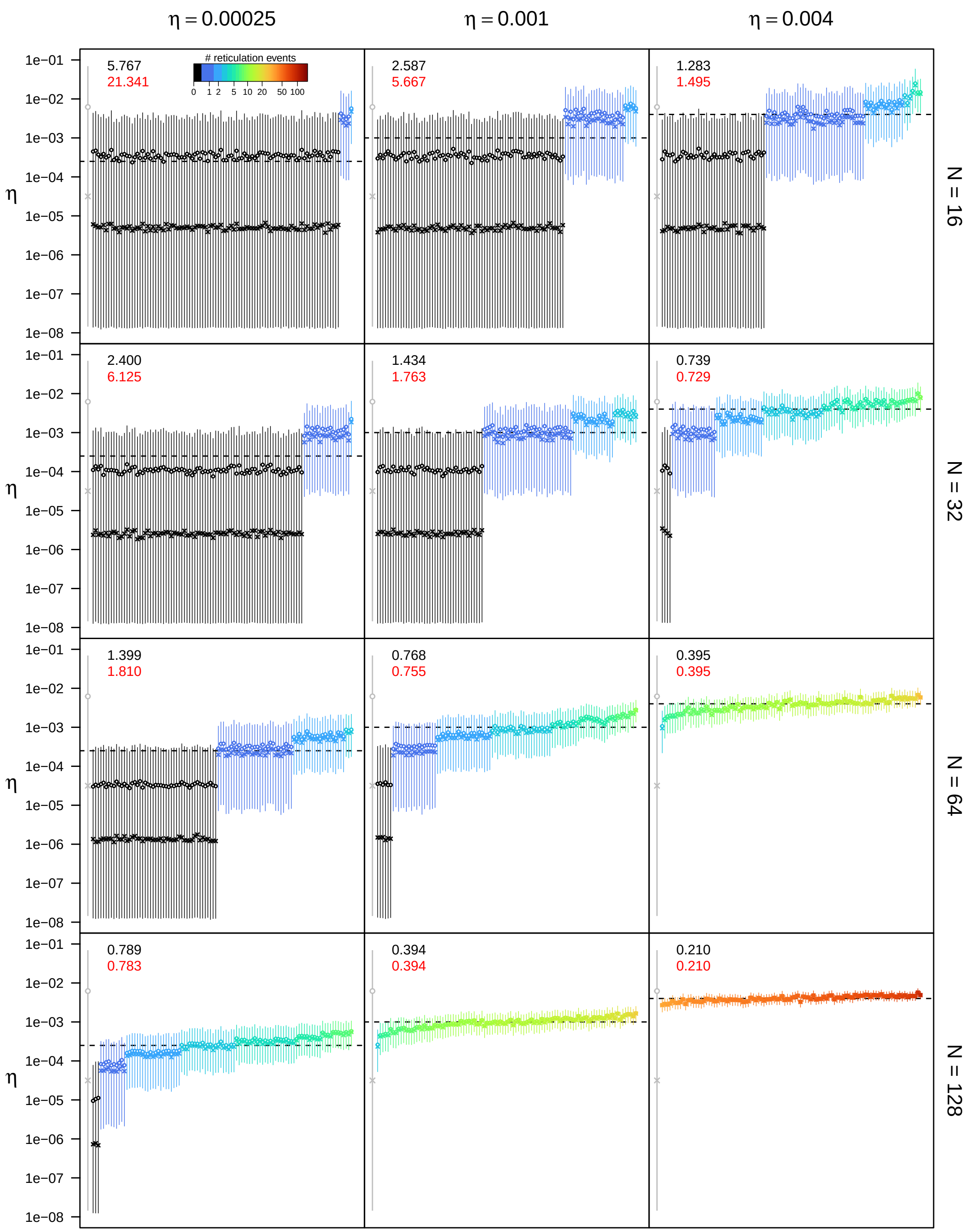


**Figure S28: Posterior estimates of the unidirectional hybridization rate $\eta$ under the unidirectional hybridization model for $\epsilon = 0.3$, $\rho = 0.5$, and $u_{\mathbf{max}} = 4N$.** See main text for interpretation.

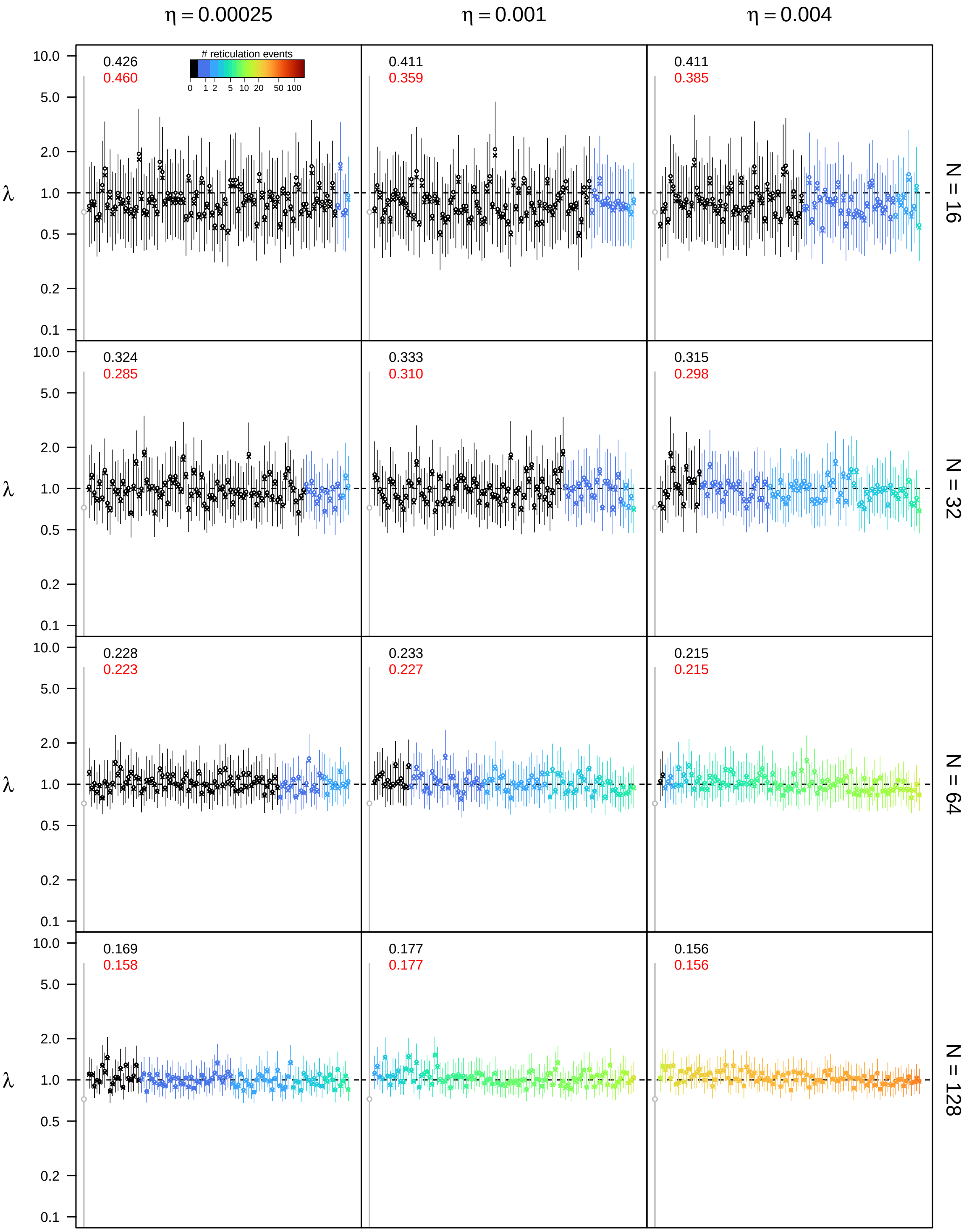


**Figure S29: Posterior estimates of the speciation rate $\lambda$ under the unidirectional hybridization model for $\epsilon = 0.3$, $\rho = 0.75$, and $u_{\mathbf{max}} = 4N$.** See main text for interpretation.

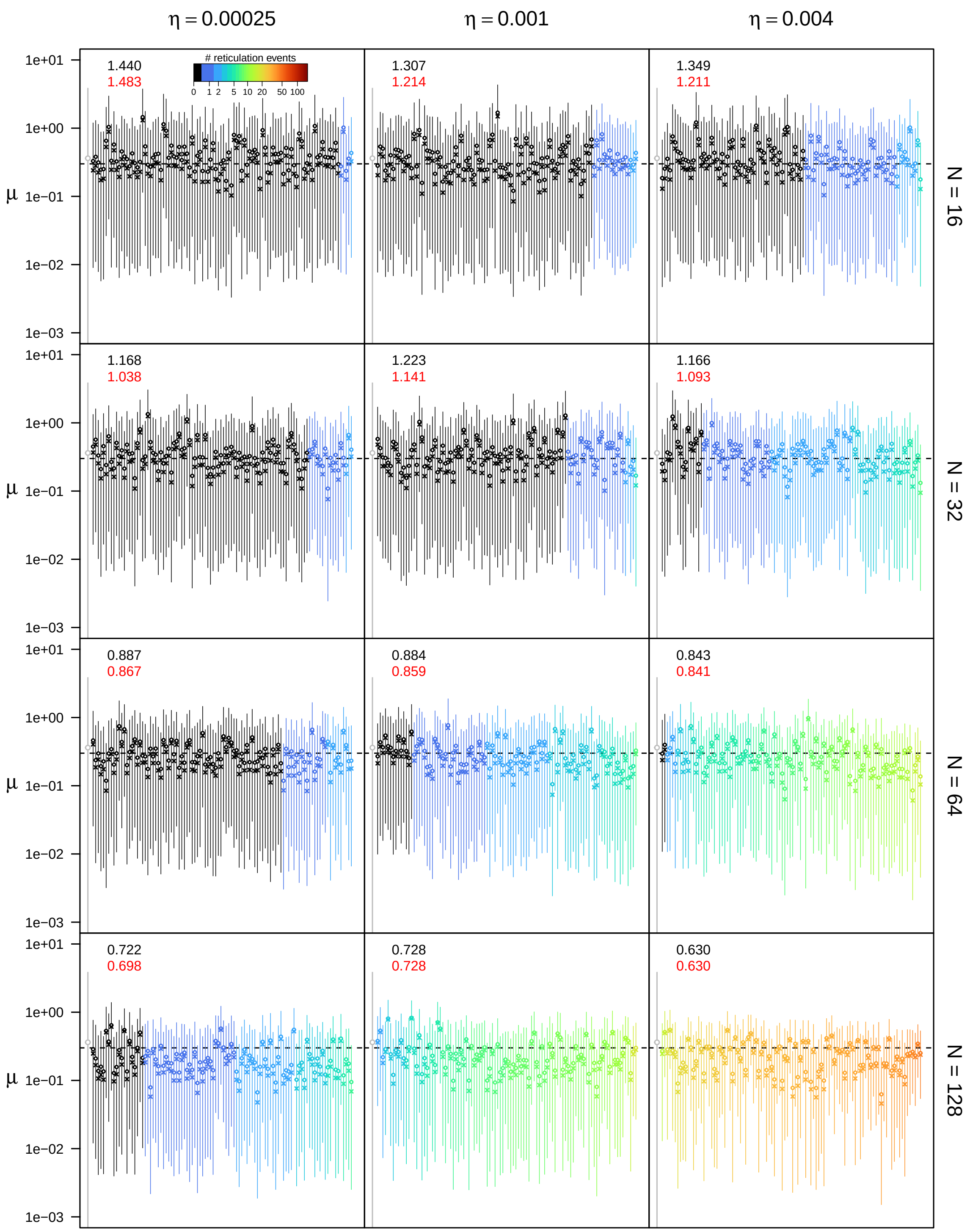


**Figure S30: Posterior estimates of the extinction rate $\mu$ under the unidirectional hybridization model for $\epsilon = 0.3$, $\rho = 0.75$, and $u_{\mathbf{max}} = 4N$.** See main text for interpretation.

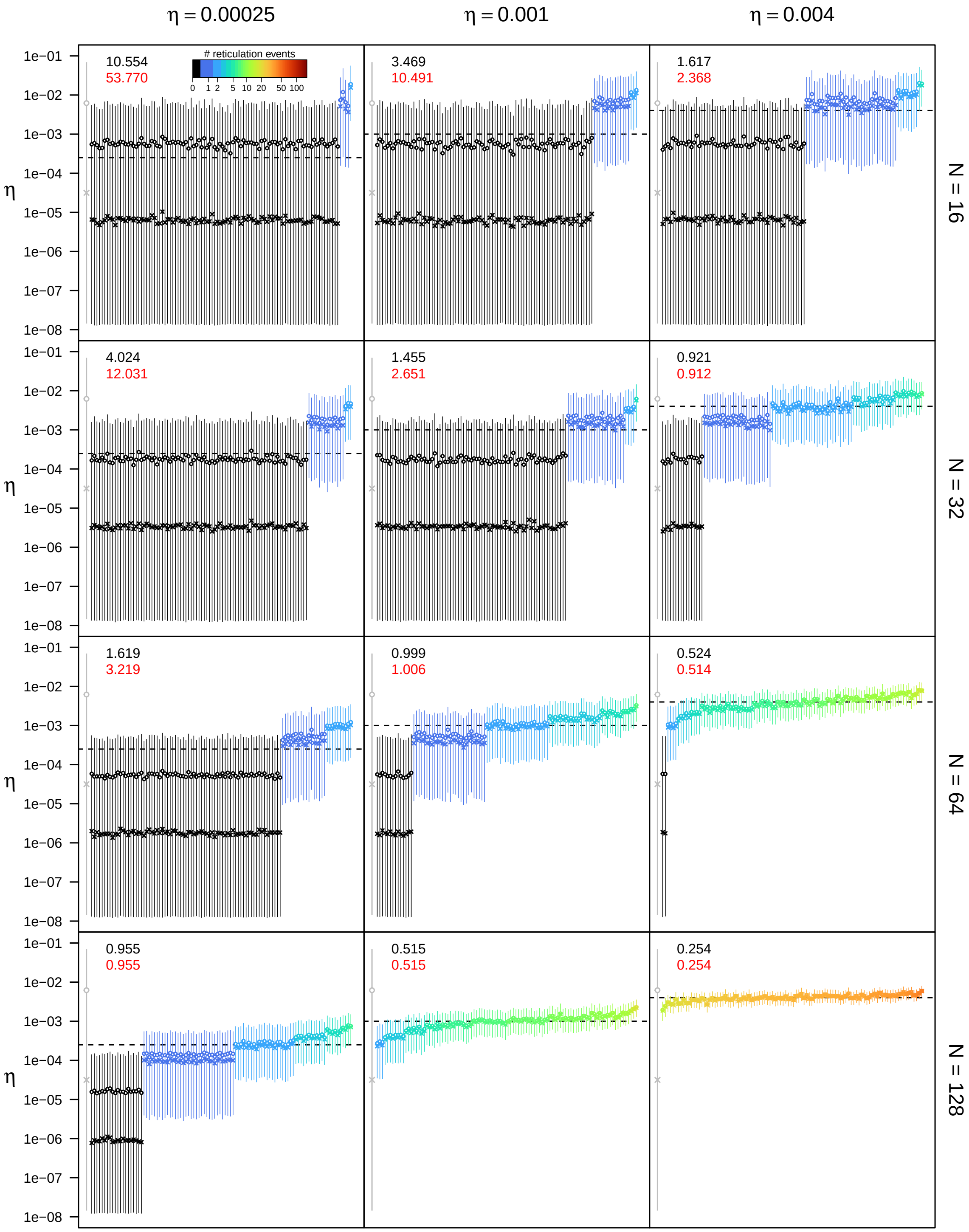


**Figure S31: Posterior estimates of the unidirectional hybridization rate $\eta$ under the unidirectional hybridization model for $\epsilon = 0.3$, $\rho = 0.75$, and $u_{\mathbf{max}} = 4N$.** See main text for interpretation.

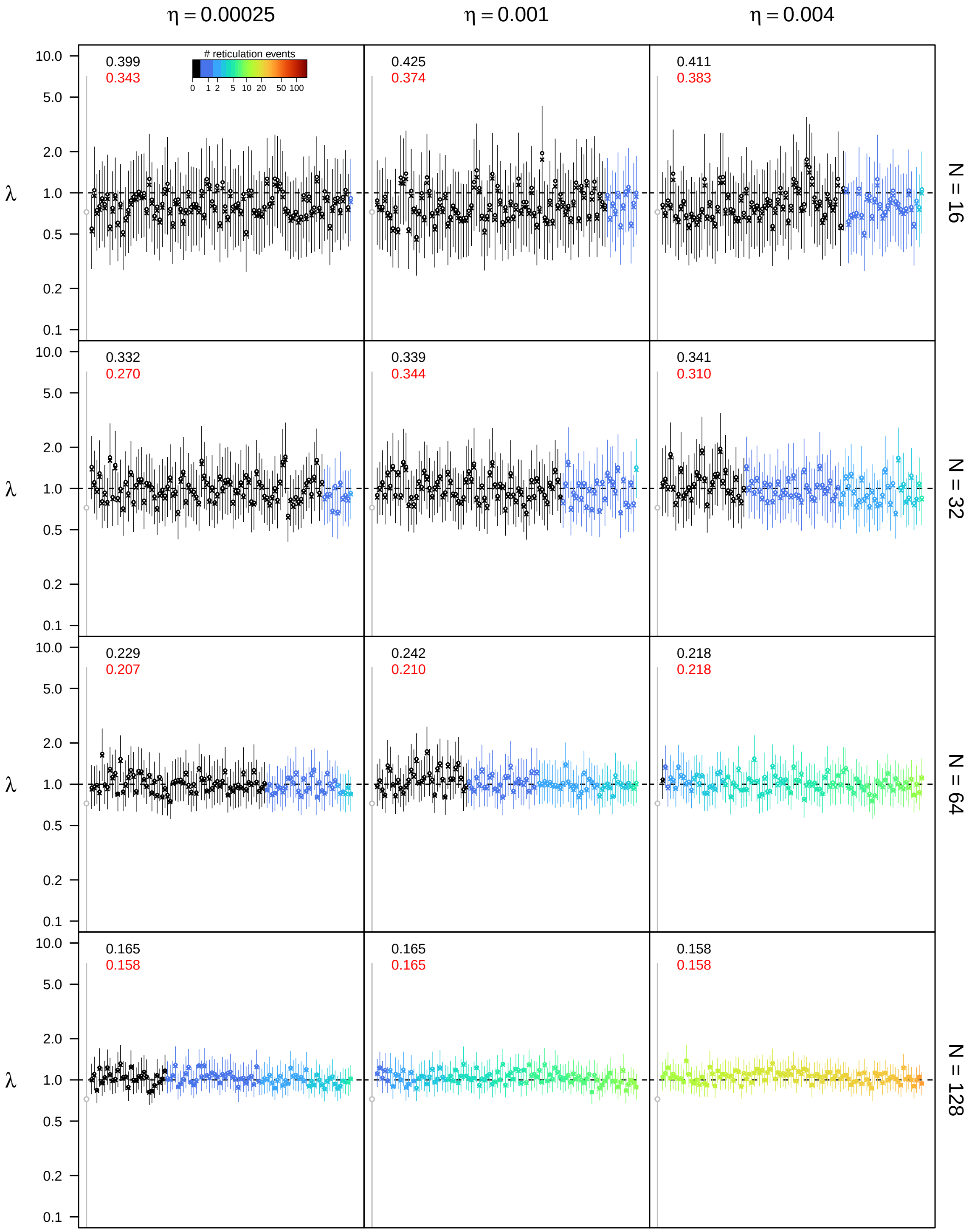


**Figure S32: Posterior estimates of the speciation rate $\lambda$ under the unidirectional hybridization model for $\epsilon = 0.3$, $\rho = 1.0$, and $u_{\mathbf{max}} = 4N$.** See main text for interpretation.

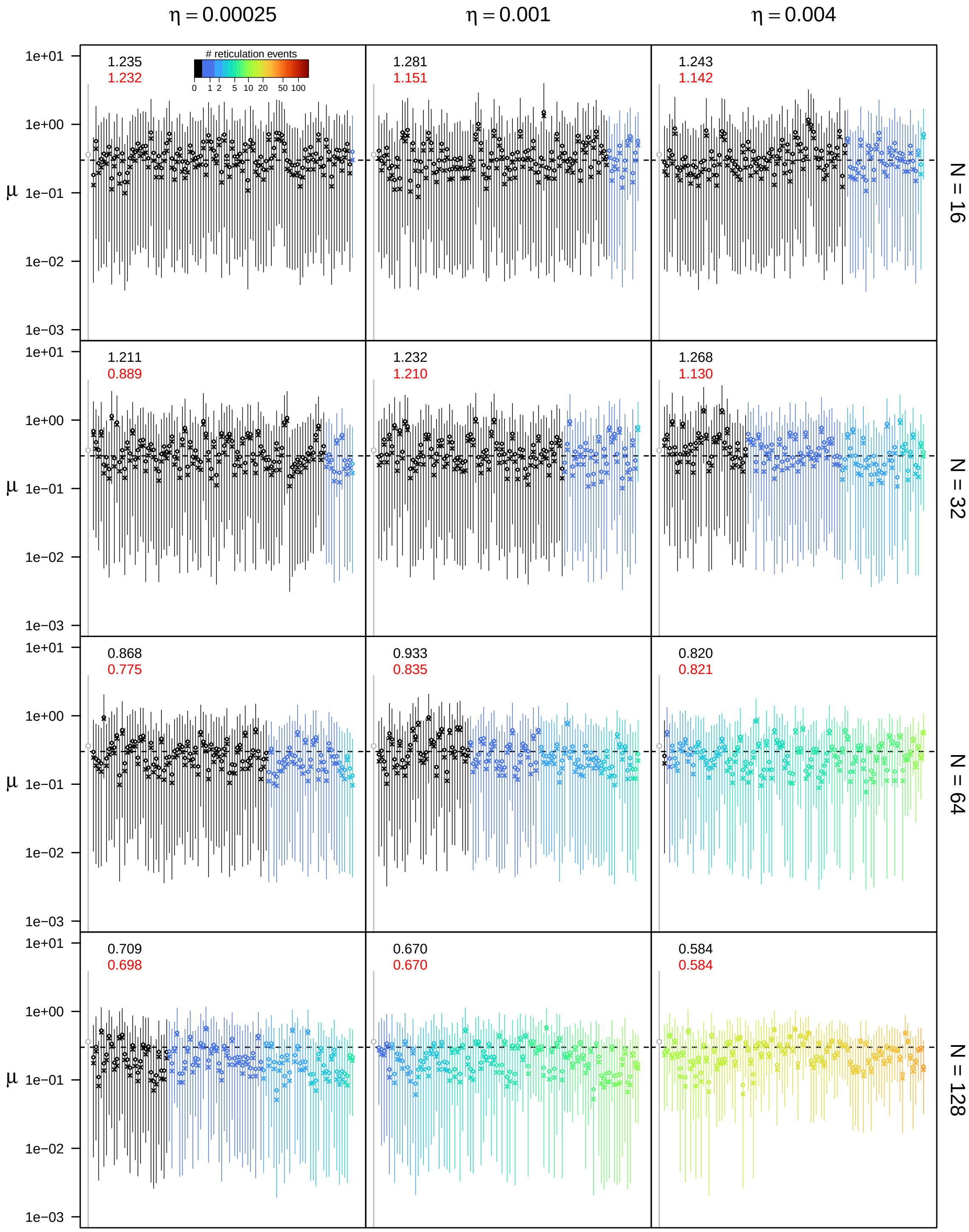


**Figure S33: Posterior estimates of the extinction rate $\mu$ under the unidirectional hybridization model for $\epsilon = 0.3$, $\rho = 1.0$, and $u_{\mathbf{max}} = 4N$.** See main text for interpretation.

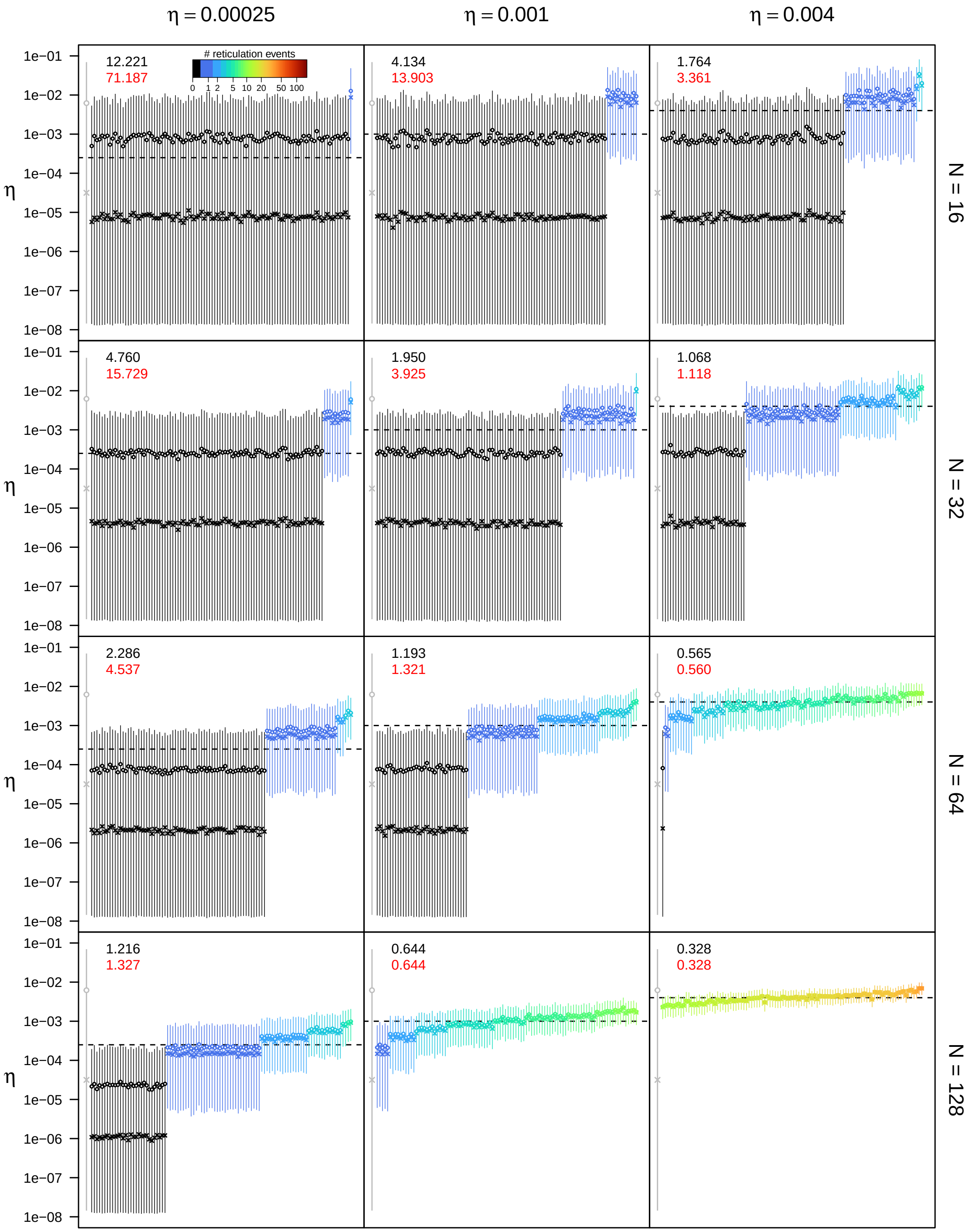


**Figure S34: Posterior estimates of the unidirectional hybridization rate $\eta$ under the unidirectional hybridization model for $\epsilon = 0.3$, $\rho = 1.0$, and $u_{\max} = 4N$.** See main text for interpretation.

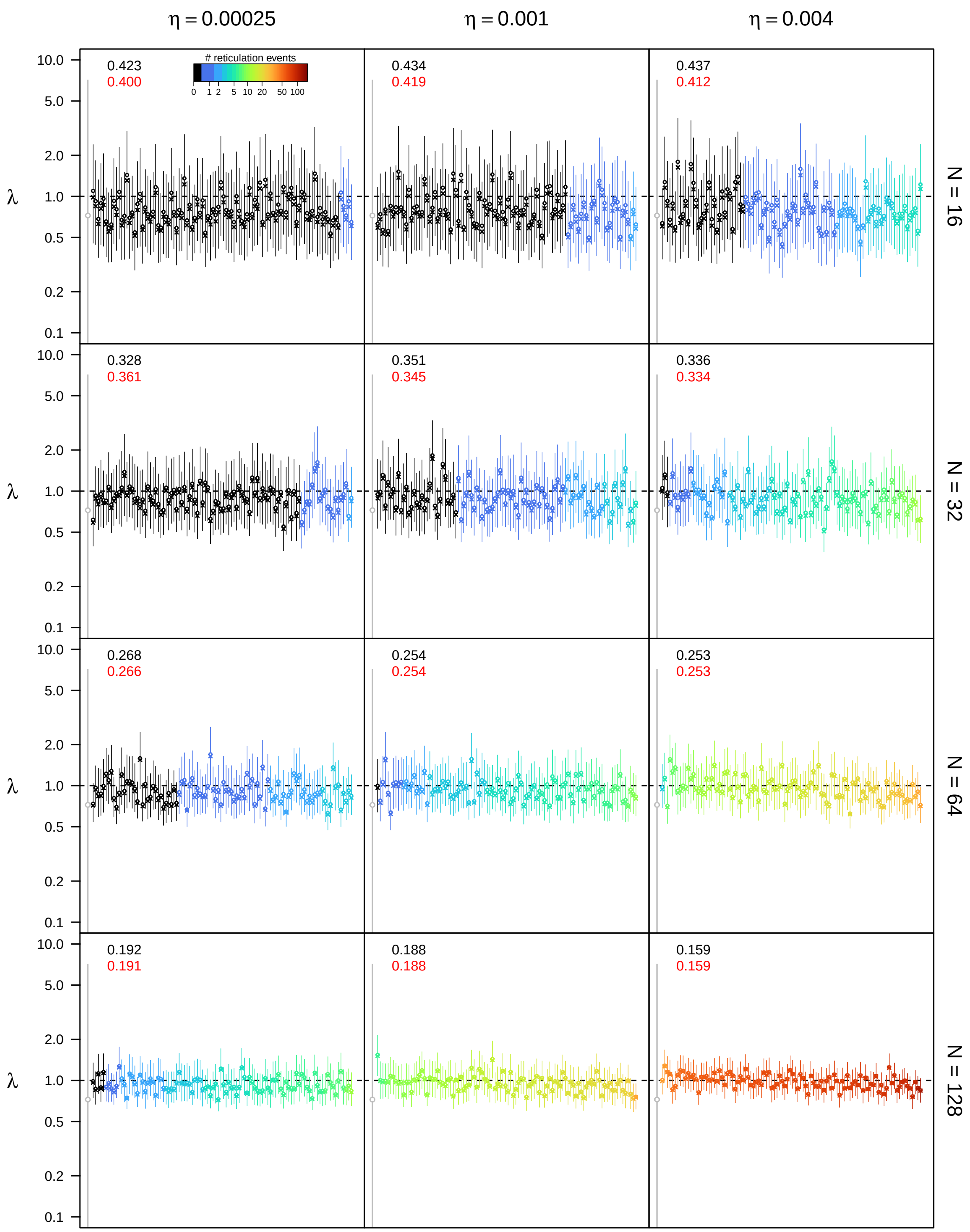


**Figure S35: Posterior estimates of the speciation rate $\lambda$ under the unidirectional hybridization model for $\epsilon = 0.5$, $\rho = 0.5$, and $u_{\mathbf{max}} = 4N$.** See main text for interpretation.

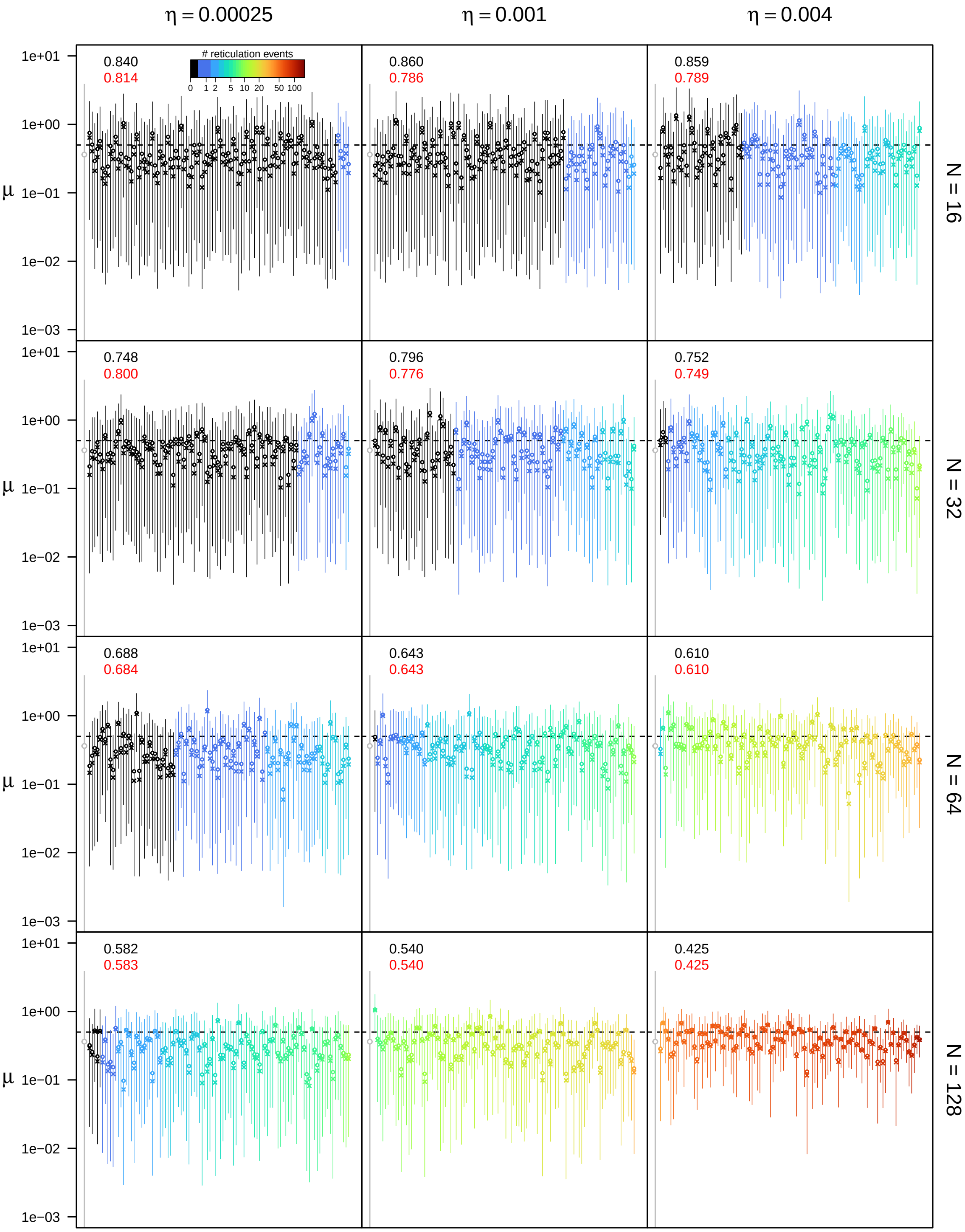


**Figure S36: Posterior estimates of the extinction rate $\mu$ under the unidirectional hybridization model for $\epsilon = 0.5$, $\rho = 0.5$, and $u_{\mathbf{max}} = 4N$.** See main text for interpretation.

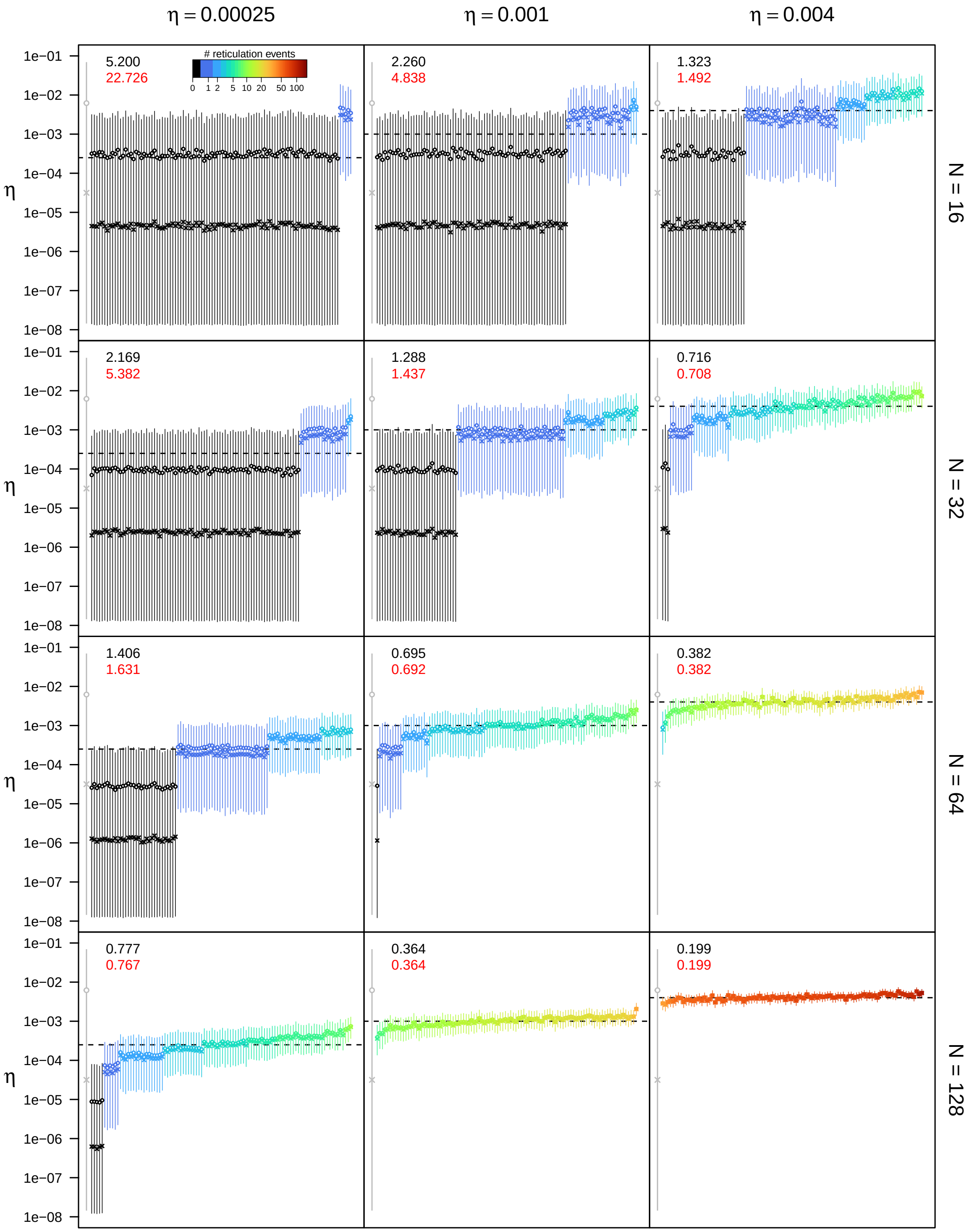


**Figure S37: Posterior estimates of the unidirectional hybridization rate $\eta$ under the unidirectional hybridization model for $\epsilon = 0.5$, $\rho = 0.5$, and $u_{\mathbf{max}} = 4N$.** See main text for interpretation.

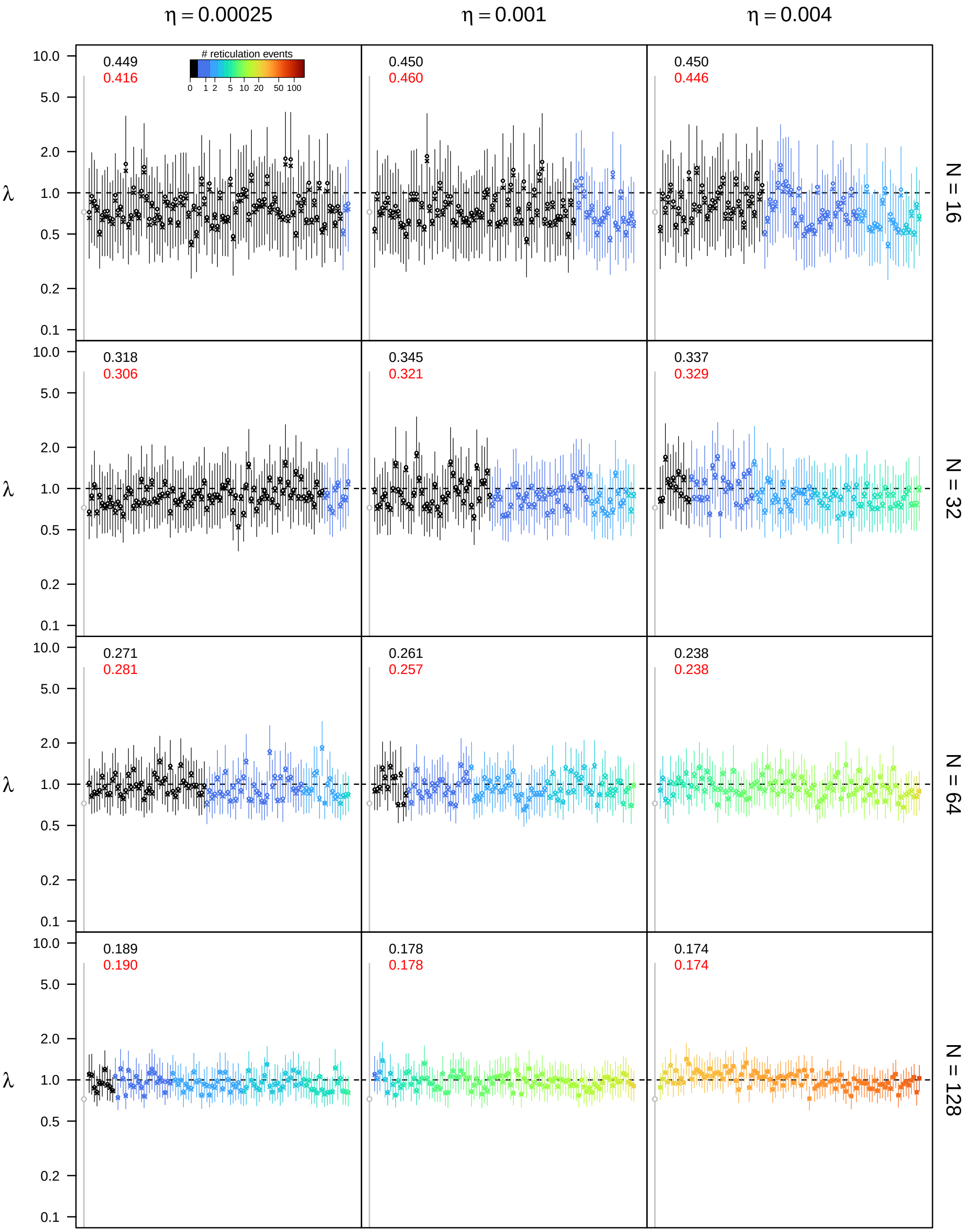


**Figure S38: Posterior estimates of the speciation rate $\lambda$ under the unidirectional hybridization model for $\epsilon = 0.5$, $\rho = 0.75$, and $u_{\mathbf{max}} = 4N$.** See main text for interpretation.

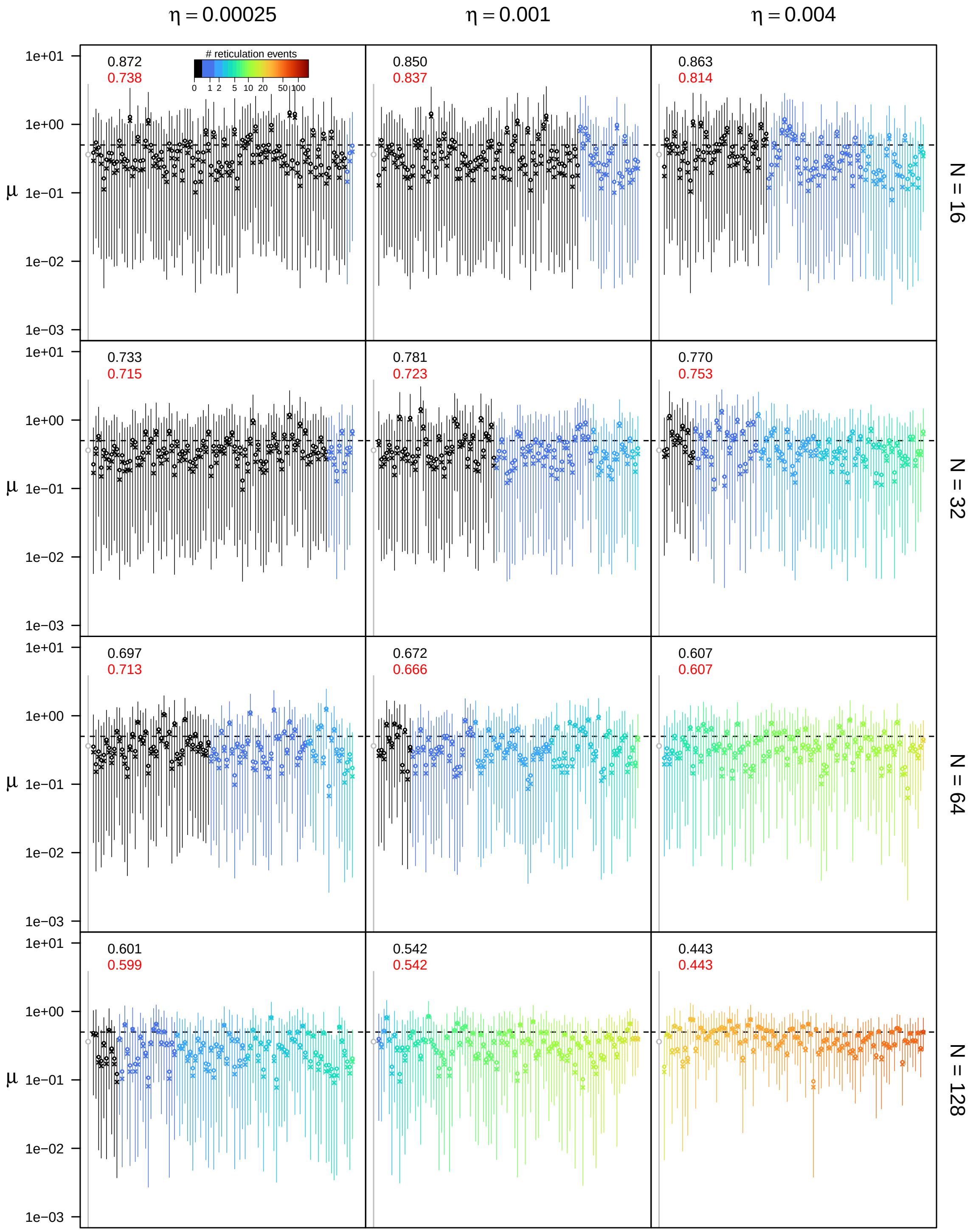


**Figure S39: Posterior estimates of the extinction rate $\mu$ under the unidirectional hybridization model for $\epsilon = 0.5$, $\rho = 0.75$, and $u_{\mathbf{max}} = 4N$.** See main text for interpretation.

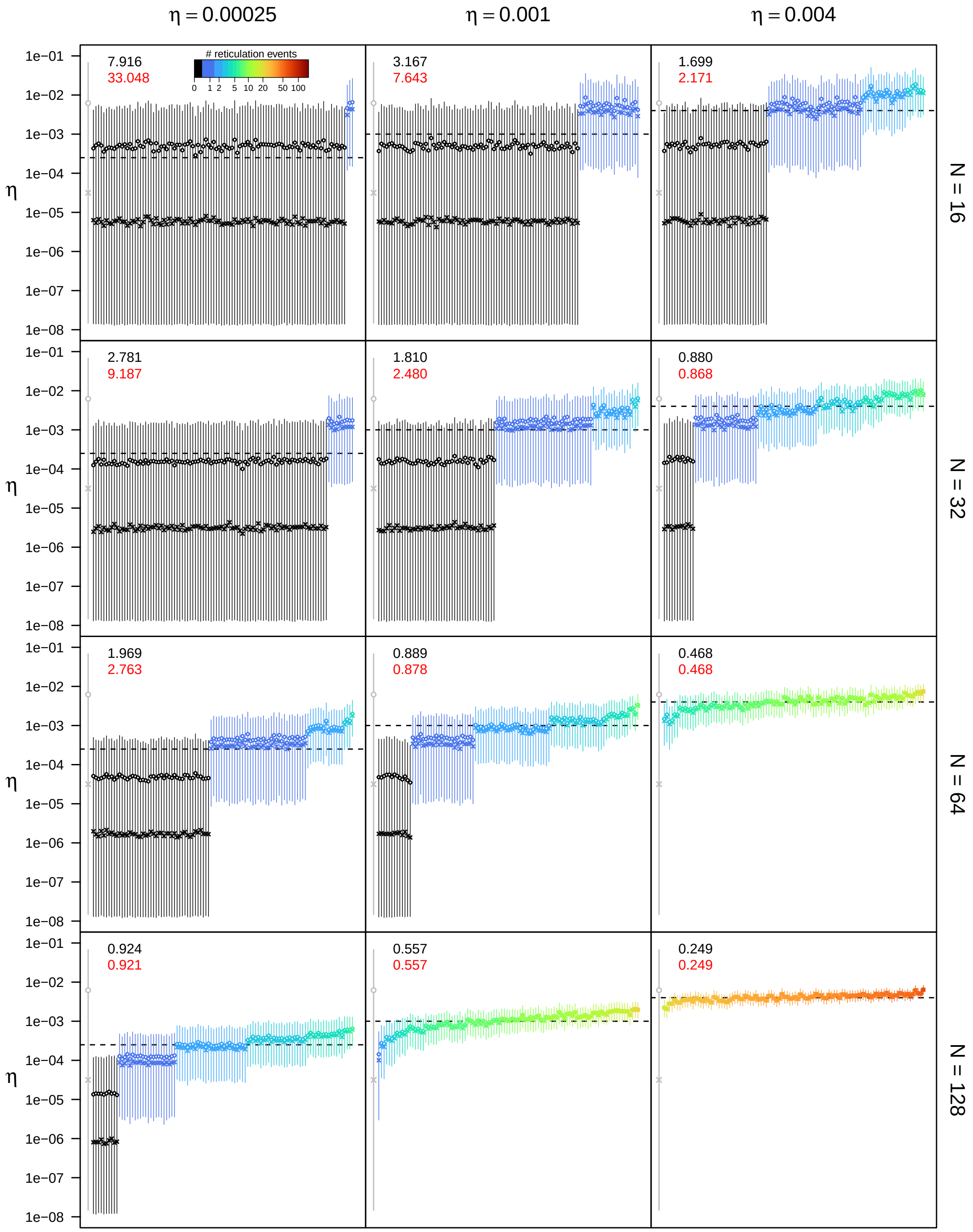


**Figure S40: Posterior estimates of the unidirectional hybridization rate $\eta$ under the unidirectional hybridization model for $\epsilon = 0.5$, $\rho = 0.75$, and $u_{\mathbf{max}} = 4N$.** See main text for interpretation.

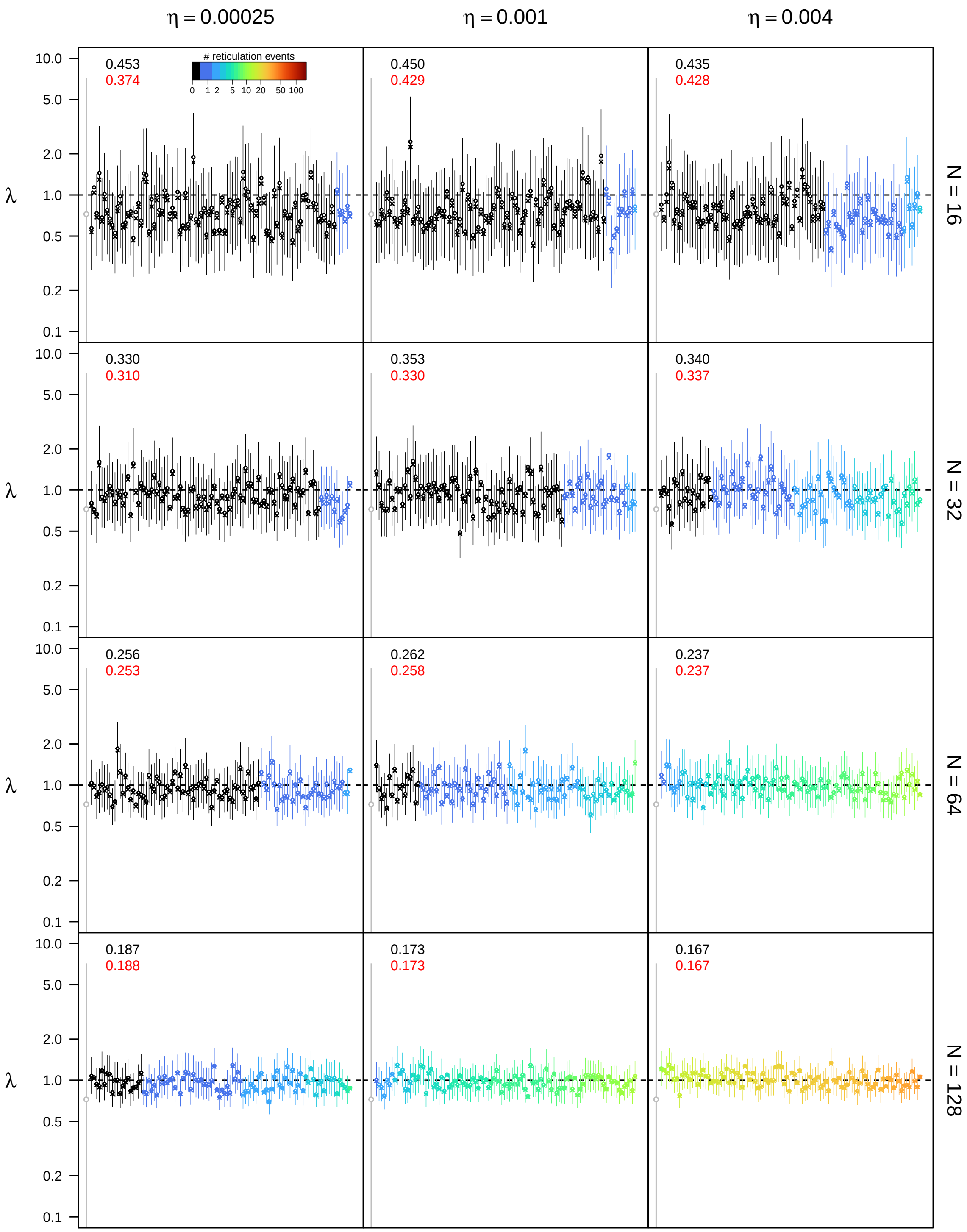


**Figure S41: Posterior estimates of the speciation rate $\lambda$ under the unidirectional hybridization model for $\epsilon = 0.5$, $\rho = 1.0$, and $u_{\mathbf{max}} = 4N$.** See main text for interpretation.

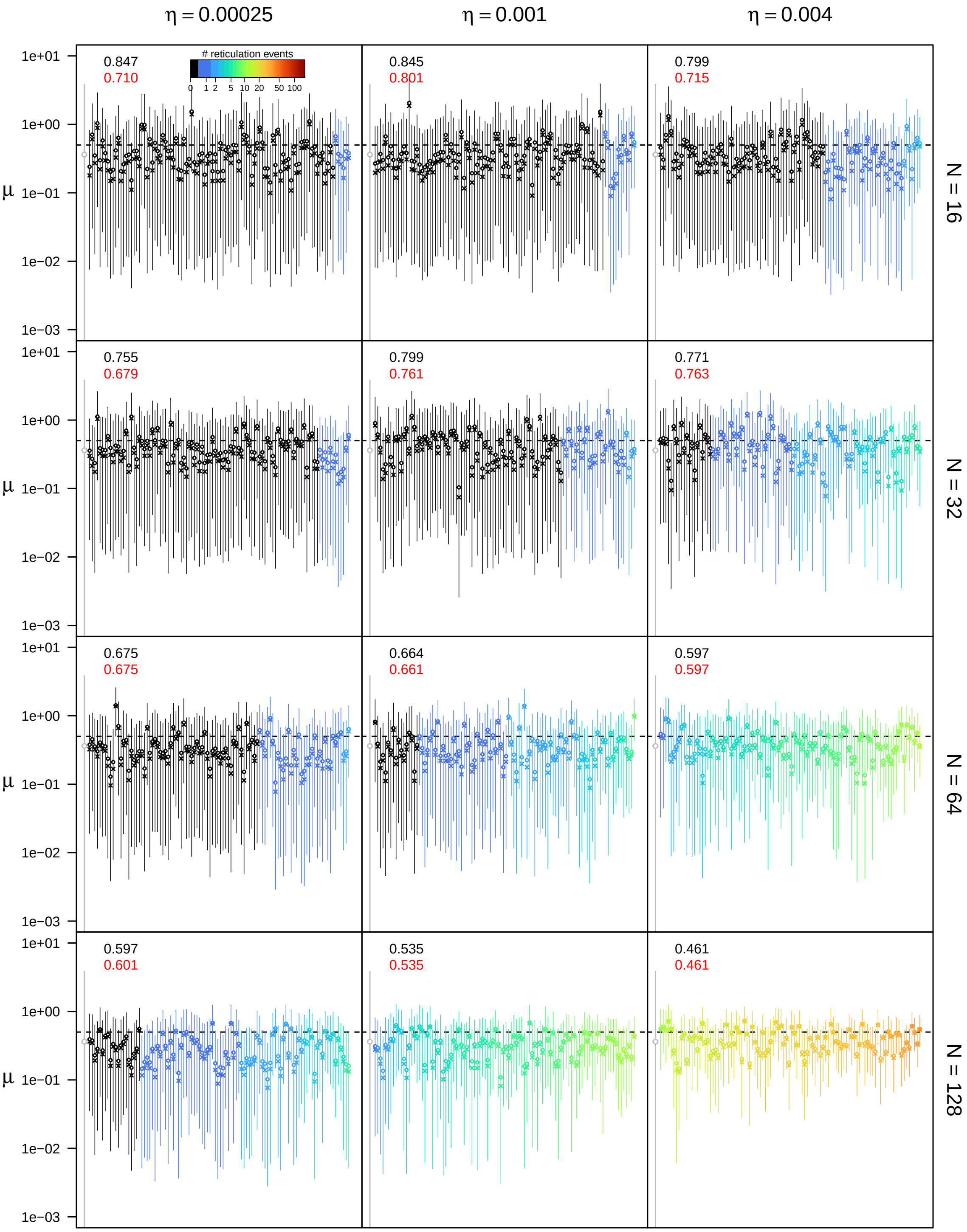


**Figure S42: Posterior estimates of the extinction rate $\mu$ under the unidirectional hybridization model for $\epsilon = 0.5$, $\rho = 1.0$, and $u_{\mathbf{max}} = 4N$.** See main text for interpretation.

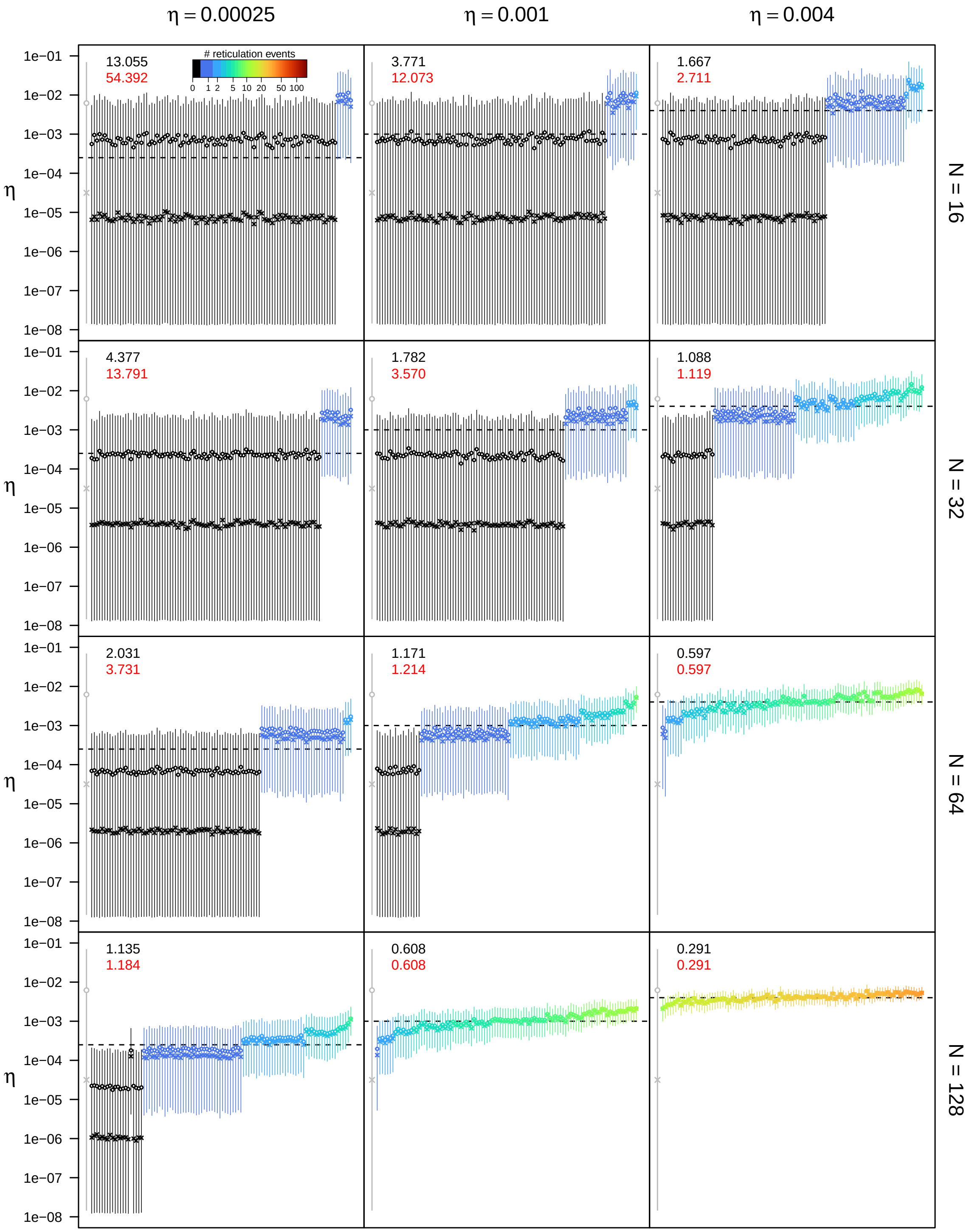


**Figure S43: Posterior estimates of the unidirectional hybridization rate $\eta$ under the unidirectional hybridization model for $\epsilon = 0.5$, $\rho = 1.0$, and $u_{\max} = 4N$.** See main text for interpretation.

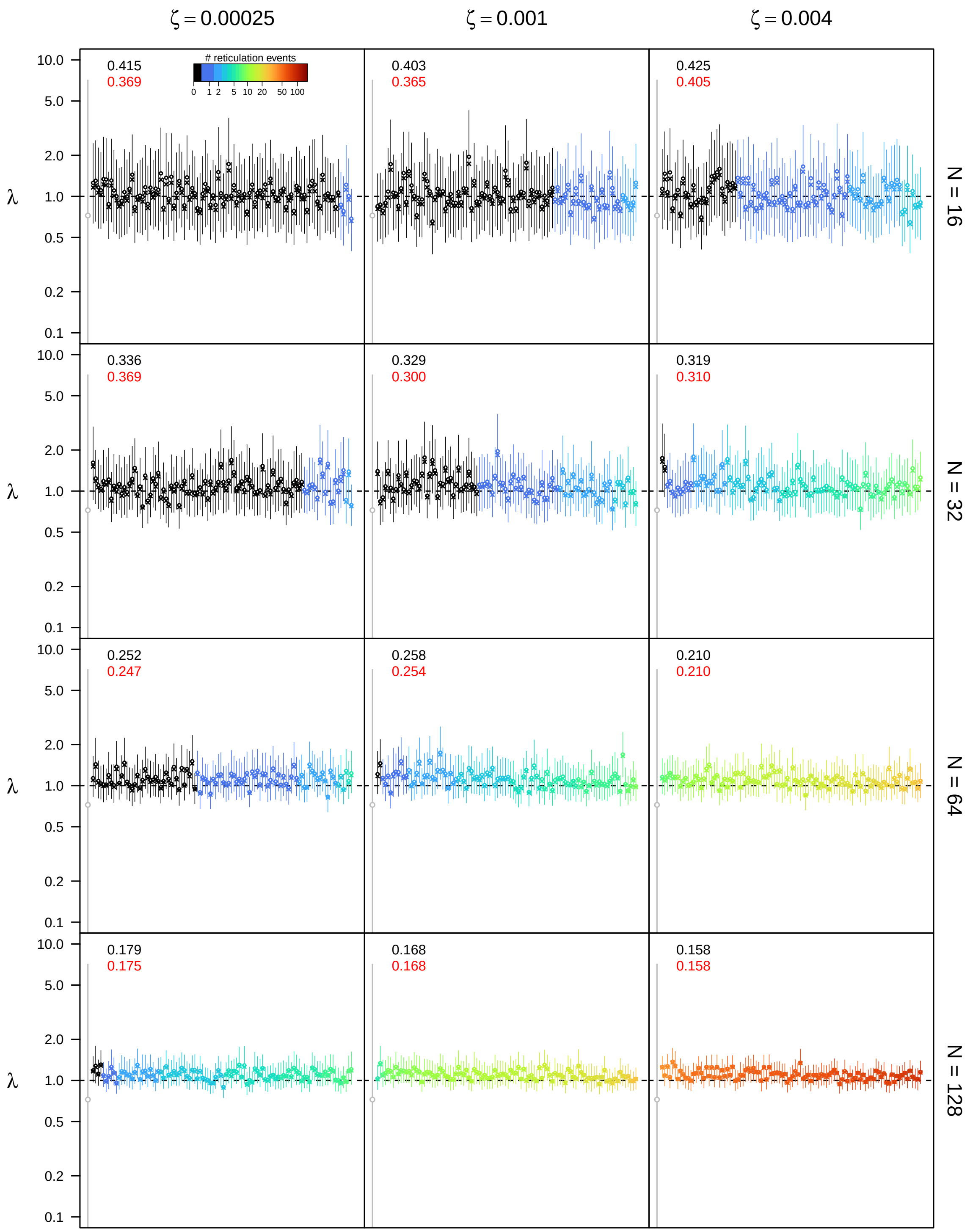


**Figure S44: Posterior estimates of the speciation rate $\lambda$ under the bidirectional hybridization model for $\epsilon = 0.1$, $\rho = 0.5$, and $u_{\mathbf{max}} = 4N$.** See main text for interpretation.

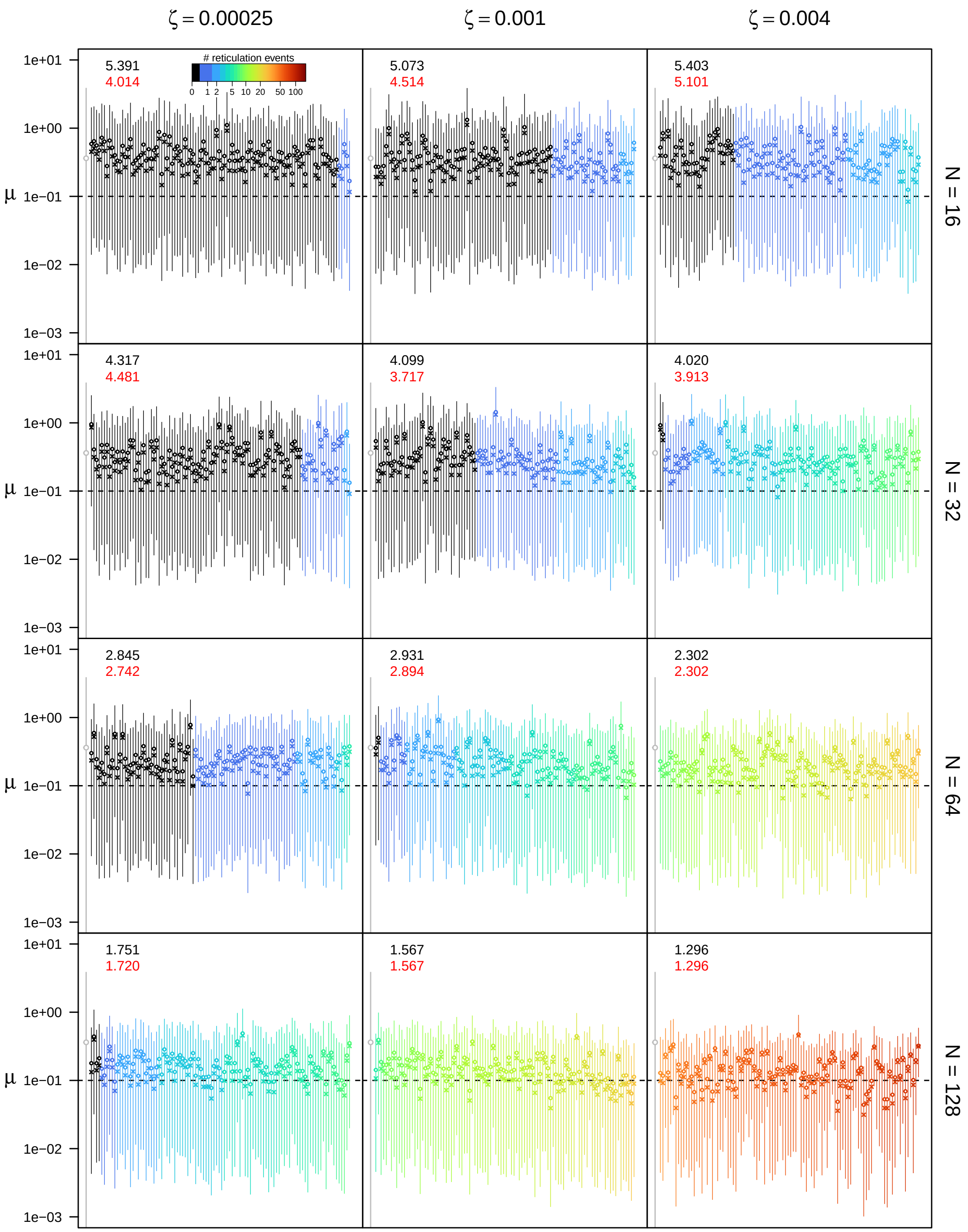


**Figure S45: Posterior estimates of the extinction rate $\mu$ under the bidirectional hybridization model for $\epsilon = 0.1$, $\rho = 0.5$, and $u_{\text{max}} = 4N$.** See main text for interpretation.

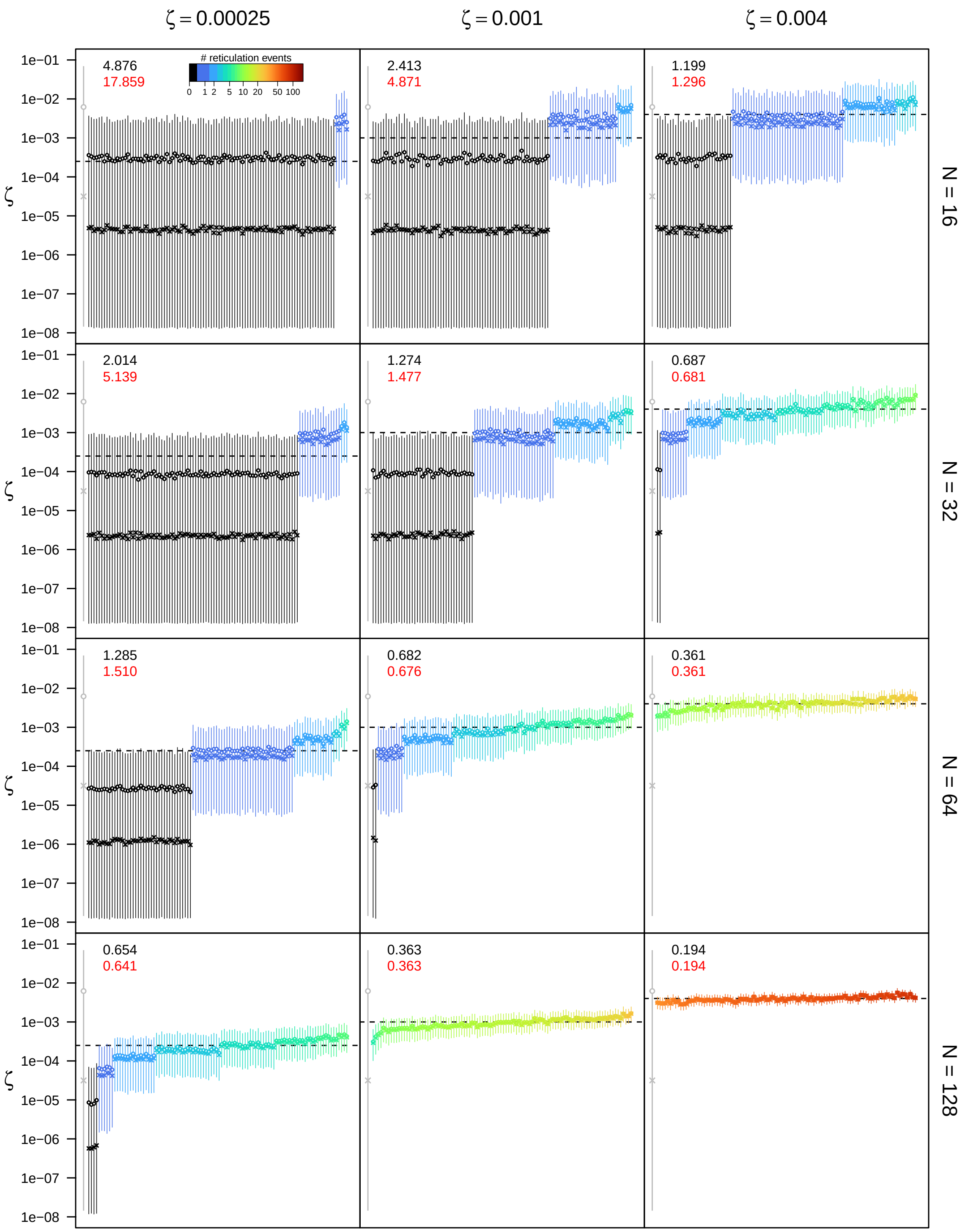


**Figure S46: Posterior estimates of the bidirectional hybridization rate $\zeta$ under the bidirectional hybridization model for $\epsilon = 0.1$, $\rho = 0.5$, and $u_{\mathbf{max}} = 4N$.** See main text for interpretation.

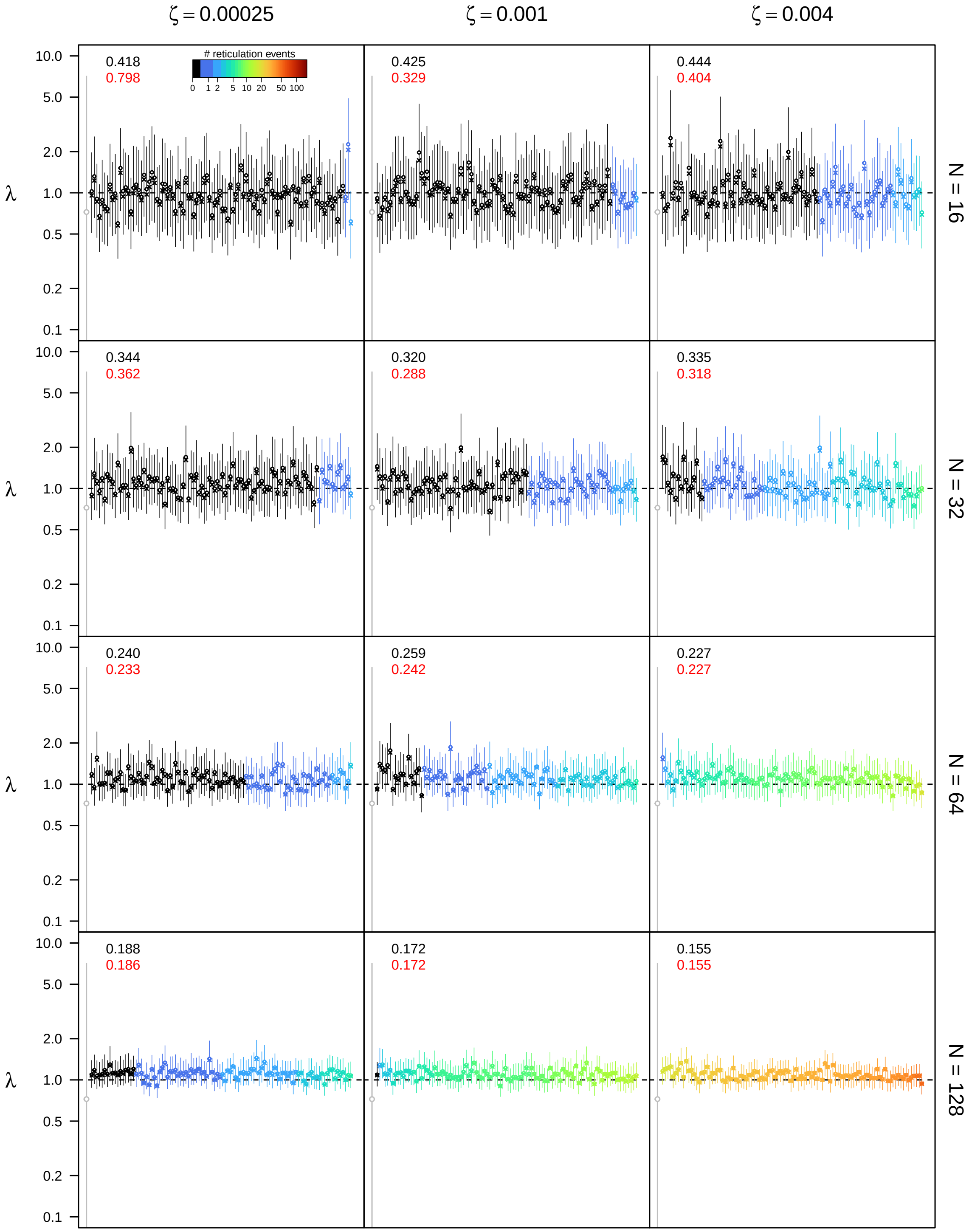


**Figure S47: Posterior estimates of the speciation rate $\lambda$ under the bidirectional hybridization model for $\epsilon = 0.1$, $\rho = 0.75$, and $u_{\mathbf{max}} = 4N$.** See main text for interpretation.

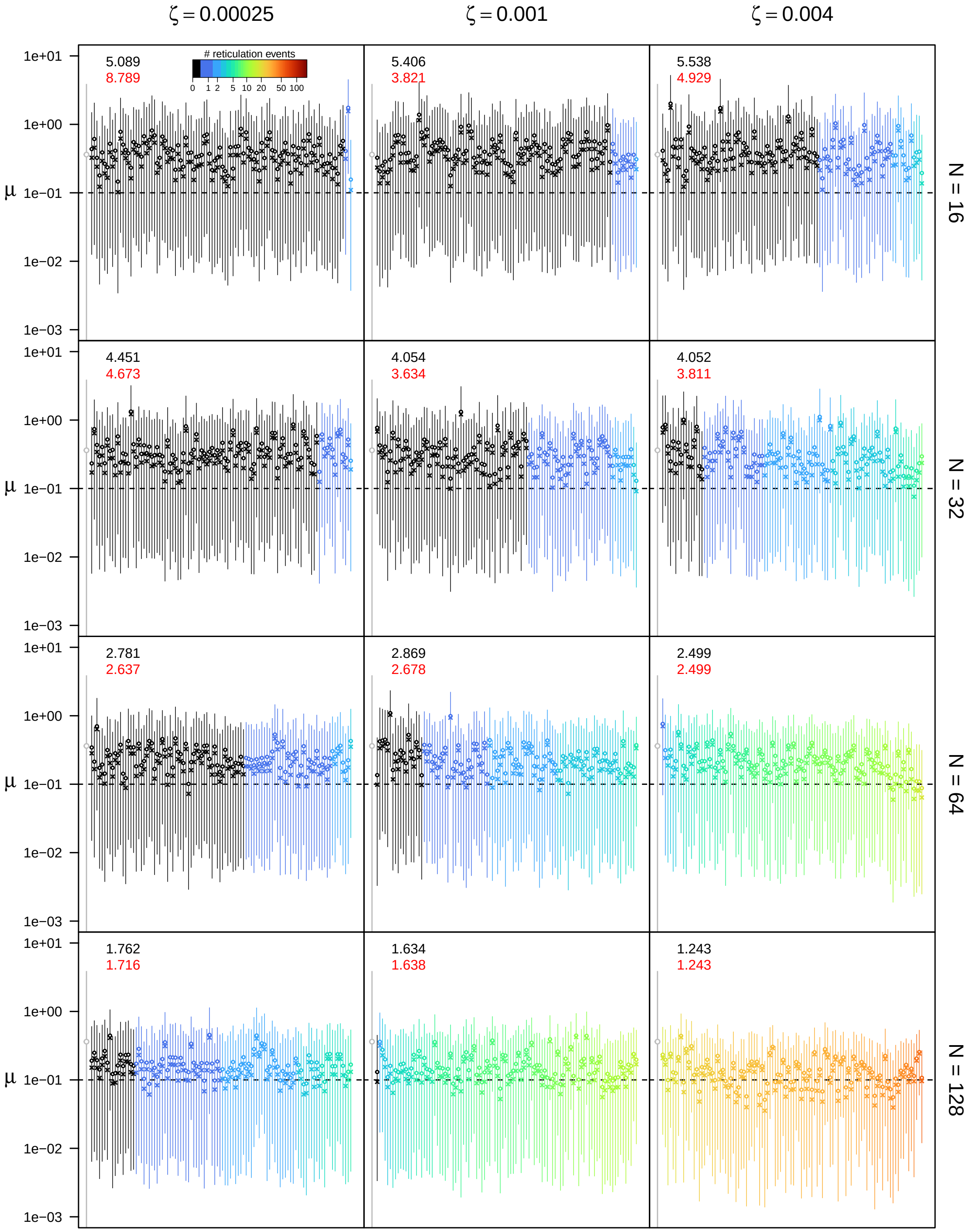


**Figure S48: Posterior estimates of the extinction rate $\mu$ under the bidirectional hybridization model for $\epsilon = 0.1$, $\rho = 0.75$, and $u_{\max} = 4N$.** See main text for interpretation.

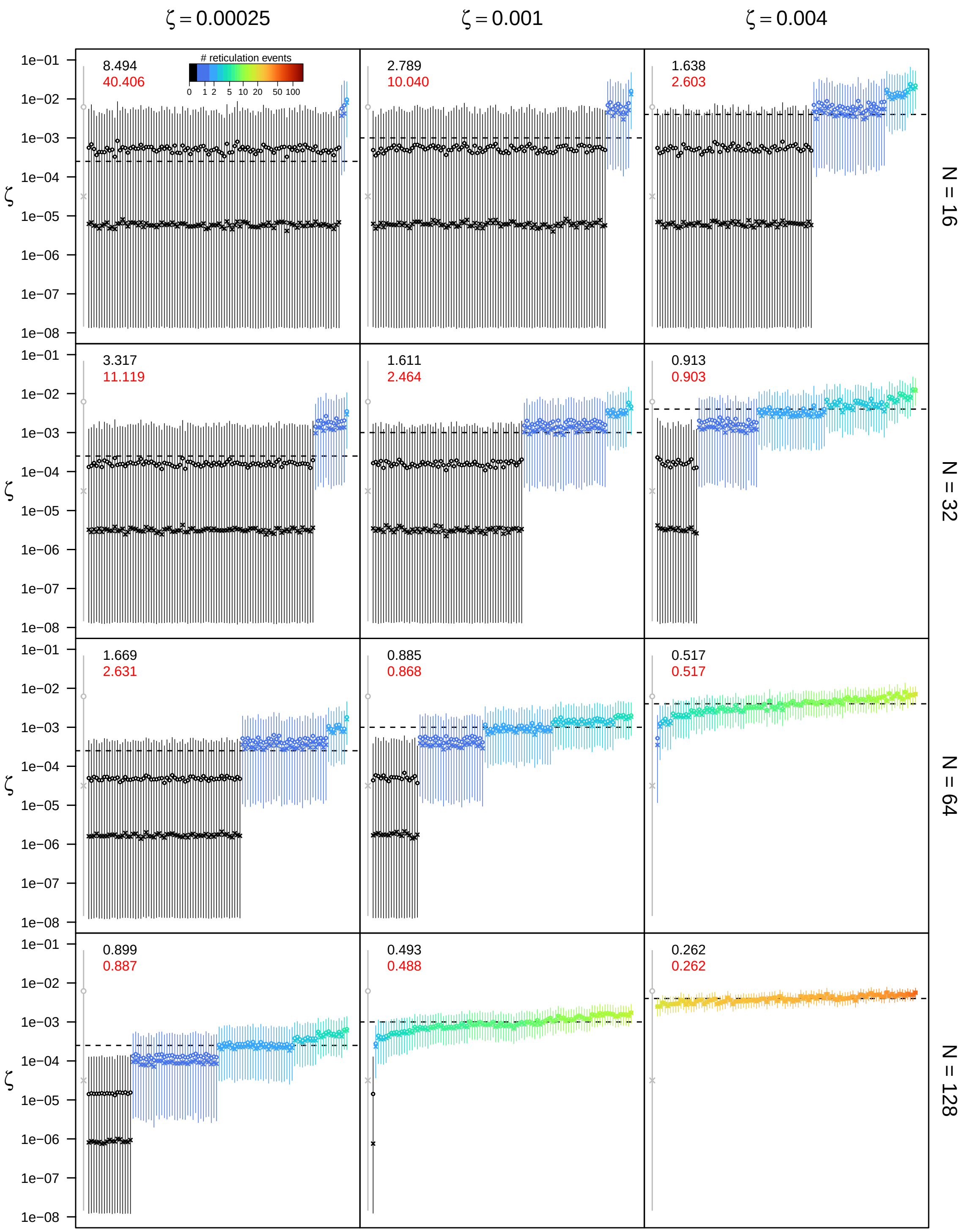


**Figure S49: Posterior estimates of the bidirectional hybridization rate $\zeta$ under the bidirectional hybridization model for $\epsilon = 0.1$, $\rho = 0.75$, and $u_{\mathbf{max}} = 4N$.** See main text for interpretation.

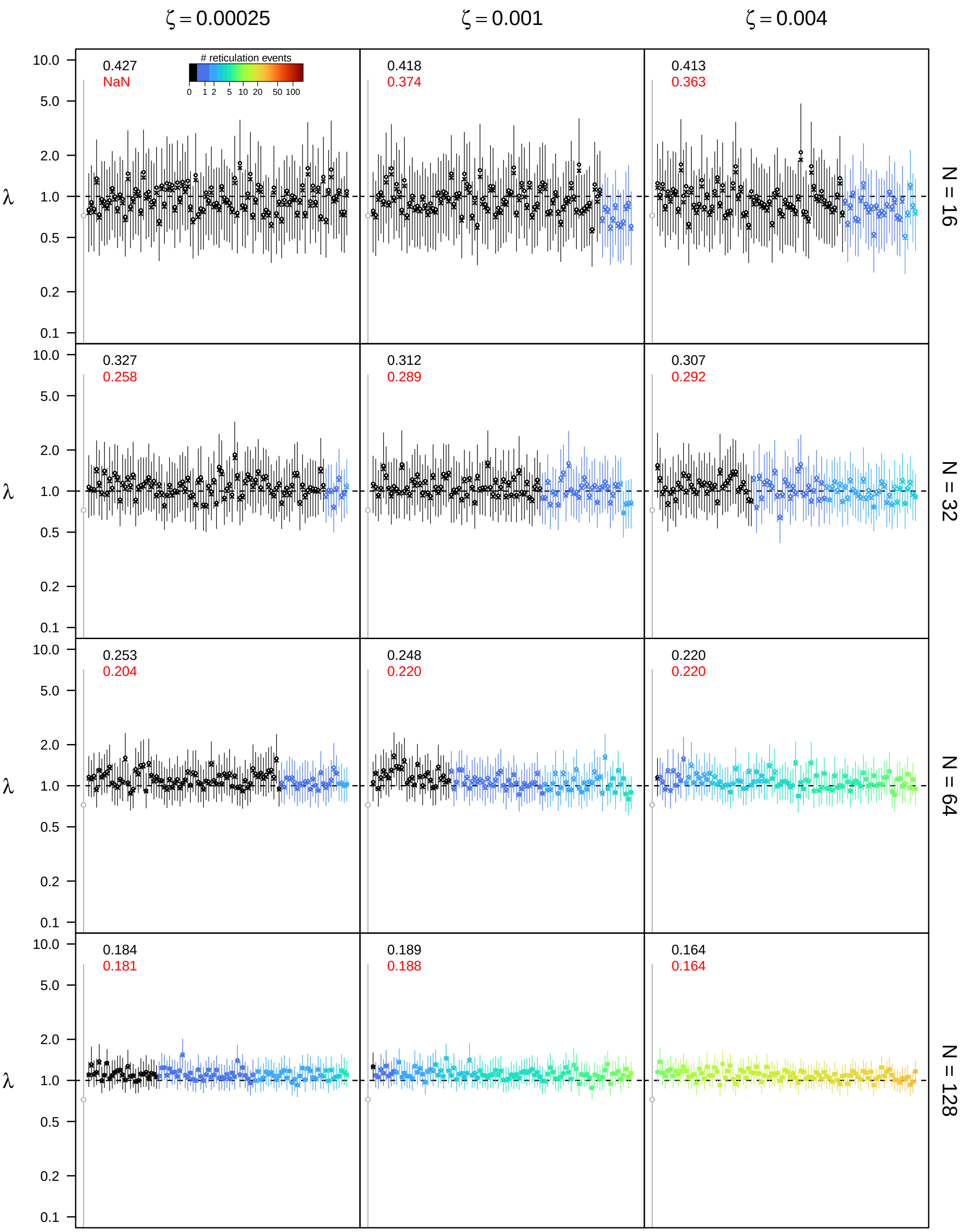


**Figure S50: Posterior estimates of the speciation rate $\lambda$ under the bidirectional hybridization model for $\epsilon = 0.1$, $\rho = 1.0$, and $u_{\mathbf{max}} = 4N$.** See main text for interpretation.

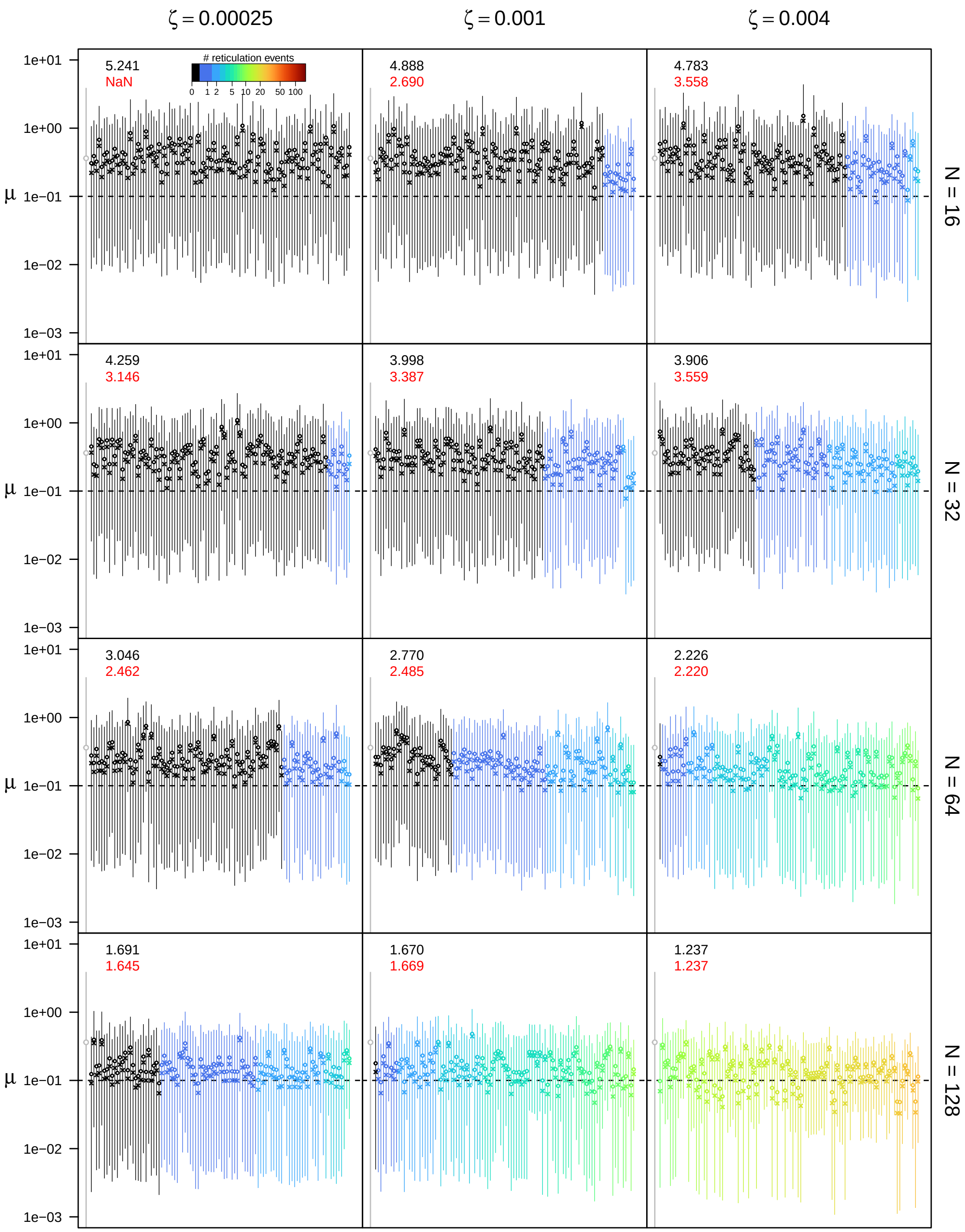


**Figure S51: Posterior estimates of the extinction rate $\mu$ under the bidirectional hybridization model for $\epsilon = 0.1$, $\rho = 1.0$, and $u_{\mathbf{max}} = 4N$.** See main text for interpretation.

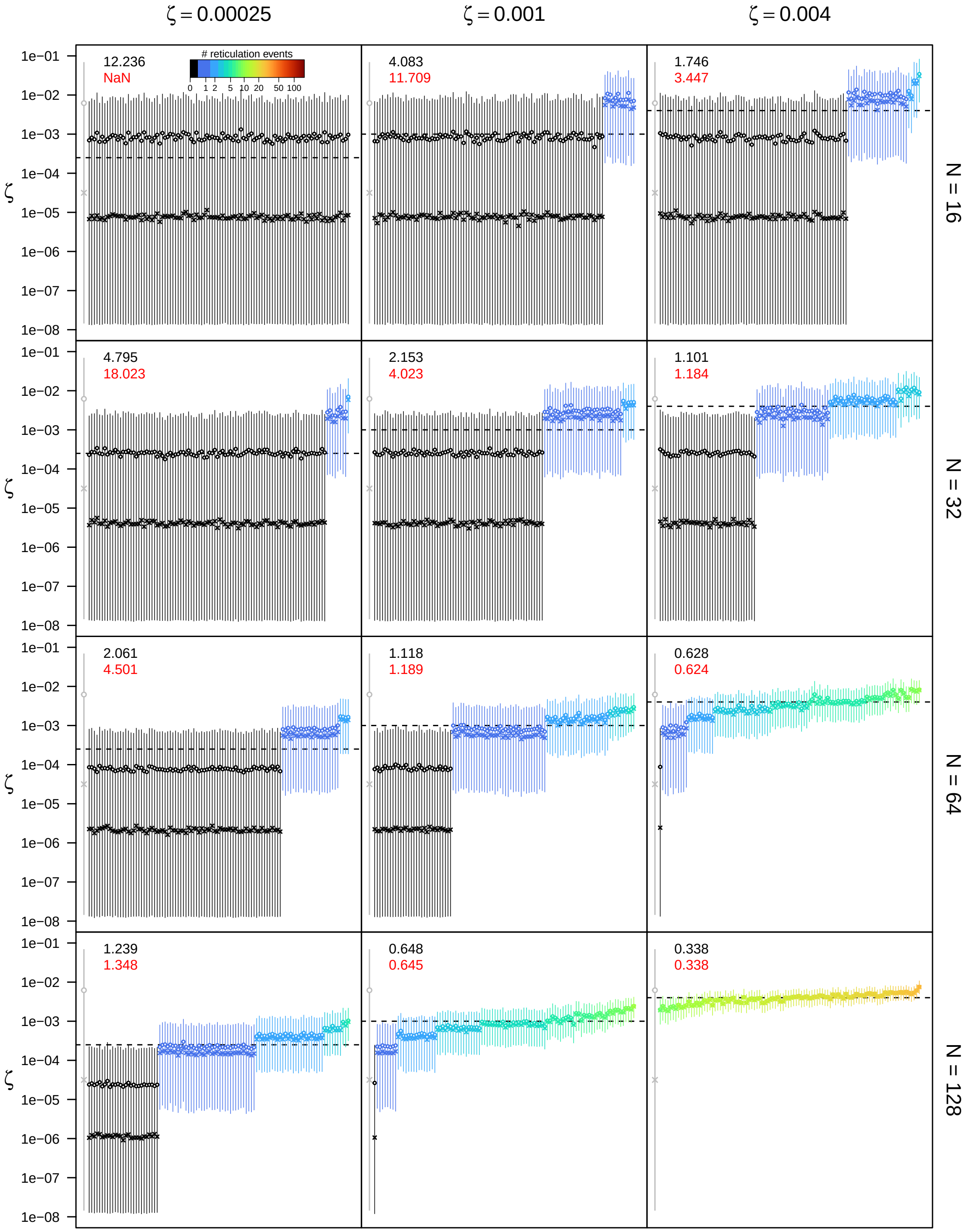


**Figure S52: Posterior estimates of the bidirectional hybridization rate $\zeta$ under the bidirectional hybridization model for $\epsilon = 0.1$, $\rho = 1.0$, and $u_{\mathbf{max}} = 4N$.** See main text for interpretation.

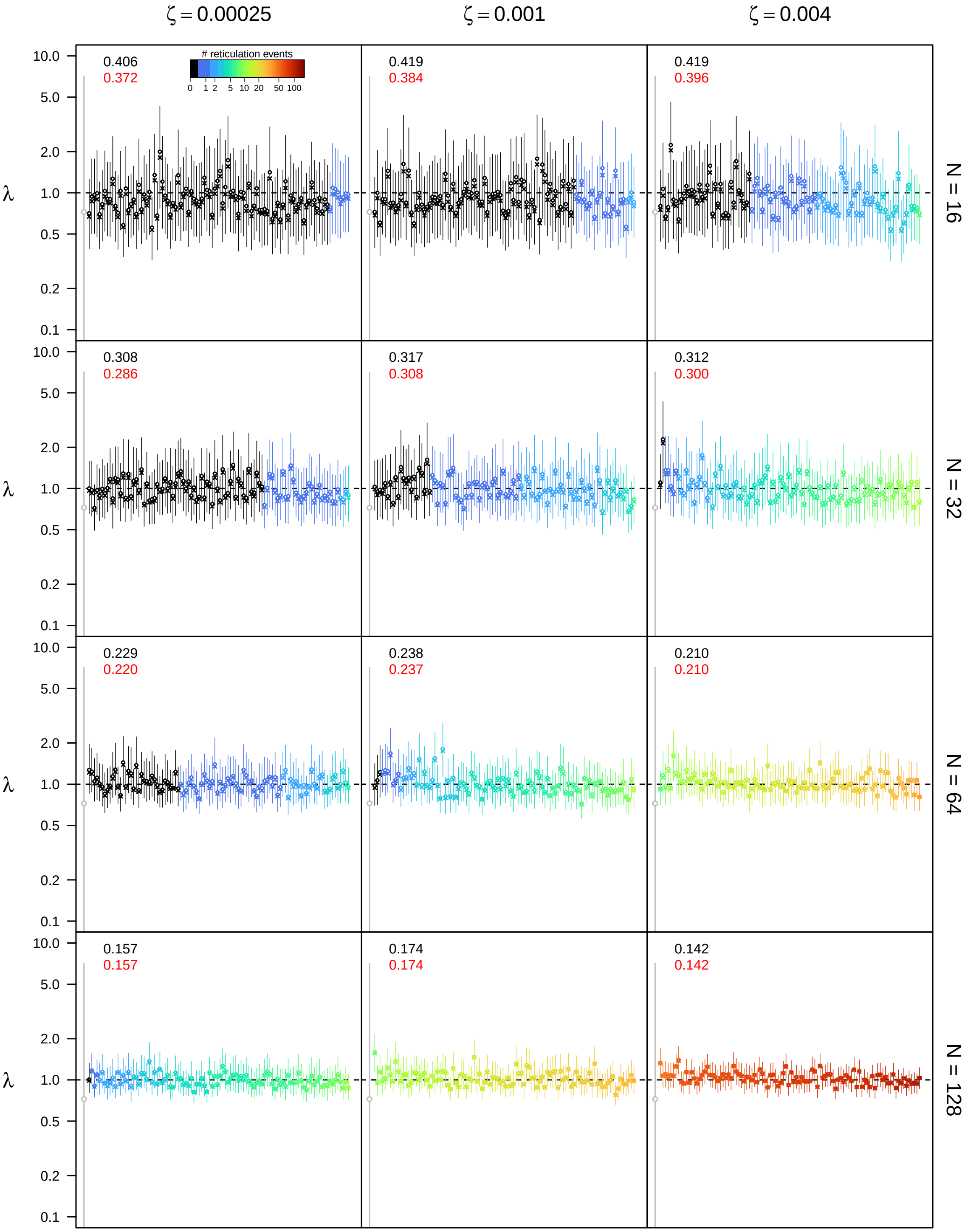


**Figure S53: Posterior estimates of the speciation rate $\lambda$ under the bidirectional hybridization model for $\epsilon = 0.3$, $\rho = 0.5$, and $u_{\max} = 4N$.** See main text for interpretation.

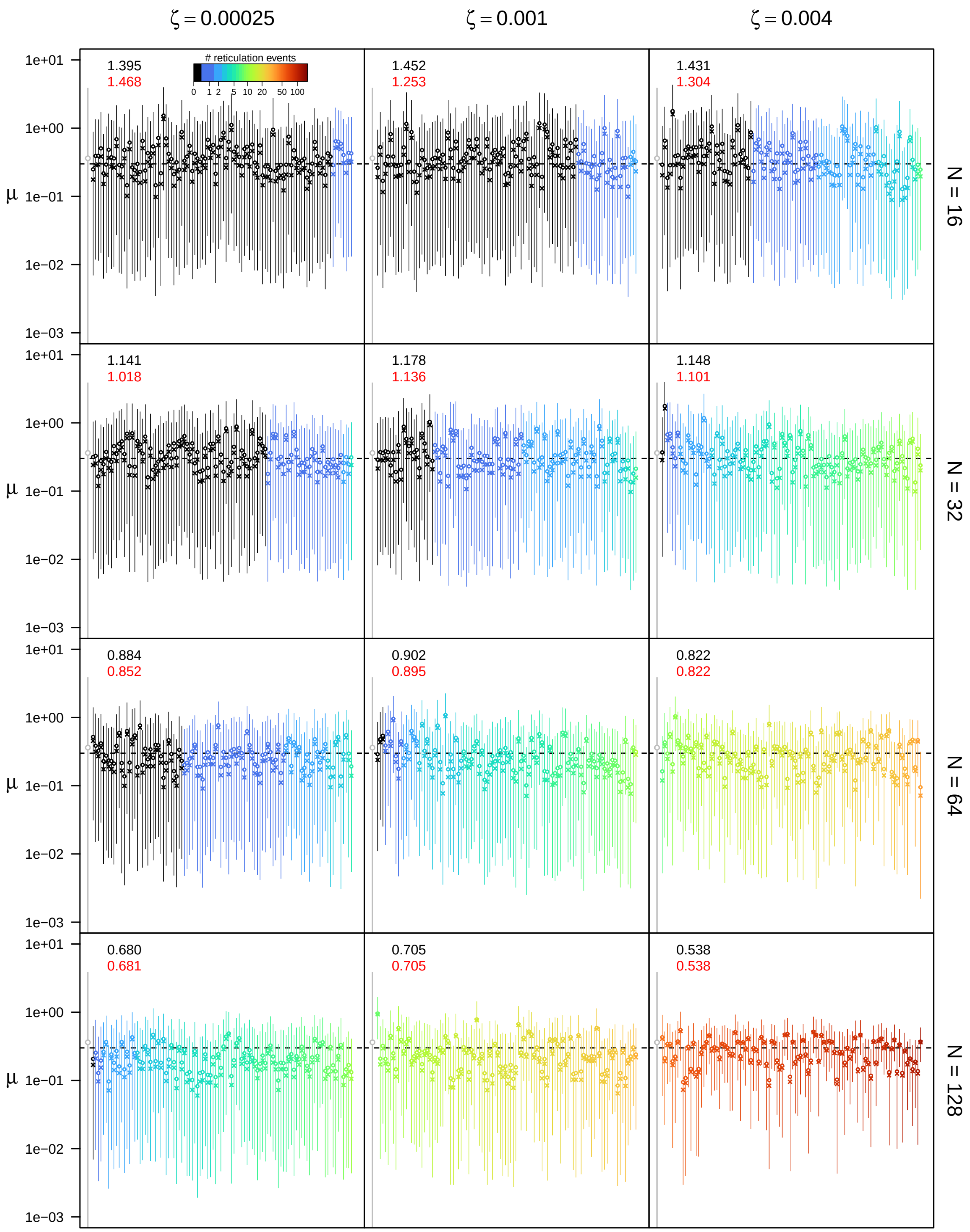


**Figure S54: Posterior estimates of the extinction rate $\mu$ under the bidirectional hybridization model for $\epsilon = 0.3$, $\rho = 0.5$, and $u_{\text{max}} = 4N$.** See main text for interpretation.

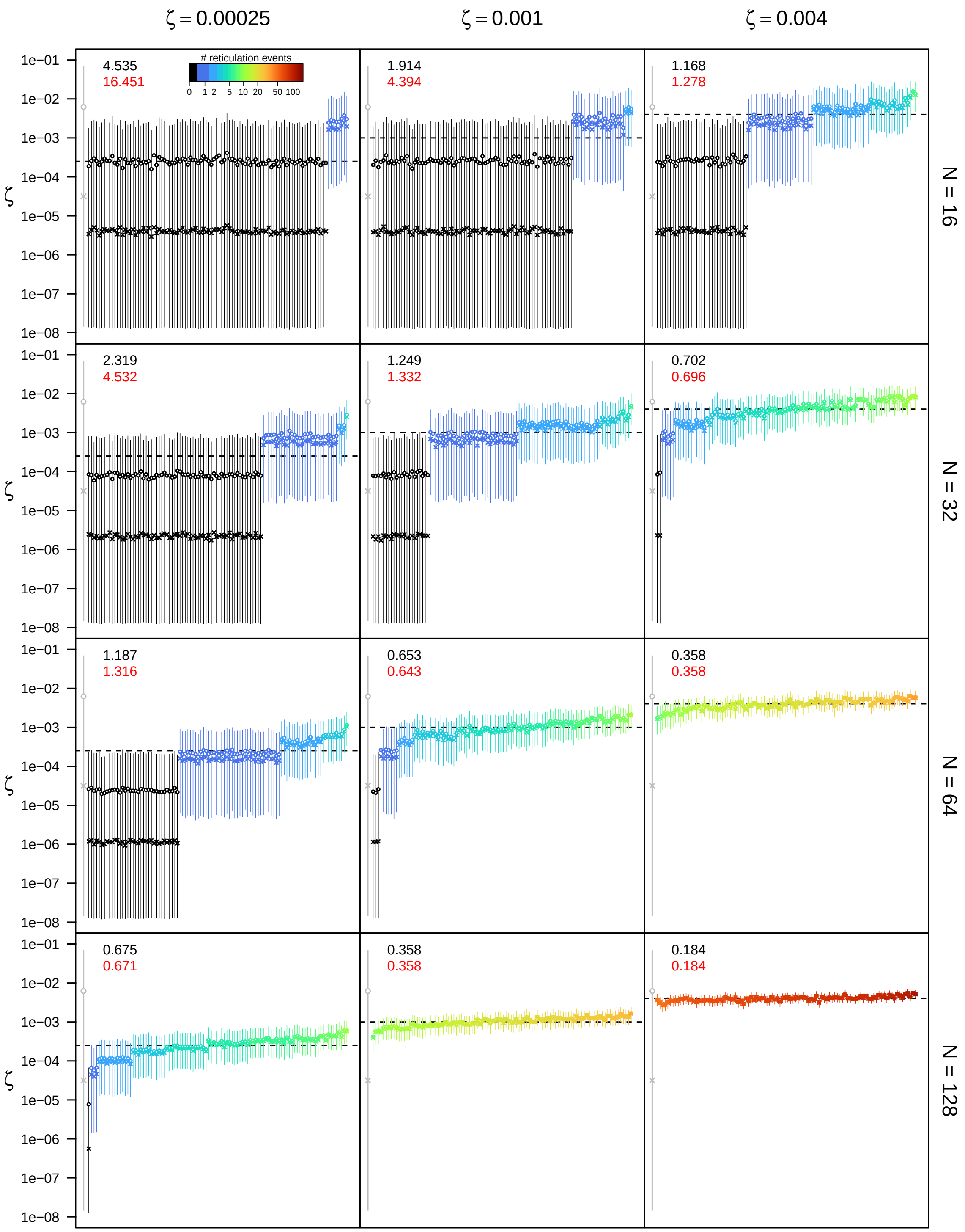


**Figure S55: Posterior estimates of the bidirectional hybridization rate $\zeta$ under the bidirectional hybridization model for** $\epsilon = 0.3$**,** $\rho = 0.5$**, and** $u_{\mathbf{max}} = 4N$**.** See main text for interpretation.

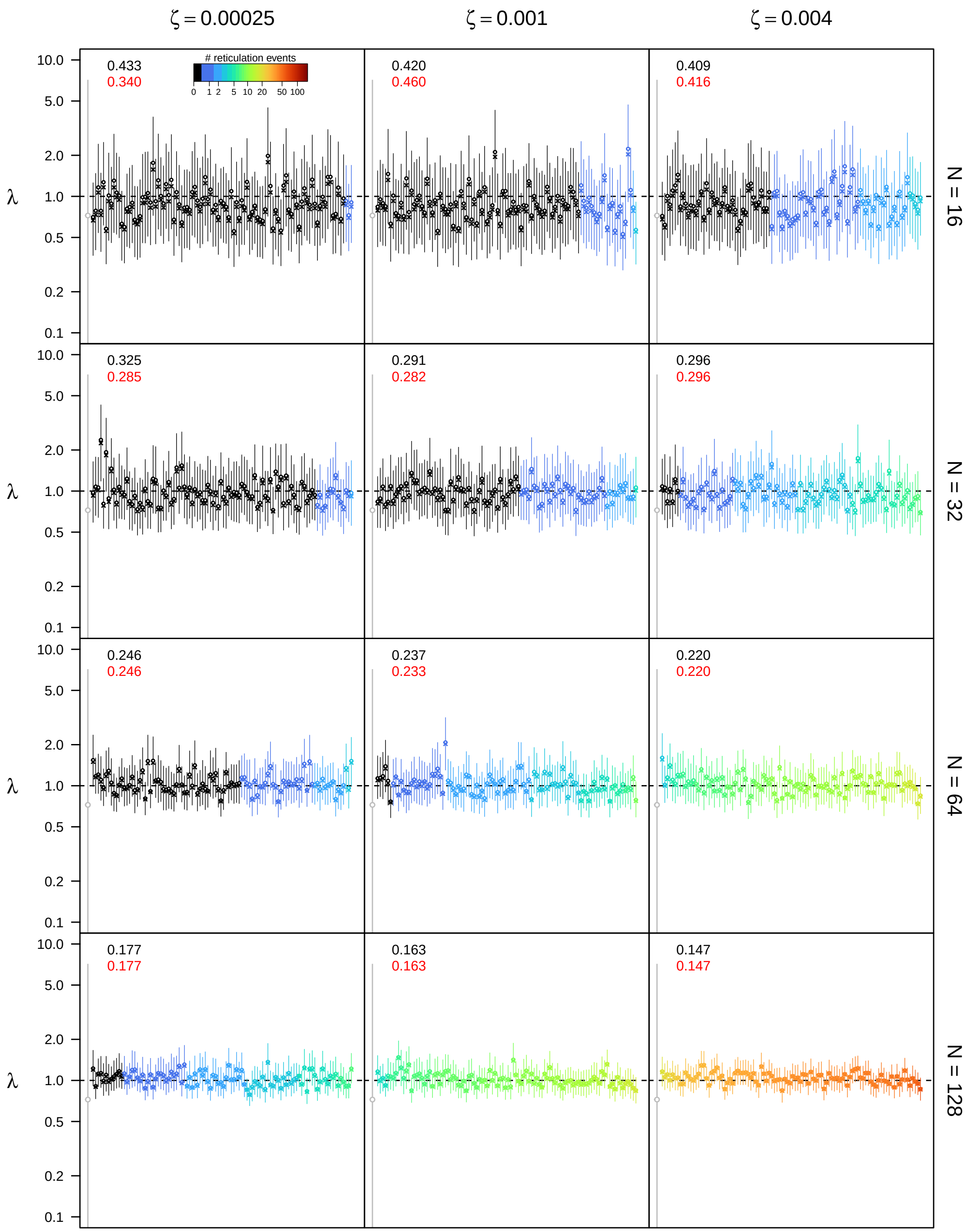


**Figure S56: Posterior estimates of the speciation rate $\lambda$ under the bidirectional hybridization model for $\epsilon = 0.3$, $\rho = 0.75$, and $u_{\max} = 4N$.** See main text for interpretation.

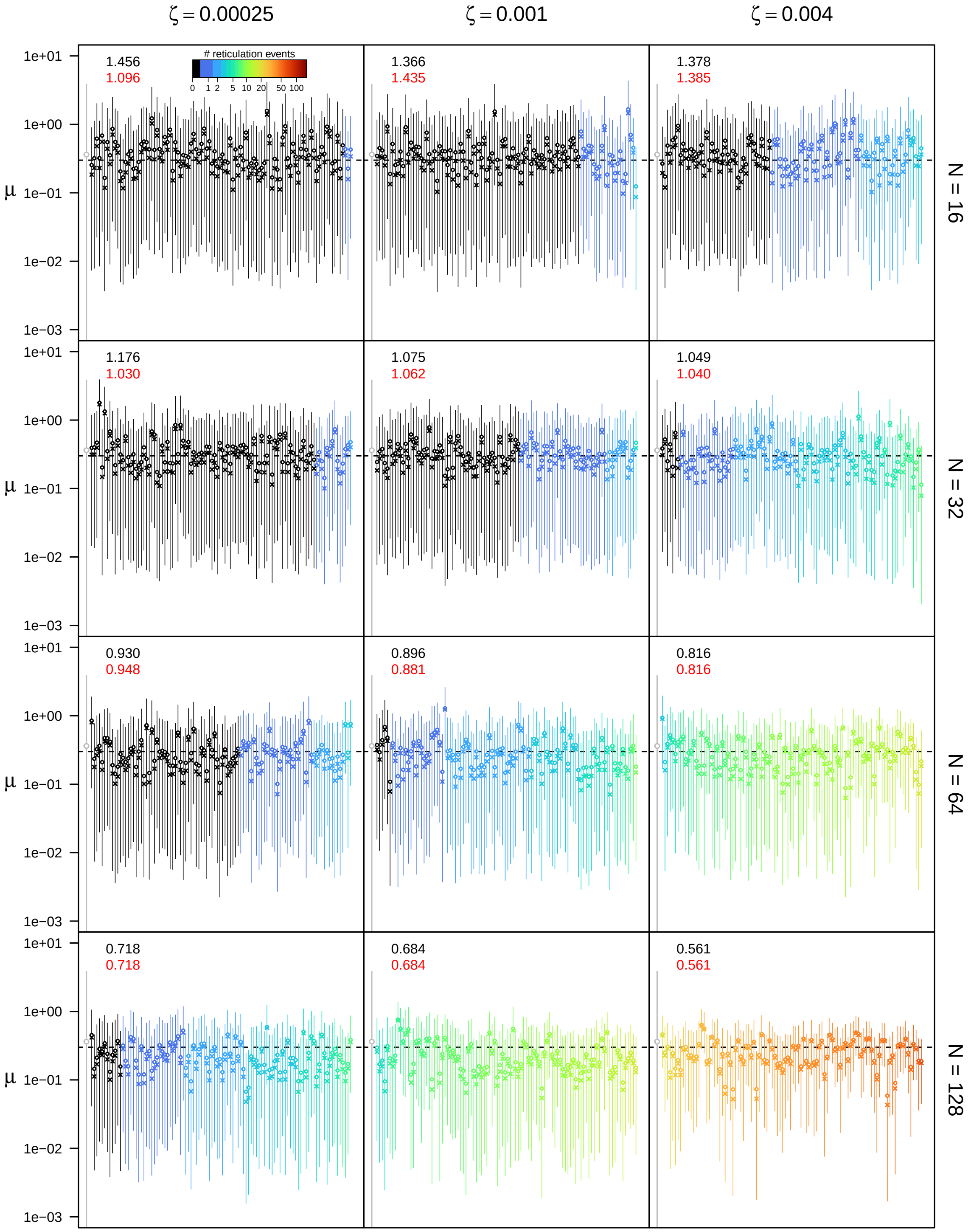


**Figure S57: Posterior estimates of the extinction rate $\mu$ under the bidirectional hybridization model for $\epsilon = 0.3$, $\rho = 0.75$, and $u_{\mathbf{max}} = 4N$.** See main text for interpretation.

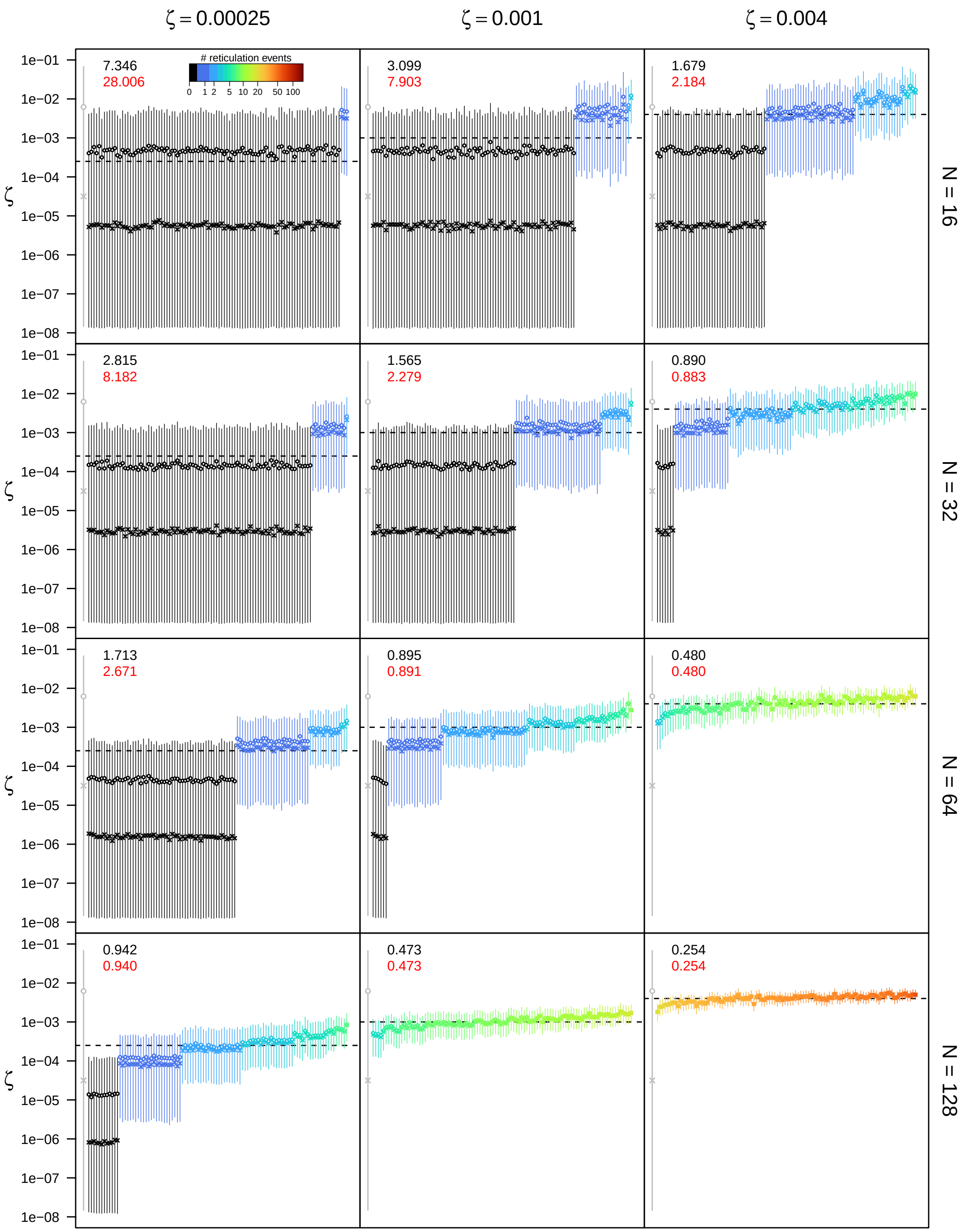


**Figure S58: Posterior estimates of the bidirectional hybridization rate $\zeta$ under the bidirectional hybridization model for $\epsilon = 0.3$, $\rho = 0.75$, and $u_{\mathbf{max}} = 4N$.** See main text for interpretation.

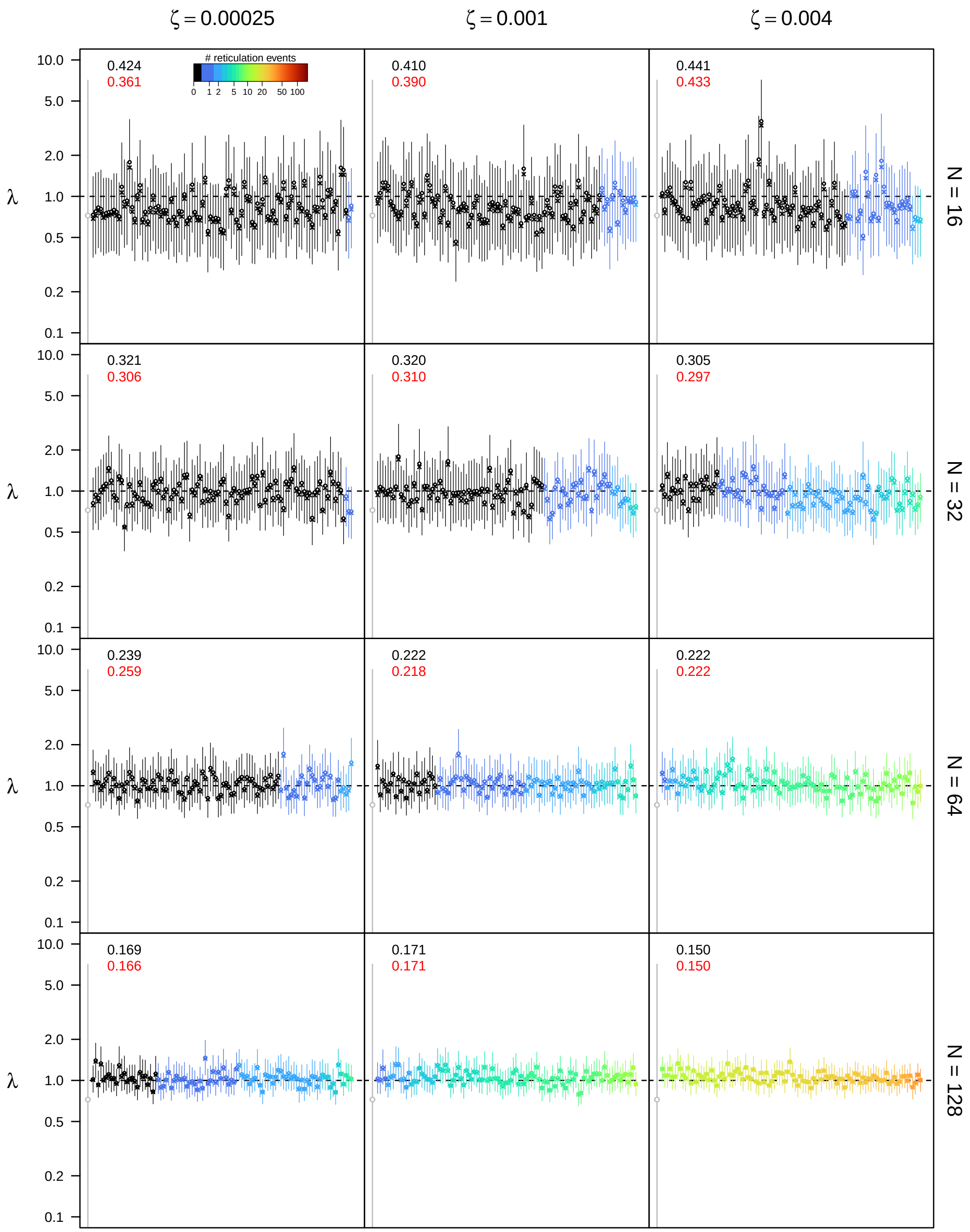


**Figure S59: Posterior estimates of the speciation rate $\lambda$ under the bidirectional hybridization model for $\epsilon = 0.3$, $\rho = 1.0$, and $u_{\textbf{max}} = 4N$.** See main text for interpretation.

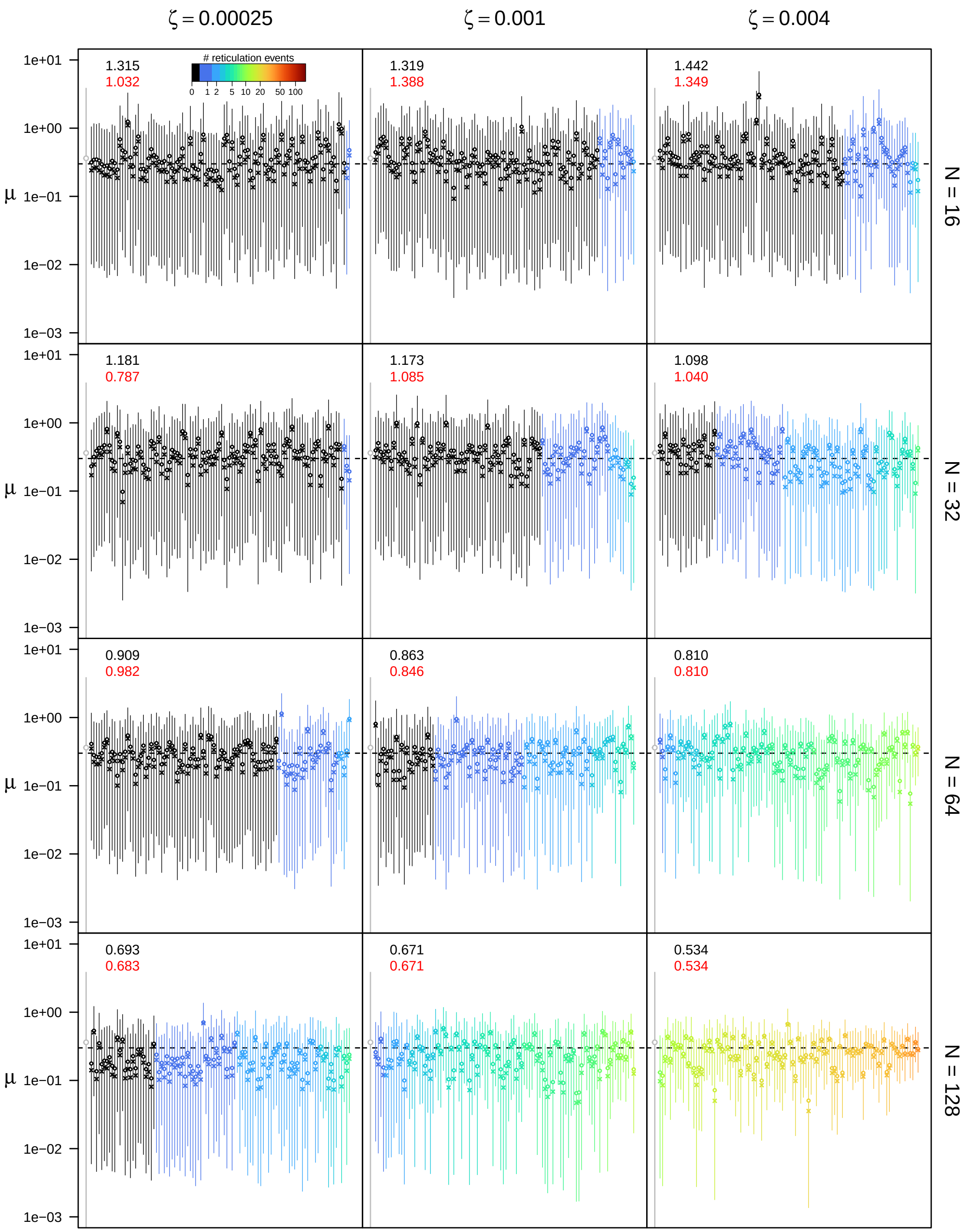


**Figure S60: Posterior estimates of the extinction rate $\mu$ under the bidirectional hybridization model for $\epsilon = 0.3$, $\rho = 1.0$, and $u_{\mathbf{max}} = 4N$.** See main text for interpretation.

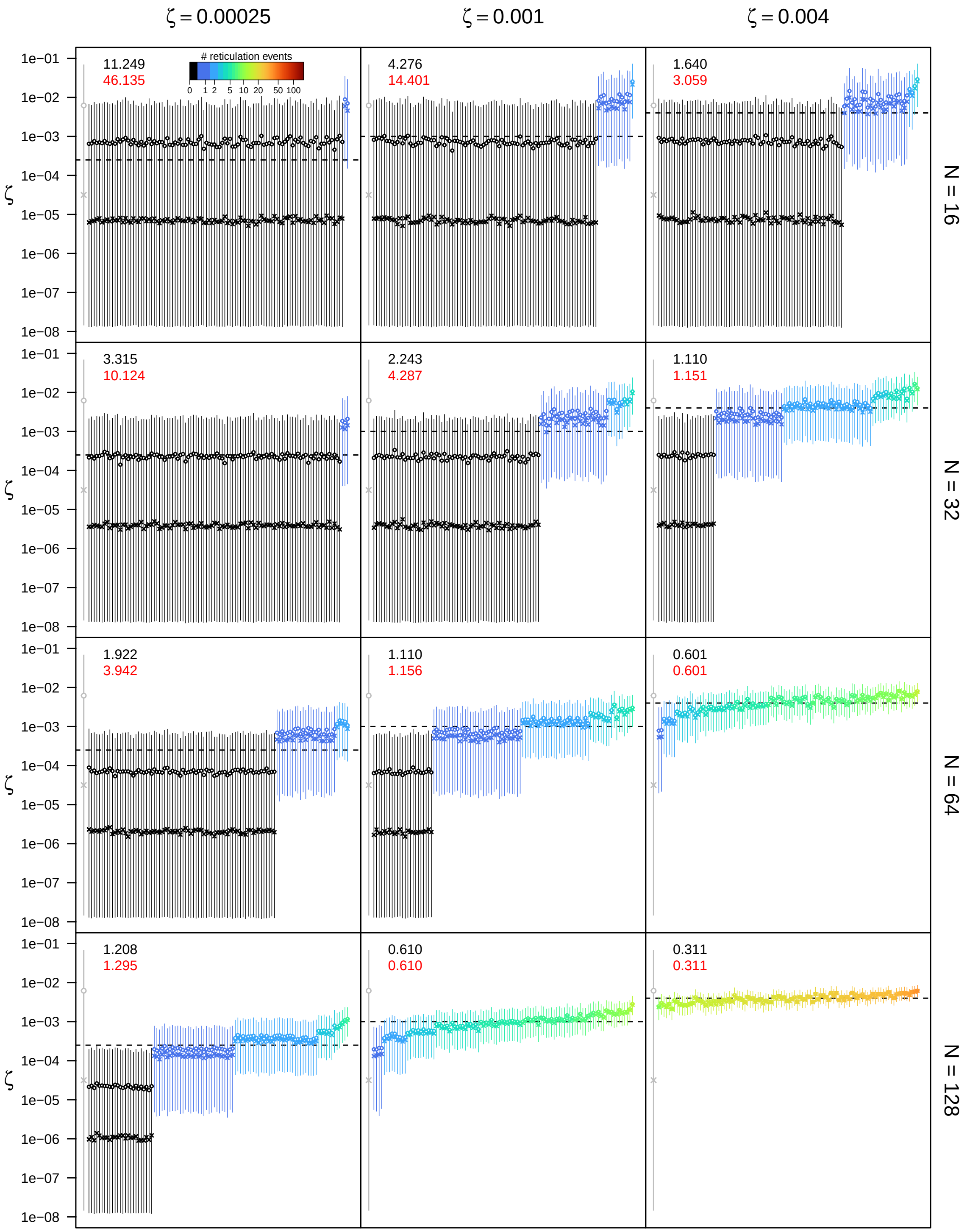


**Figure S61: Posterior estimates of the bidirectional hybridization rate $\zeta$ under the bidirectional hybridization model for $\epsilon = 0.3$, $\rho = 1.0$, and $u_{\max} = 4N$.** See main text for interpretation.

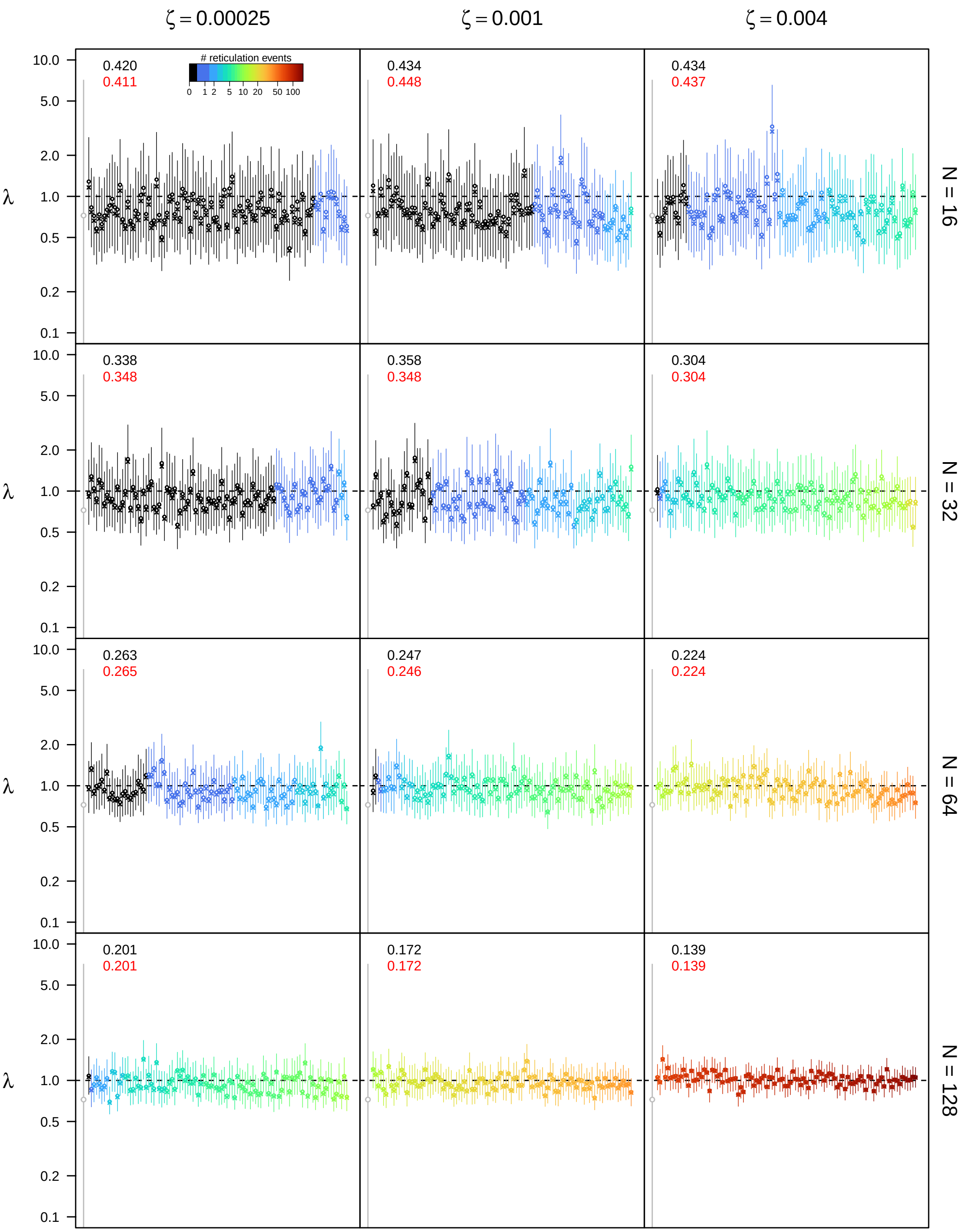


**Figure S62: Posterior estimates of the speciation rate $\lambda$ under the bidirectional hybridization model for $\epsilon = 0.5$, $\rho = 0.5$, and $u_{\max} = 4N$.** See main text for interpretation.

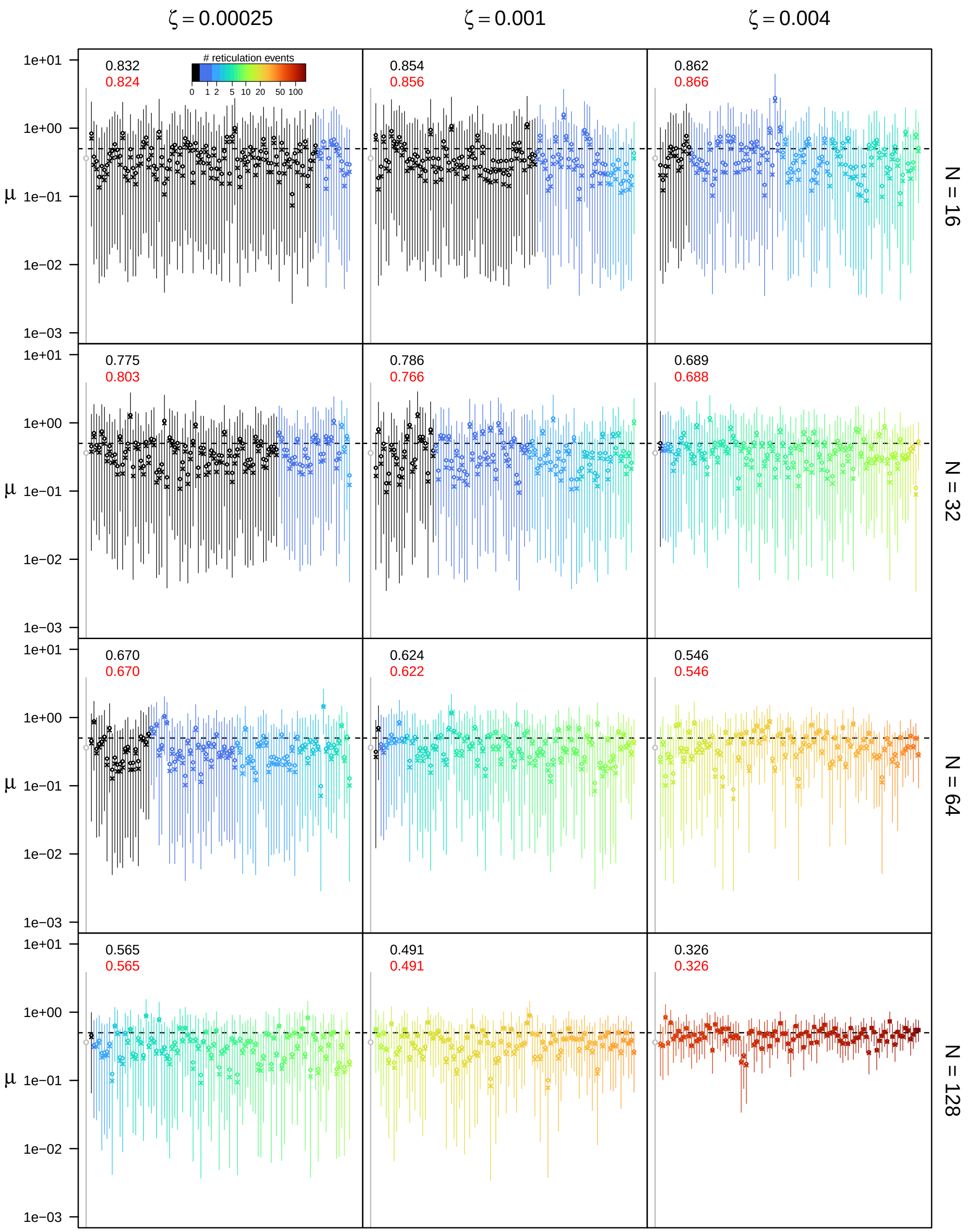


**Figure S63: Posterior estimates of the extinction rate $\mu$ under the bidirectional hybridization model for $\epsilon = 0.5$, $\rho = 0.5$, and $u_{\mathbf{max}} = 4N$.** See main text for interpretation.

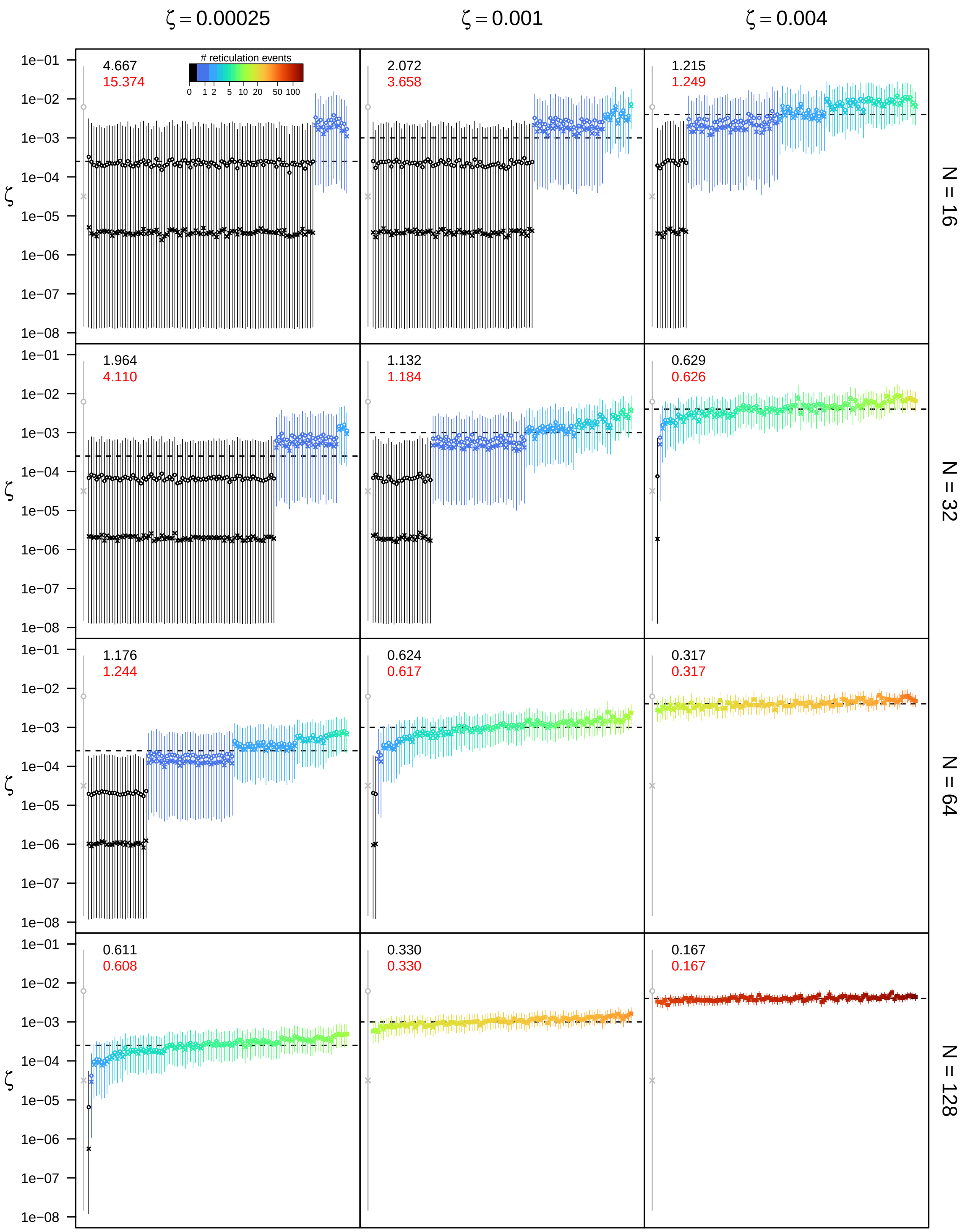


**Figure S64: Posterior estimates of the bidirectional hybridization rate $\zeta$ under the bidirectional hybridization model for** $\epsilon = 0.5$**,** $\rho = 0.5$**, and** $u_{\mathbf{max}} = 4N$**.** See main text for interpretation.

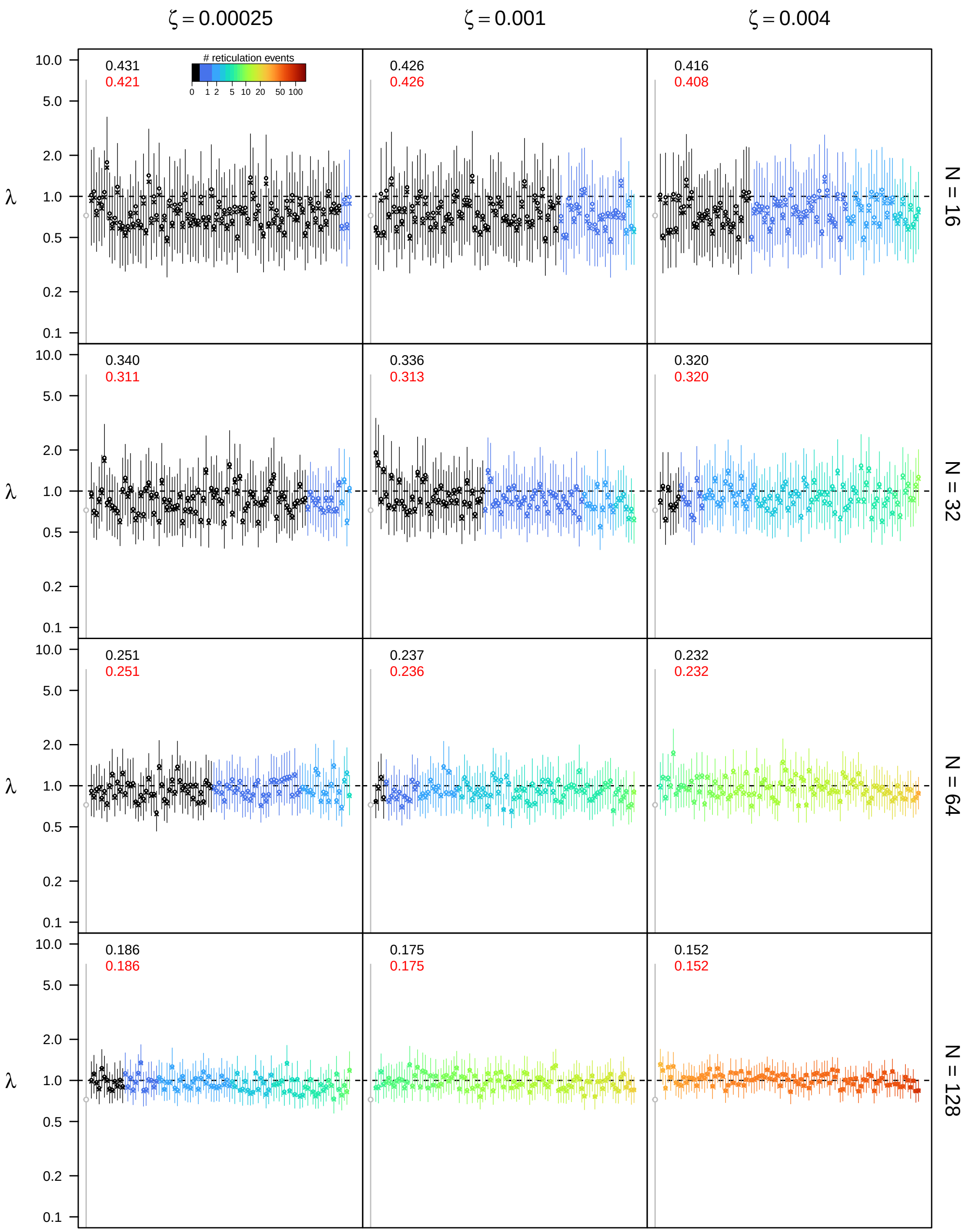


**Figure S65: Posterior estimates of the speciation rate $\lambda$ under the bidirectional hybridization model for $\epsilon = 0.5$, $\rho = 0.75$, and $u_{\mathbf{max}} = 4N$.** See main text for interpretation.

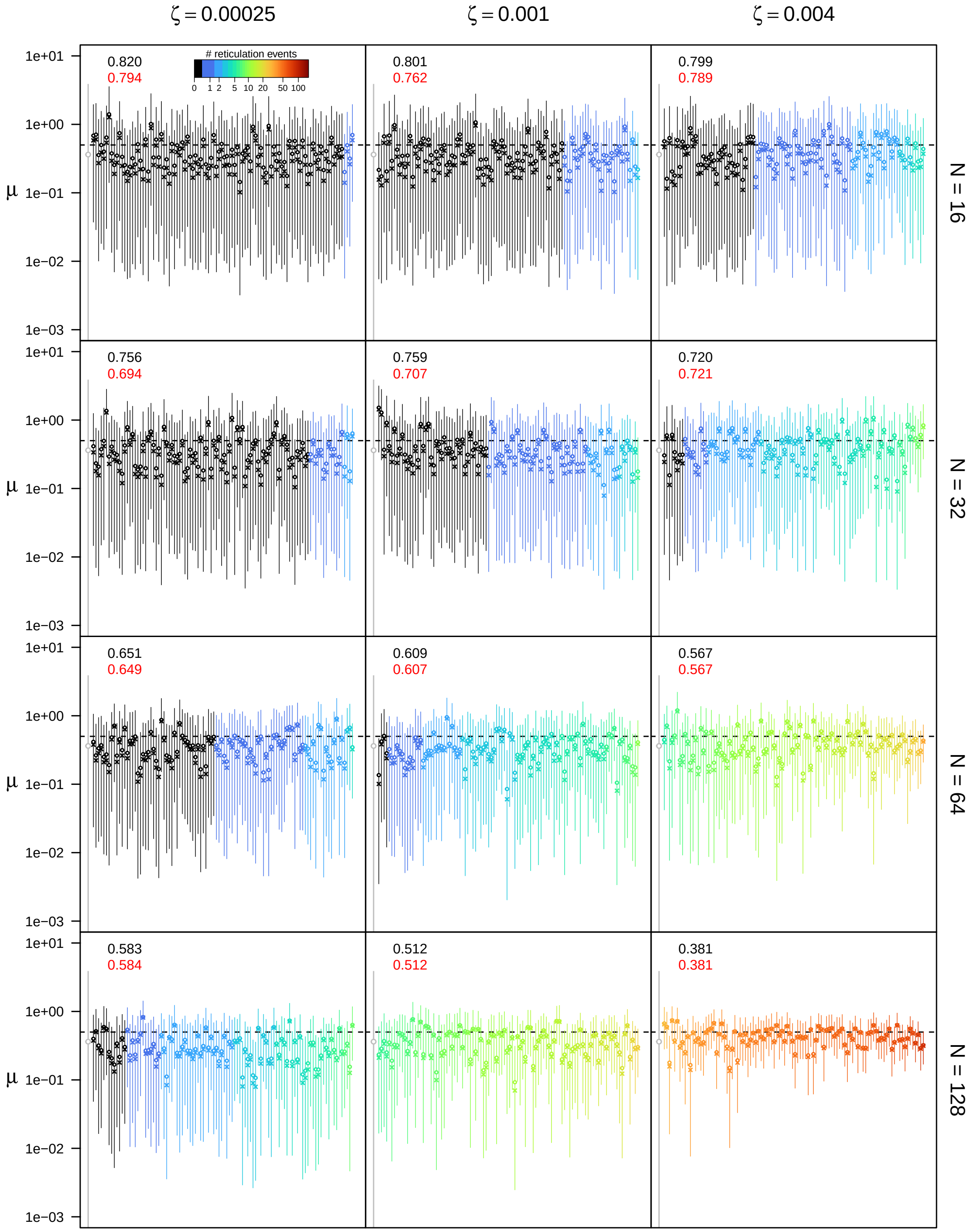


**Figure S66: Posterior estimates of the extinction rate $\mu$ under the bidirectional hybridization model for $\epsilon = 0.5$, $\rho = 0.75$, and $u_{\mathbf{max}} = 4N$.** See main text for interpretation.

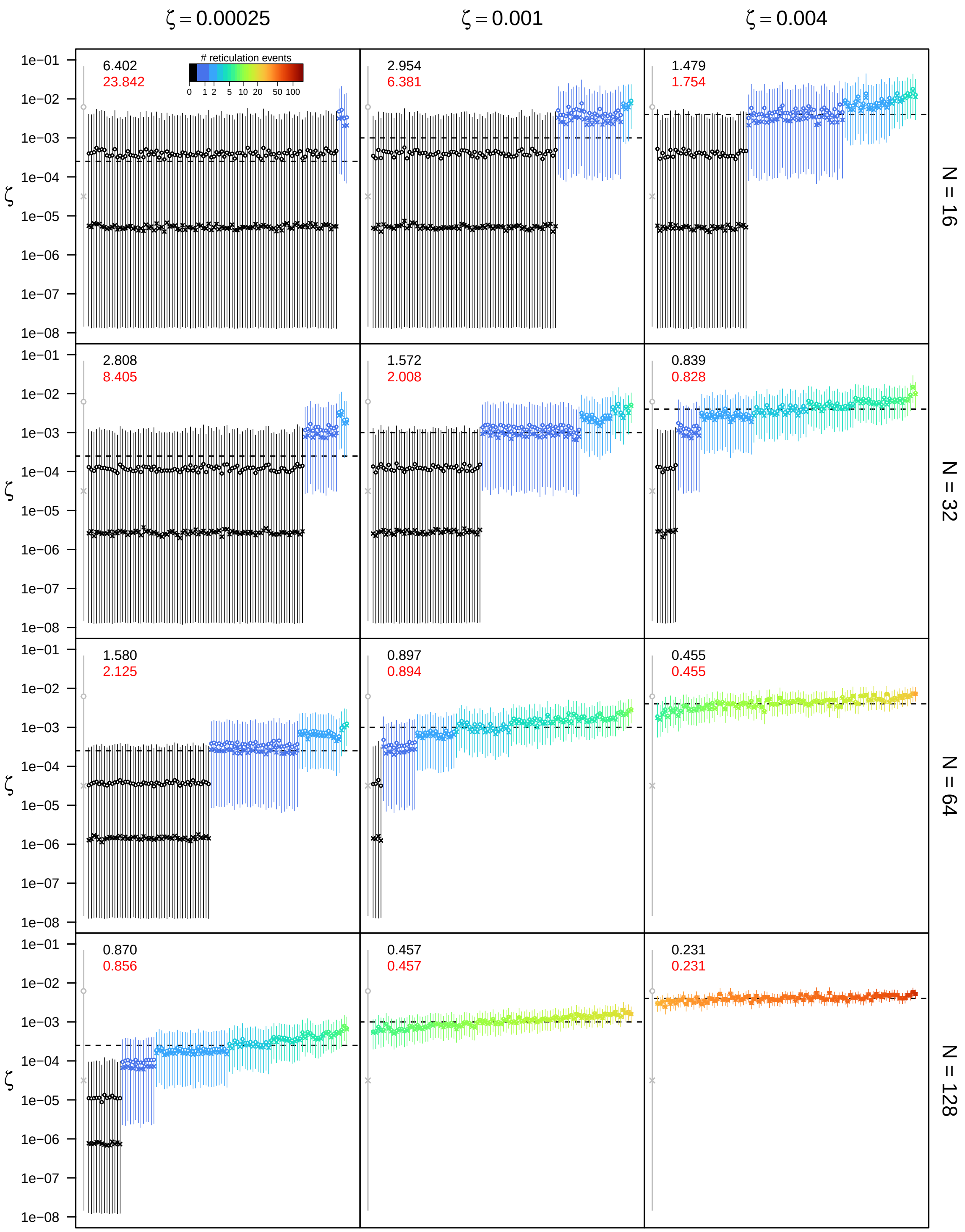


**Figure S67: Posterior estimates of the bidirectional hybridization rate $\zeta$ under the bidirectional hybridization model for $\epsilon = 0.5$, $\rho = 0.75$, and $u_{\mathbf{max}} = 4N$.** See main text for interpretation.

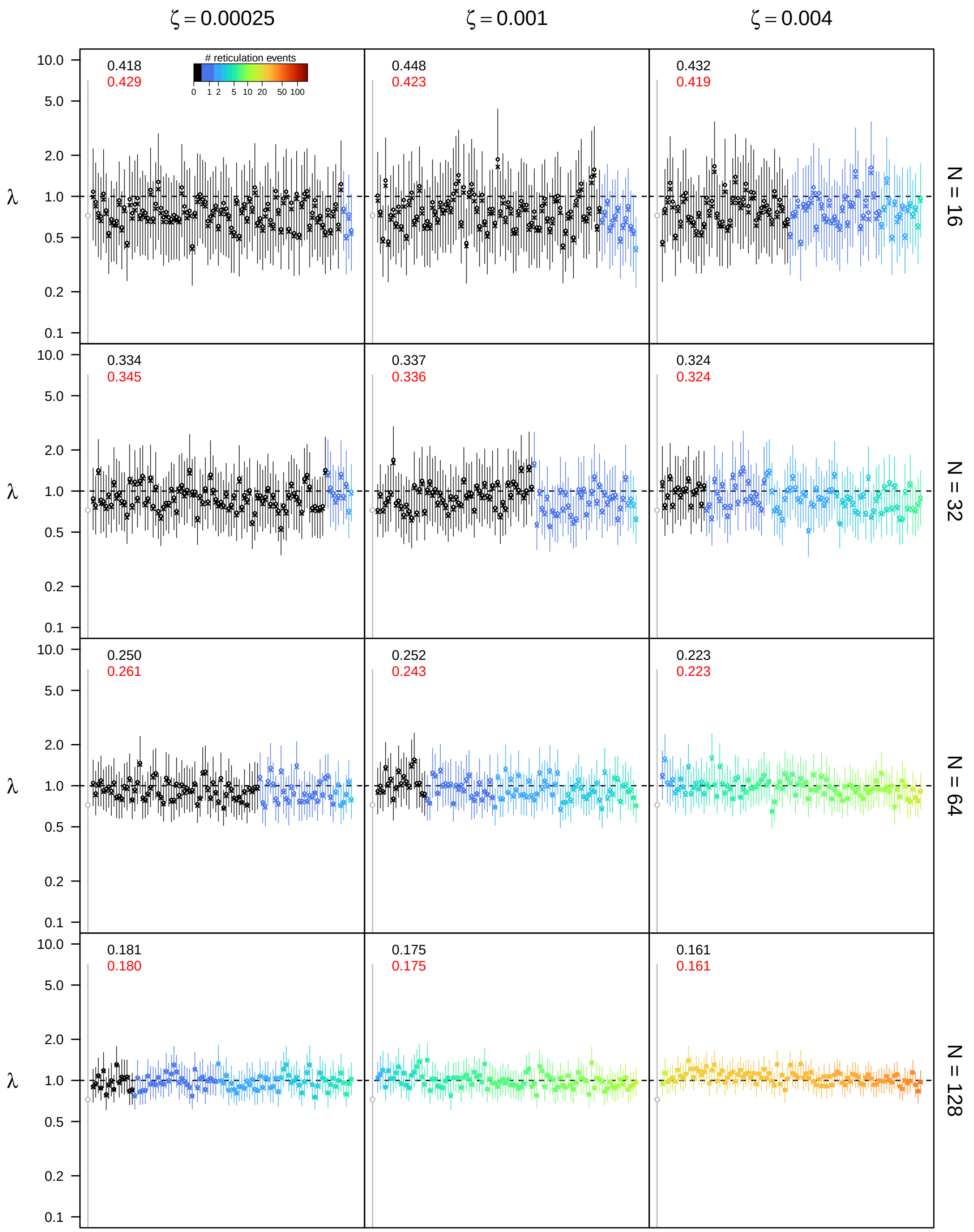


**Figure S68: Posterior estimates of the speciation rate $\lambda$ under the bidirectional hybridization model for $\epsilon = 0.5$, $\rho = 1.0$, and $u_{\mathbf{max}} = 4N$.** See main text for interpretation.

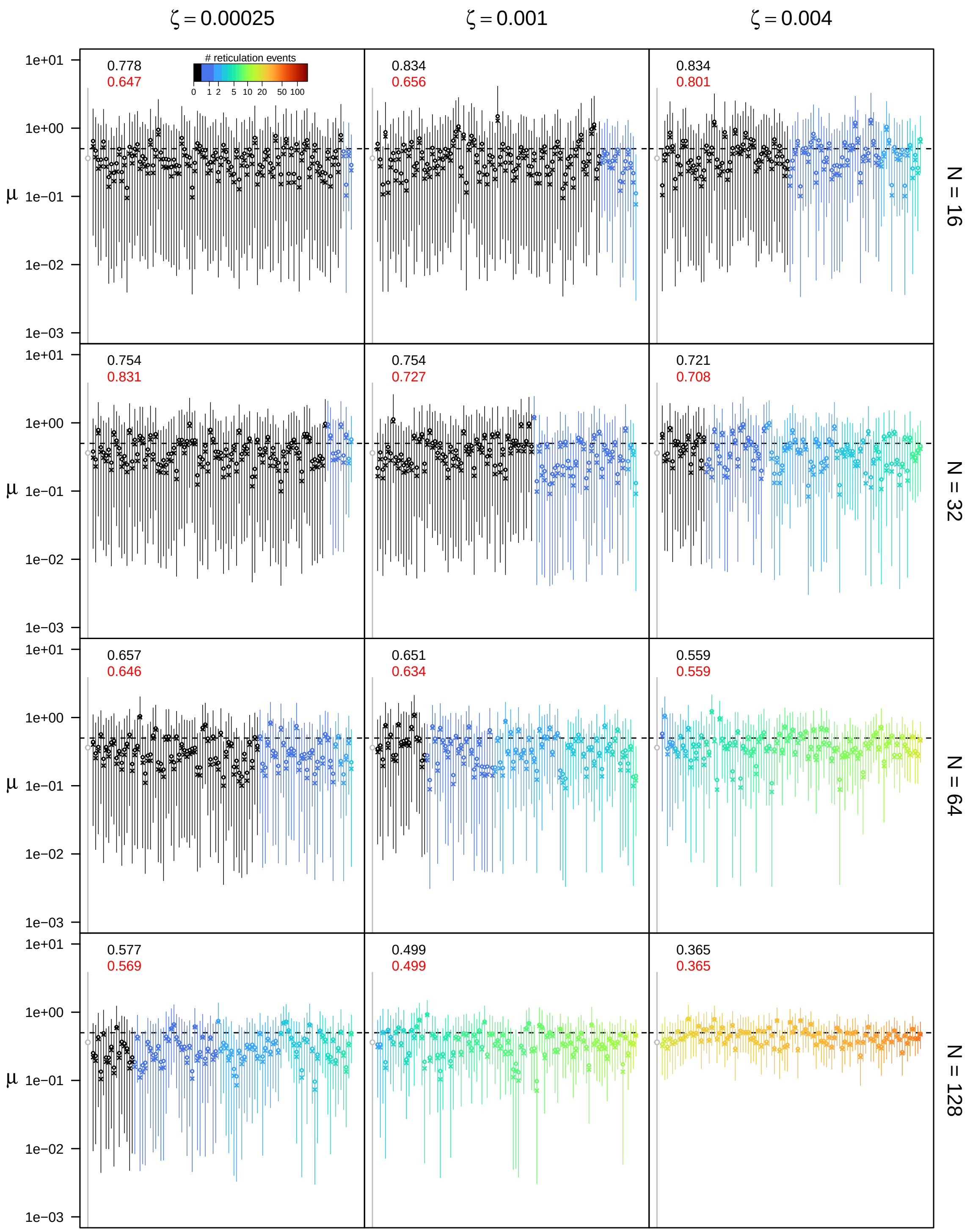


**Figure S69: Posterior estimates of the extinction rate $\mu$ under the bidirectional hybridization model for $\epsilon = 0.5$, $\rho = 1.0$, and $u_{\mathbf{max}} = 4N$.** See main text for interpretation.

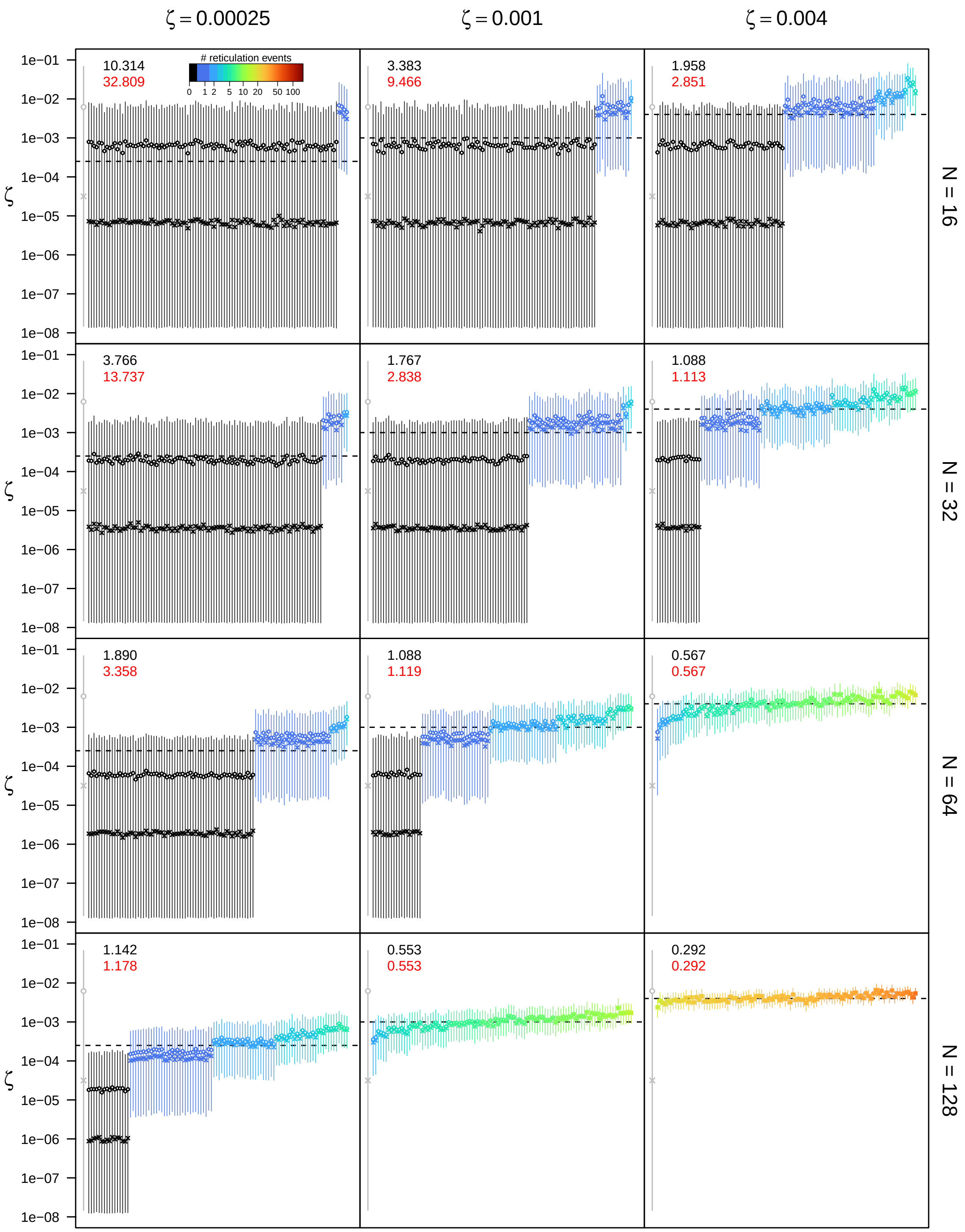


**Figure S70: Posterior estimates of the bidirectional hybridization rate $\zeta$ under the bidirectional hybridization model for $\epsilon = 0.5$, $\rho = 1.0$, and $u_{\textbf{max}} = 4N$.** See main text for interpretation.

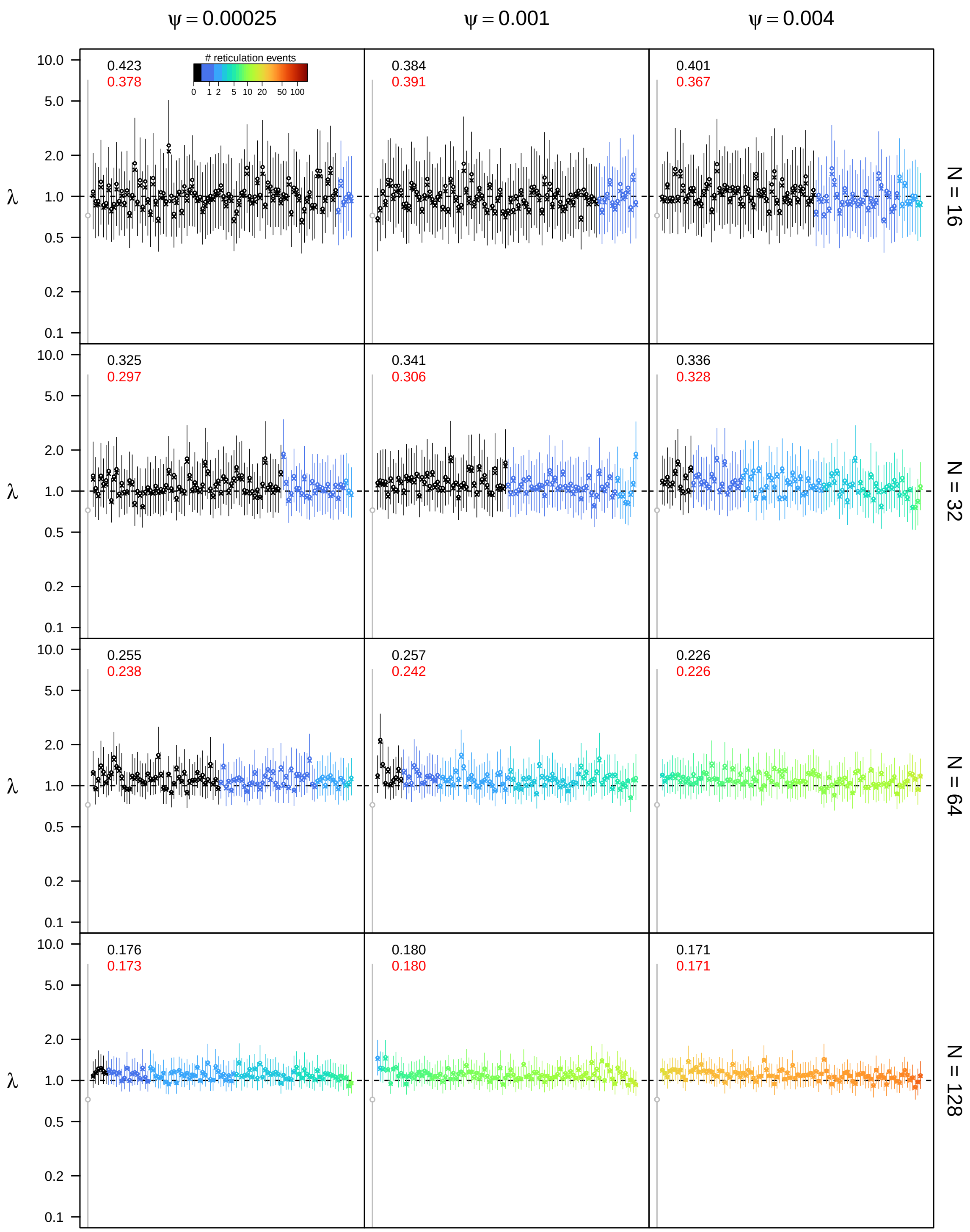


**Figure S71: Posterior estimates of the speciation rate $\lambda$ under the allopolyploidization model for $\epsilon = 0.1$, $\rho = 0.5$, and $u_{\mathbf{max}} = 4N$.** See main text for interpretation.

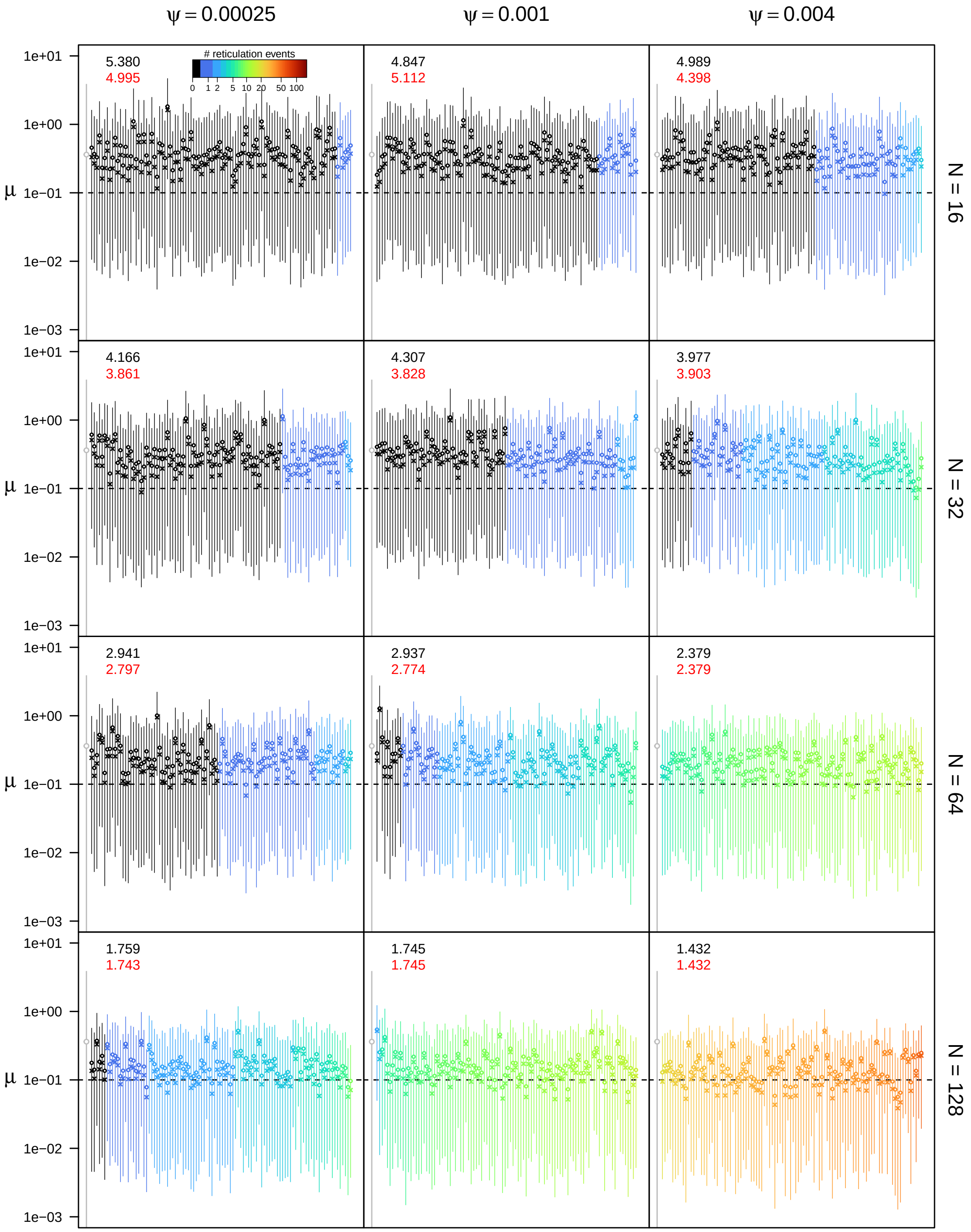


**Figure S72: Posterior estimates of the extinction rate $\mu$ under the allopolyploidization model for $\epsilon = 0.1$, $\rho = 0.5$, and $u_{\mathbf{max}} = 4N$.** See main text for interpretation.

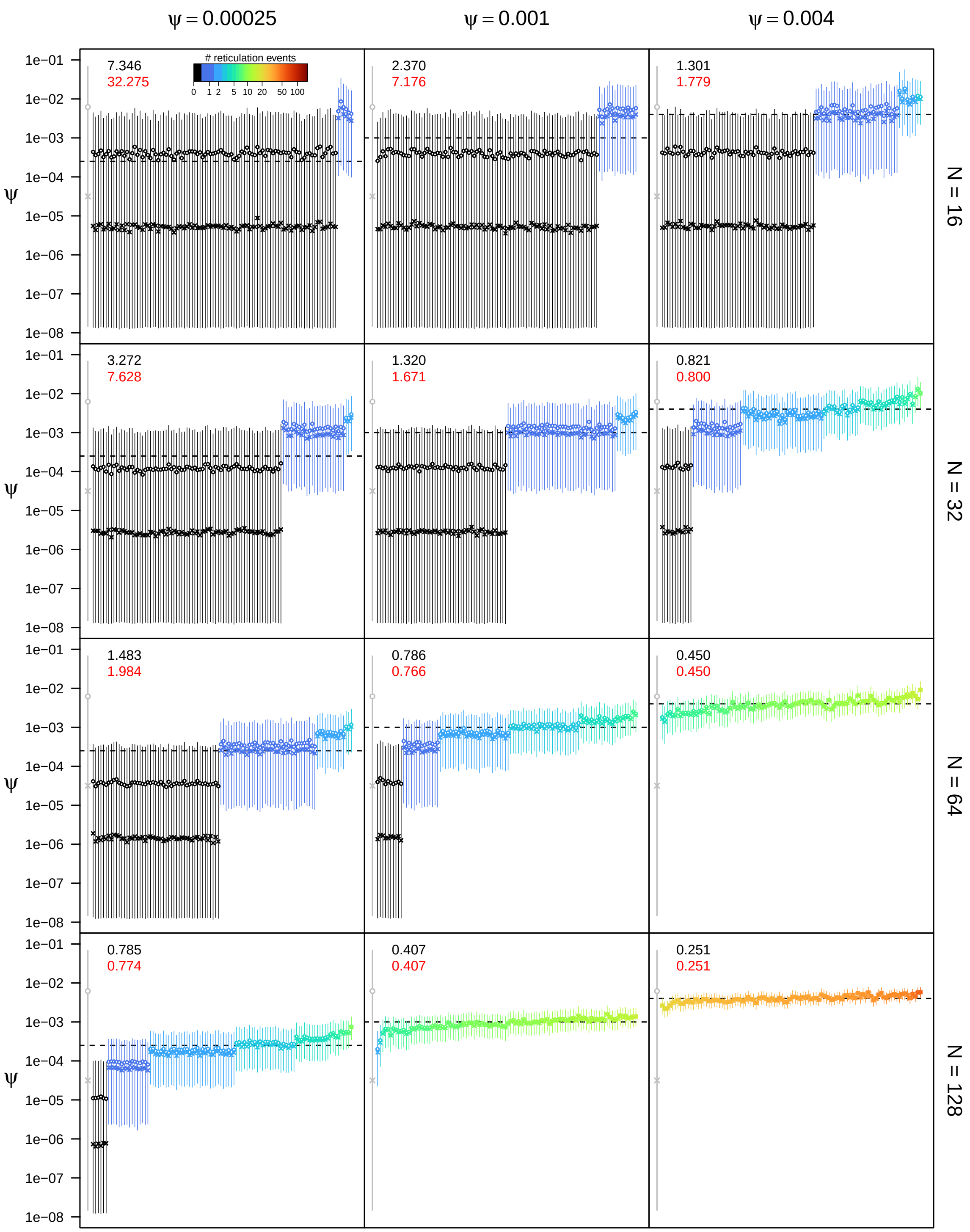


**Figure S73: Posterior estimates of the allopolyploidization rate $\psi$ under the allopolyploidization model for $\epsilon = 0.1$, $\rho = 0.5$, and $u_{\max} = 4N$.** See main text for interpretation.

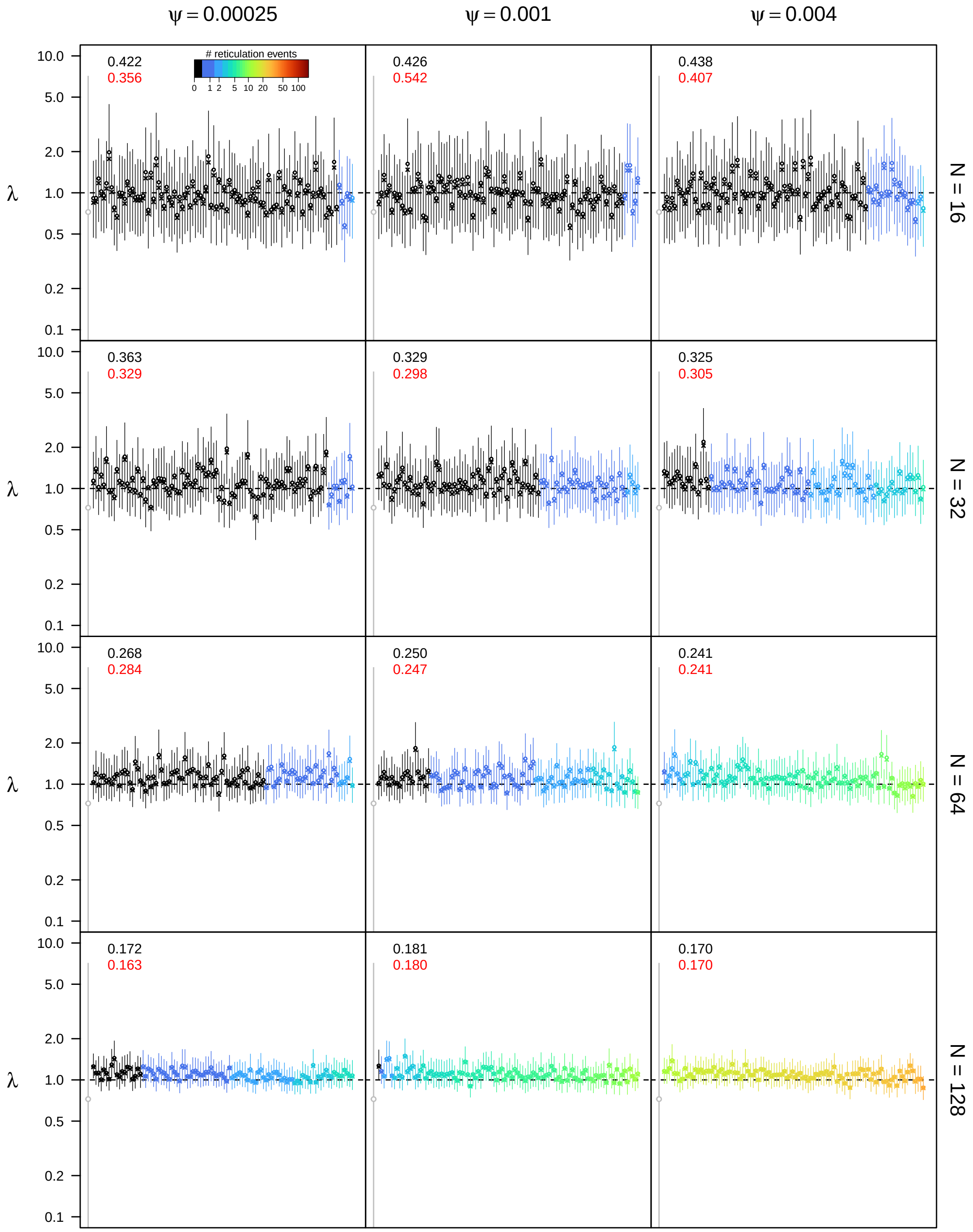


**Figure S74: Posterior estimates of the speciation rate $\lambda$ under the allopolyploidization model for $\epsilon = 0.1$, $\rho = 0.75$, and $u_{\mathbf{max}} = 4N$.** See main text for interpretation.

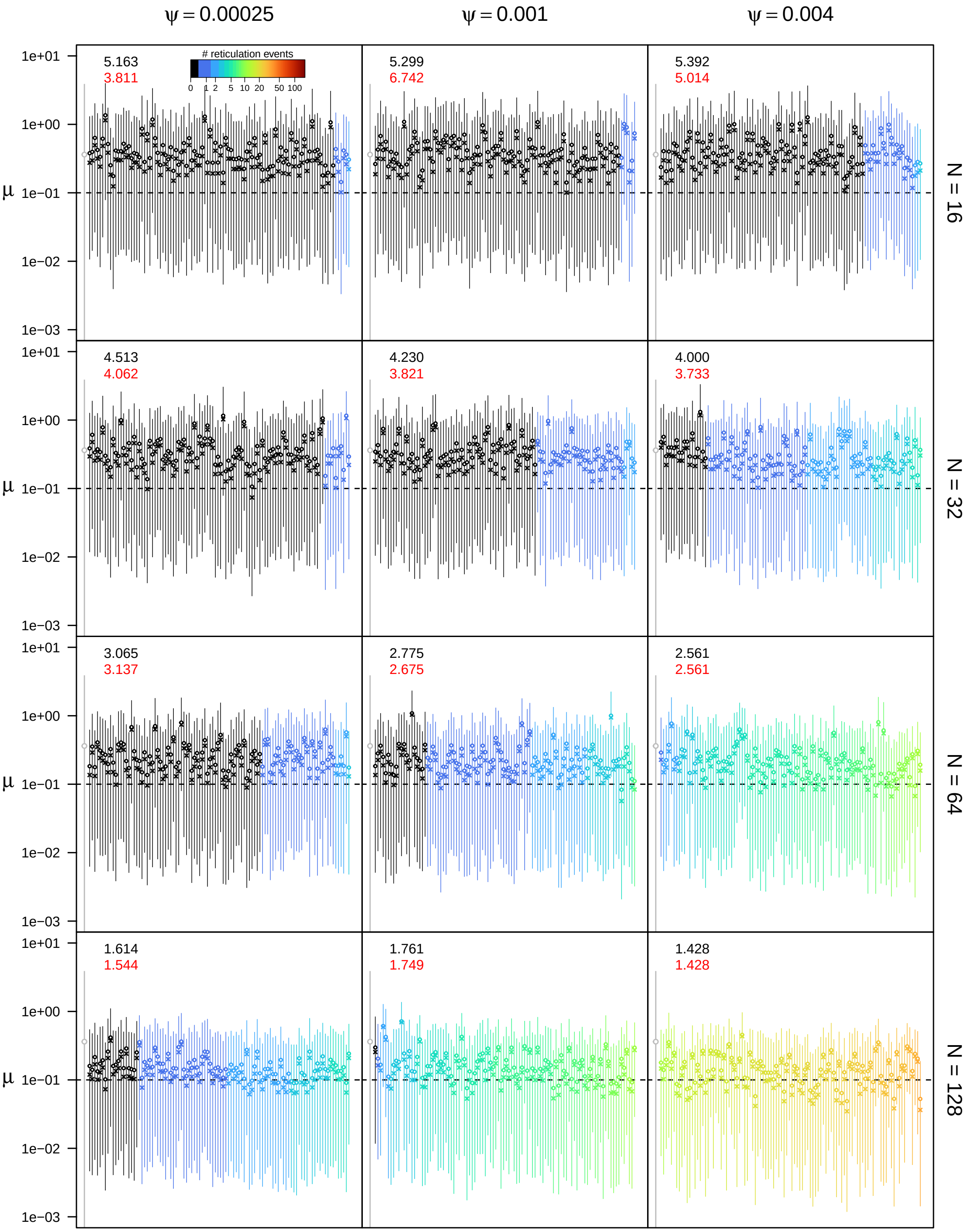


**Figure S75: Posterior estimates of the extinction rate $\mu$ under the allopolyploidization model for $\epsilon = 0.1$, $\rho = 0.75$, and $u_{\mathbf{max}} = 4N$.** See main text for interpretation.

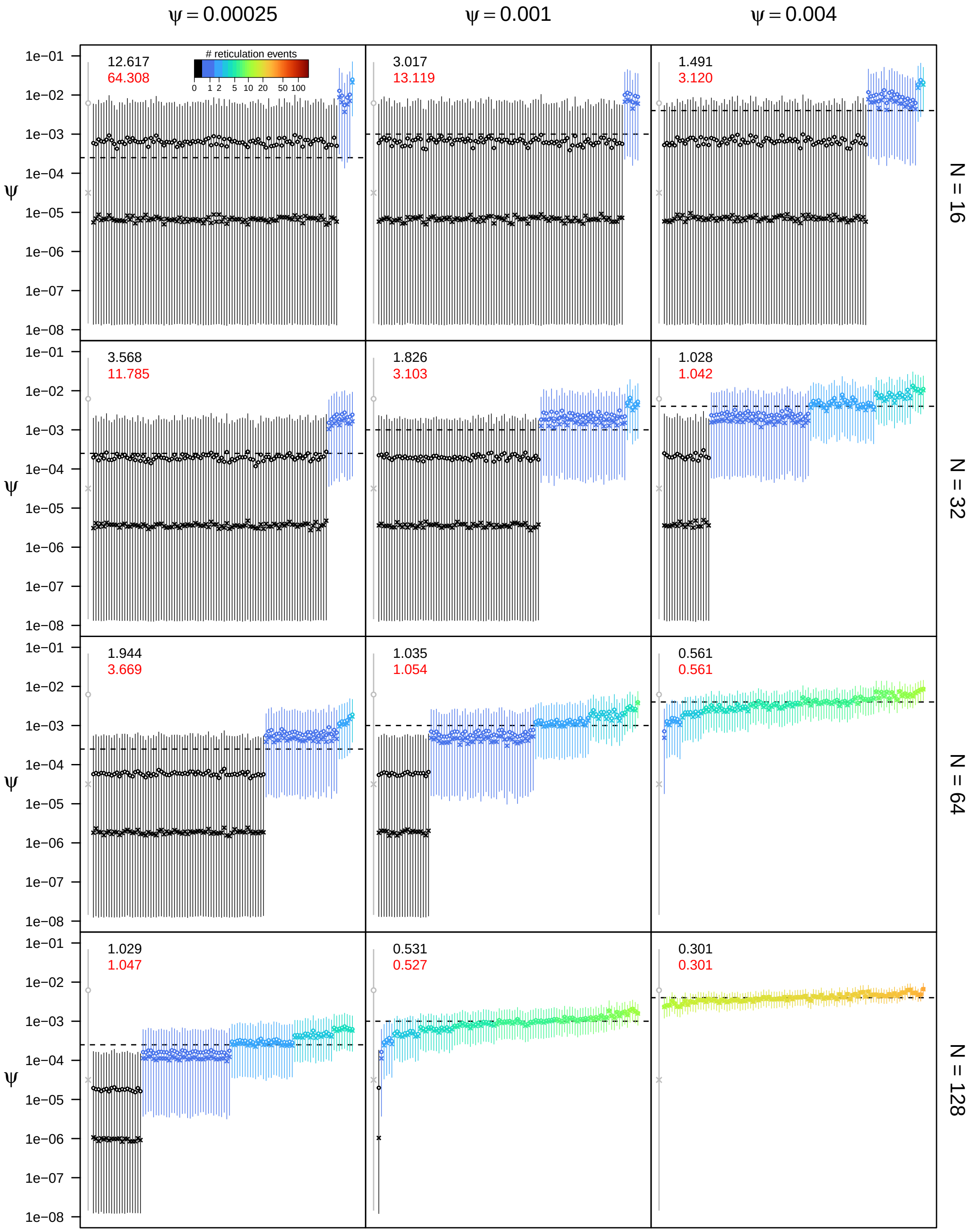


**Figure S76: Posterior estimates of the allopolyploidization rate $\psi$ under the allopolyploidization model for $\epsilon = 0.1$, $\rho = 0.75$, and $u_{\mathbf{max}} = 4N$.** See main text for interpretation.

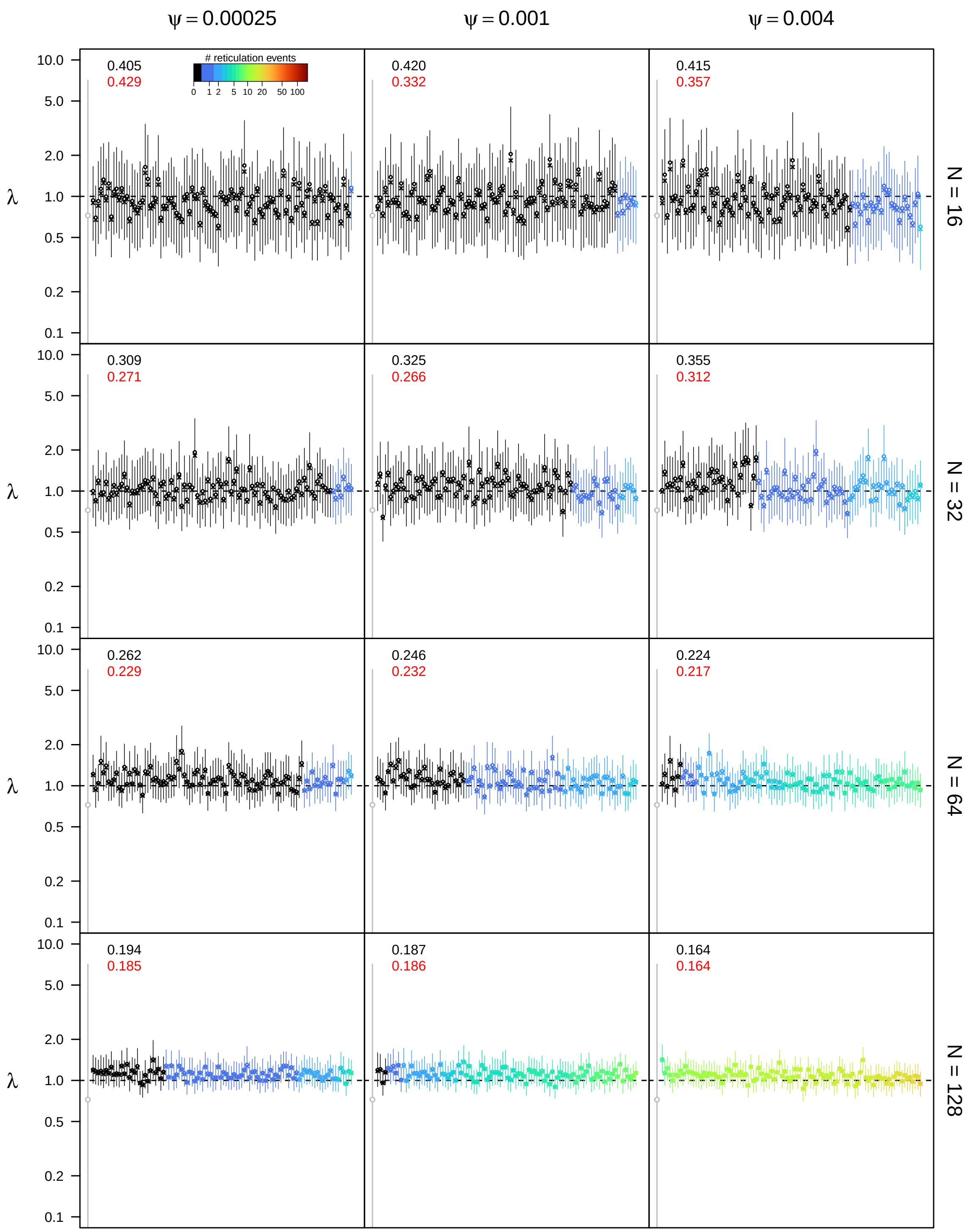


**Figure S77: Posterior estimates of the speciation rate $\lambda$ under the allopolyploidization model for $\epsilon = 0.1$, $\rho = 1.0$, and $u_{\mathbf{max}} = 4N$.** See main text for interpretation.

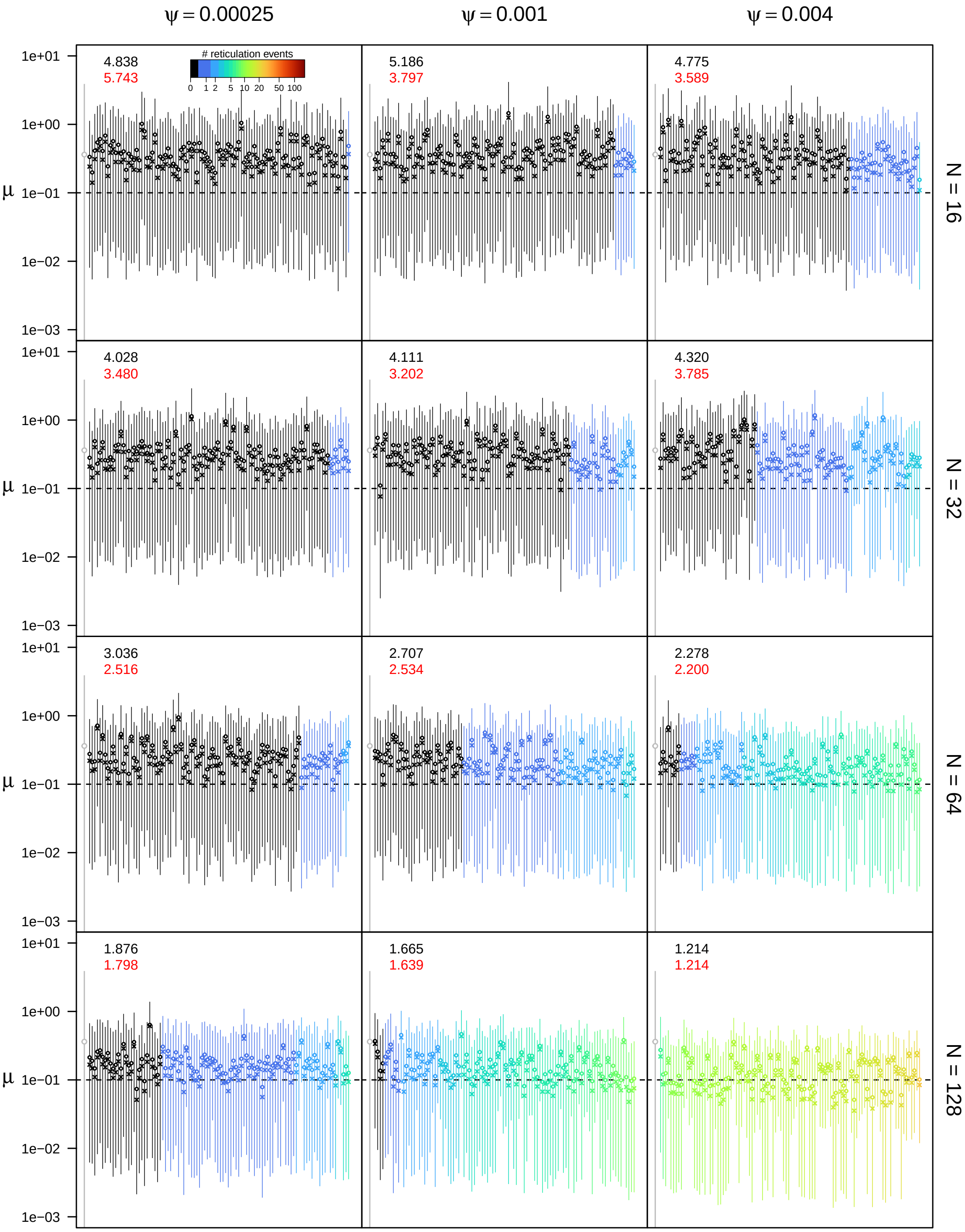


**Figure S78: Posterior estimates of the extinction rate $\mu$ under the allopolyploidization model for $\epsilon = 0.1$, $\rho = 1.0$, and $u_{\mathbf{max}} = 4N$.** See main text for interpretation.

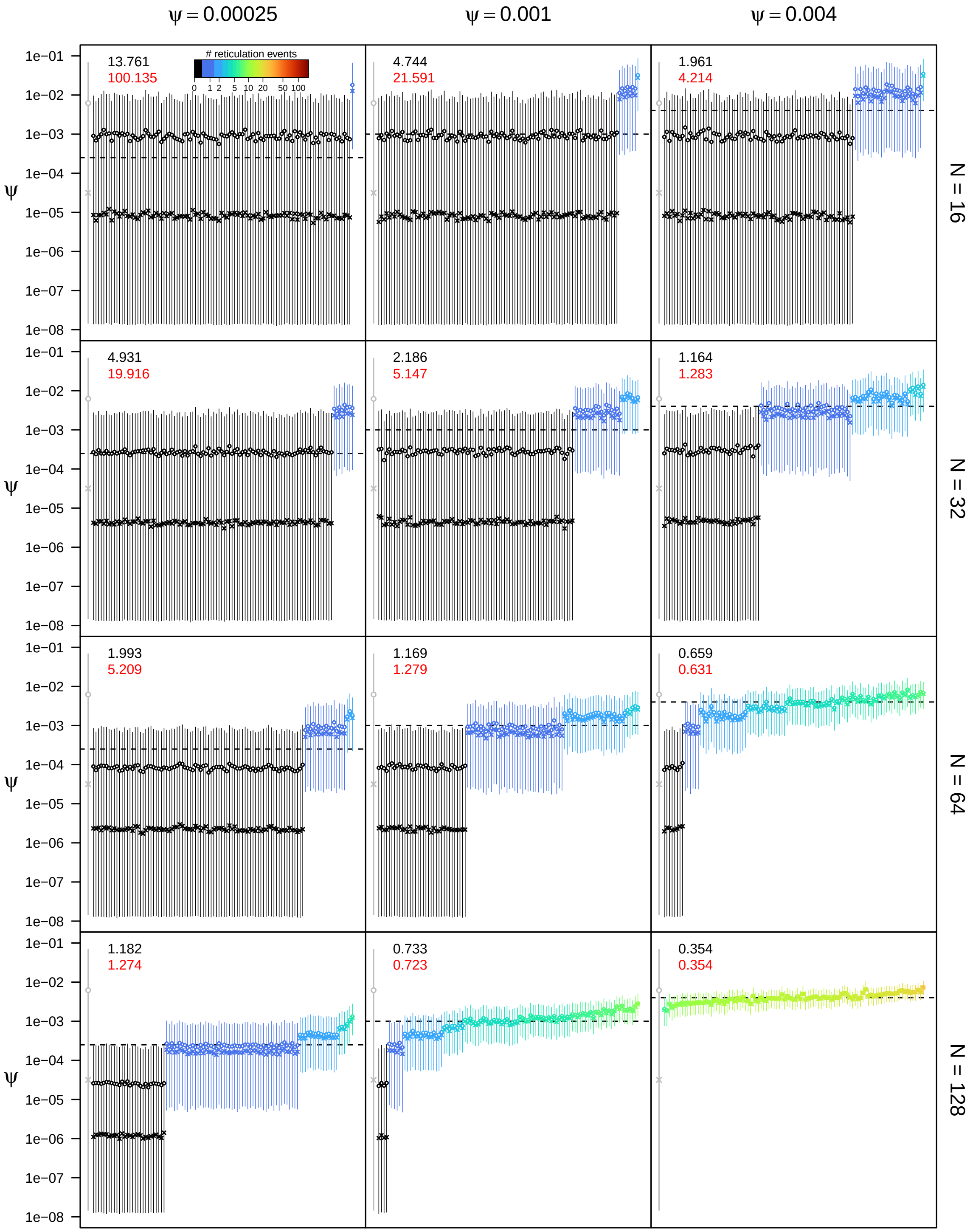


**Figure S79: Posterior estimates of the allopolyploidization rate $\psi$ under the allopolyploidization model for $\epsilon = 0.1$, $\rho = 1.0$, and $u_{\mathbf{max}} = 4N$.** See main text for interpretation.

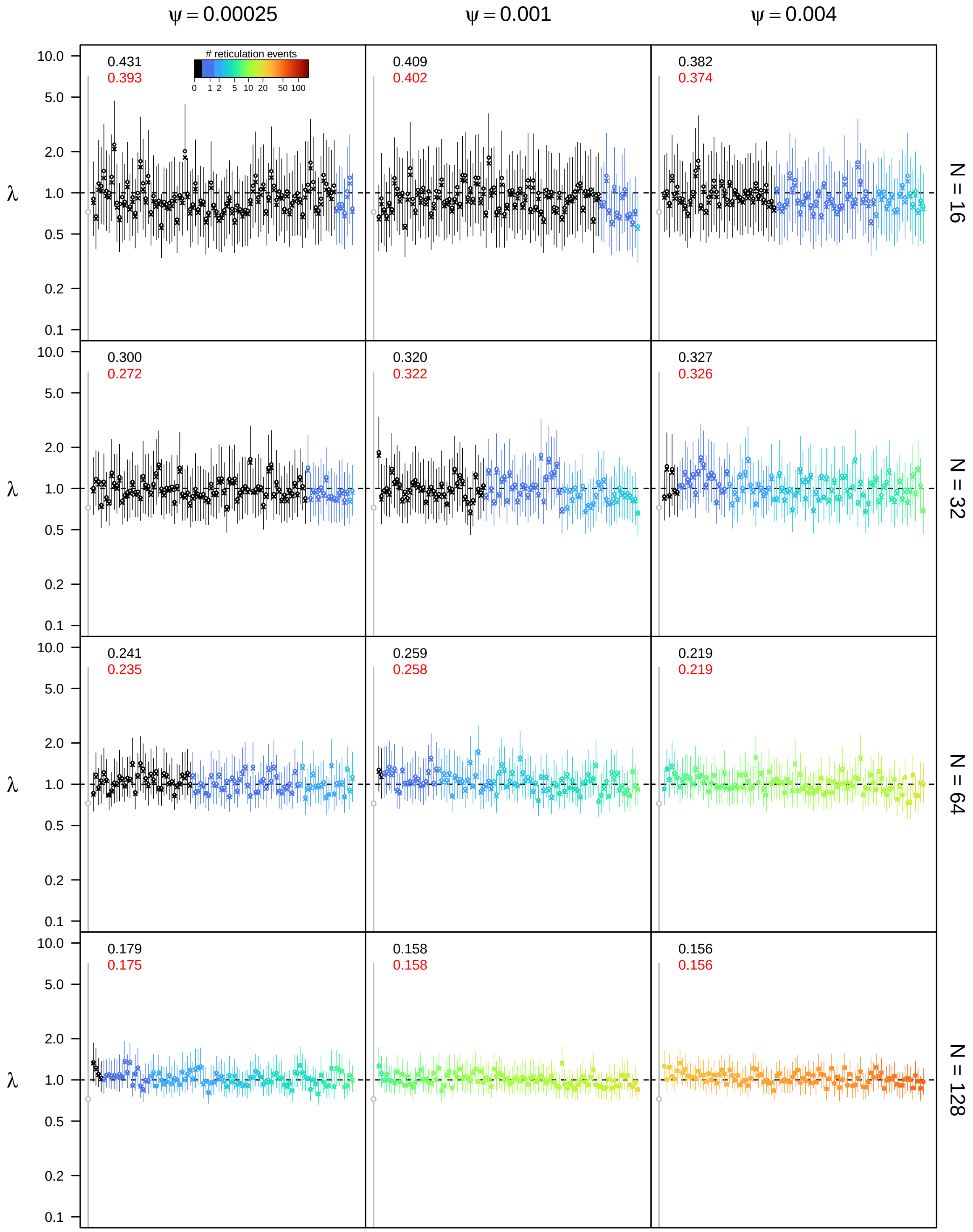


**Figure S80: Posterior estimates of the speciation rate $\lambda$ under the allopolyploidization model for $\epsilon = 0.3$, $\rho = 0.5$, and $u_{\mathbf{max}} = 4N$.** See main text for interpretation.

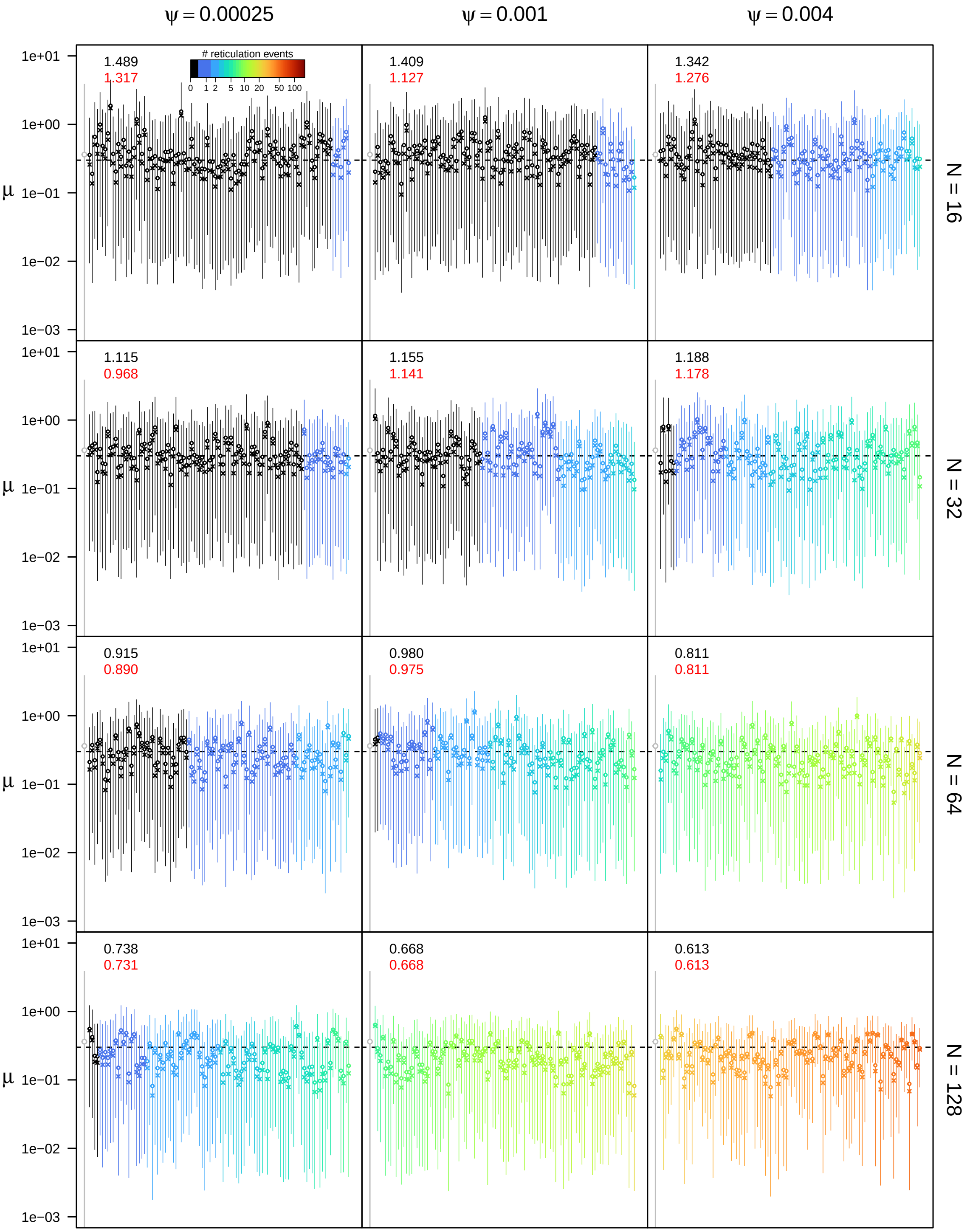


**Figure S81: Posterior estimates of the extinction rate $\mu$ under the allopolyploidization model for $\epsilon = 0.3$, $\rho = 0.5$, and $u_{\mathbf{max}} = 4N$.** See main text for interpretation.

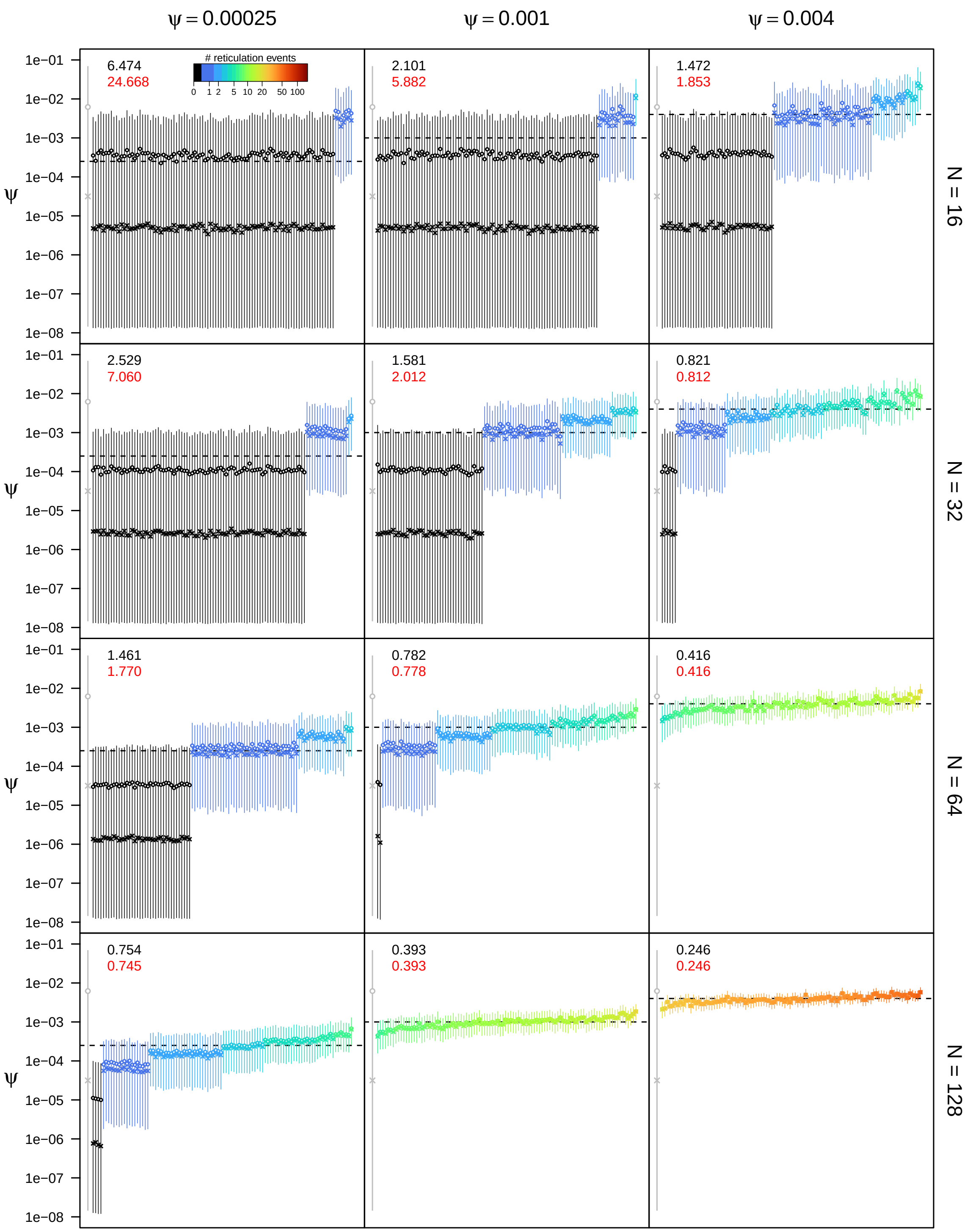


**Figure S82: Posterior estimates of the allopolyploidization rate $\psi$ under the allopolyploidization model for $\epsilon = 0.3$, $\rho = 0.5$, and $u_{\max} = 4N$.** See main text for interpretation.

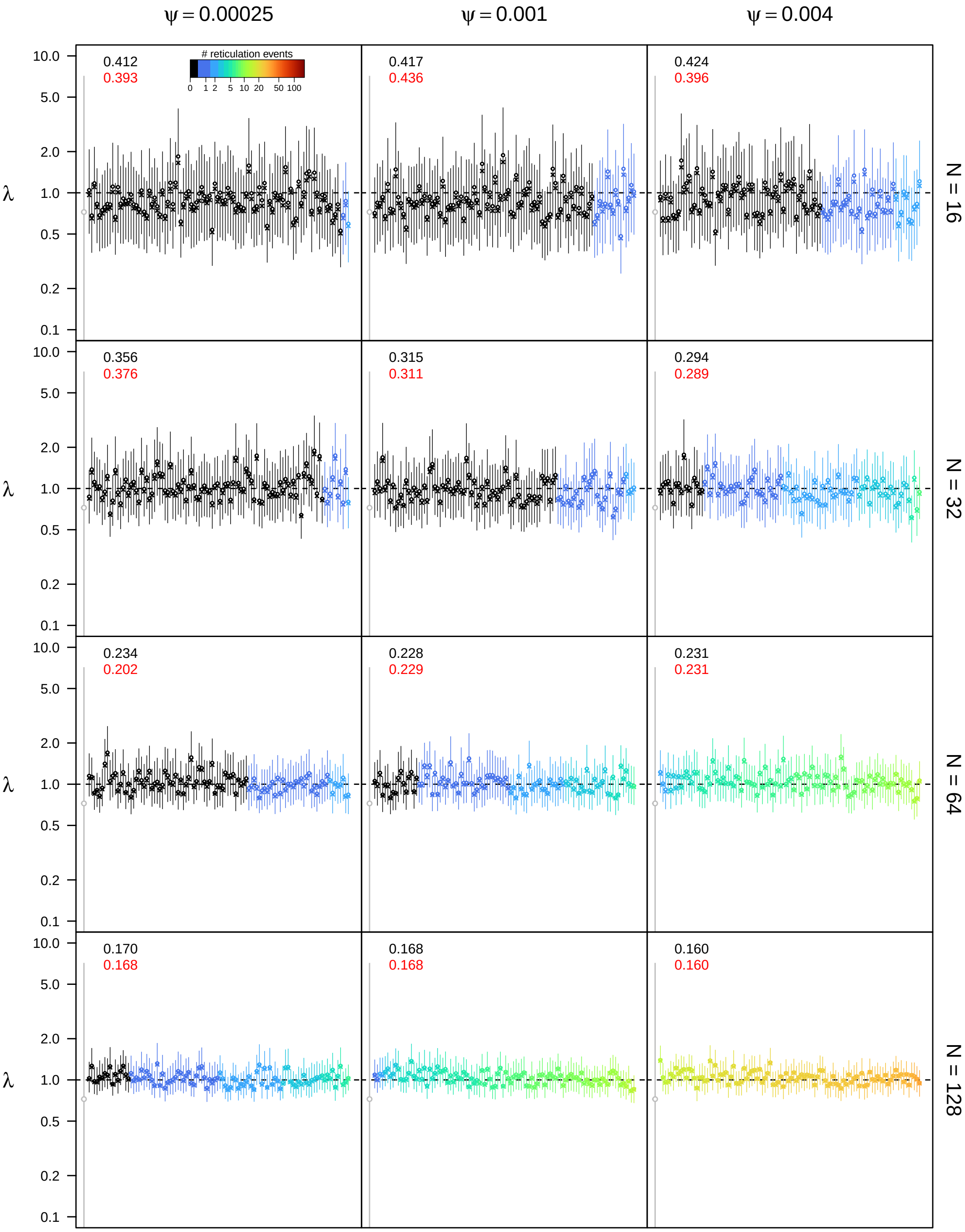


**Figure S83: Posterior estimates of the speciation rate $\lambda$ under the allopolyploidization model for $\epsilon = 0.3$, $\rho = 0.75$, and** $u_{\mathbf{max}} = 4N$. See main text for interpretation.

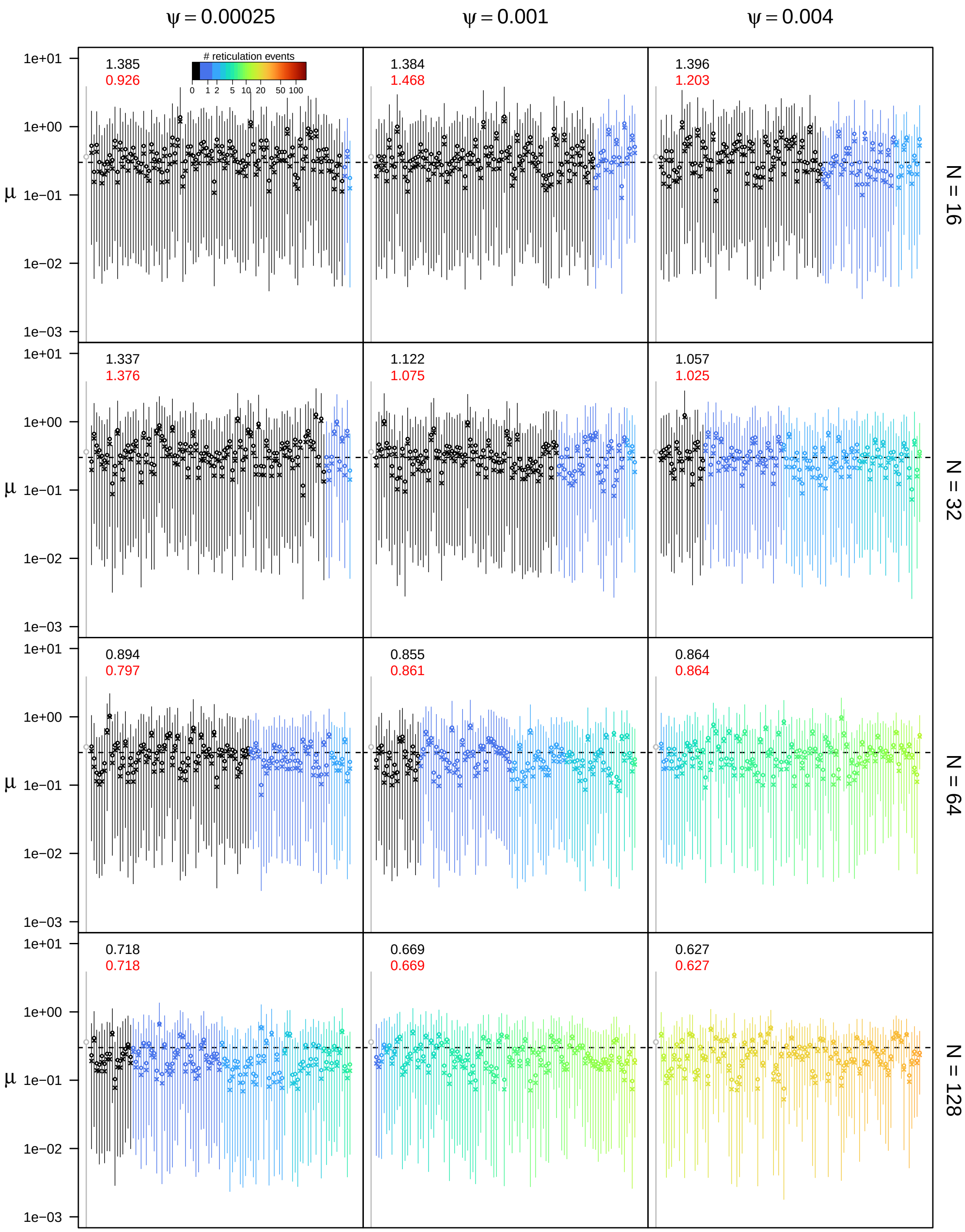


**Figure S84: Posterior estimates of the extinction rate $\mu$ under the allopolyploidization model for $\epsilon = 0.3$, $\rho = 0.75$, and** $u_{\max} = 4N$**.** See main text for interpretation.

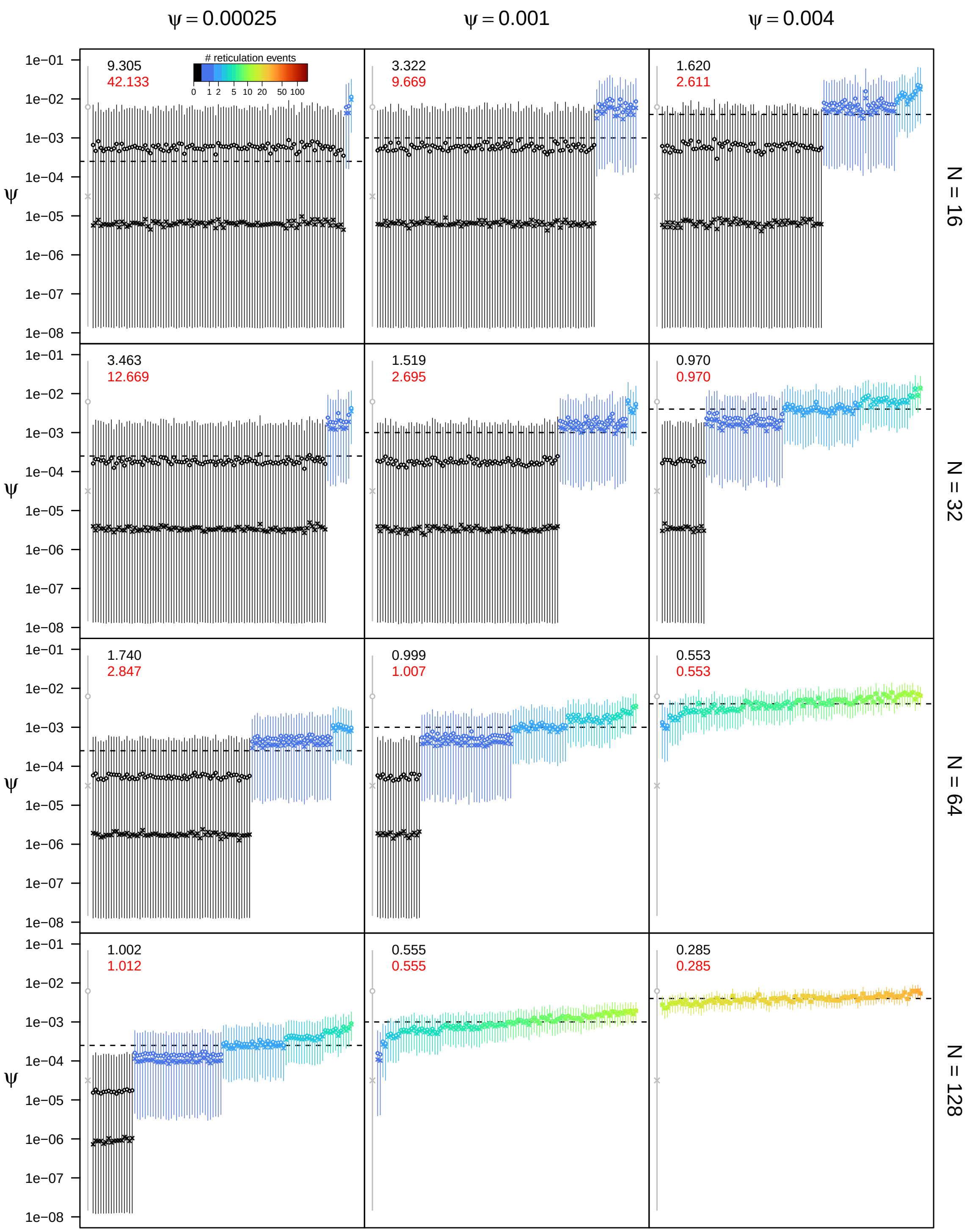


**Figure S85: Posterior estimates of the allopolyploidization rate $\psi$ under the allopolyploidization model for $\epsilon = 0.3$, $\rho = 0.75$, and $u_{\mathbf{max}} = 4N$.** See main text for interpretation.

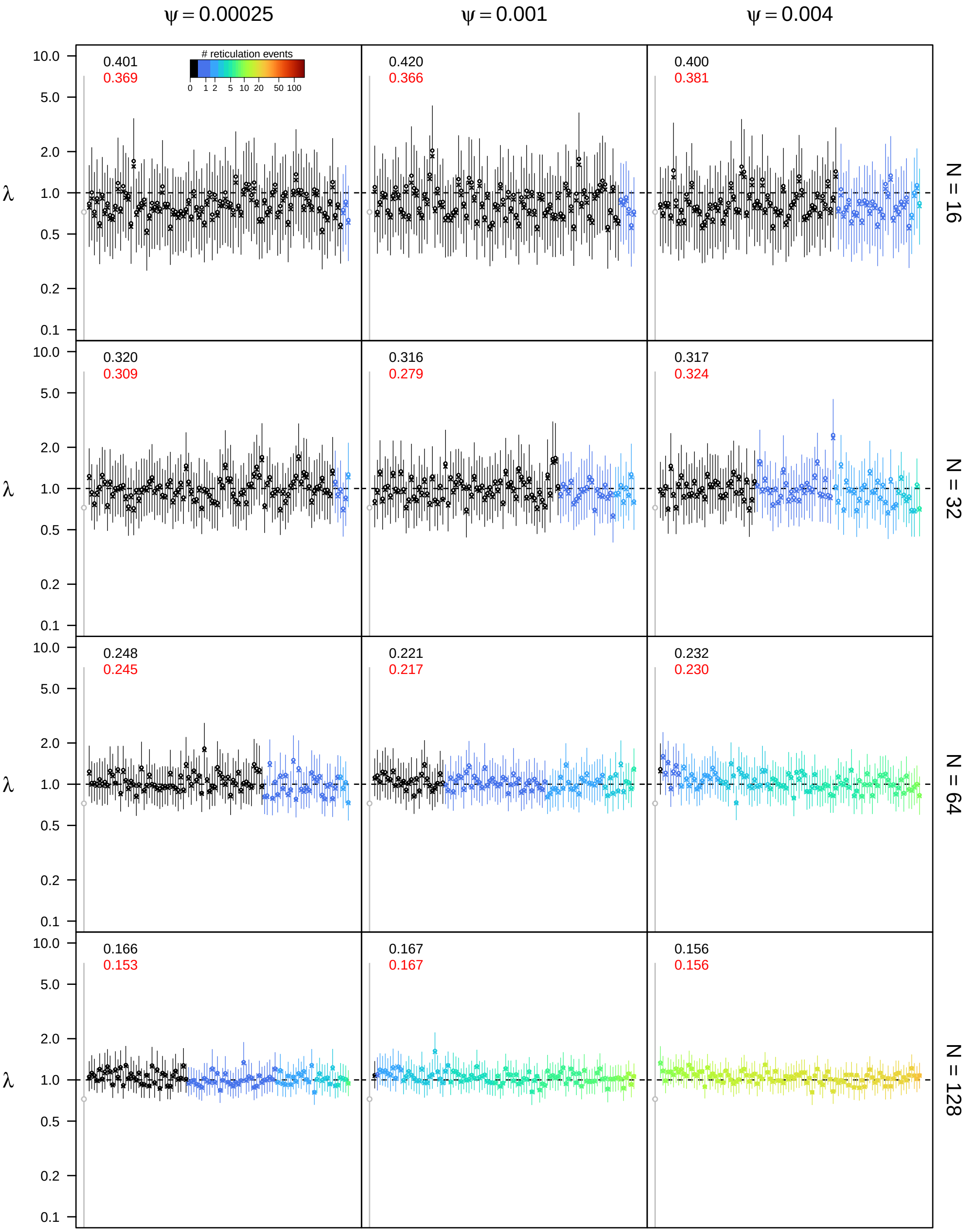


**Figure S86: Posterior estimates of the speciation rate $\lambda$ under the allopolyploidization model for $\epsilon = 0.3$, $\rho = 1.0$, and $u_{\mathbf{max}} = 4N$.** See main text for interpretation.

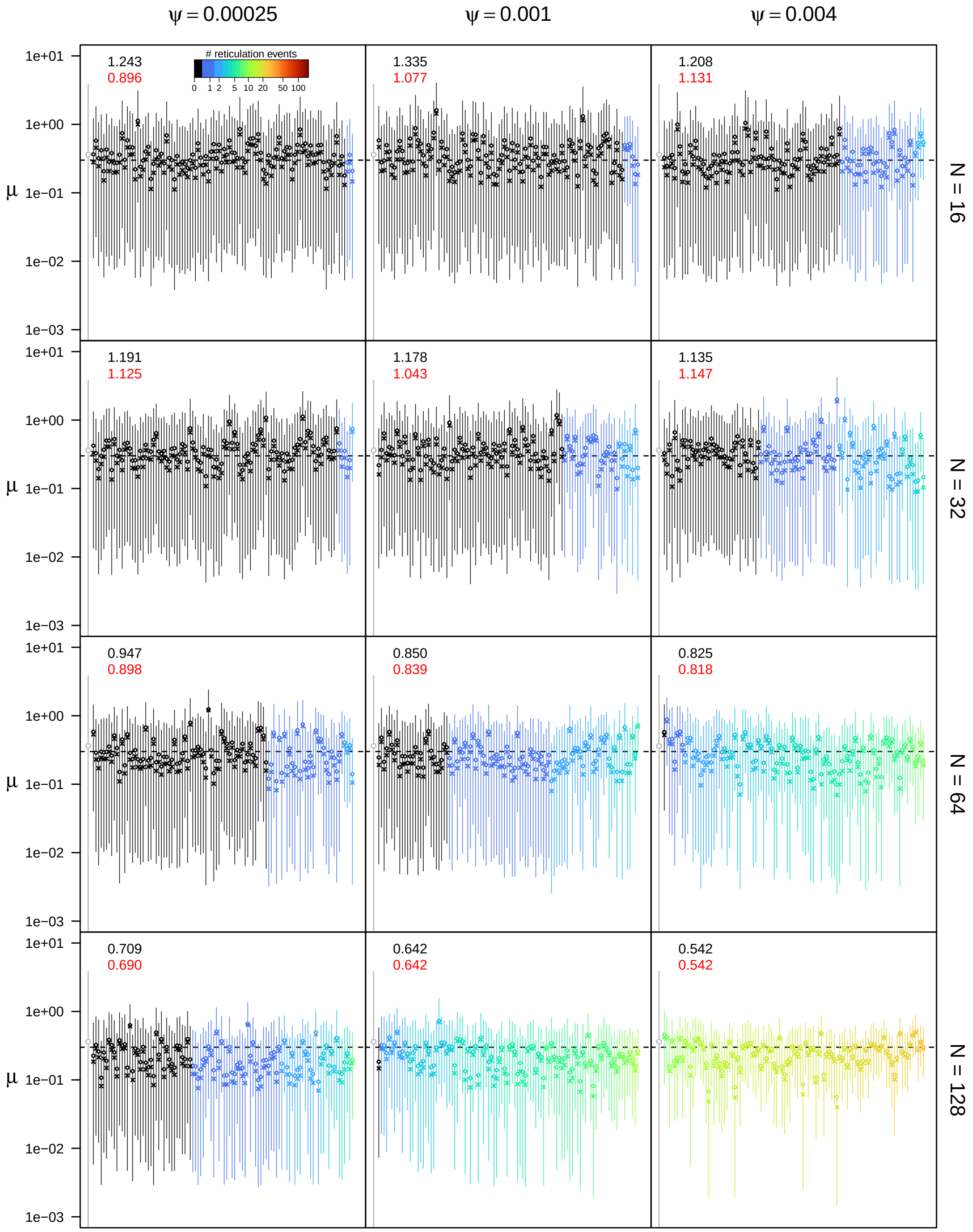


**Figure S87: Posterior estimates of the extinction rate $\mu$ under the allopolyploidization model for $\epsilon = 0.3$, $\rho = 1.0$, and $u_{\mathbf{max}} = 4N$.** See main text for interpretation.

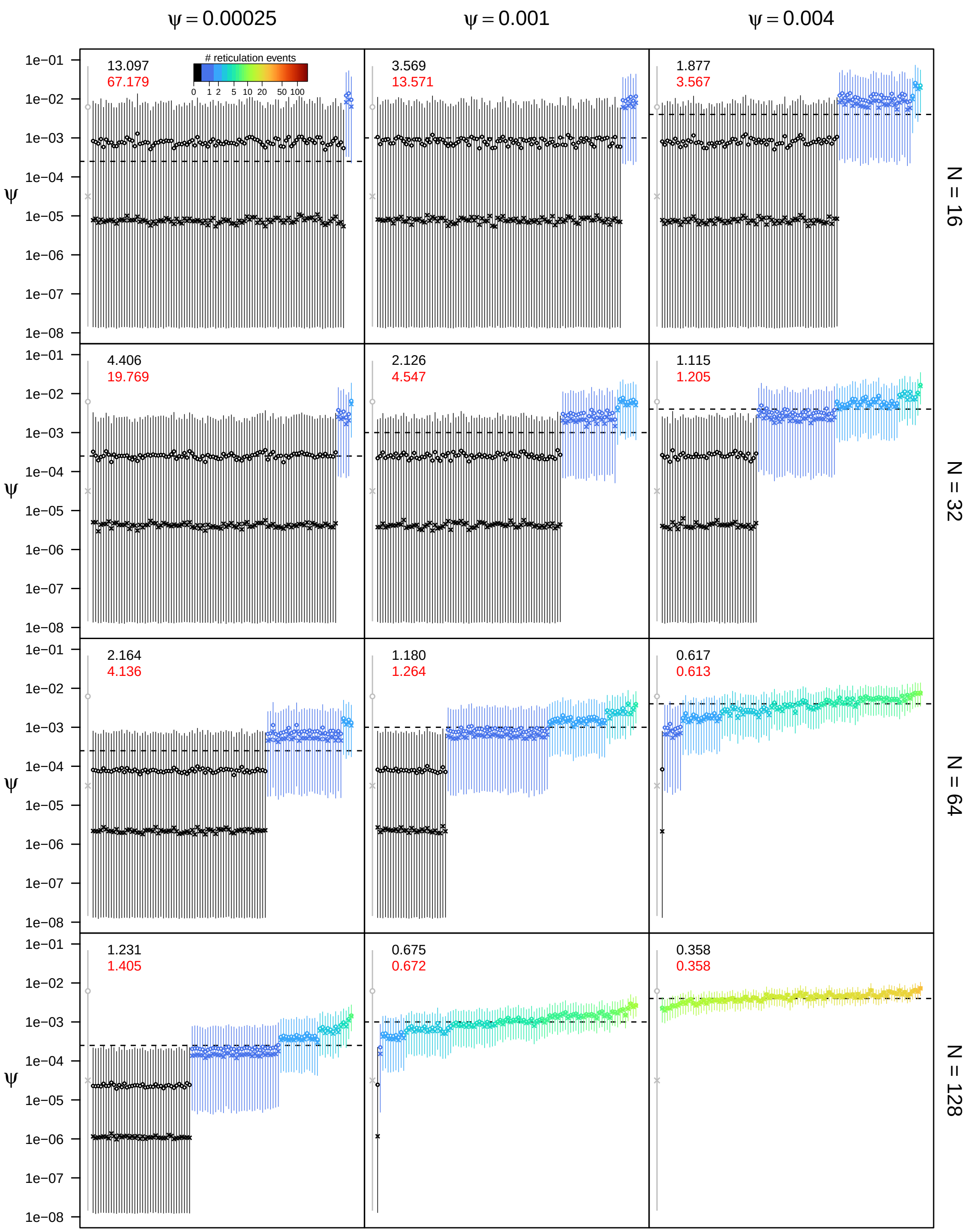


**Figure S88: Posterior estimates of the allopolyploidization rate $\psi$ under the allopolyploidization model for $\epsilon = 0.3$, $\rho = 1.0$, and $u_{\max} = 4N$.** See main text for interpretation.

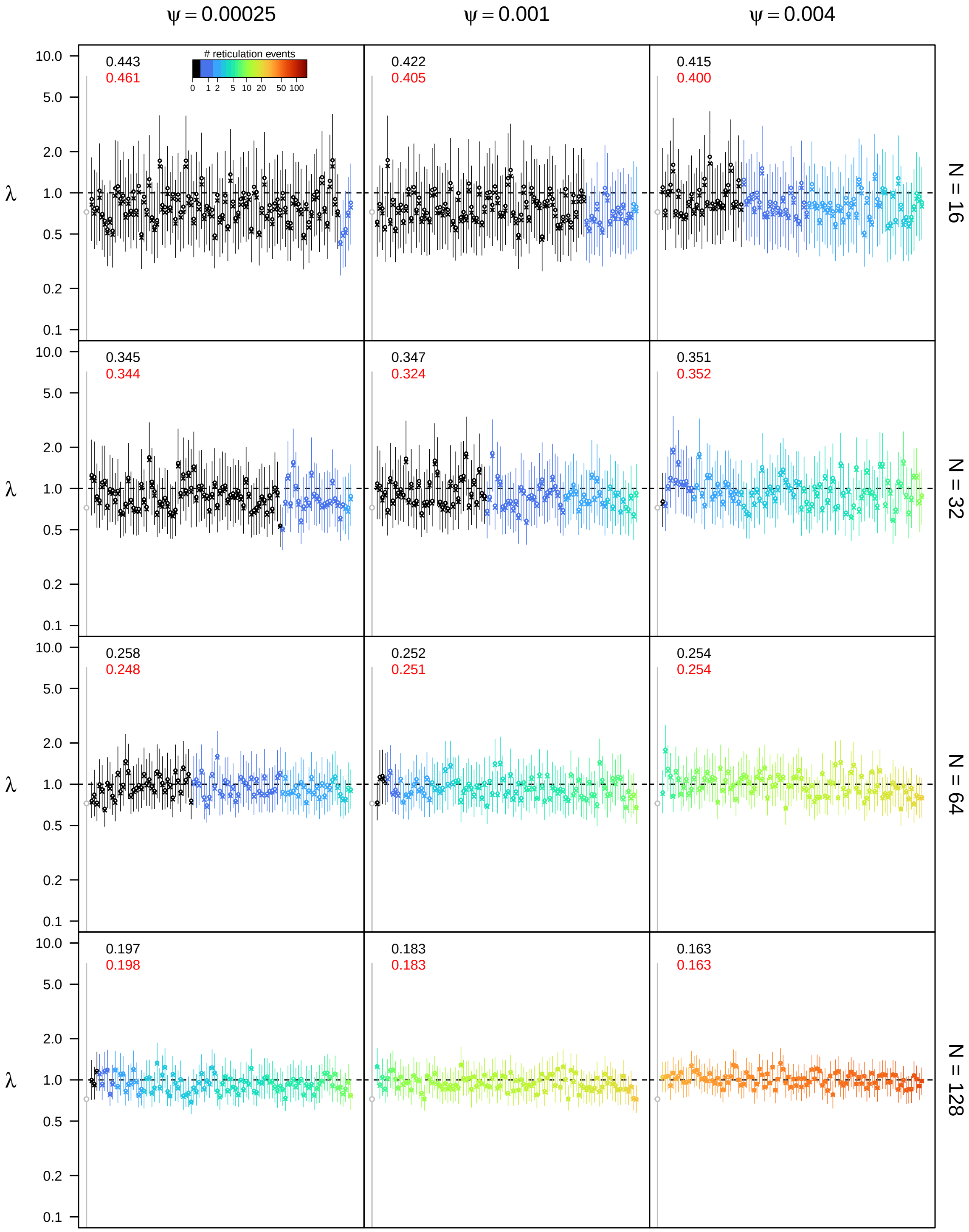


**Figure S89: Posterior estimates of the speciation rate $\lambda$ under the allopolyploidization model for $\epsilon = 0.5$, $\rho = 0.5$, and $u_{\mathbf{max}} = 4N$.** See main text for interpretation.

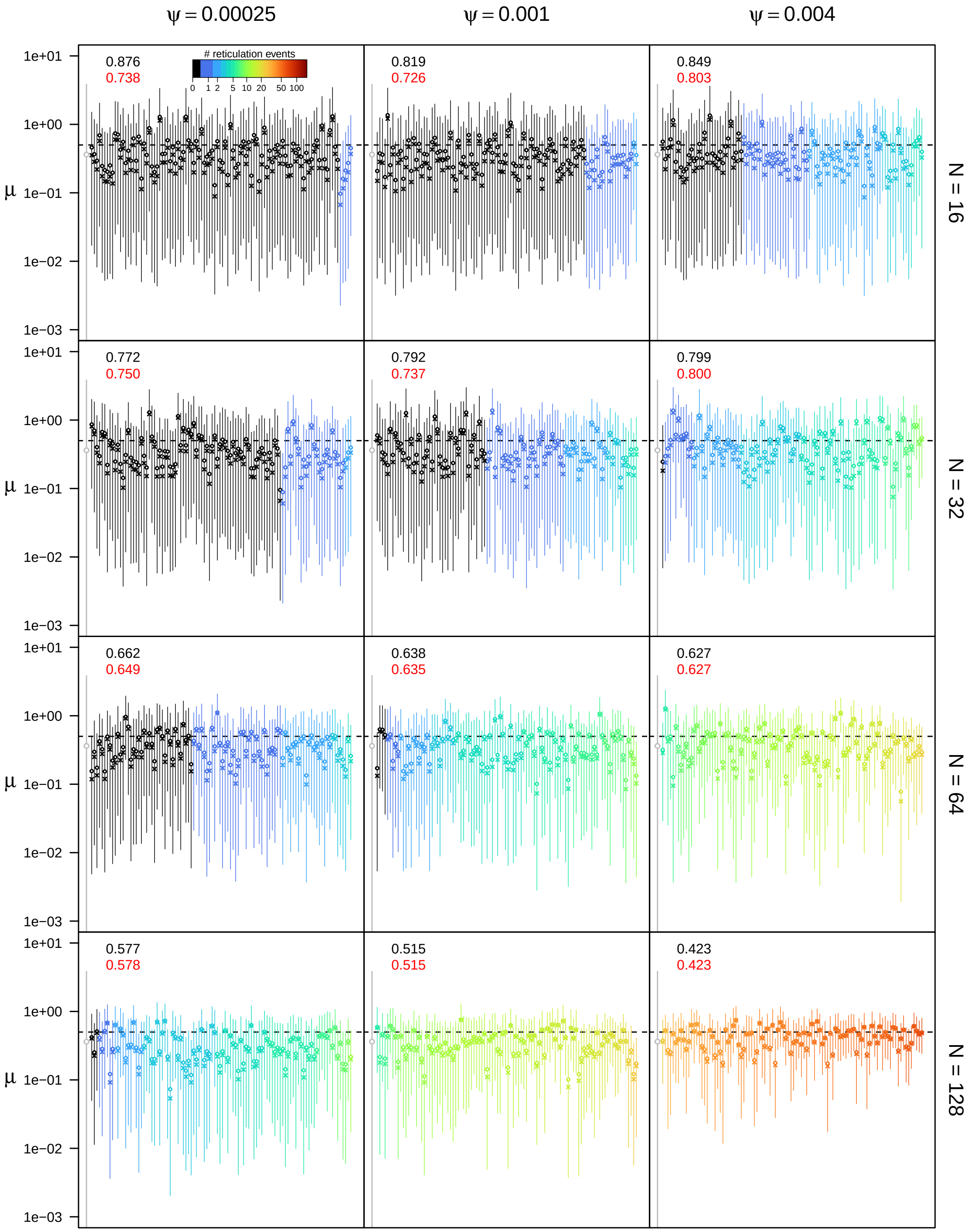


**Figure S90: Posterior estimates of the extinction rate $\mu$ under the allopolyploidization model for $\epsilon = 0.5$, $\rho = 0.5$, and $u_{\mathbf{max}} = 4N$.** See main text for interpretation.

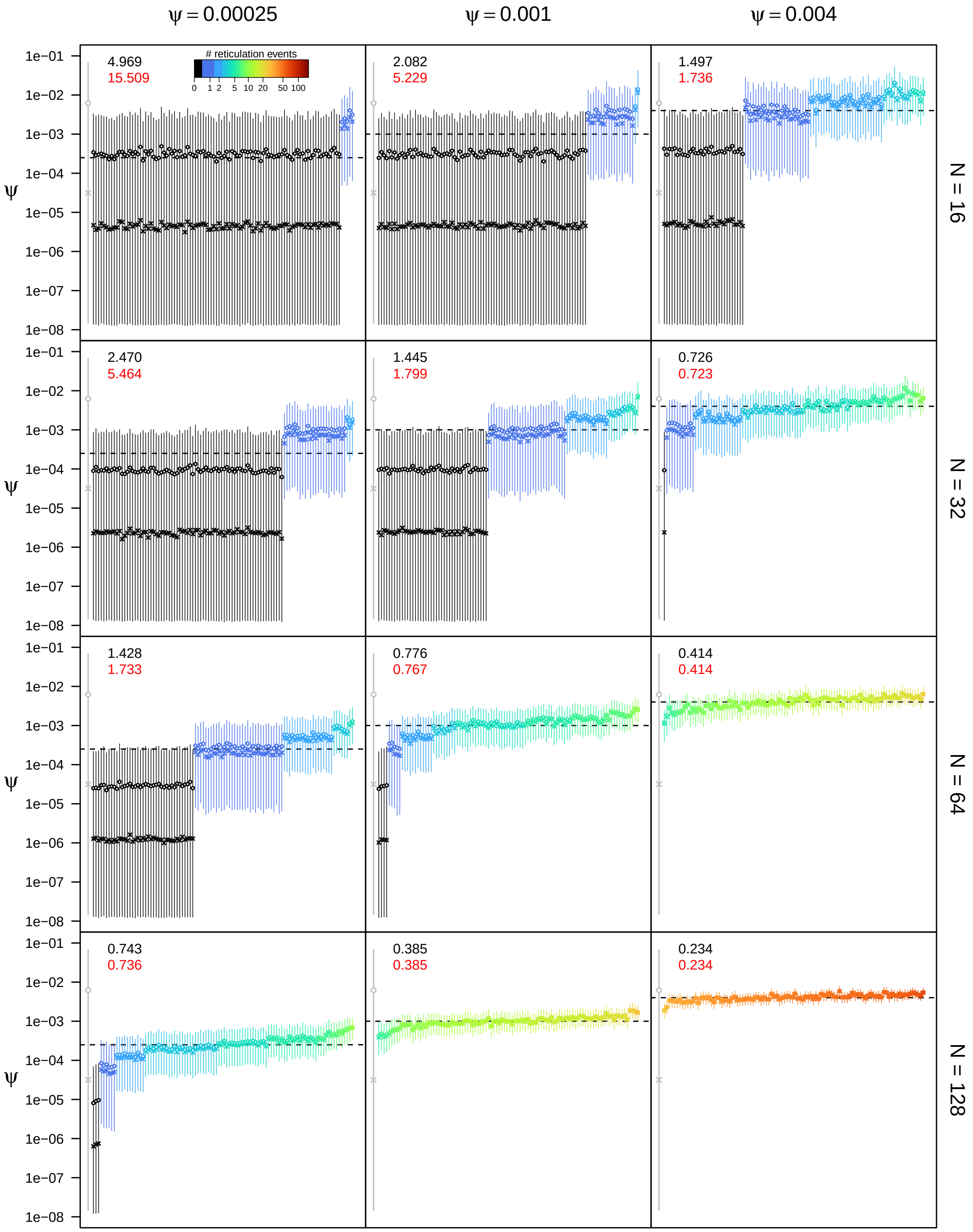


**Figure S91: Posterior estimates of the allopolyploidization rate $\psi$ under the allopolyploidization model for $\epsilon = 0.5$, $\rho = 0.5$, and $u_{\mathbf{max}} = 4N$.** See main text for interpretation.

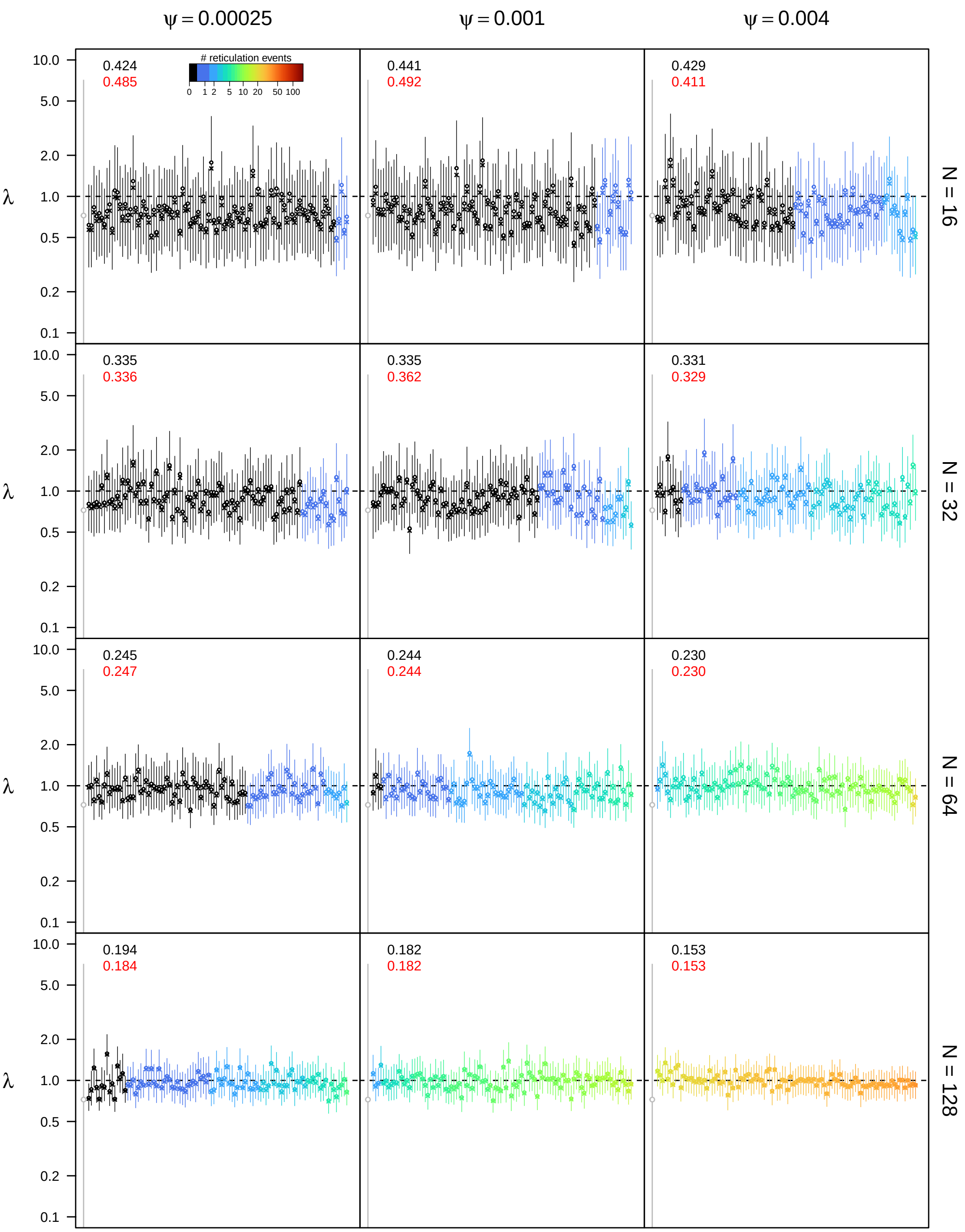


**Figure S92: Posterior estimates of the speciation rate $\lambda$ under the allopolyploidization model for $\epsilon = 0.5$, $\rho = 0.75$, and $u_{\mathbf{max}} = 4N$.** See main text for interpretation.

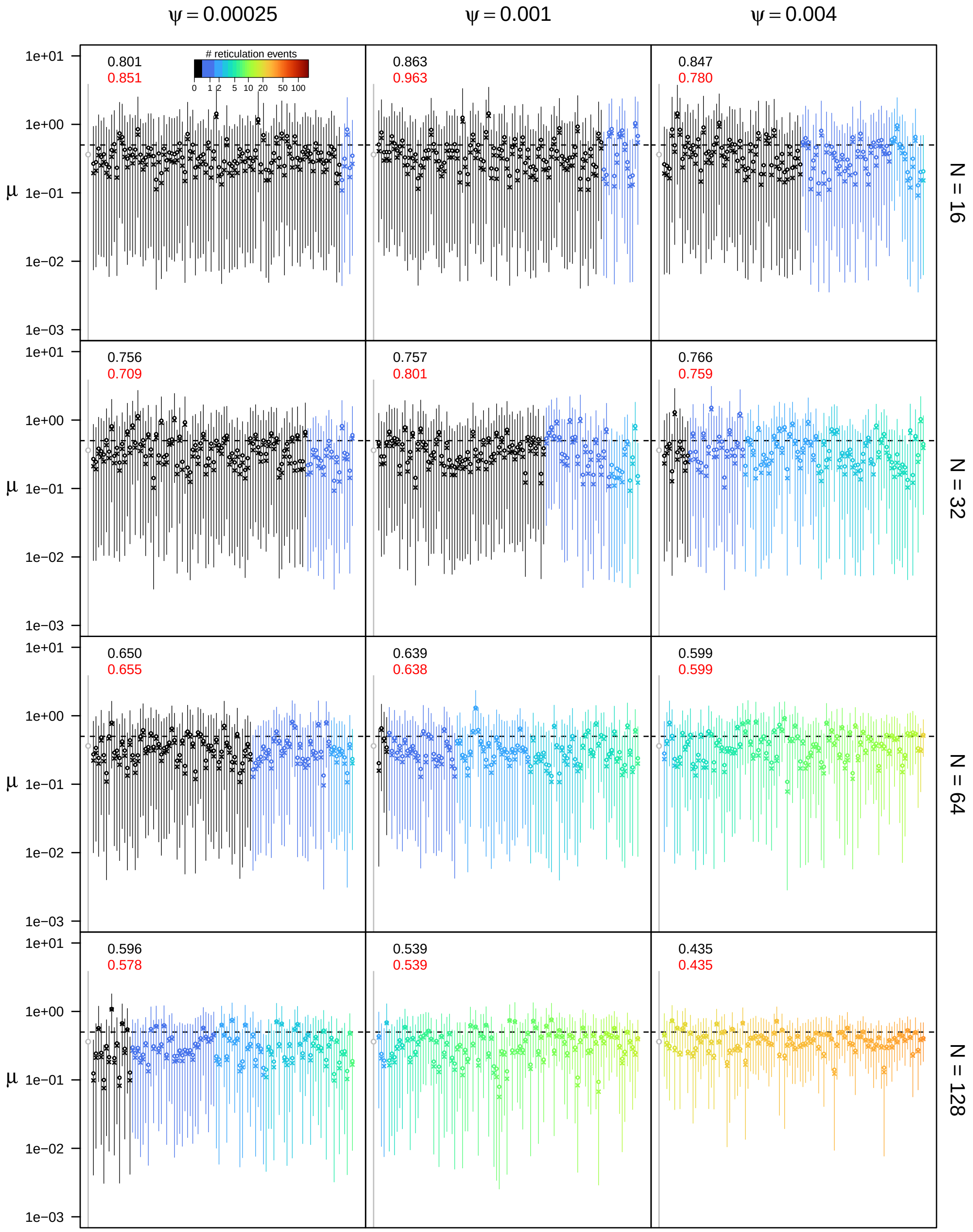


**Figure S93: Posterior estimates of the extinction rate $\mu$ under the allopolyploidization model for $\epsilon = 0.5$, $\rho = 0.75$, and $u_{\mathbf{max}} = 4N$.** See main text for interpretation.

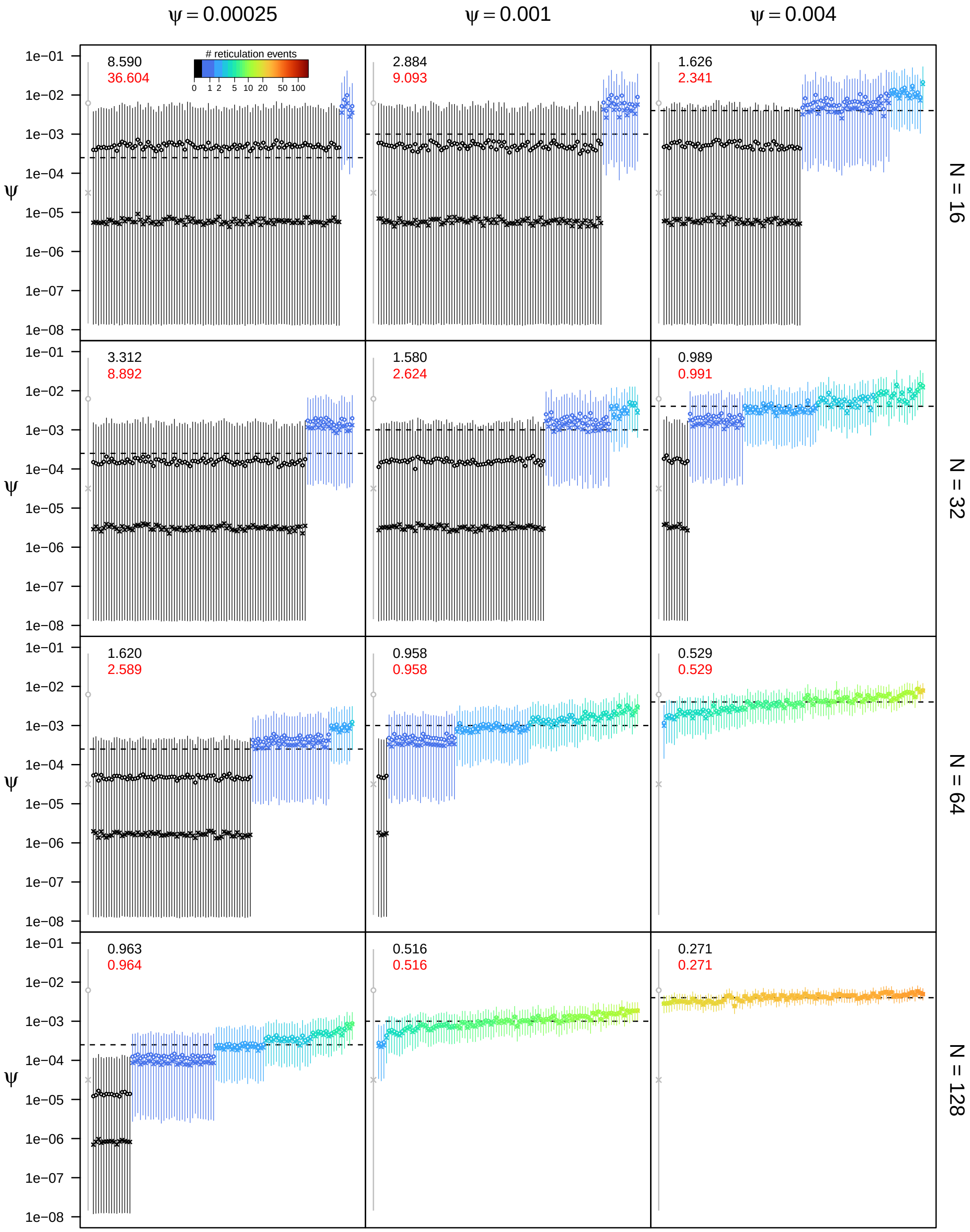


**Figure S94: Posterior estimates of the allopolyploidization rate $\psi$ under the allopolyploidization model for $\epsilon = 0.5$, $\rho = 0.75$, and $u_{\mathbf{max}} = 4N$.** See main text for interpretation.

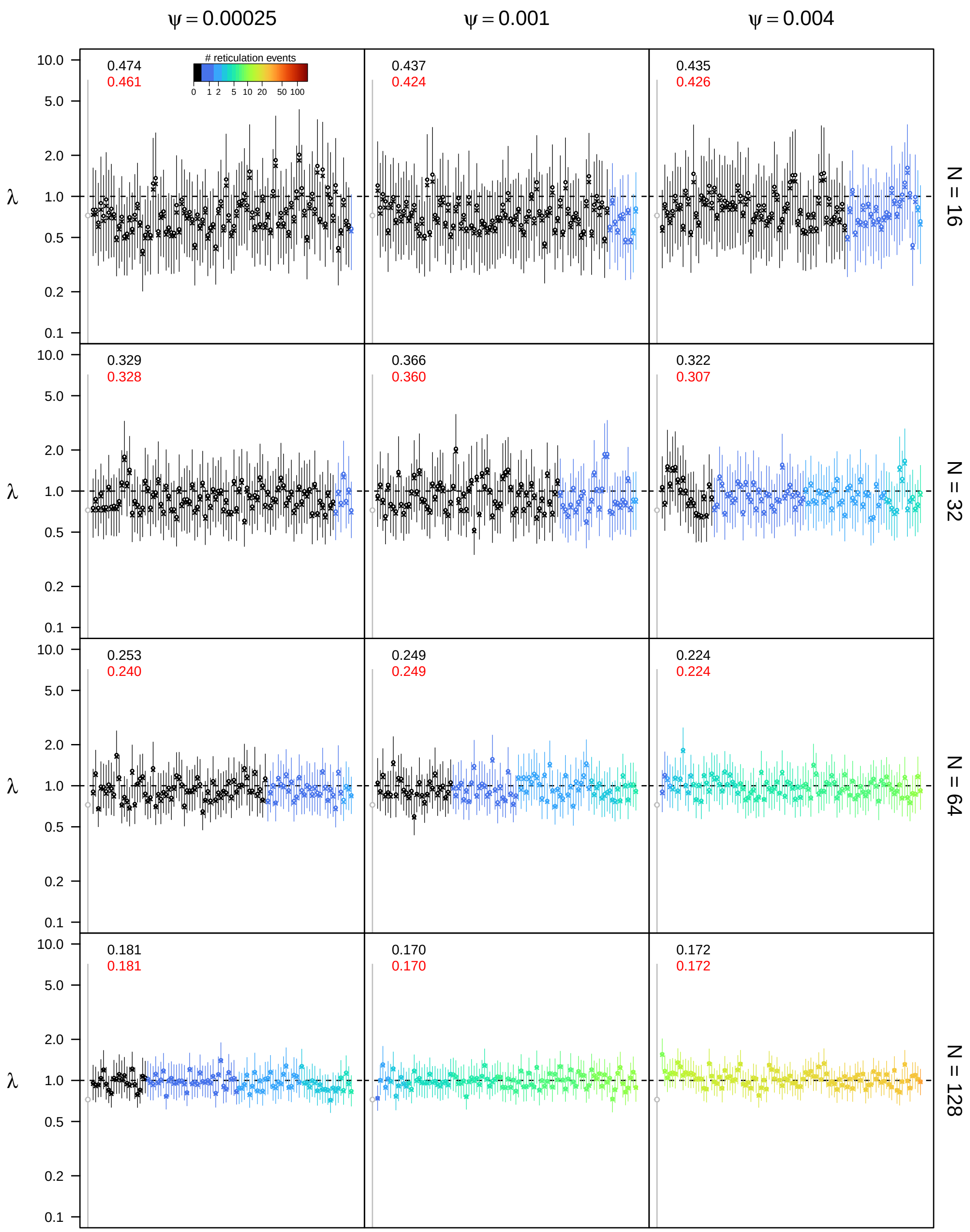


**Figure S95: Posterior estimates of the speciation rate $\lambda$ under the allopolyploidization model for $\epsilon = 0.5$, $\rho = 1.0$, and $u_{\mathbf{max}} = 4N$.** See main text for interpretation.

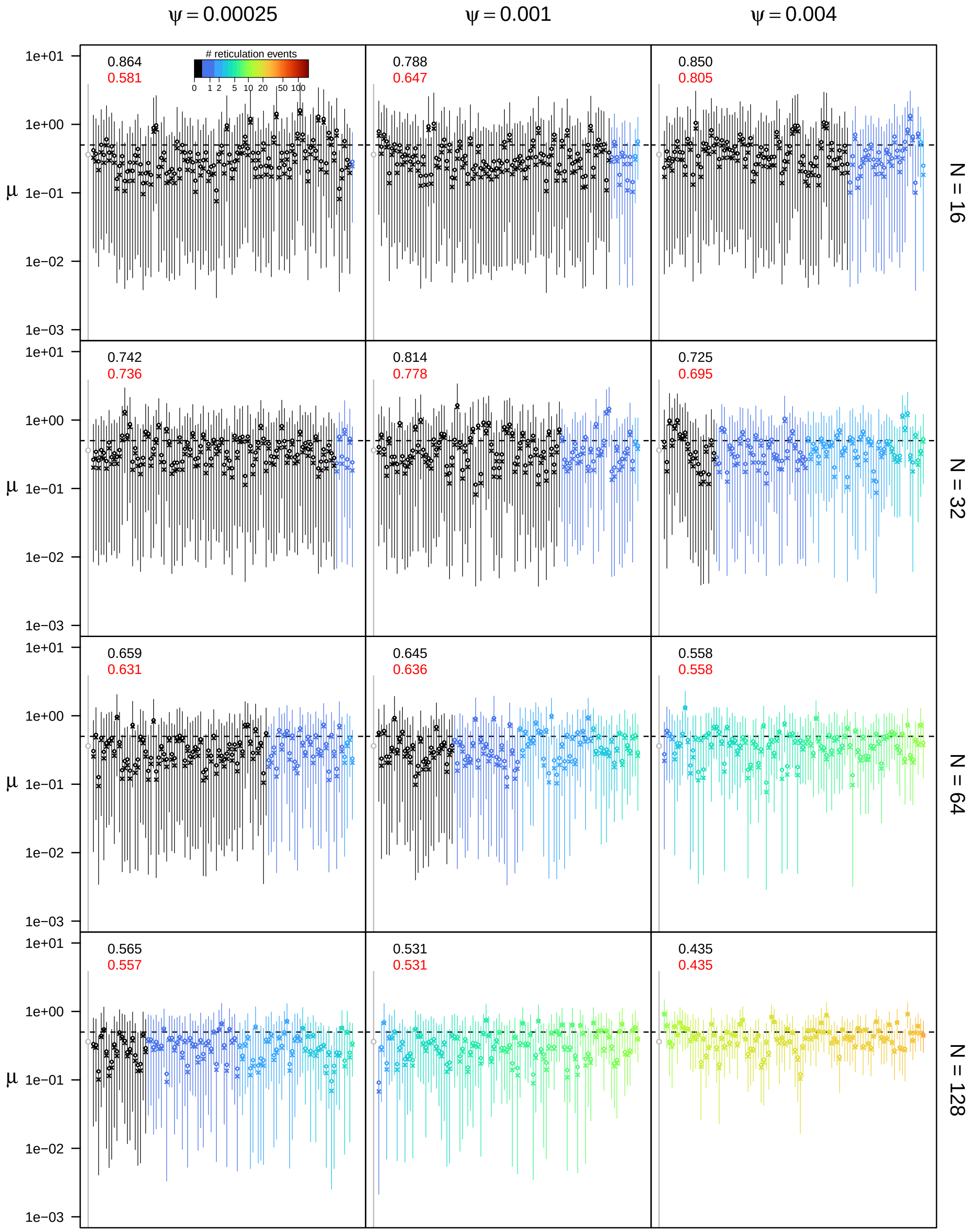


**Figure S96: Posterior estimates of the extinction rate $\mu$ under the allopolyploidization model for $\epsilon = 0.5$, $\rho = 1.0$, and $u_{\mathbf{max}} = 4N$.** See main text for interpretation.

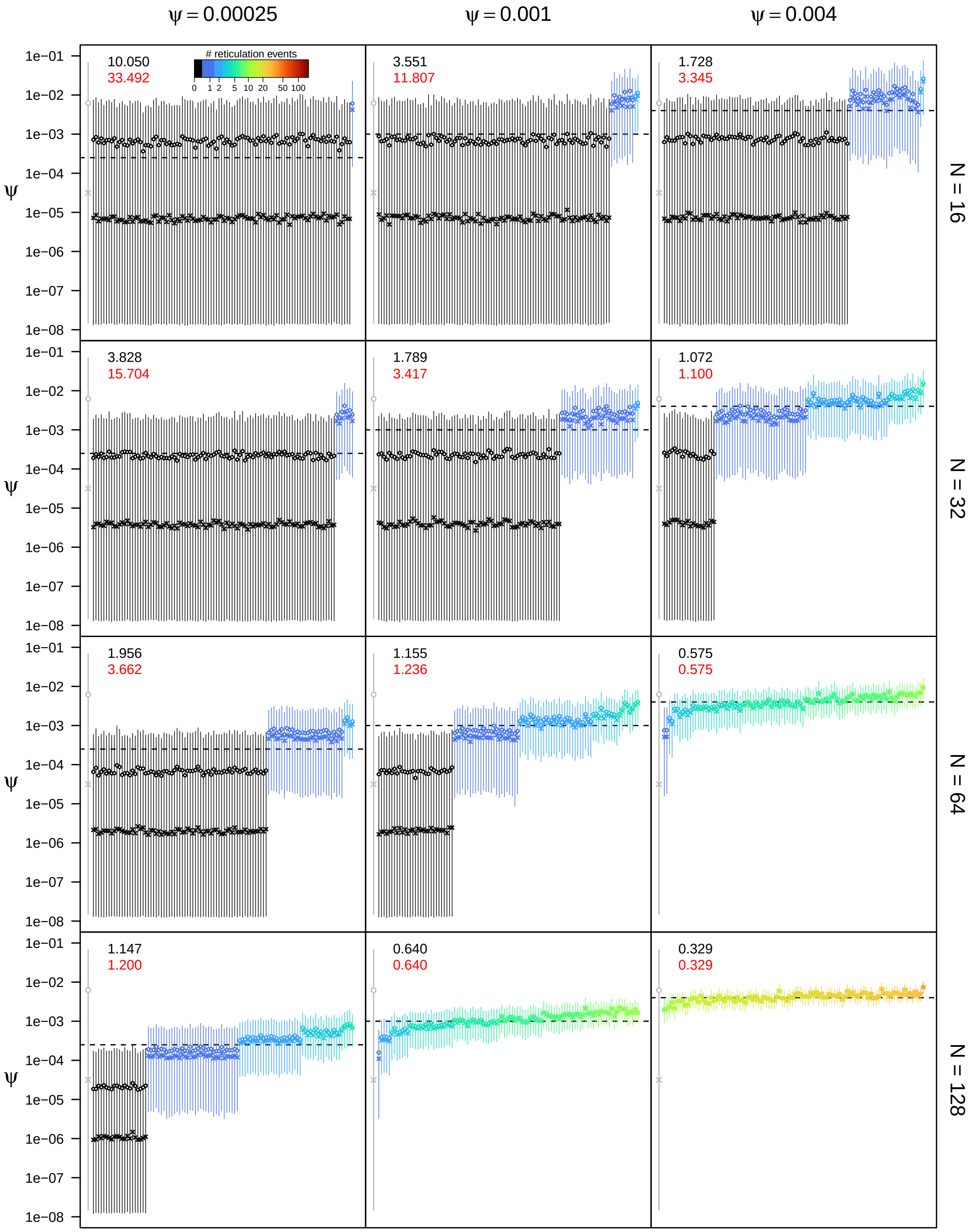


**Figure S97: Posterior estimates of the allopolyploidization rate $\psi$ under the allopolyploidization model for $\epsilon = 0.5$, $\rho = 1.0$, and $u_{\mathbf{max}} = 4N$.** See main text for interpretation.

**Table S2:** Simulation results for $\eta$ with $k = 1$.

| | | | | $\lambda$ | | | $\mu$ | | | $\theta$ | | |
|---|---|---|---|---|---|---|---|---|---|---|---|---|
| $\epsilon$ | $\rho$ | $\theta$ | $N$ | RMSE | $RMSE_c$ | CV | RMSE | $RMSE_c$ | CV | RMSE | $RMSE_c$ | CV |
| 0.1 | 0.5 | 0.00025 | 16 | 0.305 | 0.301 | 0.264 | 3.100 | 3.941 | 0.847 | 8.036 | 36.942 | 3.326 |
| | | | 32 | 0.268 | 0.249 | 0.201 | 3.127 | 2.751 | 0.834 | 3.020 | 7.822 | 2.782 |
| | | | 64 | 0.213 | 0.200 | 0.151 | 2.503 | 2.290 | 0.838 | 1.465 | 2.278 | 2.199 |
| | | | 128 | 0.154 | 0.153 | 0.106 | 1.597 | 1.580 | 0.852 | 0.798 | 0.788 | 0.798 |
| | | 0.00100 | 16 | 0.305 | 0.302 | 0.263 | 2.962 | 2.651 | 0.850 | 3.046 | 7.553 | 2.872 |
| | | | 32 | 0.250 | 0.241 | 0.200 | 3.006 | 2.811 | 0.838 | 1.631 | 2.265 | 2.026 |
| | | | 64 | 0.196 | 0.193 | 0.145 | 2.215 | 2.176 | 0.853 | 0.834 | 0.824 | 0.808 |
| | | | 128 | 0.153 | 0.153 | 0.101 | 1.481 | 1.481 | 0.848 | 0.446 | 0.446 | 0.339 |
| | | 0.00400 | 16 | 0.308 | 0.308 | 0.263 | 3.069 | 2.903 | 0.845 | 1.494 | 2.030 | 2.209 |
| | | | 32 | 0.248 | 0.243 | 0.197 | 2.890 | 2.841 | 0.837 | 0.788 | 0.767 | 0.921 |
| | | | 64 | 0.191 | 0.191 | 0.140 | 2.103 | 2.103 | 0.841 | 0.415 | 0.415 | 0.334 |
| | | | 128 | 0.135 | 0.135 | 0.095 | 1.307 | 1.307 | 0.792 | 0.214 | 0.214 | 0.160 |
| | 0.75 | 0.00025 | 16 | 0.355 | 0.315 | 0.304 | 4.203 | 4.576 | 0.837 | 10.187 | 44.992 | 3.467 |
| | | | 32 | 0.299 | 0.368 | 0.234 | 3.852 | 4.215 | 0.859 | 3.889 | 13.816 | 3.022 |
| | | | 64 | 0.274 | 0.274 | 0.171 | 3.148 | 3.151 | 0.845 | 1.817 | 3.492 | 2.425 |
| | | | 128 | 0.194 | 0.186 | 0.116 | 1.873 | 1.818 | 0.851 | 1.074 | 1.121 | 1.362 |
| | | 0.00100 | 16 | 0.329 | 0.321 | 0.301 | 3.832 | 4.013 | 0.857 | 3.308 | 13.900 | 3.268 |
| | | | 32 | 0.337 | 0.328 | 0.237 | 4.296 | 3.889 | 0.840 | 1.648 | 2.939 | 2.557 |
| | | | 64 | 0.247 | 0.242 | 0.166 | 2.871 | 2.823 | 0.851 | 0.946 | 0.945 | 1.035 |
| | | | 128 | 0.176 | 0.176 | 0.110 | 1.645 | 1.645 | 0.841 | 0.562 | 0.562 | 0.430 |
| | | 0.00400 | 16 | 0.355 | 0.333 | 0.303 | 4.143 | 3.252 | 0.845 | 1.698 | 2.977 | 2.629 |
| | | | 32 | 0.315 | 0.300 | 0.232 | 4.009 | 3.778 | 0.845 | 0.992 | 0.999 | 1.272 |
| | | | 64 | 0.243 | 0.241 | 0.160 | 2.691 | 2.661 | 0.840 | 0.528 | 0.523 | 0.449 |
| | | | 128 | 0.153 | 0.153 | 0.101 | 1.321 | 1.321 | 0.831 | 0.285 | 0.285 | 0.208 |
| | 1.0 | 0.00025 | 16 | 0.377 | 0.310 | 0.331 | 4.473 | 3.941 | 0.853 | 13.826 | 55.190 | 3.504 |
| | | | 32 | 0.323 | 0.249 | 0.243 | 4.157 | 2.960 | 0.865 | 4.866 | 19.897 | 3.194 |
| | | | 64 | 0.257 | 0.230 | 0.172 | 3.005 | 2.833 | 0.855 | 2.094 | 4.919 | 2.635 |
| | | | 128 | 0.194 | 0.175 | 0.116 | 1.827 | 1.646 | 0.857 | 1.192 | 1.362 | 1.706 |
| | | 0.00100 | 16 | 0.370 | 0.359 | 0.327 | 4.322 | 3.653 | 0.865 | 4.359 | 17.231 | 3.336 |
| | | | 32 | 0.333 | 0.321 | 0.247 | 4.331 | 3.871 | 0.850 | 1.809 | 4.452 | 2.904 |
| | | | 64 | 0.265 | 0.243 | 0.170 | 3.002 | 2.713 | 0.856 | 1.048 | 1.132 | 1.978 |
| | | | 128 | 0.181 | 0.180 | 0.112 | 1.625 | 1.604 | 0.854 | 0.661 | 0.651 | 0.625 |
| | | 0.00400 | 16 | 0.375 | 0.350 | 0.327 | 4.325 | 3.676 | 0.862 | 2.145 | 4.262 | 2.738 |
| | | | 32 | 0.345 | 0.330 | 0.241 | 4.275 | 4.041 | 0.854 | 1.084 | 1.163 | 1.858 |
| | | | 64 | 0.236 | 0.236 | 0.161 | 2.567 | 2.570 | 0.853 | 0.631 | 0.628 | 0.542 |
| | | | 128 | 0.169 | 0.169 | 0.103 | 1.382 | 1.382 | 0.784 | 0.317 | 0.317 | 0.255 |
| 0.3 | 0.5 | 0.00025 | 16 | 0.340 | 0.410 | 0.268 | 0.784 | 0.698 | 0.835 | 6.626 | 24.569 | 3.319 |
| | | | 32 | 0.265 | 0.269 | 0.210 | 0.842 | 0.768 | 0.797 | 2.615 | 6.790 | 2.794 |
| | | | 64 | 0.209 | 0.204 | 0.159 | 0.767 | 0.735 | 0.798 | 1.475 | 1.951 | 1.893 |
| | | | 128 | 0.159 | 0.160 | 0.121 | 0.682 | 0.684 | 0.759 | 0.824 | 0.820 | 0.663 |
| | | 0.00100 | 16 | 0.346 | 0.361 | 0.265 | 0.768 | 0.736 | 0.837 | 2.947 | 6.589 | 2.734 |
| | | | 32 | 0.258 | 0.261 | 0.204 | 0.812 | 0.774 | 0.809 | 1.551 | 1.960 | 1.829 |
| | | | 64 | 0.188 | 0.188 | 0.157 | 0.719 | 0.712 | 0.792 | 0.802 | 0.792 | 0.784 |
| | | | 128 | 0.149 | 0.149 | 0.116 | 0.661 | 0.661 | 0.740 | 0.408 | 0.408 | 0.301 |
| | | 0.00400 | 16 | 0.350 | 0.358 | 0.264 | 0.770 | 0.763 | 0.830 | 1.425 | 1.731 | 1.906 |
| | | | 32 | 0.258 | 0.256 | 0.206 | 0.820 | 0.810 | 0.785 | 0.787 | 0.780 | 0.740 |
| | | | 64 | 0.191 | 0.191 | 0.155 | 0.725 | 0.725 | 0.720 | 0.407 | 0.407 | 0.304 |
| | | | 128 | 0.151 | 0.151 | 0.108 | 0.629 | 0.629 | 0.586 | 0.217 | 0.217 | 0.149 |
| | 0.75 | 0.00025 | 16 | 0.354 | 0.395 | 0.309 | 1.054 | 1.151 | 0.811 | 10.964 | 54.781 | 3.392 |
| | | | 32 | 0.305 | 0.270 | 0.242 | 1.080 | 0.969 | 0.821 | 4.027 | 12.082 | 2.888 |
| | | | 64 | 0.227 | 0.222 | 0.181 | 0.880 | 0.857 | 0.803 | 1.613 | 3.201 | 2.495 |
| | | | 128 | 0.169 | 0.158 | 0.126 | 0.721 | 0.698 | 0.784 | 0.954 | 0.954 | 1.210 |
| | | 0.00100 | 16 | 0.353 | 0.316 | 0.306 | 0.983 | 0.931 | 0.821 | 3.538 | 10.651 | 3.073 |
| | | | 32 | 0.311 | 0.290 | 0.246 | 1.118 | 1.049 | 0.801 | 1.453 | 2.635 | 2.664 |
| | | | 64 | 0.232 | 0.227 | 0.180 | 0.881 | 0.857 | 0.789 | 0.997 | 1.003 | 1.073 |
| | | | 128 | 0.176 | 0.176 | 0.126 | 0.726 | 0.726 | 0.718 | 0.515 | 0.515 | 0.402 |
| | | 0.00400 | 16 | 0.351 | 0.341 | 0.307 | 1.015 | 0.950 | 0.810 | 1.646 | 2.426 | 2.355 |
| | | | 32 | 0.294 | 0.282 | 0.239 | 1.073 | 1.021 | 0.797 | 0.923 | 0.915 | 1.165 |
| | | | 64 | 0.214 | 0.214 | 0.170 | 0.839 | 0.837 | 0.752 | 0.524 | 0.514 | 0.461 |
| | | | 128 | 0.156 | 0.156 | 0.114 | 0.630 | 0.630 | 0.636 | 0.254 | 0.254 | 0.196 |
| | 1.0 | 0.00025 | 16 | 0.366 | 0.320 | 0.333 | 1.066 | 1.123 | 0.845 | 12.259 | 71.319 | 3.507 |
| | | | 32 | 0.323 | 0.268 | 0.253 | 1.174 | 0.877 | 0.822 | 4.756 | 15.646 | 3.079 |
| | | | 64 | 0.229 | 0.207 | 0.180 | 0.868 | 0.777 | 0.811 | 2.289 | 4.543 | 2.416 |
| | | | 128 | 0.165 | 0.158 | 0.127 | 0.710 | 0.699 | 0.771 | 1.213 | 1.322 | 1.426 |
| | | 0.00100 | 16 | 0.385 | 0.353 | 0.331 | 1.084 | 1.044 | 0.839 | 4.196 | 14.067 | 3.227 |
| | | | 32 | 0.331 | 0.335 | 0.254 | 1.193 | 1.169 | 0.824 | 1.949 | 3.927 | 2.657 |
| | | | 64 | 0.240 | 0.209 | 0.181 | 0.924 | 0.830 | 0.790 | 1.195 | 1.325 | 1.620 |
| | | | 128 | 0.165 | 0.165 | 0.121 | 0.670 | 0.670 | 0.730 | 0.644 | 0.644 | 0.499 |
| | | 0.00400 | 16 | 0.376 | 0.358 | 0.329 | 1.067 | 1.003 | 0.837 | 1.780 | 3.407 | 2.766 |
| | | | 32 | 0.329 | 0.304 | 0.253 | 1.221 | 1.102 | 0.799 | 1.070 | 1.122 | 1.666 |
| | | | 64 | 0.218 | 0.218 | 0.168 | 0.819 | 0.820 | 0.736 | 0.566 | 0.561 | 0.521 |
| | | | 128 | 0.158 | 0.158 | 0.112 | 0.584 | 0.584 | 0.555 | 0.328 | 0.328 | 0.237 |
| 0.5 | 0.5 | 0.00025 | 16 | 0.399 | 0.377 | 0.276 | 0.661 | 0.639 | 0.793 | 6.080 | 26.478 | 3.292 |
| | | | 32 | 0.293 | 0.309 | 0.218 | 0.632 | 0.642 | 0.748 | 2.400 | 6.118 | 2.762 |
| | | | 64 | 0.246 | 0.245 | 0.171 | 0.639 | 0.635 | 0.715 | 1.498 | 1.767 | 1.551 |
| | | | 128 | 0.189 | 0.189 | 0.136 | 0.590 | 0.592 | 0.638 | 0.814 | 0.806 | 0.658 |
| | | 0.00100 | 16 | 0.397 | 0.421 | 0.275 | 0.667 | 0.675 | 0.793 | 2.586 | 5.679 | 2.756 |
| | | | 32 | 0.303 | 0.303 | 0.216 | 0.638 | 0.627 | 0.725 | 1.428 | 1.639 | 1.590 |
| | | | 64 | 0.228 | 0.229 | 0.174 | 0.591 | 0.592 | 0.661 | 0.741 | 0.739 | 0.588 |
| | | | 128 | 0.180 | 0.180 | 0.131 | 0.534 | 0.534 | 0.542 | 0.381 | 0.381 | 0.271 |
| | | 0.00400 | 16 | 0.406 | 0.413 | 0.272 | 0.660 | 0.654 | 0.765 | 1.512 | 1.769 | 1.659 |
| | | | 32 | 0.296 | 0.297 | 0.211 | 0.620 | 0.621 | 0.700 | 0.779 | 0.773 | 0.640 |
| | | | 64 | 0.222 | 0.222 | 0.165 | 0.537 | 0.537 | 0.545 | 0.402 | 0.402 | 0.270 |
| | | | 128 | 0.151 | 0.151 | 0.113 | 0.414 | 0.414 | 0.371 | 0.205 | 0.205 | 0.136 |
| | 0.75 | 0.00025 | 16 | 0.379 | 0.398 | 0.314 | 0.672 | 0.637 | 0.774 | 8.360 | 34.104 | 3.436 |
| | | | 32 | 0.294 | 0.281 | 0.255 | 0.672 | 0.650 | 0.758 | 2.798 | 9.354 | 3.028 |
| | | | 64 | 0.264 | 0.275 | 0.200 | 0.682 | 0.696 | 0.690 | 1.972 | 2.769 | 1.875 |
| | | | 128 | 0.190 | 0.190 | 0.145 | 0.601 | 0.600 | 0.653 | 0.926 | 0.922 | 0.920 |
| | | 0.00100 | 16 | 0.391 | 0.408 | 0.312 | 0.672 | 0.665 | 0.777 | 3.288 | 7.915 | 2.925 |
| | | | 32 | 0.310 | 0.298 | 0.253 | 0.695 | 0.662 | 0.729 | 1.810 | 2.479 | 1.978 |
| | | | 64 | 0.256 | 0.252 | 0.196 | 0.661 | 0.656 | 0.686 | 0.888 | 0.877 | 1.033 |
| | | | 128 | 0.178 | 0.178 | 0.141 | 0.542 | 0.542 | 0.566 | 0.558 | 0.558 | 0.371 |
| | | 0.00400 | 16 | 0.385 | 0.401 | 0.309 | 0.670 | 0.670 | 0.758 | 1.747 | 2.248 | 1.935 |
| | | | 32 | 0.302 | 0.296 | 0.248 | 0.683 | 0.669 | 0.687 | 0.882 | 0.871 | 1.026 |
| | | | 64 | 0.237 | 0.237 | 0.183 | 0.603 | 0.603 | 0.626 | 0.467 | 0.467 | 0.365 |
| | | | 128 | 0.174 | 0.174 | 0.122 | 0.443 | 0.443 | 0.401 | 0.249 | 0.249 | 0.181 |
| | 1.0 | 0.00025 | 16 | 0.403 | 0.357 | 0.339 | 0.711 | 0.651 | 0.790 | 13.329 | 55.057 | 3.370 |
| | | | 32 | 0.318 | 0.309 | 0.266 | 0.728 | 0.674 | 0.767 | 4.395 | 13.749 | 3.056 |
| | | | 64 | 0.255 | 0.252 | 0.198 | 0.672 | 0.674 | 0.727 | 2.032 | 3.728 | 2.379 |
| | | | 128 | 0.187 | 0.188 | 0.141 | 0.596 | 0.601 | 0.655 | 1.131 | 1.179 | 1.205 |
| | | 0.00100 | 16 | 0.402 | 0.388 | 0.338 | 0.711 | 0.672 | 0.786 | 3.836 | 12.381 | 3.223 |
| | | | 32 | 0.336 | 0.316 | 0.269 | 0.757 | 0.726 | 0.729 | 1.783 | 3.559 | 2.677 |
| | | | 64 | 0.261 | 0.256 | 0.194 | 0.662 | 0.658 | 0.680 | 1.168 | 1.212 | 1.191 |
| | | | 128 | 0.173 | 0.173 | 0.134 | 0.534 | 0.534 | 0.561 | 0.609 | 0.609 | 0.446 |
| | | 0.00400 | 16 | 0.395 | 0.413 | 0.334 | 0.683 | 0.653 | 0.770 | 1.684 | 2.742 | 2.593 |
| | | | 32 | 0.323 | 0.321 | 0.259 | 0.732 | 0.724 | 0.692 | 1.092 | 1.124 | 1.299 |
| | | | 64 | 0.237 | 0.237 | 0.183 | 0.596 | 0.596 | 0.601 | 0.598 | 0.598 | 0.463 |
| | | | 128 | 0.168 | 0.168 | 0.119 | 0.461 | 0.461 | 0.415 | 0.291 | 0.291 | 0.221 |

**Table S3:** Simulation results for $\eta$ with $k = 2$.

| | | | | $\lambda$ | | | $\mu$ | | | $\theta$ | | |
|---|---|---|---|---|---|---|---|---|---|---|---|---|
| $\epsilon$ | $\rho$ | $\theta$ | $N$ | RMSE | $RMSE_c$ | CV | RMSE | $RMSE_c$ | CV | RMSE | $RMSE_c$ | CV |
| 0.1 | 0.5 | 0.00025 | 16 | 0.386 | 0.460 | 0.318 | 4.969 | 6.289 | 0.895 | 7.119 | 33.067 | 3.367 |
| | | | 32 | 0.360 | 0.319 | 0.241 | 4.471 | 3.875 | 0.872 | 2.785 | 7.141 | 2.790 |
| | | | 64 | 0.270 | 0.249 | 0.171 | 3.142 | 2.841 | 0.850 | 1.402 | 2.123 | 2.189 |
| | | | 128 | 0.184 | 0.183 | 0.115 | 1.844 | 1.822 | 0.850 | 0.770 | 0.758 | 0.801 |
| | | 0.00100 | 16 | 0.379 | 0.343 | 0.314 | 4.748 | 4.092 | 0.898 | 2.716 | 6.689 | 2.897 |
| | | | 32 | 0.338 | 0.311 | 0.239 | 4.290 | 3.957 | 0.878 | 1.512 | 2.038 | 2.031 |
| | | | 64 | 0.241 | 0.236 | 0.162 | 2.716 | 2.660 | 0.864 | 0.795 | 0.782 | 0.814 |
| | | | 128 | 0.180 | 0.180 | 0.108 | 1.663 | 1.663 | 0.846 | 0.435 | 0.435 | 0.342 |
| | | 0.00400 | 16 | 0.387 | 0.366 | 0.313 | 4.835 | 4.520 | 0.890 | 1.369 | 1.783 | 2.224 |
| | | | 32 | 0.319 | 0.311 | 0.230 | 3.977 | 3.874 | 0.874 | 0.744 | 0.718 | 0.926 |
| | | | 64 | 0.226 | 0.226 | 0.152 | 2.453 | 2.453 | 0.853 | 0.405 | 0.405 | 0.340 |
| | | | 128 | 0.151 | 0.151 | 0.098 | 1.369 | 1.369 | 0.799 | 0.213 | 0.213 | 0.163 |
| | 0.75 | 0.00025 | 16 | 0.420 | 0.377 | 0.344 | 5.169 | 5.512 | 0.906 | 9.972 | 43.835 | 3.490 |
| | | | 32 | 0.314 | 0.389 | 0.244 | 4.059 | 4.460 | 0.884 | 3.871 | 13.799 | 3.018 |
| | | | 64 | 0.275 | 0.276 | 0.171 | 3.157 | 3.161 | 0.846 | 1.807 | 3.467 | 2.414 |
| | | | 128 | 0.194 | 0.186 | 0.116 | 1.875 | 1.820 | 0.852 | 1.074 | 1.121 | 1.360 |
| | | 0.00100 | 16 | 0.374 | 0.361 | 0.335 | 4.588 | 4.669 | 0.922 | 3.238 | 13.514 | 3.268 |
| | | | 32 | 0.360 | 0.344 | 0.249 | 4.598 | 4.092 | 0.866 | 1.656 | 2.957 | 2.567 |
| | | | 64 | 0.247 | 0.243 | 0.167 | 2.883 | 2.836 | 0.853 | 0.949 | 0.948 | 1.027 |
| | | | 128 | 0.180 | 0.180 | 0.110 | 1.743 | 1.743 | 0.835 | 0.564 | 0.564 | 0.430 |
| | | 0.00400 | 16 | 0.409 | 0.358 | 0.338 | 4.985 | 3.754 | 0.908 | 1.681 | 2.938 | 2.638 |
| | | | 32 | 0.327 | 0.310 | 0.239 | 4.171 | 3.906 | 0.865 | 0.993 | 1.000 | 1.283 |
| | | | 64 | 0.243 | 0.241 | 0.159 | 2.693 | 2.663 | 0.837 | 0.527 | 0.523 | 0.447 |
| | | | 128 | 0.153 | 0.153 | 0.101 | 1.320 | 1.320 | 0.831 | 0.285 | 0.285 | 0.208 |
| | 1.0 | 0.00025 | 16 | 0.414 | 0.322 | 0.356 | 4.988 | 4.246 | 0.900 | 13.821 | 53.698 | 3.523 |
| | | | 32 | 0.330 | 0.247 | 0.247 | 4.243 | 2.905 | 0.872 | 4.863 | 20.084 | 3.195 |
| | | | 64 | 0.257 | 0.230 | 0.172 | 3.011 | 2.850 | 0.854 | 2.097 | 4.917 | 2.659 |
| | | | 128 | 0.194 | 0.176 | 0.116 | 1.826 | 1.647 | 0.860 | 1.189 | 1.357 | 1.701 |
| | | 0.00100 | 16 | 0.401 | 0.375 | 0.350 | 4.764 | 3.878 | 0.906 | 4.345 | 17.186 | 3.336 |
| | | | 32 | 0.338 | 0.324 | 0.250 | 4.396 | 3.912 | 0.858 | 1.814 | 4.446 | 2.907 |
| | | | 64 | 0.264 | 0.242 | 0.169 | 2.992 | 2.711 | 0.854 | 1.046 | 1.128 | 1.964 |
| | | | 128 | 0.181 | 0.180 | 0.111 | 1.622 | 1.601 | 0.854 | 0.662 | 0.652 | 0.627 |
| | | 0.00400 | 16 | 0.406 | 0.365 | 0.349 | 4.757 | 3.916 | 0.901 | 2.130 | 4.248 | 2.716 |
| | | | 32 | 0.350 | 0.333 | 0.244 | 4.342 | 4.096 | 0.861 | 1.080 | 1.156 | 1.873 |
| | | | 64 | 0.236 | 0.236 | 0.161 | 2.569 | 2.571 | 0.853 | 0.632 | 0.628 | 0.541 |
| | | | 128 | 0.169 | 0.169 | 0.104 | 1.382 | 1.382 | 0.784 | 0.317 | 0.317 | 0.256 |
| 0.3 | 0.5 | 0.00025 | 16 | 0.366 | 0.369 | 0.325 | 1.197 | 0.839 | 0.879 | 5.886 | 21.983 | 3.325 |
| | | | 32 | 0.331 | 0.304 | 0.258 | 1.221 | 1.039 | 0.835 | 2.407 | 6.116 | 2.817 |
| | | | 64 | 0.243 | 0.228 | 0.182 | 0.911 | 0.848 | 0.812 | 1.405 | 1.820 | 1.888 |
| | | | 128 | 0.179 | 0.179 | 0.132 | 0.727 | 0.728 | 0.755 | 0.791 | 0.785 | 0.665 |
| | | 0.00100 | 16 | 0.359 | 0.357 | 0.319 | 1.139 | 1.057 | 0.885 | 2.597 | 5.708 | 2.755 |
| | | | 32 | 0.314 | 0.300 | 0.248 | 1.133 | 1.034 | 0.848 | 1.434 | 1.762 | 1.849 |
| | | | 64 | 0.216 | 0.213 | 0.178 | 0.836 | 0.820 | 0.803 | 0.769 | 0.756 | 0.785 |
| | | | 128 | 0.159 | 0.159 | 0.125 | 0.688 | 0.688 | 0.735 | 0.394 | 0.394 | 0.304 |
| | | 0.00400 | 16 | 0.367 | 0.368 | 0.317 | 1.140 | 1.095 | 0.875 | 1.289 | 1.506 | 1.928 |
| | | | 32 | 0.311 | 0.306 | 0.246 | 1.140 | 1.119 | 0.822 | 0.740 | 0.730 | 0.753 |
| | | | 64 | 0.216 | 0.216 | 0.171 | 0.832 | 0.832 | 0.735 | 0.395 | 0.395 | 0.311 |
| | | | 128 | 0.161 | 0.161 | 0.112 | 0.655 | 0.655 | 0.598 | 0.210 | 0.210 | 0.152 |
| | 0.75 | 0.00025 | 16 | 0.408 | 0.446 | 0.354 | 1.367 | 1.436 | 0.879 | 10.564 | 53.304 | 3.396 |
| | | | 32 | 0.324 | 0.284 | 0.255 | 1.165 | 1.036 | 0.848 | 4.011 | 12.043 | 2.898 |
| | | | 64 | 0.229 | 0.222 | 0.182 | 0.889 | 0.862 | 0.807 | 1.615 | 3.192 | 2.506 |
| | | | 128 | 0.168 | 0.159 | 0.126 | 0.721 | 0.698 | 0.785 | 0.955 | 0.956 | 1.198 |
| | | 0.00100 | 16 | 0.396 | 0.349 | 0.346 | 1.246 | 1.172 | 0.887 | 3.456 | 10.451 | 3.072 |
| | | | 32 | 0.333 | 0.310 | 0.260 | 1.222 | 1.142 | 0.829 | 1.452 | 2.637 | 2.666 |
| | | | 64 | 0.234 | 0.228 | 0.181 | 0.889 | 0.863 | 0.794 | 1.002 | 1.008 | 1.074 |
| | | | 128 | 0.177 | 0.177 | 0.127 | 0.727 | 0.727 | 0.719 | 0.515 | 0.515 | 0.402 |
| | | 0.00400 | 16 | 0.396 | 0.375 | 0.348 | 1.290 | 1.171 | 0.875 | 1.623 | 2.379 | 2.375 |
| | | | 32 | 0.316 | 0.298 | 0.252 | 1.167 | 1.096 | 0.821 | 0.921 | 0.913 | 1.176 |
| | | | 64 | 0.215 | 0.215 | 0.170 | 0.844 | 0.842 | 0.754 | 0.526 | 0.516 | 0.456 |
| | | | 128 | 0.156 | 0.156 | 0.114 | 0.630 | 0.630 | 0.636 | 0.253 | 0.253 | 0.195 |
| | 1.0 | 0.00025 | 16 | 0.394 | 0.333 | 0.359 | 1.221 | 1.215 | 0.888 | 12.236 | 70.705 | 3.534 |
| | | | 32 | 0.333 | 0.271 | 0.259 | 1.212 | 0.891 | 0.833 | 4.763 | 15.630 | 3.084 |
| | | | 64 | 0.230 | 0.206 | 0.180 | 0.870 | 0.774 | 0.809 | 2.293 | 4.557 | 2.395 |
| | | | 128 | 0.165 | 0.158 | 0.126 | 0.709 | 0.698 | 0.771 | 1.218 | 1.328 | 1.430 |
| | | 0.00100 | 16 | 0.417 | 0.375 | 0.357 | 1.251 | 1.156 | 0.882 | 4.119 | 13.818 | 3.231 |
| | | | 32 | 0.339 | 0.345 | 0.259 | 1.232 | 1.212 | 0.833 | 1.951 | 3.929 | 2.663 |
| | | | 64 | 0.241 | 0.210 | 0.182 | 0.929 | 0.835 | 0.792 | 1.195 | 1.324 | 1.626 |
| | | | 128 | 0.164 | 0.164 | 0.121 | 0.671 | 0.671 | 0.731 | 0.645 | 0.645 | 0.500 |
| | | 0.00400 | 16 | 0.404 | 0.377 | 0.354 | 1.216 | 1.122 | 0.880 | 1.767 | 3.365 | 2.766 |
| | | | 32 | 0.341 | 0.312 | 0.259 | 1.270 | 1.135 | 0.810 | 1.066 | 1.115 | 1.661 |
| | | | 64 | 0.217 | 0.218 | 0.168 | 0.819 | 0.819 | 0.737 | 0.566 | 0.561 | 0.520 |
| | | | 128 | 0.158 | 0.158 | 0.112 | 0.586 | 0.586 | 0.558 | 0.329 | 0.329 | 0.238 |
| 0.5 | 0.5 | 0.00025 | 16 | 0.388 | 0.365 | 0.341 | 0.744 | 0.722 | 0.829 | 5.300 | 23.286 | 3.319 |
| | | | 32 | 0.318 | 0.347 | 0.273 | 0.726 | 0.771 | 0.783 | 2.172 | 5.417 | 2.738 |
| | | | 64 | 0.267 | 0.266 | 0.201 | 0.686 | 0.684 | 0.729 | 1.410 | 1.636 | 1.556 |
| | | | 128 | 0.192 | 0.192 | 0.151 | 0.583 | 0.584 | 0.633 | 0.778 | 0.767 | 0.661 |
| | | 0.00100 | 16 | 0.396 | 0.396 | 0.338 | 0.762 | 0.721 | 0.829 | 2.301 | 4.947 | 2.782 |
| | | | 32 | 0.340 | 0.333 | 0.269 | 0.770 | 0.750 | 0.757 | 1.289 | 1.439 | 1.595 |
| | | | 64 | 0.253 | 0.253 | 0.202 | 0.641 | 0.641 | 0.674 | 0.696 | 0.693 | 0.593 |
| | | | 128 | 0.188 | 0.188 | 0.143 | 0.540 | 0.540 | 0.545 | 0.364 | 0.364 | 0.275 |
| | | 0.00400 | 16 | 0.400 | 0.387 | 0.332 | 0.759 | 0.713 | 0.804 | 1.340 | 1.517 | 1.684 |
| | | | 32 | 0.327 | 0.326 | 0.256 | 0.732 | 0.730 | 0.736 | 0.716 | 0.708 | 0.648 |
| | | | 64 | 0.253 | 0.253 | 0.187 | 0.610 | 0.610 | 0.566 | 0.382 | 0.382 | 0.280 |
| | | | 128 | 0.159 | 0.159 | 0.117 | 0.426 | 0.426 | 0.382 | 0.199 | 0.199 | 0.140 |
| | 0.75 | 0.00025 | 16 | 0.424 | 0.404 | 0.367 | 0.813 | 0.702 | 0.842 | 8.076 | 33.166 | 3.446 |
| | | | 32 | 0.317 | 0.303 | 0.276 | 0.730 | 0.707 | 0.791 | 2.775 | 9.147 | 3.050 |
| | | | 64 | 0.270 | 0.281 | 0.203 | 0.696 | 0.711 | 0.696 | 1.980 | 2.779 | 1.888 |
| | | | 128 | 0.189 | 0.190 | 0.145 | 0.600 | 0.600 | 0.652 | 0.924 | 0.921 | 0.919 |
| | | 0.00100 | 16 | 0.428 | 0.438 | 0.362 | 0.798 | 0.784 | 0.847 | 3.179 | 7.665 | 2.932 |
| | | | 32 | 0.343 | 0.320 | 0.275 | 0.776 | 0.720 | 0.764 | 1.801 | 2.465 | 1.983 |
| | | | 64 | 0.261 | 0.256 | 0.199 | 0.670 | 0.665 | 0.691 | 0.887 | 0.876 | 1.037 |
| | | | 128 | 0.179 | 0.179 | 0.142 | 0.543 | 0.543 | 0.566 | 0.558 | 0.558 | 0.372 |
| | | 0.00400 | 16 | 0.427 | 0.429 | 0.358 | 0.810 | 0.773 | 0.820 | 1.701 | 2.175 | 1.951 |
| | | | 32 | 0.335 | 0.327 | 0.269 | 0.767 | 0.749 | 0.719 | 0.881 | 0.869 | 1.028 |
| | | | 64 | 0.238 | 0.238 | 0.184 | 0.607 | 0.607 | 0.628 | 0.468 | 0.468 | 0.366 |
| | | | 128 | 0.174 | 0.174 | 0.122 | 0.444 | 0.444 | 0.402 | 0.249 | 0.249 | 0.181 |
| | 1.0 | 0.00025 | 16 | 0.442 | 0.372 | 0.376 | 0.821 | 0.702 | 0.844 | 13.193 | 54.515 | 3.411 |
| | | | 32 | 0.330 | 0.310 | 0.274 | 0.756 | 0.679 | 0.782 | 4.378 | 13.848 | 3.052 |
| | | | 64 | 0.256 | 0.252 | 0.198 | 0.674 | 0.672 | 0.726 | 2.026 | 3.723 | 2.356 |
| | | | 128 | 0.187 | 0.188 | 0.141 | 0.597 | 0.601 | 0.655 | 1.134 | 1.182 | 1.215 |
| | | 0.00100 | 16 | 0.437 | 0.423 | 0.374 | 0.814 | 0.787 | 0.838 | 3.795 | 12.267 | 3.235 |
| | | | 32 | 0.352 | 0.330 | 0.280 | 0.795 | 0.759 | 0.747 | 1.781 | 3.567 | 2.677 |
| | | | 64 | 0.261 | 0.257 | 0.194 | 0.663 | 0.659 | 0.680 | 1.169 | 1.212 | 1.190 |
| | | | 128 | 0.173 | 0.173 | 0.135 | 0.535 | 0.535 | 0.562 | 0.608 | 0.608 | 0.446 |
| | | 0.00400 | 16 | 0.425 | 0.425 | 0.369 | 0.776 | 0.707 | 0.821 | 1.673 | 2.720 | 2.600 |
| | | | 32 | 0.339 | 0.336 | 0.269 | 0.770 | 0.760 | 0.707 | 1.092 | 1.124 | 1.298 |
| | | | 64 | 0.237 | 0.237 | 0.182 | 0.598 | 0.598 | 0.601 | 0.598 | 0.598 | 0.464 |
| | | | 128 | 0.168 | 0.168 | 0.120 | 0.462 | 0.462 | 0.415 | 0.291 | 0.291 | 0.221 |

**Table S4:** Simulation results for $\eta$ with $k = 4$.

| | | | | $\lambda$ | | | $\mu$ | | | $\theta$ | | |
|---|---|---|---|---|---|---|---|---|---|---|---|---|
| $\epsilon$ | $\rho$ | $\theta$ | $N$ | RMSE | RMSE$_c$ | CV | RMSE | RMSE$_c$ | CV | RMSE | RMSE$_c$ | CV |
| 0.1 | 0.5 | 0.00025 | 16 | 0.430 | 0.512 | 0.344 | 5.495 | 6.849 | 0.941 | 6.991 | 32.821 | 3.335 |
| | | | 32 | 0.366 | 0.323 | 0.244 | 4.541 | 3.933 | 0.882 | 2.783 | 7.115 | 2.795 |
| | | | 64 | 0.270 | 0.249 | 0.171 | 3.143 | 2.842 | 0.850 | 1.404 | 2.128 | 2.190 |
| | | | 128 | 0.184 | 0.183 | 0.115 | 1.842 | 1.820 | 0.850 | 0.773 | 0.761 | 0.806 |
| | | 0.00100 | 16 | 0.423 | 0.370 | 0.341 | 5.284 | 4.427 | 0.943 | 2.686 | 6.584 | 2.892 |
| | | | 32 | 0.346 | 0.315 | 0.242 | 4.377 | 4.001 | 0.884 | 1.510 | 2.033 | 2.039 |
| | | | 64 | 0.241 | 0.236 | 0.162 | 2.720 | 2.666 | 0.865 | 0.796 | 0.783 | 0.807 |
| | | | 128 | 0.179 | 0.179 | 0.108 | 1.662 | 1.662 | 0.849 | 0.436 | 0.436 | 0.342 |
| | | 0.00400 | 16 | 0.429 | 0.403 | 0.339 | 5.337 | 4.969 | 0.934 | 1.361 | 1.768 | 2.234 |
| | | | 32 | 0.323 | 0.314 | 0.233 | 4.016 | 3.905 | 0.878 | 0.743 | 0.717 | 0.917 |
| | | | 64 | 0.225 | 0.225 | 0.151 | 2.446 | 2.446 | 0.852 | 0.404 | 0.404 | 0.339 |
| | | | 128 | 0.152 | 0.152 | 0.098 | 1.372 | 1.372 | 0.801 | 0.213 | 0.213 | 0.163 |
| | 0.75 | 0.00025 | 16 | 0.435 | 0.383 | 0.353 | 5.335 | 5.607 | 0.918 | 9.857 | 43.764 | 3.473 |
| | | | 32 | 0.315 | 0.389 | 0.244 | 4.065 | 4.464 | 0.885 | 3.875 | 13.735 | 3.034 |
| | | | 64 | 0.274 | 0.273 | 0.171 | 3.152 | 3.132 | 0.848 | 1.813 | 3.475 | 2.428 |
| | | | 128 | 0.194 | 0.186 | 0.116 | 1.876 | 1.818 | 0.850 | 1.075 | 1.124 | 1.365 |
| | | 0.00100 | 16 | 0.382 | 0.366 | 0.341 | 4.670 | 4.746 | 0.932 | 3.270 | 13.457 | 3.316 |
| | | | 32 | 0.360 | 0.342 | 0.249 | 4.588 | 4.079 | 0.867 | 1.655 | 2.956 | 2.550 |
| | | | 64 | 0.247 | 0.243 | 0.167 | 2.877 | 2.830 | 0.851 | 0.948 | 0.947 | 1.037 |
| | | | 128 | 0.176 | 0.176 | 0.110 | 1.646 | 1.646 | 0.842 | 0.562 | 0.562 | 0.430 |
| | | 0.00400 | 16 | 0.419 | 0.361 | 0.345 | 5.078 | 3.774 | 0.920 | 1.685 | 2.953 | 2.649 |
| | | | 32 | 0.329 | 0.311 | 0.240 | 4.187 | 3.916 | 0.864 | 0.994 | 1.001 | 1.271 |
| | | | 64 | 0.244 | 0.242 | 0.160 | 2.700 | 2.669 | 0.838 | 0.528 | 0.523 | 0.446 |
| | | | 128 | 0.153 | 0.153 | 0.101 | 1.320 | 1.320 | 0.832 | 0.285 | 0.285 | 0.208 |
| | 1.0 | 0.00025 | 16 | 0.422 | 0.327 | 0.361 | 5.070 | 4.165 | 0.906 | 13.741 | 55.488 | 3.499 |
| | | | 32 | 0.331 | 0.249 | 0.247 | 4.249 | 2.939 | 0.872 | 4.855 | 19.739 | 3.195 |
| | | | 64 | 0.257 | 0.230 | 0.172 | 3.002 | 2.834 | 0.854 | 2.088 | 4.899 | 2.636 |
| | | | 128 | 0.194 | 0.175 | 0.116 | 1.823 | 1.643 | 0.858 | 1.187 | 1.355 | 1.695 |
| | | 0.00100 | 16 | 0.408 | 0.376 | 0.354 | 4.827 | 3.890 | 0.912 | 4.314 | 17.080 | 3.351 |
| | | | 32 | 0.338 | 0.324 | 0.250 | 4.396 | 3.910 | 0.857 | 1.811 | 4.440 | 2.915 |
| | | | 64 | 0.265 | 0.243 | 0.170 | 3.000 | 2.717 | 0.856 | 1.047 | 1.130 | 1.973 |
| | | | 128 | 0.181 | 0.180 | 0.112 | 1.625 | 1.604 | 0.856 | 0.663 | 0.653 | 0.629 |
| | | 0.00400 | 16 | 0.410 | 0.365 | 0.351 | 4.799 | 3.912 | 0.907 | 2.123 | 4.234 | 2.710 |
| | | | 32 | 0.351 | 0.335 | 0.245 | 4.354 | 4.109 | 0.863 | 1.084 | 1.162 | 1.874 |
| | | | 64 | 0.235 | 0.236 | 0.160 | 2.562 | 2.565 | 0.851 | 0.630 | 0.627 | 0.539 |
| | | | 128 | 0.168 | 0.168 | 0.103 | 1.379 | 1.379 | 0.784 | 0.316 | 0.316 | 0.255 |
| 0.3 | 0.5 | 0.00025 | 16 | 0.401 | 0.375 | 0.352 | 1.359 | 0.876 | 0.925 | 5.767 | 21.341 | 3.321 |
| | | | 32 | 0.340 | 0.312 | 0.263 | 1.254 | 1.068 | 0.845 | 2.400 | 6.125 | 2.809 |
| | | | 64 | 0.245 | 0.230 | 0.183 | 0.918 | 0.855 | 0.811 | 1.399 | 1.810 | 1.884 |
| | | | 128 | 0.179 | 0.179 | 0.132 | 0.726 | 0.728 | 0.755 | 0.789 | 0.783 | 0.665 |
| | | 0.00100 | 16 | 0.392 | 0.382 | 0.347 | 1.295 | 1.197 | 0.932 | 2.587 | 5.667 | 2.762 |
| | | | 32 | 0.319 | 0.304 | 0.250 | 1.153 | 1.051 | 0.853 | 1.434 | 1.763 | 1.849 |
| | | | 64 | 0.216 | 0.212 | 0.178 | 0.835 | 0.819 | 0.804 | 0.768 | 0.755 | 0.785 |
| | | | 128 | 0.160 | 0.160 | 0.125 | 0.690 | 0.690 | 0.737 | 0.394 | 0.394 | 0.304 |
| | | 0.00400 | 16 | 0.397 | 0.395 | 0.345 | 1.287 | 1.225 | 0.923 | 1.283 | 1.495 | 1.932 |
| | | | 32 | 0.315 | 0.309 | 0.249 | 1.155 | 1.133 | 0.826 | 0.739 | 0.729 | 0.750 |
| | | | 64 | 0.215 | 0.215 | 0.170 | 0.829 | 0.829 | 0.733 | 0.395 | 0.395 | 0.311 |
| | | | 128 | 0.161 | 0.161 | 0.111 | 0.655 | 0.655 | 0.598 | 0.210 | 0.210 | 0.152 |
| | 0.75 | 0.00025 | 16 | 0.426 | 0.460 | 0.365 | 1.440 | 1.483 | 0.898 | 10.554 | 53.770 | 3.403 |
| | | | 32 | 0.324 | 0.285 | 0.255 | 1.168 | 1.038 | 0.846 | 4.024 | 12.031 | 2.895 |
| | | | 64 | 0.228 | 0.223 | 0.182 | 0.887 | 0.867 | 0.806 | 1.619 | 3.219 | 2.485 |
| | | | 128 | 0.169 | 0.158 | 0.126 | 0.722 | 0.698 | 0.784 | 0.955 | 0.955 | 1.203 |
| | | 0.00100 | 16 | 0.411 | 0.359 | 0.357 | 1.307 | 1.214 | 0.905 | 3.469 | 10.491 | 3.085 |
| | | | 32 | 0.333 | 0.310 | 0.260 | 1.223 | 1.141 | 0.832 | 1.455 | 2.651 | 2.653 |
| | | | 64 | 0.233 | 0.227 | 0.180 | 0.884 | 0.859 | 0.790 | 0.999 | 1.006 | 1.073 |
| | | | 128 | 0.177 | 0.177 | 0.127 | 0.728 | 0.728 | 0.719 | 0.515 | 0.515 | 0.402 |
| | | 0.00400 | 16 | 0.411 | 0.385 | 0.358 | 1.349 | 1.211 | 0.890 | 1.617 | 2.368 | 2.359 |
| | | | 32 | 0.315 | 0.298 | 0.252 | 1.166 | 1.093 | 0.823 | 0.921 | 0.912 | 1.172 |
| | | | 64 | 0.215 | 0.215 | 0.170 | 0.843 | 0.841 | 0.753 | 0.524 | 0.514 | 0.459 |
| | | | 128 | 0.156 | 0.156 | 0.114 | 0.630 | 0.630 | 0.635 | 0.254 | 0.254 | 0.196 |
| | 1.0 | 0.00025 | 16 | 0.399 | 0.343 | 0.363 | 1.235 | 1.232 | 0.897 | 12.221 | 71.187 | 3.510 |
| | | | 32 | 0.332 | 0.270 | 0.258 | 1.211 | 0.889 | 0.832 | 4.760 | 15.729 | 3.070 |
| | | | 64 | 0.229 | 0.207 | 0.180 | 0.868 | 0.775 | 0.811 | 2.286 | 4.537 | 2.413 |
| | | | 128 | 0.165 | 0.158 | 0.126 | 0.709 | 0.698 | 0.771 | 1.216 | 1.327 | 1.422 |
| | | 0.00100 | 16 | 0.425 | 0.374 | 0.362 | 1.281 | 1.151 | 0.891 | 4.134 | 13.903 | 3.234 |
| | | | 32 | 0.339 | 0.344 | 0.260 | 1.232 | 1.210 | 0.832 | 1.950 | 3.925 | 2.674 |
| | | | 64 | 0.242 | 0.210 | 0.182 | 0.933 | 0.835 | 0.793 | 1.193 | 1.321 | 1.623 |
| | | | 128 | 0.165 | 0.165 | 0.121 | 0.670 | 0.670 | 0.730 | 0.644 | 0.644 | 0.500 |
| | | 0.00400 | 16 | 0.411 | 0.383 | 0.359 | 1.243 | 1.142 | 0.886 | 1.764 | 3.361 | 2.774 |
| | | | 32 | 0.341 | 0.310 | 0.259 | 1.268 | 1.130 | 0.810 | 1.068 | 1.118 | 1.662 |
| | | | 64 | 0.218 | 0.218 | 0.169 | 0.820 | 0.821 | 0.737 | 0.565 | 0.560 | 0.522 |
| | | | 128 | 0.158 | 0.158 | 0.112 | 0.584 | 0.584 | 0.556 | 0.328 | 0.328 | 0.237 |
| 0.5 | 0.5 | 0.00025 | 16 | 0.423 | 0.400 | 0.380 | 0.840 | 0.814 | 0.882 | 5.200 | 22.726 | 3.330 |
| | | | 32 | 0.328 | 0.361 | 0.282 | 0.748 | 0.800 | 0.797 | 2.169 | 5.382 | 2.760 |
| | | | 64 | 0.268 | 0.266 | 0.202 | 0.688 | 0.684 | 0.730 | 1.406 | 1.631 | 1.543 |
| | | | 128 | 0.192 | 0.191 | 0.150 | 0.582 | 0.583 | 0.631 | 0.777 | 0.767 | 0.661 |
| | | 0.00100 | 16 | 0.434 | 0.419 | 0.375 | 0.860 | 0.786 | 0.883 | 2.260 | 4.838 | 2.789 |
| | | | 32 | 0.351 | 0.345 | 0.278 | 0.796 | 0.776 | 0.771 | 1.288 | 1.437 | 1.610 |
| | | | 64 | 0.254 | 0.254 | 0.203 | 0.643 | 0.643 | 0.676 | 0.695 | 0.692 | 0.593 |
| | | | 128 | 0.188 | 0.188 | 0.143 | 0.540 | 0.540 | 0.544 | 0.364 | 0.364 | 0.275 |
| | | 0.00400 | 16 | 0.437 | 0.412 | 0.370 | 0.859 | 0.789 | 0.855 | 1.323 | 1.492 | 1.691 |
| | | | 32 | 0.336 | 0.334 | 0.263 | 0.752 | 0.749 | 0.747 | 0.716 | 0.708 | 0.650 |
| | | | 64 | 0.253 | 0.253 | 0.187 | 0.610 | 0.610 | 0.566 | 0.382 | 0.382 | 0.280 |
| | | | 128 | 0.159 | 0.159 | 0.117 | 0.425 | 0.425 | 0.381 | 0.199 | 0.199 | 0.140 |
| | 0.75 | 0.00025 | 16 | 0.449 | 0.416 | 0.384 | 0.872 | 0.738 | 0.866 | 7.916 | 33.048 | 3.431 |
| | | | 32 | 0.318 | 0.306 | 0.277 | 0.733 | 0.715 | 0.794 | 2.781 | 9.187 | 3.059 |
| | | | 64 | 0.271 | 0.281 | 0.204 | 0.697 | 0.713 | 0.699 | 1.969 | 2.763 | 1.871 |
| | | | 128 | 0.189 | 0.190 | 0.145 | 0.601 | 0.599 | 0.652 | 0.924 | 0.921 | 0.918 |
| | | 0.00100 | 16 | 0.450 | 0.460 | 0.377 | 0.850 | 0.837 | 0.868 | 3.167 | 7.643 | 2.947 |
| | | | 32 | 0.345 | 0.321 | 0.276 | 0.781 | 0.723 | 0.767 | 1.810 | 2.480 | 1.981 |
| | | | 64 | 0.261 | 0.257 | 0.199 | 0.672 | 0.666 | 0.692 | 0.889 | 0.878 | 1.034 |
| | | | 128 | 0.178 | 0.178 | 0.141 | 0.542 | 0.542 | 0.566 | 0.557 | 0.557 | 0.372 |
| | | 0.00400 | 16 | 0.450 | 0.446 | 0.377 | 0.863 | 0.814 | 0.846 | 1.699 | 2.171 | 1.960 |
| | | | 32 | 0.337 | 0.329 | 0.271 | 0.770 | 0.753 | 0.721 | 0.880 | 0.868 | 1.026 |
| | | | 64 | 0.238 | 0.238 | 0.184 | 0.607 | 0.607 | 0.628 | 0.468 | 0.468 | 0.365 |
| | | | 128 | 0.174 | 0.174 | 0.122 | 0.443 | 0.443 | 0.401 | 0.249 | 0.249 | 0.181 |
| | 1.0 | 0.00025 | 16 | 0.453 | 0.374 | 0.384 | 0.847 | 0.710 | 0.854 | 13.055 | 54.392 | 3.399 |
| | | | 32 | 0.330 | 0.310 | 0.274 | 0.755 | 0.679 | 0.778 | 4.377 | 13.791 | 3.057 |
| | | | 64 | 0.256 | 0.253 | 0.198 | 0.675 | 0.675 | 0.728 | 2.031 | 3.731 | 2.364 |
| | | | 128 | 0.187 | 0.188 | 0.141 | 0.597 | 0.601 | 0.655 | 1.135 | 1.184 | 1.210 |
| | | 0.00100 | 16 | 0.450 | 0.429 | 0.384 | 0.845 | 0.801 | 0.852 | 3.771 | 12.073 | 3.240 |
| | | | 32 | 0.353 | 0.330 | 0.280 | 0.799 | 0.761 | 0.746 | 1.782 | 3.570 | 2.691 |
| | | | 64 | 0.262 | 0.258 | 0.194 | 0.664 | 0.661 | 0.681 | 1.171 | 1.214 | 1.195 |
| | | | 128 | 0.173 | 0.173 | 0.135 | 0.535 | 0.535 | 0.561 | 0.608 | 0.608 | 0.446 |
| | | 0.00400 | 16 | 0.435 | 0.428 | 0.376 | 0.799 | 0.715 | 0.832 | 1.667 | 2.711 | 2.584 |
| | | | 32 | 0.340 | 0.337 | 0.268 | 0.771 | 0.763 | 0.705 | 1.088 | 1.119 | 1.298 |
| | | | 64 | 0.237 | 0.237 | 0.183 | 0.597 | 0.597 | 0.600 | 0.597 | 0.597 | 0.463 |
| | | | 128 | 0.167 | 0.167 | 0.120 | 0.461 | 0.461 | 0.415 | 0.291 | 0.291 | 0.221 |

**Table S5:** Simulation results for $\zeta$ with $k = 1$.

| | | | | $\lambda$ | | | $\mu$ | | | $\theta$ | | |
|---|---|---|---|---|---|---|---|---|---|---|---|---|
| $\epsilon$ | $\rho$ | $\theta$ | $N$ | RMSE | $RMSE_c$ | CV | RMSE | $RMSE_c$ | CV | RMSE | $RMSE_c$ | CV |
| 0.1 | 0.5 | 0.00025 | 16 | 0.290 | 0.323 | 0.265 | 3.070 | 2.372 | 0.849 | 5.783 | 21.273 | 3.345 |
| | | | 32 | 0.245 | 0.269 | 0.201 | 2.945 | 3.017 | 0.838 | 2.252 | 5.925 | 2.781 |
| | | | 64 | 0.202 | 0.200 | 0.147 | 2.294 | 2.227 | 0.854 | 1.364 | 1.640 | 1.722 |
| | | | 128 | 0.152 | 0.148 | 0.105 | 1.533 | 1.509 | 0.854 | 0.681 | 0.669 | 0.659 |
| | | 0.00100 | 16 | 0.301 | 0.297 | 0.261 | 2.958 | 2.722 | 0.854 | 2.842 | 5.921 | 2.651 |
| | | | 32 | 0.246 | 0.234 | 0.198 | 2.877 | 2.663 | 0.845 | 1.395 | 1.674 | 1.772 |
| | | | 64 | 0.208 | 0.205 | 0.147 | 2.380 | 2.355 | 0.839 | 0.723 | 0.718 | 0.636 |
| | | | 128 | 0.144 | 0.144 | 0.100 | 1.408 | 1.408 | 0.844 | 0.374 | 0.374 | 0.283 |
| | | 0.00400 | 16 | 0.297 | 0.294 | 0.263 | 3.152 | 3.027 | 0.839 | 1.380 | 1.550 | 1.639 |
| | | | 32 | 0.241 | 0.235 | 0.196 | 2.949 | 2.888 | 0.830 | 0.743 | 0.738 | 0.638 |
| | | | 64 | 0.180 | 0.180 | 0.136 | 2.039 | 2.039 | 0.804 | 0.381 | 0.381 | 0.276 |
| | | | 128 | 0.142 | 0.142 | 0.090 | 1.276 | 1.276 | 0.717 | 0.195 | 0.195 | 0.144 |
| | 0.75 | 0.00025 | 16 | 0.351 | 0.513 | 0.304 | 4.100 | 5.571 | 0.845 | 8.693 | 43.220 | 3.445 |
| | | | 32 | 0.323 | 0.347 | 0.236 | 4.182 | 4.469 | 0.842 | 3.306 | 11.013 | 3.012 |
| | | | 64 | 0.239 | 0.232 | 0.167 | 2.777 | 2.627 | 0.858 | 1.674 | 2.644 | 2.201 |
| | | | 128 | 0.188 | 0.186 | 0.113 | 1.760 | 1.713 | 0.855 | 0.902 | 0.891 | 1.125 |
| | | 0.00100 | 16 | 0.348 | 0.305 | 0.305 | 4.299 | 3.335 | 0.835 | 2.837 | 10.170 | 3.299 |
| | | | 32 | 0.305 | 0.280 | 0.232 | 3.862 | 3.523 | 0.857 | 1.615 | 2.473 | 2.324 |
| | | | 64 | 0.259 | 0.242 | 0.164 | 2.866 | 2.679 | 0.856 | 0.885 | 0.868 | 1.182 |
| | | | 128 | 0.172 | 0.172 | 0.109 | 1.633 | 1.637 | 0.827 | 0.493 | 0.488 | 0.420 |
| | | 0.00400 | 16 | 0.360 | 0.342 | 0.304 | 4.396 | 4.061 | 0.829 | 1.658 | 2.646 | 2.518 |
| | | | 32 | 0.321 | 0.306 | 0.227 | 3.876 | 3.667 | 0.855 | 0.918 | 0.908 | 1.164 |
| | | | 64 | 0.226 | 0.226 | 0.155 | 2.500 | 2.500 | 0.824 | 0.517 | 0.517 | 0.404 |
| | | | 128 | 0.155 | 0.155 | 0.098 | 1.248 | 1.248 | 0.778 | 0.262 | 0.262 | 0.199 |
| | 1.0 | 0.00025 | 16 | 0.381 | – | 0.331 | 4.655 | – | 0.856 | 12.316 | – | 3.570 |
| | | | 32 | 0.322 | 0.255 | 0.245 | 4.199 | 3.084 | 0.860 | 4.756 | 17.977 | 3.172 |
| | | | 64 | 0.253 | 0.203 | 0.172 | 3.053 | 2.448 | 0.847 | 2.057 | 4.497 | 2.588 |
| | | | 128 | 0.184 | 0.181 | 0.113 | 1.691 | 1.647 | 0.859 | 1.242 | 1.352 | 1.402 |
| | | 0.00100 | 16 | 0.377 | 0.369 | 0.329 | 4.353 | 2.507 | 0.865 | 4.094 | 11.857 | 3.304 |
| | | | 32 | 0.309 | 0.287 | 0.241 | 3.960 | 3.366 | 0.869 | 2.159 | 4.046 | 2.531 |
| | | | 64 | 0.247 | 0.220 | 0.166 | 2.761 | 2.475 | 0.852 | 1.122 | 1.194 | 1.543 |
| | | | 128 | 0.189 | 0.188 | 0.111 | 1.670 | 1.669 | 0.826 | 0.649 | 0.646 | 0.581 |
| | | 0.00400 | 16 | 0.371 | 0.350 | 0.325 | 4.240 | 3.368 | 0.859 | 1.743 | 3.414 | 2.855 |
| | | | 32 | 0.303 | 0.289 | 0.238 | 3.851 | 3.524 | 0.863 | 1.099 | 1.180 | 1.782 |
| | | | 64 | 0.220 | 0.220 | 0.153 | 2.224 | 2.218 | 0.862 | 0.627 | 0.623 | 0.589 |
| | | | 128 | 0.164 | 0.164 | 0.099 | 1.233 | 1.233 | 0.712 | 0.339 | 0.339 | 0.254 |
| 0.3 | 0.5 | 0.00025 | 16 | 0.341 | 0.307 | 0.268 | 0.795 | 0.770 | 0.829 | 5.522 | 20.424 | 3.237 |
| | | | 32 | 0.250 | 0.256 | 0.206 | 0.803 | 0.757 | 0.815 | 2.601 | 5.199 | 2.464 |
| | | | 64 | 0.194 | 0.191 | 0.159 | 0.741 | 0.724 | 0.793 | 1.268 | 1.440 | 1.603 |
| | | | 128 | 0.146 | 0.147 | 0.116 | 0.654 | 0.654 | 0.766 | 0.716 | 0.714 | 0.529 |
| | | 0.00100 | 16 | 0.337 | 0.333 | 0.269 | 0.818 | 0.790 | 0.832 | 2.229 | 5.337 | 2.876 |
| | | | 32 | 0.251 | 0.251 | 0.205 | 0.823 | 0.805 | 0.800 | 1.402 | 1.528 | 1.349 |
| | | | 64 | 0.208 | 0.208 | 0.156 | 0.766 | 0.764 | 0.768 | 0.697 | 0.688 | 0.598 |
| | | | 128 | 0.157 | 0.157 | 0.115 | 0.665 | 0.665 | 0.674 | 0.382 | 0.382 | 0.254 |
| | | 0.00400 | 16 | 0.340 | 0.340 | 0.265 | 0.821 | 0.789 | 0.815 | 1.349 | 1.556 | 1.727 |
| | | | 32 | 0.253 | 0.248 | 0.202 | 0.828 | 0.808 | 0.764 | 0.773 | 0.769 | 0.572 |
| | | | 64 | 0.187 | 0.187 | 0.148 | 0.729 | 0.729 | 0.663 | 0.375 | 0.375 | 0.261 |
| | | | 128 | 0.134 | 0.134 | 0.097 | 0.514 | 0.514 | 0.446 | 0.191 | 0.191 | 0.133 |
| | 0.75 | 0.00025 | 16 | 0.354 | 0.313 | 0.308 | 1.056 | 0.883 | 0.813 | 7.524 | 28.818 | 3.458 |
| | | | 32 | 0.303 | 0.269 | 0.242 | 1.077 | 0.949 | 0.818 | 2.849 | 8.335 | 2.990 |
| | | | 64 | 0.244 | 0.243 | 0.180 | 0.921 | 0.936 | 0.798 | 1.715 | 2.680 | 2.133 |
| | | | 128 | 0.177 | 0.177 | 0.129 | 0.718 | 0.719 | 0.762 | 0.941 | 0.938 | 0.971 |
| | | 0.00100 | 16 | 0.362 | 0.391 | 0.307 | 1.030 | 1.063 | 0.810 | 3.189 | 8.197 | 2.983 |
| | | | 32 | 0.277 | 0.271 | 0.241 | 1.012 | 1.005 | 0.824 | 1.570 | 2.289 | 2.243 |
| | | | 64 | 0.235 | 0.231 | 0.176 | 0.887 | 0.873 | 0.784 | 0.893 | 0.889 | 0.854 |
| | | | 128 | 0.163 | 0.163 | 0.121 | 0.685 | 0.685 | 0.702 | 0.473 | 0.473 | 0.351 |
| | | 0.00400 | 16 | 0.345 | 0.351 | 0.304 | 1.037 | 1.043 | 0.792 | 1.728 | 2.267 | 2.018 |
| | | | 32 | 0.282 | 0.283 | 0.231 | 0.985 | 0.978 | 0.803 | 0.891 | 0.885 | 0.890 |
| | | | 64 | 0.218 | 0.218 | 0.163 | 0.810 | 0.810 | 0.705 | 0.480 | 0.480 | 0.359 |
| | | | 128 | 0.147 | 0.147 | 0.106 | 0.562 | 0.562 | 0.525 | 0.254 | 0.254 | 0.185 |
| | 1.0 | 0.00025 | 16 | 0.384 | 0.345 | 0.333 | 1.113 | 0.904 | 0.840 | 11.582 | 44.864 | 3.571 |
| | | | 32 | 0.312 | 0.305 | 0.255 | 1.144 | 0.761 | 0.820 | 3.310 | 10.068 | 3.301 |
| | | | 64 | 0.238 | 0.257 | 0.182 | 0.906 | 0.974 | 0.796 | 1.921 | 3.933 | 2.532 |
| | | | 128 | 0.170 | 0.166 | 0.125 | 0.694 | 0.685 | 0.769 | 1.207 | 1.293 | 1.334 |
| | | 0.00100 | 16 | 0.371 | 0.346 | 0.333 | 1.121 | 1.190 | 0.832 | 4.332 | 14.547 | 3.212 |
| | | | 32 | 0.313 | 0.306 | 0.252 | 1.142 | 1.068 | 0.809 | 2.252 | 4.302 | 2.514 |
| | | | 64 | 0.222 | 0.217 | 0.175 | 0.860 | 0.845 | 0.752 | 1.110 | 1.155 | 1.350 |
| | | | 128 | 0.171 | 0.171 | 0.120 | 0.670 | 0.670 | 0.662 | 0.609 | 0.609 | 0.498 |
| | | 0.00400 | 16 | 0.383 | 0.380 | 0.329 | 1.173 | 1.115 | 0.809 | 1.657 | 3.101 | 2.836 |
| | | | 32 | 0.300 | 0.292 | 0.243 | 1.075 | 1.019 | 0.795 | 1.113 | 1.155 | 1.365 |
| | | | 64 | 0.221 | 0.221 | 0.162 | 0.807 | 0.807 | 0.663 | 0.601 | 0.601 | 0.465 |
| | | | 128 | 0.150 | 0.150 | 0.105 | 0.534 | 0.534 | 0.492 | 0.311 | 0.311 | 0.236 |
| 0.5 | 0.5 | 0.00025 | 16 | 0.397 | 0.381 | 0.277 | 0.656 | 0.644 | 0.790 | 5.623 | 18.664 | 3.104 |
| | | | 32 | 0.290 | 0.274 | 0.218 | 0.638 | 0.624 | 0.741 | 2.204 | 4.805 | 2.546 |
| | | | 64 | 0.237 | 0.240 | 0.172 | 0.617 | 0.617 | 0.681 | 1.285 | 1.386 | 1.290 |
| | | | 128 | 0.193 | 0.193 | 0.136 | 0.563 | 0.564 | 0.590 | 0.656 | 0.653 | 0.477 |
| | | 0.00100 | 16 | 0.405 | 0.422 | 0.274 | 0.668 | 0.676 | 0.789 | 2.475 | 4.573 | 2.472 |
| | | | 32 | 0.315 | 0.312 | 0.214 | 0.647 | 0.641 | 0.730 | 1.269 | 1.362 | 1.362 |
| | | | 64 | 0.219 | 0.219 | 0.172 | 0.563 | 0.563 | 0.611 | 0.683 | 0.677 | 0.514 |
| | | | 128 | 0.166 | 0.166 | 0.126 | 0.488 | 0.488 | 0.480 | 0.356 | 0.356 | 0.231 |
| | | 0.00400 | 16 | 0.398 | 0.401 | 0.271 | 0.655 | 0.655 | 0.754 | 1.459 | 1.527 | 1.111 |
| | | | 32 | 0.276 | 0.277 | 0.208 | 0.578 | 0.578 | 0.648 | 0.715 | 0.712 | 0.462 |
| | | | 64 | 0.208 | 0.208 | 0.152 | 0.500 | 0.500 | 0.487 | 0.336 | 0.336 | 0.233 |
| | | | 128 | 0.130 | 0.130 | 0.096 | 0.314 | 0.314 | 0.256 | 0.170 | 0.170 | 0.121 |
| | 0.75 | 0.00025 | 16 | 0.382 | 0.379 | 0.314 | 0.664 | 0.609 | 0.779 | 6.679 | 24.944 | 3.419 |
| | | | 32 | 0.314 | 0.297 | 0.251 | 0.691 | 0.653 | 0.764 | 2.827 | 8.437 | 2.878 |
| | | | 64 | 0.248 | 0.247 | 0.200 | 0.644 | 0.639 | 0.691 | 1.586 | 2.137 | 1.913 |
| | | | 128 | 0.186 | 0.186 | 0.144 | 0.583 | 0.584 | 0.627 | 0.868 | 0.854 | 0.934 |
| | | 0.00100 | 16 | 0.382 | 0.396 | 0.311 | 0.657 | 0.640 | 0.775 | 3.061 | 6.654 | 2.781 |
| | | | 32 | 0.306 | 0.294 | 0.250 | 0.685 | 0.657 | 0.743 | 1.584 | 2.028 | 1.915 |
| | | | 64 | 0.235 | 0.235 | 0.190 | 0.605 | 0.603 | 0.664 | 0.900 | 0.897 | 0.720 |
| | | | 128 | 0.175 | 0.175 | 0.136 | 0.513 | 0.513 | 0.513 | 0.458 | 0.458 | 0.327 |
| | | 0.00400 | 16 | 0.370 | 0.362 | 0.308 | 0.634 | 0.616 | 0.734 | 1.533 | 1.836 | 1.830 |
| | | | 32 | 0.298 | 0.298 | 0.246 | 0.665 | 0.664 | 0.677 | 0.848 | 0.837 | 0.850 |
| | | | 64 | 0.230 | 0.230 | 0.172 | 0.561 | 0.561 | 0.537 | 0.455 | 0.455 | 0.325 |
| | | | 128 | 0.152 | 0.152 | 0.110 | 0.381 | 0.381 | 0.329 | 0.230 | 0.230 | 0.171 |
| | 1.0 | 0.00025 | 16 | 0.386 | 0.424 | 0.338 | 0.680 | 0.597 | 0.798 | 10.562 | 33.347 | 3.491 |
| | | | 32 | 0.320 | 0.317 | 0.268 | 0.720 | 0.762 | 0.753 | 3.772 | 13.819 | 3.140 |
| | | | 64 | 0.250 | 0.260 | 0.194 | 0.656 | 0.645 | 0.720 | 1.893 | 3.360 | 2.358 |
| | | | 128 | 0.181 | 0.180 | 0.139 | 0.577 | 0.569 | 0.618 | 1.139 | 1.176 | 1.117 |
| | | 0.00100 | 16 | 0.401 | 0.418 | 0.337 | 0.706 | 0.621 | 0.787 | 3.437 | 9.584 | 3.241 |
| | | | 32 | 0.325 | 0.325 | 0.262 | 0.725 | 0.700 | 0.757 | 1.767 | 2.843 | 2.429 |
| | | | 64 | 0.251 | 0.243 | 0.190 | 0.650 | 0.633 | 0.662 | 1.086 | 1.116 | 1.217 |
| | | | 128 | 0.175 | 0.175 | 0.129 | 0.498 | 0.498 | 0.481 | 0.553 | 0.553 | 0.416 |
| | | 0.00400 | 16 | 0.378 | 0.368 | 0.332 | 0.679 | 0.649 | 0.729 | 2.007 | 2.941 | 2.256 |
| | | | 32 | 0.314 | 0.315 | 0.251 | 0.696 | 0.686 | 0.691 | 1.087 | 1.113 | 1.195 |
| | | | 64 | 0.223 | 0.223 | 0.172 | 0.557 | 0.557 | 0.533 | 0.567 | 0.567 | 0.433 |
| | | | 128 | 0.161 | 0.161 | 0.110 | 0.365 | 0.365 | 0.312 | 0.292 | 0.292 | 0.209 |

**Table S6:** Simulation results for $\zeta$ with $k = 2$.

| | | | | $\lambda$ | | | $\mu$ | | | $\theta$ | | |
|---|---|---|---|---|---|---|---|---|---|---|---|---|
| $\epsilon$ | $\rho$ | $\theta$ | $N$ | RMSE | $RMSE_c$ | CV | RMSE | $RMSE_c$ | CV | RMSE | $RMSE_c$ | CV |
| 0.1 | 0.5 | 0.00025 | 16 | 0.374 | 0.342 | 0.319 | 4.891 | 3.631 | 0.896 | 4.951 | 17.919 | 3.368 |
| | | | 32 | 0.330 | 0.363 | 0.241 | 4.252 | 4.428 | 0.878 | 2.018 | 5.119 | 2.787 |
| | | | 64 | 0.251 | 0.248 | 0.165 | 2.845 | 2.743 | 0.864 | 1.283 | 1.506 | 1.723 |
| | | | 128 | 0.179 | 0.176 | 0.112 | 1.753 | 1.723 | 0.854 | 0.653 | 0.640 | 0.661 |
| | | 0.00100 | 16 | 0.369 | 0.341 | 0.311 | 4.640 | 4.177 | 0.905 | 2.438 | 4.937 | 2.692 |
| | | | 32 | 0.325 | 0.297 | 0.234 | 4.049 | 3.687 | 0.884 | 1.277 | 1.482 | 1.784 |
| | | | 64 | 0.257 | 0.253 | 0.164 | 2.921 | 2.885 | 0.852 | 0.683 | 0.677 | 0.645 |
| | | | 128 | 0.167 | 0.167 | 0.106 | 1.564 | 1.564 | 0.843 | 0.363 | 0.363 | 0.288 |
| | | 0.00400 | 16 | 0.386 | 0.370 | 0.313 | 4.931 | 4.682 | 0.887 | 1.206 | 1.305 | 1.678 |
| | | | 32 | 0.318 | 0.308 | 0.227 | 4.002 | 3.901 | 0.869 | 0.686 | 0.680 | 0.655 |
| | | | 64 | 0.209 | 0.209 | 0.145 | 2.294 | 2.294 | 0.818 | 0.362 | 0.362 | 0.286 |
| | | | 128 | 0.158 | 0.158 | 0.092 | 1.299 | 1.299 | 0.734 | 0.194 | 0.194 | 0.150 |
| | 0.75 | 0.00025 | 16 | 0.405 | 0.702 | 0.340 | 4.943 | 7.842 | 0.911 | 8.467 | 40.770 | 3.464 |
| | | | 32 | 0.344 | 0.361 | 0.247 | 4.448 | 4.638 | 0.866 | 3.301 | 11.080 | 3.010 |
| | | | 64 | 0.240 | 0.232 | 0.167 | 2.778 | 2.620 | 0.861 | 1.673 | 2.639 | 2.197 |
| | | | 128 | 0.188 | 0.186 | 0.113 | 1.759 | 1.712 | 0.855 | 0.899 | 0.888 | 1.137 |
| | | 0.00100 | 16 | 0.410 | 0.326 | 0.344 | 5.237 | 3.738 | 0.900 | 2.794 | 10.082 | 3.350 |
| | | | 32 | 0.320 | 0.289 | 0.241 | 4.059 | 3.657 | 0.875 | 1.613 | 2.471 | 2.315 |
| | | | 64 | 0.259 | 0.241 | 0.165 | 2.865 | 2.669 | 0.860 | 0.886 | 0.869 | 1.181 |
| | | | 128 | 0.171 | 0.171 | 0.109 | 1.630 | 1.634 | 0.826 | 0.493 | 0.488 | 0.419 |
| | | 0.00400 | 16 | 0.426 | 0.395 | 0.341 | 5.355 | 4.841 | 0.893 | 1.632 | 2.589 | 2.527 |
| | | | 32 | 0.335 | 0.318 | 0.234 | 4.056 | 3.814 | 0.871 | 0.914 | 0.903 | 1.164 |
| | | | 64 | 0.227 | 0.227 | 0.155 | 2.509 | 2.509 | 0.827 | 0.517 | 0.517 | 0.405 |
| | | | 128 | 0.155 | 0.155 | 0.097 | 1.244 | 1.244 | 0.776 | 0.263 | 0.263 | 0.199 |
| | 1.0 | 0.00025 | 16 | 0.420 | – | 0.356 | 5.163 | – | 0.898 | 12.254 | – | 3.618 |
| | | | 32 | 0.328 | 0.260 | 0.249 | 4.279 | 3.171 | 0.867 | 4.761 | 18.011 | 3.164 |
| | | | 64 | 0.253 | 0.204 | 0.172 | 3.055 | 2.462 | 0.847 | 2.064 | 4.514 | 2.597 |
| | | | 128 | 0.184 | 0.180 | 0.113 | 1.685 | 1.642 | 0.858 | 1.240 | 1.349 | 1.397 |
| | | 0.00100 | 16 | 0.413 | 0.371 | 0.351 | 4.833 | 2.619 | 0.906 | 4.045 | 11.755 | 3.291 |
| | | | 32 | 0.313 | 0.289 | 0.244 | 3.997 | 3.379 | 0.878 | 2.173 | 4.069 | 2.551 |
| | | | 64 | 0.247 | 0.221 | 0.166 | 2.766 | 2.487 | 0.855 | 1.125 | 1.198 | 1.547 |
| | | | 128 | 0.189 | 0.188 | 0.111 | 1.666 | 1.665 | 0.825 | 0.648 | 0.645 | 0.581 |
| | | 0.00400 | 16 | 0.407 | 0.362 | 0.347 | 4.721 | 3.572 | 0.901 | 1.750 | 3.438 | 2.870 |
| | | | 32 | 0.307 | 0.290 | 0.241 | 3.898 | 3.548 | 0.870 | 1.097 | 1.177 | 1.789 |
| | | | 64 | 0.221 | 0.220 | 0.153 | 2.227 | 2.220 | 0.865 | 0.628 | 0.624 | 0.588 |
| | | | 128 | 0.164 | 0.164 | 0.099 | 1.236 | 1.236 | 0.712 | 0.338 | 0.338 | 0.254 |
| 0.3 | 0.5 | 0.00025 | 16 | 0.369 | 0.331 | 0.325 | 1.223 | 1.271 | 0.874 | 4.598 | 16.720 | 3.238 |
| | | | 32 | 0.303 | 0.283 | 0.250 | 1.121 | 1.005 | 0.852 | 2.321 | 4.539 | 2.475 |
| | | | 64 | 0.228 | 0.219 | 0.182 | 0.880 | 0.850 | 0.805 | 1.190 | 1.320 | 1.603 |
| | | | 128 | 0.157 | 0.157 | 0.127 | 0.679 | 0.679 | 0.763 | 0.674 | 0.671 | 0.535 |
| | | 0.00100 | 16 | 0.376 | 0.355 | 0.324 | 1.265 | 1.127 | 0.873 | 1.923 | 4.403 | 2.900 |
| | | | 32 | 0.312 | 0.303 | 0.248 | 1.158 | 1.118 | 0.837 | 1.251 | 1.335 | 1.369 |
| | | | 64 | 0.238 | 0.237 | 0.176 | 0.904 | 0.898 | 0.781 | 0.652 | 0.642 | 0.607 |
| | | | 128 | 0.173 | 0.173 | 0.123 | 0.704 | 0.704 | 0.676 | 0.359 | 0.359 | 0.261 |
| | | 0.00400 | 16 | 0.379 | 0.364 | 0.317 | 1.258 | 1.166 | 0.860 | 1.173 | 1.286 | 1.767 |
| | | | 32 | 0.306 | 0.296 | 0.237 | 1.127 | 1.086 | 0.799 | 0.702 | 0.696 | 0.592 |
| | | | 64 | 0.210 | 0.210 | 0.160 | 0.823 | 0.823 | 0.681 | 0.357 | 0.357 | 0.272 |
| | | | 128 | 0.142 | 0.142 | 0.099 | 0.537 | 0.537 | 0.466 | 0.184 | 0.184 | 0.139 |
| | 0.75 | 0.00025 | 16 | 0.413 | 0.337 | 0.354 | 1.379 | 1.093 | 0.881 | 7.255 | 28.074 | 3.469 |
| | | | 32 | 0.323 | 0.286 | 0.255 | 1.170 | 1.032 | 0.846 | 2.800 | 8.221 | 2.975 |
| | | | 64 | 0.246 | 0.246 | 0.181 | 0.930 | 0.950 | 0.800 | 1.710 | 2.668 | 2.130 |
| | | | 128 | 0.177 | 0.177 | 0.129 | 0.718 | 0.718 | 0.763 | 0.943 | 0.940 | 0.967 |
| | | 0.00100 | 16 | 0.407 | 0.442 | 0.350 | 1.309 | 1.353 | 0.879 | 3.101 | 7.888 | 3.000 |
| | | | 32 | 0.292 | 0.283 | 0.252 | 1.078 | 1.063 | 0.849 | 1.569 | 2.290 | 2.237 |
| | | | 64 | 0.236 | 0.232 | 0.177 | 0.894 | 0.879 | 0.788 | 0.893 | 0.889 | 0.853 |
| | | | 128 | 0.162 | 0.162 | 0.121 | 0.684 | 0.684 | 0.701 | 0.473 | 0.473 | 0.350 |
| | | 0.00400 | 16 | 0.395 | 0.402 | 0.345 | 1.325 | 1.333 | 0.856 | 1.686 | 2.196 | 2.047 |
| | | | 32 | 0.295 | 0.295 | 0.240 | 1.045 | 1.036 | 0.823 | 0.889 | 0.883 | 0.890 |
| | | | 64 | 0.219 | 0.219 | 0.163 | 0.813 | 0.813 | 0.705 | 0.479 | 0.479 | 0.359 |
| | | | 128 | 0.147 | 0.147 | 0.106 | 0.561 | 0.561 | 0.524 | 0.253 | 0.253 | 0.185 |
| | 1.0 | 0.00025 | 16 | 0.416 | 0.358 | 0.359 | 1.279 | 1.001 | 0.883 | 11.312 | 44.169 | 3.552 |
| | | | 32 | 0.321 | 0.306 | 0.261 | 1.182 | 0.791 | 0.832 | 3.316 | 10.293 | 3.338 |
| | | | 64 | 0.239 | 0.259 | 0.183 | 0.908 | 0.983 | 0.796 | 1.922 | 3.931 | 2.535 |
| | | | 128 | 0.169 | 0.166 | 0.124 | 0.694 | 0.684 | 0.768 | 1.205 | 1.291 | 1.338 |
| | | 0.00100 | 16 | 0.402 | 0.380 | 0.362 | 1.288 | 1.343 | 0.879 | 4.278 | 14.240 | 3.248 |
| | | | 32 | 0.321 | 0.312 | 0.257 | 1.177 | 1.090 | 0.819 | 2.249 | 4.302 | 2.494 |
| | | | 64 | 0.222 | 0.217 | 0.175 | 0.860 | 0.845 | 0.753 | 1.109 | 1.155 | 1.353 |
| | | | 128 | 0.172 | 0.172 | 0.121 | 0.673 | 0.673 | 0.662 | 0.608 | 0.608 | 0.498 |
| | | 0.00400 | 16 | 0.426 | 0.418 | 0.356 | 1.385 | 1.299 | 0.852 | 1.649 | 3.082 | 2.865 |
| | | | 32 | 0.305 | 0.297 | 0.247 | 1.097 | 1.039 | 0.804 | 1.111 | 1.153 | 1.362 |
| | | | 64 | 0.222 | 0.222 | 0.163 | 0.808 | 0.808 | 0.665 | 0.600 | 0.600 | 0.464 |
| | | | 128 | 0.150 | 0.150 | 0.105 | 0.535 | 0.535 | 0.492 | 0.312 | 0.312 | 0.236 |
| 0.5 | 0.5 | 0.00025 | 16 | 0.386 | 0.379 | 0.342 | 0.737 | 0.742 | 0.826 | 4.713 | 15.606 | 3.134 |
| | | | 32 | 0.330 | 0.338 | 0.274 | 0.755 | 0.780 | 0.776 | 1.963 | 4.121 | 2.570 |
| | | | 64 | 0.262 | 0.265 | 0.204 | 0.669 | 0.669 | 0.697 | 1.177 | 1.245 | 1.307 |
| | | | 128 | 0.201 | 0.201 | 0.150 | 0.566 | 0.567 | 0.589 | 0.611 | 0.607 | 0.483 |
| | | 0.00100 | 16 | 0.396 | 0.410 | 0.337 | 0.754 | 0.759 | 0.828 | 2.112 | 3.746 | 2.527 |
| | | | 32 | 0.347 | 0.338 | 0.265 | 0.762 | 0.742 | 0.763 | 1.132 | 1.184 | 1.380 |
| | | | 64 | 0.246 | 0.246 | 0.200 | 0.624 | 0.622 | 0.628 | 0.624 | 0.616 | 0.528 |
| | | | 128 | 0.172 | 0.172 | 0.136 | 0.491 | 0.491 | 0.483 | 0.330 | 0.330 | 0.239 |
| | | 0.00400 | 16 | 0.397 | 0.400 | 0.329 | 0.765 | 0.767 | 0.795 | 1.233 | 1.269 | 1.151 |
| | | | 32 | 0.298 | 0.298 | 0.251 | 0.675 | 0.675 | 0.690 | 0.630 | 0.626 | 0.487 |
| | | | 64 | 0.225 | 0.225 | 0.167 | 0.547 | 0.547 | 0.511 | 0.318 | 0.318 | 0.246 |
| | | | 128 | 0.139 | 0.139 | 0.099 | 0.326 | 0.326 | 0.265 | 0.167 | 0.167 | 0.128 |
| | 0.75 | 0.00025 | 16 | 0.415 | 0.395 | 0.366 | 0.782 | 0.725 | 0.850 | 6.488 | 23.710 | 3.461 |
| | | | 32 | 0.338 | 0.310 | 0.272 | 0.752 | 0.691 | 0.796 | 2.800 | 8.370 | 2.902 |
| | | | 64 | 0.251 | 0.250 | 0.203 | 0.652 | 0.649 | 0.697 | 1.586 | 2.137 | 1.916 |
| | | | 128 | 0.186 | 0.186 | 0.145 | 0.584 | 0.585 | 0.628 | 0.869 | 0.855 | 0.932 |
| | | 0.00100 | 16 | 0.411 | 0.416 | 0.361 | 0.764 | 0.735 | 0.842 | 2.964 | 6.415 | 2.818 |
| | | | 32 | 0.333 | 0.313 | 0.271 | 0.756 | 0.709 | 0.776 | 1.576 | 2.015 | 1.917 |
| | | | 64 | 0.237 | 0.237 | 0.192 | 0.609 | 0.607 | 0.666 | 0.899 | 0.896 | 0.721 |
| | | | 128 | 0.175 | 0.175 | 0.135 | 0.512 | 0.512 | 0.512 | 0.459 | 0.459 | 0.328 |
| | | 0.00400 | 16 | 0.399 | 0.391 | 0.359 | 0.756 | 0.745 | 0.804 | 1.479 | 1.753 | 1.843 |
| | | | 32 | 0.319 | 0.320 | 0.262 | 0.720 | 0.720 | 0.704 | 0.841 | 0.829 | 0.851 |
| | | | 64 | 0.231 | 0.231 | 0.173 | 0.565 | 0.565 | 0.540 | 0.455 | 0.455 | 0.325 |
| | | | 128 | 0.152 | 0.152 | 0.110 | 0.381 | 0.381 | 0.329 | 0.230 | 0.230 | 0.170 |
| | 1.0 | 0.00025 | 16 | 0.410 | 0.429 | 0.372 | 0.759 | 0.634 | 0.852 | 10.341 | 32.461 | 3.507 |
| | | | 32 | 0.332 | 0.339 | 0.279 | 0.751 | 0.818 | 0.769 | 3.761 | 13.812 | 3.147 |
| | | | 64 | 0.251 | 0.261 | 0.195 | 0.658 | 0.646 | 0.721 | 1.891 | 3.357 | 2.355 |
| | | | 128 | 0.182 | 0.180 | 0.139 | 0.577 | 0.569 | 0.618 | 1.141 | 1.177 | 1.119 |
| | | 0.00100 | 16 | 0.436 | 0.422 | 0.371 | 0.806 | 0.652 | 0.836 | 3.377 | 9.421 | 3.253 |
| | | | 32 | 0.337 | 0.335 | 0.271 | 0.755 | 0.726 | 0.771 | 1.763 | 2.836 | 2.422 |
| | | | 64 | 0.252 | 0.243 | 0.190 | 0.652 | 0.633 | 0.664 | 1.085 | 1.115 | 1.222 |
| | | | 128 | 0.175 | 0.175 | 0.130 | 0.498 | 0.498 | 0.483 | 0.554 | 0.554 | 0.417 |
| | | 0.00400 | 16 | 0.417 | 0.404 | 0.370 | 0.801 | 0.768 | 0.782 | 1.965 | 2.865 | 2.277 |
| | | | 32 | 0.323 | 0.323 | 0.257 | 0.720 | 0.707 | 0.703 | 1.092 | 1.118 | 1.207 |
| | | | 64 | 0.224 | 0.224 | 0.173 | 0.560 | 0.560 | 0.535 | 0.567 | 0.567 | 0.434 |
| | | | 128 | 0.160 | 0.160 | 0.110 | 0.364 | 0.364 | 0.311 | 0.292 | 0.292 | 0.209 |

**Table S7:** Simulation results for $\zeta$ with $k = 4$.

| | | | | $\lambda$ | | | $\mu$ | | | $\theta$ | | |
|---|---|---|---|---|---|---|---|---|---|---|---|---|
| $\epsilon$ | $\rho$ | $\theta$ | $N$ | RMSE | $RMSE_c$ | CV | RMSE | $RMSE_c$ | CV | RMSE | $RMSE_c$ | CV |
| 0.1 | 0.5 | 0.00025 | 16 | 0.415 | 0.369 | 0.345 | 5.391 | 4.014 | 0.945 | 4.876 | 17.859 | 3.363 |
| | | | 32 | 0.336 | 0.369 | 0.245 | 4.317 | 4.481 | 0.886 | 2.014 | 5.139 | 2.779 |
| | | | 64 | 0.252 | 0.247 | 0.165 | 2.845 | 2.742 | 0.866 | 1.285 | 1.510 | 1.725 |
| | | | 128 | 0.179 | 0.175 | 0.112 | 1.751 | 1.720 | 0.851 | 0.654 | 0.641 | 0.661 |
| | | 0.00100 | 16 | 0.403 | 0.365 | 0.333 | 5.073 | 4.514 | 0.944 | 2.413 | 4.871 | 2.687 |
| | | | 32 | 0.329 | 0.300 | 0.236 | 4.099 | 3.717 | 0.888 | 1.274 | 1.477 | 1.791 |
| | | | 64 | 0.258 | 0.254 | 0.164 | 2.931 | 2.894 | 0.852 | 0.682 | 0.676 | 0.644 |
| | | | 128 | 0.168 | 0.168 | 0.106 | 1.567 | 1.567 | 0.844 | 0.363 | 0.363 | 0.288 |
| | | 0.00400 | 16 | 0.425 | 0.405 | 0.337 | 5.403 | 5.101 | 0.928 | 1.199 | 1.296 | 1.677 |
| | | | 32 | 0.319 | 0.310 | 0.229 | 4.020 | 3.913 | 0.875 | 0.687 | 0.681 | 0.656 |
| | | | 64 | 0.210 | 0.210 | 0.145 | 2.302 | 2.302 | 0.817 | 0.361 | 0.361 | 0.286 |
| | | | 128 | 0.158 | 0.158 | 0.092 | 1.296 | 1.296 | 0.733 | 0.194 | 0.194 | 0.150 |
| | 0.75 | 0.00025 | 16 | 0.418 | 0.798 | 0.349 | 5.089 | 8.789 | 0.926 | 8.494 | 40.406 | 3.502 |
| | | | 32 | 0.344 | 0.362 | 0.248 | 4.451 | 4.673 | 0.867 | 3.317 | 11.119 | 3.016 |
| | | | 64 | 0.240 | 0.233 | 0.167 | 2.781 | 2.637 | 0.862 | 1.669 | 2.631 | 2.187 |
| | | | 128 | 0.188 | 0.186 | 0.113 | 1.762 | 1.716 | 0.855 | 0.899 | 0.887 | 1.137 |
| | | 0.00100 | 16 | 0.425 | 0.329 | 0.354 | 5.406 | 3.821 | 0.918 | 2.789 | 10.040 | 3.340 |
| | | | 32 | 0.320 | 0.288 | 0.240 | 4.054 | 3.634 | 0.876 | 1.611 | 2.464 | 2.323 |
| | | | 64 | 0.259 | 0.242 | 0.165 | 2.869 | 2.678 | 0.858 | 0.885 | 0.868 | 1.174 |
| | | | 128 | 0.172 | 0.172 | 0.109 | 1.634 | 1.638 | 0.828 | 0.493 | 0.488 | 0.420 |
| | | 0.00400 | 16 | 0.444 | 0.404 | 0.350 | 5.538 | 4.929 | 0.909 | 1.638 | 2.603 | 2.531 |
| | | | 32 | 0.335 | 0.318 | 0.235 | 4.052 | 3.811 | 0.872 | 0.913 | 0.903 | 1.162 |
| | | | 64 | 0.227 | 0.227 | 0.155 | 2.499 | 2.499 | 0.825 | 0.517 | 0.517 | 0.404 |
| | | | 128 | 0.155 | 0.155 | 0.098 | 1.243 | 1.243 | 0.776 | 0.262 | 0.262 | 0.199 |
| | 1.0 | 0.00025 | 16 | 0.427 | – | 0.360 | 5.241 | – | 0.906 | 12.236 | – | 3.618 |
| | | | 32 | 0.327 | 0.258 | 0.248 | 4.259 | 3.146 | 0.868 | 4.795 | 18.023 | 3.187 |
| | | | 64 | 0.253 | 0.204 | 0.172 | 3.046 | 2.462 | 0.848 | 2.061 | 4.501 | 2.599 |
| | | | 128 | 0.184 | 0.181 | 0.114 | 1.691 | 1.645 | 0.858 | 1.239 | 1.348 | 1.398 |
| | | 0.00100 | 16 | 0.418 | 0.374 | 0.355 | 4.888 | 2.690 | 0.914 | 4.083 | 11.709 | 3.306 |
| | | | 32 | 0.312 | 0.289 | 0.243 | 3.998 | 3.387 | 0.874 | 2.153 | 4.023 | 2.551 |
| | | | 64 | 0.248 | 0.220 | 0.166 | 2.770 | 2.485 | 0.853 | 1.118 | 1.189 | 1.540 |
| | | | 128 | 0.189 | 0.188 | 0.111 | 1.670 | 1.669 | 0.824 | 0.648 | 0.645 | 0.582 |
| | | 0.00400 | 16 | 0.413 | 0.363 | 0.351 | 4.783 | 3.558 | 0.905 | 1.746 | 3.447 | 2.842 |
| | | | 32 | 0.307 | 0.292 | 0.241 | 3.906 | 3.559 | 0.870 | 1.101 | 1.184 | 1.778 |
| | | | 64 | 0.220 | 0.220 | 0.153 | 2.226 | 2.220 | 0.859 | 0.628 | 0.624 | 0.585 |
| | | | 128 | 0.164 | 0.164 | 0.099 | 1.237 | 1.237 | 0.713 | 0.338 | 0.338 | 0.253 |
| 0.3 | 0.5 | 0.00025 | 16 | 0.406 | 0.372 | 0.355 | 1.395 | 1.468 | 0.921 | 4.535 | 16.451 | 3.272 |
| | | | 32 | 0.308 | 0.286 | 0.254 | 1.141 | 1.018 | 0.860 | 2.319 | 4.532 | 2.461 |
| | | | 64 | 0.229 | 0.220 | 0.182 | 0.884 | 0.852 | 0.805 | 1.187 | 1.316 | 1.604 |
| | | | 128 | 0.157 | 0.157 | 0.127 | 0.680 | 0.681 | 0.764 | 0.675 | 0.671 | 0.535 |
| | | 0.00100 | 16 | 0.419 | 0.384 | 0.355 | 1.452 | 1.253 | 0.921 | 1.914 | 4.394 | 2.908 |
| | | | 32 | 0.317 | 0.308 | 0.252 | 1.178 | 1.136 | 0.845 | 1.249 | 1.332 | 1.379 |
| | | | 64 | 0.238 | 0.237 | 0.176 | 0.902 | 0.895 | 0.780 | 0.653 | 0.643 | 0.606 |
| | | | 128 | 0.174 | 0.174 | 0.123 | 0.705 | 0.705 | 0.676 | 0.358 | 0.358 | 0.261 |
| | | 0.00400 | 16 | 0.419 | 0.396 | 0.344 | 1.431 | 1.304 | 0.902 | 1.168 | 1.278 | 1.792 |
| | | | 32 | 0.312 | 0.300 | 0.240 | 1.148 | 1.101 | 0.806 | 0.702 | 0.696 | 0.593 |
| | | | 64 | 0.210 | 0.210 | 0.160 | 0.822 | 0.822 | 0.681 | 0.358 | 0.358 | 0.272 |
| | | | 128 | 0.142 | 0.142 | 0.099 | 0.538 | 0.538 | 0.467 | 0.184 | 0.184 | 0.139 |
| | 0.75 | 0.00025 | 16 | 0.433 | 0.340 | 0.367 | 1.456 | 1.096 | 0.903 | 7.346 | 28.006 | 3.496 |
| | | | 32 | 0.325 | 0.285 | 0.256 | 1.176 | 1.030 | 0.846 | 2.815 | 8.182 | 2.988 |
| | | | 64 | 0.246 | 0.246 | 0.181 | 0.930 | 0.948 | 0.802 | 1.713 | 2.671 | 2.147 |
| | | | 128 | 0.177 | 0.177 | 0.129 | 0.718 | 0.718 | 0.763 | 0.942 | 0.940 | 0.970 |
| | | 0.00100 | 16 | 0.420 | 0.460 | 0.360 | 1.366 | 1.435 | 0.896 | 3.099 | 7.903 | 2.999 |
| | | | 32 | 0.291 | 0.282 | 0.251 | 1.075 | 1.062 | 0.848 | 1.565 | 2.279 | 2.247 |
| | | | 64 | 0.237 | 0.233 | 0.177 | 0.896 | 0.881 | 0.788 | 0.895 | 0.891 | 0.851 |
| | | | 128 | 0.163 | 0.163 | 0.121 | 0.684 | 0.684 | 0.701 | 0.473 | 0.473 | 0.350 |
| | | 0.00400 | 16 | 0.409 | 0.416 | 0.355 | 1.378 | 1.385 | 0.872 | 1.679 | 2.184 | 2.057 |
| | | | 32 | 0.296 | 0.296 | 0.240 | 1.049 | 1.040 | 0.824 | 0.890 | 0.883 | 0.895 |
| | | | 64 | 0.220 | 0.220 | 0.163 | 0.816 | 0.816 | 0.707 | 0.480 | 0.480 | 0.359 |
| | | | 128 | 0.147 | 0.147 | 0.106 | 0.561 | 0.561 | 0.524 | 0.254 | 0.254 | 0.185 |
| | 1.0 | 0.00025 | 16 | 0.424 | 0.361 | 0.365 | 1.315 | 1.032 | 0.894 | 11.249 | 46.135 | 3.567 |
| | | | 32 | 0.321 | 0.306 | 0.261 | 1.181 | 0.787 | 0.831 | 3.315 | 10.124 | 3.306 |
| | | | 64 | 0.239 | 0.259 | 0.183 | 0.909 | 0.982 | 0.798 | 1.922 | 3.942 | 2.516 |
| | | | 128 | 0.169 | 0.166 | 0.124 | 0.693 | 0.683 | 0.766 | 1.208 | 1.295 | 1.339 |
| | | 0.00100 | 16 | 0.410 | 0.390 | 0.368 | 1.319 | 1.388 | 0.890 | 4.276 | 14.401 | 3.248 |
| | | | 32 | 0.320 | 0.310 | 0.257 | 1.173 | 1.085 | 0.817 | 2.243 | 4.287 | 2.500 |
| | | | 64 | 0.222 | 0.218 | 0.175 | 0.863 | 0.846 | 0.753 | 1.110 | 1.156 | 1.347 |
| | | | 128 | 0.171 | 0.171 | 0.120 | 0.671 | 0.671 | 0.661 | 0.610 | 0.610 | 0.498 |
| | | 0.00400 | 16 | 0.441 | 0.433 | 0.364 | 1.442 | 1.349 | 0.862 | 1.640 | 3.059 | 2.842 |
| | | | 32 | 0.305 | 0.297 | 0.247 | 1.098 | 1.040 | 0.805 | 1.110 | 1.151 | 1.358 |
| | | | 64 | 0.222 | 0.222 | 0.162 | 0.810 | 0.810 | 0.665 | 0.601 | 0.601 | 0.465 |
| | | | 128 | 0.150 | 0.150 | 0.105 | 0.534 | 0.534 | 0.492 | 0.311 | 0.311 | 0.236 |
| 0.5 | 0.5 | 0.00025 | 16 | 0.420 | 0.411 | 0.382 | 0.832 | 0.824 | 0.880 | 4.667 | 15.374 | 3.149 |
| | | | 32 | 0.338 | 0.348 | 0.281 | 0.775 | 0.803 | 0.788 | 1.964 | 4.110 | 2.575 |
| | | | 64 | 0.263 | 0.265 | 0.204 | 0.670 | 0.670 | 0.699 | 1.176 | 1.244 | 1.307 |
| | | | 128 | 0.201 | 0.201 | 0.150 | 0.565 | 0.565 | 0.587 | 0.611 | 0.608 | 0.484 |
| | | 0.00100 | 16 | 0.434 | 0.448 | 0.372 | 0.854 | 0.856 | 0.878 | 2.072 | 3.658 | 2.518 |
| | | | 32 | 0.358 | 0.348 | 0.272 | 0.786 | 0.766 | 0.776 | 1.132 | 1.184 | 1.377 |
| | | | 64 | 0.247 | 0.246 | 0.200 | 0.624 | 0.622 | 0.629 | 0.624 | 0.617 | 0.529 |
| | | | 128 | 0.172 | 0.172 | 0.135 | 0.491 | 0.491 | 0.483 | 0.330 | 0.330 | 0.239 |
| | | 0.00400 | 16 | 0.434 | 0.437 | 0.362 | 0.862 | 0.866 | 0.839 | 1.215 | 1.249 | 1.149 |
| | | | 32 | 0.304 | 0.304 | 0.255 | 0.689 | 0.688 | 0.695 | 0.629 | 0.626 | 0.490 |
| | | | 64 | 0.224 | 0.224 | 0.167 | 0.546 | 0.546 | 0.511 | 0.317 | 0.317 | 0.246 |
| | | | 128 | 0.139 | 0.139 | 0.099 | 0.326 | 0.326 | 0.265 | 0.167 | 0.167 | 0.128 |
| | 0.75 | 0.00025 | 16 | 0.431 | 0.421 | 0.381 | 0.820 | 0.794 | 0.871 | 6.402 | 23.842 | 3.452 |
| | | | 32 | 0.340 | 0.311 | 0.272 | 0.756 | 0.694 | 0.797 | 2.808 | 8.405 | 2.905 |
| | | | 64 | 0.251 | 0.251 | 0.202 | 0.651 | 0.649 | 0.696 | 1.580 | 2.125 | 1.918 |
| | | | 128 | 0.186 | 0.186 | 0.144 | 0.583 | 0.584 | 0.626 | 0.870 | 0.856 | 0.932 |
| | | 0.00100 | 16 | 0.426 | 0.426 | 0.376 | 0.801 | 0.762 | 0.865 | 2.954 | 6.381 | 2.806 |
| | | | 32 | 0.336 | 0.313 | 0.273 | 0.759 | 0.707 | 0.777 | 1.572 | 2.008 | 1.926 |
| | | | 64 | 0.237 | 0.236 | 0.192 | 0.609 | 0.607 | 0.667 | 0.897 | 0.894 | 0.721 |
| | | | 128 | 0.175 | 0.175 | 0.135 | 0.512 | 0.512 | 0.511 | 0.457 | 0.457 | 0.326 |
| | | 0.00400 | 16 | 0.416 | 0.408 | 0.376 | 0.799 | 0.789 | 0.828 | 1.479 | 1.754 | 1.843 |
| | | | 32 | 0.320 | 0.320 | 0.262 | 0.720 | 0.721 | 0.705 | 0.839 | 0.828 | 0.853 |
| | | | 64 | 0.232 | 0.232 | 0.174 | 0.567 | 0.567 | 0.541 | 0.455 | 0.455 | 0.326 |
| | | | 128 | 0.152 | 0.152 | 0.110 | 0.381 | 0.381 | 0.329 | 0.231 | 0.231 | 0.171 |
| | 1.0 | 0.00025 | 16 | 0.418 | 0.429 | 0.380 | 0.778 | 0.647 | 0.862 | 10.314 | 32.809 | 3.499 |
| | | | 32 | 0.334 | 0.345 | 0.279 | 0.754 | 0.831 | 0.768 | 3.766 | 13.737 | 3.131 |
| | | | 64 | 0.250 | 0.261 | 0.194 | 0.657 | 0.646 | 0.719 | 1.890 | 3.358 | 2.335 |
| | | | 128 | 0.181 | 0.180 | 0.139 | 0.577 | 0.569 | 0.619 | 1.142 | 1.178 | 1.113 |
| | | 0.00100 | 16 | 0.448 | 0.423 | 0.380 | 0.834 | 0.656 | 0.850 | 3.383 | 9.466 | 3.258 |
| | | | 32 | 0.337 | 0.336 | 0.271 | 0.754 | 0.727 | 0.772 | 1.767 | 2.838 | 2.432 |
| | | | 64 | 0.252 | 0.243 | 0.190 | 0.651 | 0.634 | 0.663 | 1.088 | 1.119 | 1.226 |
| | | | 128 | 0.175 | 0.175 | 0.130 | 0.499 | 0.499 | 0.484 | 0.553 | 0.553 | 0.416 |
| | | 0.00400 | 16 | 0.432 | 0.419 | 0.381 | 0.834 | 0.801 | 0.795 | 1.958 | 2.851 | 2.269 |
| | | | 32 | 0.324 | 0.324 | 0.258 | 0.721 | 0.708 | 0.703 | 1.088 | 1.113 | 1.200 |
| | | | 64 | 0.223 | 0.223 | 0.172 | 0.559 | 0.559 | 0.534 | 0.567 | 0.567 | 0.433 |
| | | | 128 | 0.161 | 0.161 | 0.110 | 0.365 | 0.365 | 0.311 | 0.292 | 0.292 | 0.209 |

**Table S8:** Simulation results for $\psi$ with $k = 1$.

| | | | | $\lambda$ | | | $\mu$ | | | $\theta$ | | |
|---|---|---|---|---|---|---|---|---|---|---|---|---|
| $\epsilon$ | $\rho$ | $\theta$ | $N$ | RMSE | RMSE$_c$ | CV | RMSE | RMSE$_c$ | CV | RMSE | RMSE$_c$ | CV |
| 0.1 | 0.5 | 0.00025 | 16 | 0.305 | 0.304 | 0.264 | 3.053 | 3.102 | 0.848 | 8.339 | 36.055 | 3.321 |
| | | | 32 | 0.241 | 0.234 | 0.199 | 2.899 | 2.794 | 0.843 | 3.591 | 8.464 | 2.634 |
| | | | 64 | 0.204 | 0.195 | 0.149 | 2.367 | 2.300 | 0.850 | 1.554 | 2.117 | 1.953 |
| | | | 128 | 0.148 | 0.146 | 0.105 | 1.543 | 1.531 | 0.849 | 0.810 | 0.800 | 0.798 |
| | | 0.00100 | 16 | 0.297 | 0.297 | 0.263 | 2.816 | 3.106 | 0.859 | 2.674 | 8.273 | 3.129 |
| | | | 32 | 0.246 | 0.235 | 0.201 | 3.035 | 2.785 | 0.841 | 1.419 | 1.862 | 2.117 |
| | | | 64 | 0.208 | 0.197 | 0.148 | 2.420 | 2.303 | 0.840 | 0.819 | 0.803 | 0.905 |
| | | | 128 | 0.156 | 0.156 | 0.105 | 1.590 | 1.590 | 0.830 | 0.416 | 0.416 | 0.343 |
| | | 0.00400 | 16 | 0.295 | 0.305 | 0.263 | 2.985 | 2.824 | 0.862 | 1.412 | 2.033 | 2.451 |
| | | | 32 | 0.256 | 0.253 | 0.199 | 2.983 | 2.956 | 0.846 | 0.861 | 0.846 | 1.006 |
| | | | 64 | 0.195 | 0.195 | 0.145 | 2.155 | 2.155 | 0.831 | 0.463 | 0.463 | 0.349 |
| | | | 128 | 0.157 | 0.157 | 0.102 | 1.439 | 1.439 | 0.773 | 0.255 | 0.255 | 0.184 |
| | 0.75 | 0.00025 | 16 | 0.350 | 0.340 | 0.303 | 4.123 | 3.479 | 0.847 | 12.895 | 64.985 | 3.390 |
| | | | 32 | 0.343 | 0.307 | 0.235 | 4.258 | 3.798 | 0.846 | 3.566 | 11.754 | 3.083 |
| | | | 64 | 0.267 | 0.285 | 0.169 | 3.058 | 3.150 | 0.850 | 1.948 | 3.671 | 2.374 |
| | | | 128 | 0.172 | 0.163 | 0.111 | 1.610 | 1.544 | 0.868 | 1.031 | 1.050 | 1.183 |
| | | 0.00100 | 16 | 0.352 | 0.441 | 0.305 | 4.260 | 5.495 | 0.842 | 3.099 | 13.674 | 3.410 |
| | | | 32 | 0.313 | 0.289 | 0.233 | 4.008 | 3.684 | 0.850 | 1.826 | 3.105 | 2.431 |
| | | | 64 | 0.250 | 0.247 | 0.164 | 2.779 | 2.683 | 0.863 | 1.038 | 1.058 | 1.282 |
| | | | 128 | 0.181 | 0.180 | 0.112 | 1.763 | 1.751 | 0.822 | 0.532 | 0.527 | 0.474 |
| | | 0.00400 | 16 | 0.365 | 0.357 | 0.304 | 4.377 | 4.293 | 0.841 | 1.506 | 3.167 | 2.961 |
| | | | 32 | 0.315 | 0.298 | 0.229 | 3.871 | 3.646 | 0.869 | 1.029 | 1.043 | 1.284 |
| | | | 64 | 0.241 | 0.241 | 0.158 | 2.560 | 2.560 | 0.840 | 0.561 | 0.561 | 0.463 |
| | | | 128 | 0.169 | 0.169 | 0.105 | 1.424 | 1.424 | 0.796 | 0.301 | 0.301 | 0.230 |
| | 1.0 | 0.00025 | 16 | 0.369 | 0.424 | 0.329 | 4.340 | 5.614 | 0.864 | 13.904 | 102.553 | 3.518 |
| | | | 32 | 0.304 | 0.272 | 0.244 | 3.974 | 3.520 | 0.872 | 4.921 | 19.986 | 3.173 |
| | | | 64 | 0.261 | 0.229 | 0.171 | 3.027 | 2.517 | 0.849 | 1.994 | 5.206 | 2.729 |
| | | | 128 | 0.194 | 0.185 | 0.117 | 1.876 | 1.799 | 0.839 | 1.185 | 1.278 | 1.470 |
| | | 0.00100 | 16 | 0.378 | 0.327 | 0.331 | 4.621 | 3.699 | 0.859 | 4.783 | 21.480 | 3.348 |
| | | | 32 | 0.320 | 0.263 | 0.242 | 4.060 | 3.163 | 0.865 | 2.193 | 5.167 | 2.759 |
| | | | 64 | 0.246 | 0.232 | 0.166 | 2.714 | 2.534 | 0.863 | 1.166 | 1.275 | 1.645 |
| | | | 128 | 0.188 | 0.187 | 0.111 | 1.670 | 1.643 | 0.822 | 0.732 | 0.721 | 0.621 |
| | | 0.00400 | 16 | 0.377 | 0.347 | 0.327 | 4.281 | 3.435 | 0.875 | 1.962 | 4.191 | 2.874 |
| | | | 32 | 0.352 | 0.311 | 0.240 | 4.289 | 3.785 | 0.848 | 1.164 | 1.284 | 1.802 |
| | | | 64 | 0.224 | 0.217 | 0.157 | 2.286 | 2.209 | 0.860 | 0.659 | 0.631 | 0.773 |
| | | | 128 | 0.163 | 0.163 | 0.104 | 1.213 | 1.213 | 0.784 | 0.353 | 0.353 | 0.270 |
| 0.3 | 0.5 | 0.00025 | 16 | 0.341 | 0.352 | 0.269 | 0.823 | 0.760 | 0.827 | 7.493 | 28.574 | 3.285 |
| | | | 32 | 0.249 | 0.249 | 0.207 | 0.791 | 0.743 | 0.818 | 2.739 | 7.694 | 2.817 |
| | | | 64 | 0.203 | 0.205 | 0.160 | 0.765 | 0.757 | 0.789 | 1.546 | 1.905 | 1.698 |
| | | | 128 | 0.158 | 0.156 | 0.121 | 0.688 | 0.684 | 0.750 | 0.777 | 0.769 | 0.747 |
| | | 0.00100 | 16 | 0.338 | 0.383 | 0.269 | 0.800 | 0.750 | 0.832 | 2.380 | 6.690 | 3.110 |
| | | | 32 | 0.263 | 0.270 | 0.207 | 0.827 | 0.837 | 0.811 | 1.705 | 2.220 | 1.825 |
| | | | 64 | 0.215 | 0.214 | 0.163 | 0.812 | 0.810 | 0.761 | 0.816 | 0.813 | 0.715 |
| | | | 128 | 0.148 | 0.148 | 0.117 | 0.642 | 0.642 | 0.713 | 0.405 | 0.405 | 0.311 |
| | | 0.00400 | 16 | 0.318 | 0.328 | 0.269 | 0.808 | 0.806 | 0.827 | 1.618 | 2.105 | 2.016 |
| | | | 32 | 0.265 | 0.265 | 0.208 | 0.888 | 0.887 | 0.785 | 0.874 | 0.867 | 0.809 |
| | | | 64 | 0.201 | 0.201 | 0.155 | 0.748 | 0.748 | 0.744 | 0.425 | 0.425 | 0.333 |
| | | | 128 | 0.150 | 0.150 | 0.113 | 0.599 | 0.599 | 0.578 | 0.247 | 0.247 | 0.177 |
| | 0.75 | 0.00025 | 16 | 0.347 | 0.378 | 0.309 | 1.027 | 0.788 | 0.817 | 9.544 | 41.342 | 3.450 |
| | | | 32 | 0.328 | 0.348 | 0.246 | 1.216 | 1.262 | 0.799 | 3.483 | 12.692 | 3.073 |
| | | | 64 | 0.233 | 0.202 | 0.181 | 0.890 | 0.796 | 0.797 | 1.750 | 2.874 | 2.244 |
| | | | 128 | 0.170 | 0.168 | 0.128 | 0.719 | 0.718 | 0.764 | 1.001 | 1.010 | 1.111 |
| | | 0.00100 | 16 | 0.352 | 0.363 | 0.308 | 1.028 | 1.073 | 0.813 | 3.407 | 10.001 | 3.124 |
| | | | 32 | 0.299 | 0.296 | 0.240 | 1.045 | 1.009 | 0.828 | 1.525 | 2.712 | 2.600 |
| | | | 64 | 0.228 | 0.228 | 0.175 | 0.854 | 0.859 | 0.796 | 1.001 | 1.010 | 1.177 |
| | | | 128 | 0.168 | 0.168 | 0.125 | 0.669 | 0.669 | 0.697 | 0.556 | 0.556 | 0.426 |
| | | 0.00400 | 16 | 0.357 | 0.356 | 0.307 | 1.062 | 0.973 | 0.815 | 1.648 | 2.677 | 2.557 |
| | | | 32 | 0.283 | 0.282 | 0.234 | 1.005 | 0.988 | 0.821 | 0.971 | 0.972 | 1.212 |
| | | | 64 | 0.231 | 0.231 | 0.169 | 0.863 | 0.863 | 0.717 | 0.553 | 0.553 | 0.417 |
| | | | 128 | 0.160 | 0.160 | 0.115 | 0.627 | 0.627 | 0.602 | 0.285 | 0.285 | 0.220 |
| | 1.0 | 0.00025 | 16 | 0.369 | 0.369 | 0.334 | 1.065 | 0.871 | 0.847 | 13.410 | 67.144 | 3.469 |
| | | | 32 | 0.314 | 0.306 | 0.254 | 1.165 | 1.111 | 0.823 | 4.368 | 19.682 | 3.208 |
| | | | 64 | 0.247 | 0.245 | 0.183 | 0.945 | 0.900 | 0.784 | 2.160 | 4.146 | 2.406 |
| | | | 128 | 0.167 | 0.153 | 0.126 | 0.711 | 0.691 | 0.773 | 1.229 | 1.403 | 1.621 |
| | | 0.00100 | 16 | 0.376 | 0.353 | 0.332 | 1.125 | 0.966 | 0.832 | 3.602 | 13.623 | 3.394 |
| | | | 32 | 0.306 | 0.275 | 0.252 | 1.137 | 1.020 | 0.810 | 2.128 | 4.555 | 2.630 |
| | | | 64 | 0.221 | 0.216 | 0.175 | 0.849 | 0.836 | 0.790 | 1.180 | 1.264 | 1.460 |
| | | | 128 | 0.167 | 0.167 | 0.120 | 0.642 | 0.642 | 0.681 | 0.676 | 0.673 | 0.512 |
| | | 0.00400 | 16 | 0.370 | 0.360 | 0.328 | 1.058 | 1.022 | 0.837 | 1.886 | 3.591 | 2.720 |
| | | | 32 | 0.309 | 0.314 | 0.243 | 1.103 | 1.112 | 0.821 | 1.112 | 1.201 | 1.769 |
| | | | 64 | 0.232 | 0.230 | 0.167 | 0.824 | 0.817 | 0.730 | 0.618 | 0.615 | 0.581 |
| | | | 128 | 0.156 | 0.156 | 0.110 | 0.541 | 0.541 | 0.522 | 0.358 | 0.358 | 0.250 |
| 0.5 | 0.5 | 0.00025 | 16 | 0.402 | 0.511 | 0.275 | 0.671 | 0.738 | 0.793 | 5.817 | 17.602 | 3.315 |
| | | | 32 | 0.311 | 0.327 | 0.215 | 0.656 | 0.666 | 0.760 | 2.724 | 6.130 | 2.611 |
| | | | 64 | 0.227 | 0.224 | 0.176 | 0.605 | 0.602 | 0.683 | 1.509 | 1.865 | 1.694 |
| | | | 128 | 0.191 | 0.191 | 0.136 | 0.577 | 0.578 | 0.615 | 0.776 | 0.770 | 0.621 |
| | | 0.00100 | 16 | 0.404 | 0.436 | 0.273 | 0.671 | 0.674 | 0.805 | 2.371 | 6.067 | 2.959 |
| | | | 32 | 0.297 | 0.301 | 0.218 | 0.644 | 0.637 | 0.739 | 1.566 | 2.005 | 1.838 |
| | | | 64 | 0.231 | 0.231 | 0.174 | 0.584 | 0.585 | 0.641 | 0.836 | 0.830 | 0.631 |
| | | | 128 | 0.177 | 0.177 | 0.133 | 0.508 | 0.508 | 0.517 | 0.397 | 0.397 | 0.287 |
| | | 0.00400 | 16 | 0.370 | 0.382 | 0.277 | 0.647 | 0.633 | 0.768 | 1.730 | 2.072 | 1.621 |
| | | | 32 | 0.288 | 0.288 | 0.217 | 0.638 | 0.637 | 0.682 | 0.784 | 0.782 | 0.631 |
| | | | 64 | 0.229 | 0.229 | 0.168 | 0.571 | 0.571 | 0.564 | 0.431 | 0.431 | 0.304 |
| | | | 128 | 0.157 | 0.157 | 0.120 | 0.410 | 0.410 | 0.369 | 0.241 | 0.241 | 0.162 |
| | 0.75 | 0.00025 | 16 | 0.380 | 0.434 | 0.313 | 0.659 | 0.700 | 0.788 | 8.839 | 37.512 | 3.386 |
| | | | 32 | 0.309 | 0.322 | 0.254 | 0.691 | 0.670 | 0.757 | 3.357 | 8.969 | 2.896 |
| | | | 64 | 0.242 | 0.244 | 0.197 | 0.643 | 0.647 | 0.719 | 1.610 | 2.568 | 2.230 |
| | | | 128 | 0.193 | 0.184 | 0.145 | 0.595 | 0.577 | 0.623 | 0.963 | 0.964 | 1.053 |
| | | 0.00100 | 16 | 0.378 | 0.401 | 0.315 | 0.677 | 0.705 | 0.775 | 3.049 | 9.612 | 3.194 |
| | | | 32 | 0.311 | 0.335 | 0.251 | 0.697 | 0.736 | 0.759 | 1.591 | 2.651 | 2.440 |
| | | | 64 | 0.242 | 0.242 | 0.190 | 0.633 | 0.632 | 0.688 | 0.961 | 0.961 | 0.811 |
| | | | 128 | 0.183 | 0.183 | 0.137 | 0.539 | 0.539 | 0.551 | 0.514 | 0.514 | 0.384 |
| | | 0.00400 | 16 | 0.370 | 0.373 | 0.312 | 0.678 | 0.659 | 0.774 | 1.666 | 2.416 | 2.314 |
| | | | 32 | 0.309 | 0.308 | 0.243 | 0.713 | 0.708 | 0.729 | 0.993 | 0.996 | 0.980 |
| | | | 64 | 0.230 | 0.230 | 0.181 | 0.597 | 0.597 | 0.600 | 0.529 | 0.529 | 0.404 |
| | | | 128 | 0.153 | 0.153 | 0.122 | 0.434 | 0.434 | 0.415 | 0.271 | 0.271 | 0.203 |
| | 1.0 | 0.00025 | 16 | 0.420 | 0.463 | 0.338 | 0.719 | 0.569 | 0.802 | 10.396 | 35.653 | 3.507 |
| | | | 32 | 0.315 | 0.316 | 0.265 | 0.708 | 0.710 | 0.772 | 3.822 | 15.867 | 3.163 |
| | | | 64 | 0.253 | 0.240 | 0.196 | 0.659 | 0.632 | 0.721 | 1.960 | 3.666 | 2.410 |
| | | | 128 | 0.180 | 0.181 | 0.141 | 0.565 | 0.556 | 0.616 | 1.145 | 1.197 | 1.242 |
| | | 0.00100 | 16 | 0.403 | 0.419 | 0.336 | 0.687 | 0.615 | 0.803 | 3.566 | 11.942 | 3.233 |
| | | | 32 | 0.348 | 0.348 | 0.262 | 0.773 | 0.751 | 0.754 | 1.797 | 3.443 | 2.642 |
| | | | 64 | 0.248 | 0.249 | 0.191 | 0.644 | 0.636 | 0.666 | 1.153 | 1.233 | 1.455 |
| | | | 128 | 0.171 | 0.171 | 0.130 | 0.531 | 0.531 | 0.535 | 0.639 | 0.639 | 0.445 |
| | | 0.00400 | 16 | 0.385 | 0.387 | 0.337 | 0.716 | 0.698 | 0.786 | 1.744 | 3.368 | 2.827 |
| | | | 32 | 0.313 | 0.301 | 0.251 | 0.705 | 0.680 | 0.710 | 1.074 | 1.102 | 1.332 |
| | | | 64 | 0.223 | 0.223 | 0.172 | 0.558 | 0.558 | 0.543 | 0.574 | 0.574 | 0.460 |
| | | | 128 | 0.172 | 0.172 | 0.117 | 0.435 | 0.435 | 0.356 | 0.329 | 0.329 | 0.235 |

**Table S9:** Simulation results for $\psi$ with $k = 2$.

| | | | | $\lambda$ | | | $\mu$ | | | $\theta$ | | |
|---|---|---|---|---|---|---|---|---|---|---|---|---|
| $\epsilon$ | $\rho$ | $\theta$ | $N$ | RMSE | $RMSE_c$ | CV | RMSE | $RMSE_c$ | CV | RMSE | $RMSE_c$ | CV |
| 0.1 | 0.5 | 0.00025 | 16 | 0.382 | 0.352 | 0.317 | 4.852 | 4.659 | 0.897 | 7.442 | 32.352 | 3.359 |
| | | | 32 | 0.321 | 0.297 | 0.237 | 4.104 | 3.843 | 0.885 | 3.292 | 7.682 | 2.632 |
| | | | 64 | 0.255 | 0.238 | 0.167 | 2.936 | 2.798 | 0.859 | 1.479 | 1.976 | 1.939 |
| | | | 128 | 0.176 | 0.174 | 0.112 | 1.761 | 1.747 | 0.846 | 0.785 | 0.774 | 0.803 |
| | | 0.00100 | 16 | 0.350 | 0.357 | 0.311 | 4.400 | 4.667 | 0.909 | 2.396 | 7.317 | 3.130 |
| | | | 32 | 0.337 | 0.303 | 0.238 | 4.268 | 3.802 | 0.882 | 1.325 | 1.682 | 2.117 |
| | | | 64 | 0.256 | 0.241 | 0.164 | 2.931 | 2.766 | 0.852 | 0.784 | 0.763 | 0.909 |
| | | | 128 | 0.180 | 0.180 | 0.111 | 1.745 | 1.745 | 0.830 | 0.406 | 0.406 | 0.346 |
| | | 0.00400 | 16 | 0.368 | 0.348 | 0.309 | 4.599 | 4.165 | 0.913 | 1.302 | 1.781 | 2.452 |
| | | | 32 | 0.334 | 0.326 | 0.227 | 3.949 | 3.868 | 0.881 | 0.821 | 0.800 | 1.027 |
| | | | 64 | 0.227 | 0.227 | 0.152 | 2.381 | 2.381 | 0.844 | 0.450 | 0.450 | 0.354 |
| | | | 128 | 0.171 | 0.171 | 0.103 | 1.435 | 1.435 | 0.792 | 0.251 | 0.251 | 0.186 |
| | 0.75 | 0.00025 | 16 | 0.409 | 0.360 | 0.341 | 5.019 | 3.879 | 0.912 | 12.661 | 64.500 | 3.411 |
| | | | 32 | 0.363 | 0.327 | 0.246 | 4.524 | 4.054 | 0.870 | 3.579 | 11.801 | 3.091 |
| | | | 64 | 0.268 | 0.284 | 0.169 | 3.057 | 3.136 | 0.850 | 1.943 | 3.675 | 2.350 |
| | | | 128 | 0.172 | 0.162 | 0.111 | 1.611 | 1.542 | 0.868 | 1.027 | 1.044 | 1.183 |
| | | 0.00100 | 16 | 0.414 | 0.528 | 0.342 | 5.168 | 6.600 | 0.907 | 3.016 | 13.297 | 3.385 |
| | | | 32 | 0.328 | 0.296 | 0.242 | 4.215 | 3.801 | 0.872 | 1.820 | 3.087 | 2.432 |
| | | | 64 | 0.250 | 0.247 | 0.164 | 2.774 | 2.674 | 0.861 | 1.038 | 1.057 | 1.283 |
| | | | 128 | 0.181 | 0.180 | 0.112 | 1.764 | 1.751 | 0.823 | 0.531 | 0.527 | 0.473 |
| | | 0.00400 | 16 | 0.426 | 0.403 | 0.340 | 5.261 | 4.958 | 0.907 | 1.490 | 3.125 | 2.970 |
| | | | 32 | 0.326 | 0.306 | 0.235 | 4.011 | 3.745 | 0.882 | 1.023 | 1.036 | 1.276 |
| | | | 64 | 0.241 | 0.241 | 0.158 | 2.555 | 2.555 | 0.837 | 0.560 | 0.560 | 0.462 |
| | | | 128 | 0.169 | 0.169 | 0.106 | 1.425 | 1.425 | 0.793 | 0.300 | 0.300 | 0.229 |
| | 1.0 | 0.00025 | 16 | 0.399 | 0.433 | 0.351 | 4.774 | 5.732 | 0.906 | 13.778 | 99.870 | 3.533 |
| | | | 32 | 0.308 | 0.272 | 0.247 | 4.030 | 3.502 | 0.879 | 4.924 | 20.066 | 3.168 |
| | | | 64 | 0.262 | 0.228 | 0.171 | 3.034 | 2.509 | 0.851 | 1.996 | 5.202 | 2.725 |
| | | | 128 | 0.194 | 0.185 | 0.117 | 1.879 | 1.802 | 0.839 | 1.185 | 1.278 | 1.469 |
| | | 0.00100 | 16 | 0.415 | 0.332 | 0.355 | 5.121 | 3.801 | 0.901 | 4.734 | 21.569 | 3.356 |
| | | | 32 | 0.326 | 0.267 | 0.245 | 4.126 | 3.218 | 0.873 | 2.175 | 5.109 | 2.757 |
| | | | 64 | 0.246 | 0.231 | 0.166 | 2.711 | 2.534 | 0.863 | 1.164 | 1.272 | 1.646 |
| | | | 128 | 0.187 | 0.186 | 0.111 | 1.664 | 1.637 | 0.822 | 0.733 | 0.723 | 0.624 |
| | | 0.00400 | 16 | 0.410 | 0.355 | 0.347 | 4.717 | 3.543 | 0.912 | 1.944 | 4.144 | 2.870 |
| | | | 32 | 0.355 | 0.312 | 0.241 | 4.310 | 3.777 | 0.853 | 1.166 | 1.286 | 1.806 |
| | | | 64 | 0.225 | 0.218 | 0.157 | 2.291 | 2.213 | 0.862 | 0.660 | 0.632 | 0.772 |
| | | | 128 | 0.163 | 0.163 | 0.104 | 1.210 | 1.210 | 0.783 | 0.355 | 0.355 | 0.270 |
| 0.3 | 0.5 | 0.00025 | 16 | 0.382 | 0.359 | 0.325 | 1.280 | 1.156 | 0.870 | 6.561 | 24.834 | 3.324 |
| | | | 32 | 0.296 | 0.270 | 0.250 | 1.098 | 0.960 | 0.853 | 2.519 | 7.045 | 2.835 |
| | | | 64 | 0.242 | 0.236 | 0.183 | 0.919 | 0.893 | 0.802 | 1.463 | 1.774 | 1.705 |
| | | | 128 | 0.178 | 0.175 | 0.132 | 0.737 | 0.730 | 0.747 | 0.754 | 0.745 | 0.751 |
| | | 0.00100 | 16 | 0.371 | 0.382 | 0.326 | 1.230 | 1.022 | 0.873 | 2.120 | 5.936 | 3.128 |
| | | | 32 | 0.313 | 0.316 | 0.247 | 1.130 | 1.120 | 0.850 | 1.584 | 2.018 | 1.836 |
| | | | 64 | 0.259 | 0.258 | 0.184 | 0.980 | 0.974 | 0.775 | 0.779 | 0.775 | 0.721 |
| | | | 128 | 0.157 | 0.157 | 0.124 | 0.667 | 0.667 | 0.715 | 0.392 | 0.392 | 0.313 |
| | | 0.00400 | 16 | 0.353 | 0.352 | 0.319 | 1.213 | 1.174 | 0.876 | 1.473 | 1.856 | 2.021 |
| | | | 32 | 0.324 | 0.322 | 0.240 | 1.174 | 1.164 | 0.823 | 0.820 | 0.810 | 0.822 |
| | | | 64 | 0.220 | 0.220 | 0.163 | 0.813 | 0.813 | 0.761 | 0.416 | 0.416 | 0.338 |
| | | | 128 | 0.156 | 0.156 | 0.114 | 0.613 | 0.613 | 0.599 | 0.245 | 0.245 | 0.179 |
| | 0.75 | 0.00025 | 16 | 0.397 | 0.389 | 0.354 | 1.324 | 0.900 | 0.886 | 9.453 | 43.210 | 3.488 |
| | | | 32 | 0.356 | 0.374 | 0.262 | 1.335 | 1.366 | 0.827 | 3.494 | 12.700 | 3.086 |
| | | | 64 | 0.235 | 0.203 | 0.182 | 0.894 | 0.796 | 0.800 | 1.751 | 2.873 | 2.251 |
| | | | 128 | 0.170 | 0.168 | 0.128 | 0.718 | 0.718 | 0.765 | 1.002 | 1.011 | 1.112 |
| | | 0.00100 | 16 | 0.402 | 0.419 | 0.352 | 1.328 | 1.406 | 0.882 | 3.336 | 9.804 | 3.147 |
| | | | 32 | 0.314 | 0.310 | 0.252 | 1.120 | 1.071 | 0.850 | 1.516 | 2.687 | 2.606 |
| | | | 64 | 0.229 | 0.229 | 0.175 | 0.856 | 0.861 | 0.799 | 1.001 | 1.009 | 1.179 |
| | | | 128 | 0.168 | 0.168 | 0.125 | 0.667 | 0.667 | 0.697 | 0.557 | 0.557 | 0.426 |
| | | 0.00400 | 16 | 0.409 | 0.389 | 0.346 | 1.340 | 1.173 | 0.876 | 1.630 | 2.639 | 2.565 |
| | | | 32 | 0.294 | 0.289 | 0.242 | 1.054 | 1.024 | 0.837 | 0.971 | 0.972 | 1.214 |
| | | | 64 | 0.231 | 0.231 | 0.169 | 0.862 | 0.862 | 0.717 | 0.554 | 0.554 | 0.418 |
| | | | 128 | 0.160 | 0.160 | 0.115 | 0.627 | 0.627 | 0.602 | 0.285 | 0.285 | 0.220 |
| | 1.0 | 0.00025 | 16 | 0.396 | 0.371 | 0.361 | 1.218 | 0.898 | 0.892 | 13.190 | 67.381 | 3.491 |
| | | | 32 | 0.320 | 0.307 | 0.259 | 1.191 | 1.123 | 0.836 | 4.415 | 19.671 | 3.210 |
| | | | 64 | 0.248 | 0.245 | 0.183 | 0.949 | 0.901 | 0.788 | 2.157 | 4.128 | 2.421 |
| | | | 128 | 0.166 | 0.153 | 0.125 | 0.709 | 0.691 | 0.771 | 1.229 | 1.403 | 1.610 |
| | | 0.00100 | 16 | 0.411 | 0.364 | 0.359 | 1.303 | 1.066 | 0.876 | 3.538 | 13.311 | 3.381 |
| | | | 32 | 0.315 | 0.278 | 0.258 | 1.174 | 1.036 | 0.821 | 2.129 | 4.532 | 2.674 |
| | | | 64 | 0.221 | 0.216 | 0.175 | 0.851 | 0.837 | 0.789 | 1.182 | 1.266 | 1.457 |
| | | | 128 | 0.167 | 0.168 | 0.121 | 0.643 | 0.643 | 0.682 | 0.675 | 0.672 | 0.513 |
| | | 0.00400 | 16 | 0.395 | 0.378 | 0.349 | 1.187 | 1.117 | 0.875 | 1.874 | 3.555 | 2.734 |
| | | | 32 | 0.317 | 0.324 | 0.246 | 1.136 | 1.149 | 0.825 | 1.113 | 1.203 | 1.764 |
| | | | 64 | 0.232 | 0.230 | 0.167 | 0.823 | 0.816 | 0.728 | 0.619 | 0.615 | 0.581 |
| | | | 128 | 0.156 | 0.156 | 0.110 | 0.541 | 0.541 | 0.523 | 0.358 | 0.358 | 0.250 |
| 0.5 | 0.5 | 0.00025 | 16 | 0.404 | 0.457 | 0.339 | 0.774 | 0.717 | 0.832 | 5.087 | 15.601 | 3.377 |
| | | | 32 | 0.336 | 0.337 | 0.267 | 0.751 | 0.734 | 0.795 | 2.469 | 5.464 | 2.620 |
| | | | 64 | 0.257 | 0.248 | 0.207 | 0.659 | 0.648 | 0.695 | 1.420 | 1.721 | 1.692 |
| | | | 128 | 0.197 | 0.198 | 0.149 | 0.579 | 0.580 | 0.611 | 0.743 | 0.736 | 0.627 |
| | | 0.00100 | 16 | 0.392 | 0.394 | 0.334 | 0.741 | 0.683 | 0.847 | 2.115 | 5.336 | 2.977 |
| | | | 32 | 0.334 | 0.317 | 0.268 | 0.764 | 0.719 | 0.774 | 1.446 | 1.801 | 1.848 |
| | | | 64 | 0.252 | 0.251 | 0.200 | 0.638 | 0.635 | 0.660 | 0.778 | 0.770 | 0.637 |
| | | | 128 | 0.183 | 0.183 | 0.141 | 0.515 | 0.515 | 0.523 | 0.385 | 0.385 | 0.291 |
| | | 0.00400 | 16 | 0.381 | 0.374 | 0.330 | 0.763 | 0.729 | 0.812 | 1.520 | 1.769 | 1.652 |
| | | | 32 | 0.345 | 0.345 | 0.258 | 0.785 | 0.786 | 0.723 | 0.727 | 0.724 | 0.647 |
| | | | 64 | 0.253 | 0.253 | 0.180 | 0.625 | 0.625 | 0.584 | 0.415 | 0.415 | 0.311 |
| | | | 128 | 0.164 | 0.164 | 0.122 | 0.424 | 0.424 | 0.385 | 0.234 | 0.234 | 0.165 |
| | 0.75 | 0.00025 | 16 | 0.408 | 0.460 | 0.365 | 0.762 | 0.801 | 0.859 | 8.553 | 36.582 | 3.403 |
| | | | 32 | 0.334 | 0.335 | 0.274 | 0.754 | 0.707 | 0.792 | 3.315 | 8.881 | 2.885 |
| | | | 64 | 0.246 | 0.249 | 0.200 | 0.652 | 0.660 | 0.726 | 1.613 | 2.572 | 2.239 |
| | | | 128 | 0.194 | 0.184 | 0.146 | 0.596 | 0.578 | 0.625 | 0.966 | 0.967 | 1.063 |
| | | 0.00100 | 16 | 0.421 | 0.461 | 0.366 | 0.814 | 0.891 | 0.842 | 2.898 | 9.128 | 3.183 |
| | | | 32 | 0.334 | 0.362 | 0.269 | 0.755 | 0.799 | 0.788 | 1.579 | 2.622 | 2.436 |
| | | | 64 | 0.244 | 0.244 | 0.191 | 0.639 | 0.638 | 0.690 | 0.963 | 0.962 | 0.816 |
| | | | 128 | 0.182 | 0.182 | 0.137 | 0.539 | 0.539 | 0.552 | 0.514 | 0.514 | 0.384 |
| | | 0.00400 | 16 | 0.413 | 0.404 | 0.357 | 0.807 | 0.762 | 0.834 | 1.630 | 2.349 | 2.327 |
| | | | 32 | 0.332 | 0.329 | 0.256 | 0.766 | 0.759 | 0.750 | 0.990 | 0.992 | 0.982 |
| | | | 64 | 0.230 | 0.230 | 0.181 | 0.596 | 0.596 | 0.598 | 0.528 | 0.528 | 0.404 |
| | | | 128 | 0.153 | 0.153 | 0.122 | 0.434 | 0.434 | 0.415 | 0.271 | 0.271 | 0.203 |
| | 1.0 | 0.00025 | 16 | 0.459 | 0.457 | 0.373 | 0.829 | 0.578 | 0.852 | 10.157 | 32.986 | 3.507 |
| | | | 32 | 0.327 | 0.322 | 0.274 | 0.737 | 0.718 | 0.785 | 3.823 | 15.552 | 3.192 |
| | | | 64 | 0.253 | 0.240 | 0.196 | 0.660 | 0.632 | 0.721 | 1.957 | 3.655 | 2.418 |
| | | | 128 | 0.181 | 0.181 | 0.141 | 0.566 | 0.557 | 0.618 | 1.144 | 1.196 | 1.237 |
| | | 0.00100 | 16 | 0.428 | 0.421 | 0.369 | 0.768 | 0.642 | 0.851 | 3.528 | 11.730 | 3.258 |
| | | | 32 | 0.364 | 0.358 | 0.271 | 0.810 | 0.772 | 0.767 | 1.789 | 3.418 | 2.647 |
| | | | 64 | 0.249 | 0.249 | 0.191 | 0.646 | 0.637 | 0.668 | 1.155 | 1.235 | 1.460 |
| | | | 128 | 0.170 | 0.170 | 0.130 | 0.531 | 0.531 | 0.535 | 0.639 | 0.639 | 0.445 |
| | | 0.00400 | 16 | 0.423 | 0.419 | 0.370 | 0.821 | 0.788 | 0.833 | 1.729 | 3.328 | 2.846 |
| | | | 32 | 0.322 | 0.308 | 0.257 | 0.726 | 0.697 | 0.721 | 1.070 | 1.097 | 1.333 |
| | | | 64 | 0.223 | 0.223 | 0.172 | 0.558 | 0.558 | 0.543 | 0.575 | 0.575 | 0.460 |
| | | | 128 | 0.172 | 0.172 | 0.117 | 0.434 | 0.434 | 0.356 | 0.330 | 0.330 | 0.235 |

**Table S10:** Simulation results for $\psi$ with $k = 4$.

| | | | | $\lambda$ | | | $\mu$ | | | $\theta$ | | |
|---|---|---|---|---|---|---|---|---|---|---|---|---|
| $\epsilon$ | $\rho$ | $\theta$ | $N$ | RMSE | RMSE$_c$ | CV | RMSE | RMSE$_c$ | CV | RMSE | RMSE$_c$ | CV |
| 0.1 | 0.5 | 0.00025 | 16 | 0.423 | 0.378 | 0.342 | 5.380 | 4.995 | 0.940 | 7.346 | 32.275 | 3.354 |
| | | | 32 | 0.325 | 0.297 | 0.239 | 4.166 | 3.861 | 0.890 | 3.272 | 7.628 | 2.643 |
| | | | 64 | 0.255 | 0.238 | 0.167 | 2.941 | 2.797 | 0.861 | 1.483 | 1.984 | 1.945 |
| | | | 128 | 0.176 | 0.173 | 0.112 | 1.759 | 1.743 | 0.847 | 0.785 | 0.774 | 0.801 |
| | | 0.00100 | 16 | 0.384 | 0.391 | 0.335 | 4.847 | 5.112 | 0.953 | 2.370 | 7.176 | 3.138 |
| | | | 32 | 0.341 | 0.306 | 0.241 | 4.307 | 3.828 | 0.886 | 1.320 | 1.671 | 2.121 |
| | | | 64 | 0.257 | 0.242 | 0.164 | 2.937 | 2.774 | 0.852 | 0.786 | 0.766 | 0.910 |
| | | | 128 | 0.180 | 0.180 | 0.111 | 1.745 | 1.745 | 0.828 | 0.407 | 0.407 | 0.346 |
| | | 0.00400 | 16 | 0.401 | 0.367 | 0.332 | 4.989 | 4.398 | 0.956 | 1.301 | 1.779 | 2.457 |
| | | | 32 | 0.336 | 0.328 | 0.228 | 3.977 | 3.903 | 0.887 | 0.821 | 0.800 | 1.018 |
| | | | 64 | 0.226 | 0.226 | 0.152 | 2.379 | 2.379 | 0.844 | 0.450 | 0.450 | 0.354 |
| | | | 128 | 0.171 | 0.171 | 0.103 | 1.432 | 1.432 | 0.790 | 0.251 | 0.251 | 0.186 |
| | 0.75 | 0.00025 | 16 | 0.422 | 0.356 | 0.347 | 5.163 | 3.811 | 0.922 | 12.617 | 64.308 | 3.412 |
| | | | 32 | 0.363 | 0.329 | 0.247 | 4.513 | 4.062 | 0.871 | 3.568 | 11.785 | 3.089 |
| | | | 64 | 0.268 | 0.284 | 0.170 | 3.065 | 3.137 | 0.852 | 1.944 | 3.669 | 2.357 |
| | | | 128 | 0.172 | 0.163 | 0.111 | 1.614 | 1.544 | 0.869 | 1.029 | 1.047 | 1.186 |
| | | 0.00100 | 16 | 0.426 | 0.542 | 0.350 | 5.299 | 6.742 | 0.919 | 3.017 | 13.119 | 3.414 |
| | | | 32 | 0.329 | 0.298 | 0.243 | 4.230 | 3.821 | 0.875 | 1.826 | 3.103 | 2.428 |
| | | | 64 | 0.250 | 0.247 | 0.164 | 2.775 | 2.675 | 0.863 | 1.035 | 1.054 | 1.282 |
| | | | 128 | 0.181 | 0.180 | 0.112 | 1.761 | 1.749 | 0.822 | 0.531 | 0.527 | 0.473 |
| | | 0.00400 | 16 | 0.438 | 0.407 | 0.348 | 5.392 | 5.014 | 0.921 | 1.491 | 3.120 | 2.981 |
| | | | 32 | 0.325 | 0.305 | 0.235 | 4.000 | 3.733 | 0.882 | 1.028 | 1.042 | 1.283 |
| | | | 64 | 0.241 | 0.241 | 0.158 | 2.561 | 2.561 | 0.840 | 0.561 | 0.561 | 0.462 |
| | | | 128 | 0.170 | 0.170 | 0.106 | 1.428 | 1.428 | 0.794 | 0.301 | 0.301 | 0.229 |
| | 1.0 | 0.00025 | 16 | 0.405 | 0.429 | 0.356 | 4.838 | 5.743 | 0.914 | 13.761 | 100.135 | 3.511 |
| | | | 32 | 0.309 | 0.271 | 0.247 | 4.028 | 3.480 | 0.876 | 4.931 | 19.916 | 3.175 |
| | | | 64 | 0.262 | 0.229 | 0.172 | 3.036 | 2.516 | 0.852 | 1.993 | 5.209 | 2.706 |
| | | | 128 | 0.194 | 0.185 | 0.117 | 1.876 | 1.798 | 0.836 | 1.182 | 1.274 | 1.463 |
| | | 0.00100 | 16 | 0.420 | 0.332 | 0.358 | 5.186 | 3.797 | 0.907 | 4.744 | 21.591 | 3.366 |
| | | | 32 | 0.325 | 0.266 | 0.245 | 4.111 | 3.202 | 0.871 | 2.186 | 5.147 | 2.770 |
| | | | 64 | 0.246 | 0.232 | 0.166 | 2.707 | 2.534 | 0.865 | 1.169 | 1.279 | 1.644 |
| | | | 128 | 0.187 | 0.186 | 0.111 | 1.665 | 1.639 | 0.824 | 0.733 | 0.723 | 0.628 |
| | | 0.00400 | 16 | 0.415 | 0.357 | 0.350 | 4.775 | 3.589 | 0.915 | 1.961 | 4.214 | 2.872 |
| | | | 32 | 0.355 | 0.312 | 0.242 | 4.320 | 3.785 | 0.854 | 1.164 | 1.283 | 1.797 |
| | | | 64 | 0.224 | 0.217 | 0.157 | 2.278 | 2.200 | 0.860 | 0.659 | 0.631 | 0.767 |
| | | | 128 | 0.164 | 0.164 | 0.104 | 1.214 | 1.214 | 0.783 | 0.354 | 0.354 | 0.269 |
| 0.3 | 0.5 | 0.00025 | 16 | 0.431 | 0.393 | 0.357 | 1.489 | 1.317 | 0.918 | 6.474 | 24.668 | 3.329 |
| | | | 32 | 0.300 | 0.272 | 0.254 | 1.115 | 0.968 | 0.863 | 2.529 | 7.060 | 2.841 |
| | | | 64 | 0.241 | 0.235 | 0.183 | 0.915 | 0.890 | 0.800 | 1.461 | 1.770 | 1.712 |
| | | | 128 | 0.179 | 0.175 | 0.132 | 0.738 | 0.731 | 0.747 | 0.754 | 0.745 | 0.750 |
| | | 0.00100 | 16 | 0.409 | 0.402 | 0.357 | 1.409 | 1.127 | 0.922 | 2.101 | 5.882 | 3.124 |
| | | | 32 | 0.320 | 0.322 | 0.251 | 1.155 | 1.141 | 0.856 | 1.581 | 2.012 | 1.836 |
| | | | 64 | 0.259 | 0.258 | 0.184 | 0.980 | 0.975 | 0.776 | 0.782 | 0.778 | 0.724 |
| | | | 128 | 0.158 | 0.158 | 0.124 | 0.668 | 0.668 | 0.715 | 0.393 | 0.393 | 0.314 |
| | | 0.00400 | 16 | 0.382 | 0.374 | 0.342 | 1.342 | 1.276 | 0.916 | 1.472 | 1.853 | 2.035 |
| | | | 32 | 0.327 | 0.326 | 0.242 | 1.188 | 1.178 | 0.829 | 0.821 | 0.812 | 0.820 |
| | | | 64 | 0.219 | 0.219 | 0.163 | 0.811 | 0.811 | 0.761 | 0.416 | 0.416 | 0.338 |
| | | | 128 | 0.156 | 0.156 | 0.114 | 0.613 | 0.613 | 0.598 | 0.246 | 0.246 | 0.179 |
| | 0.75 | 0.00025 | 16 | 0.412 | 0.393 | 0.365 | 1.385 | 0.926 | 0.903 | 9.305 | 42.133 | 3.459 |
| | | | 32 | 0.356 | 0.376 | 0.262 | 1.337 | 1.376 | 0.826 | 3.463 | 12.669 | 3.070 |
| | | | 64 | 0.234 | 0.202 | 0.181 | 0.894 | 0.797 | 0.800 | 1.740 | 2.847 | 2.237 |
| | | | 128 | 0.170 | 0.168 | 0.128 | 0.718 | 0.718 | 0.763 | 1.002 | 1.012 | 1.107 |
| | | 0.00100 | 16 | 0.417 | 0.436 | 0.361 | 1.384 | 1.468 | 0.897 | 3.322 | 9.669 | 3.143 |
| | | | 32 | 0.315 | 0.311 | 0.252 | 1.122 | 1.075 | 0.853 | 1.519 | 2.695 | 2.607 |
| | | | 64 | 0.228 | 0.229 | 0.175 | 0.855 | 0.861 | 0.796 | 0.999 | 1.007 | 1.175 |
| | | | 128 | 0.168 | 0.168 | 0.125 | 0.669 | 0.669 | 0.696 | 0.555 | 0.555 | 0.425 |
| | | 0.00400 | 16 | 0.424 | 0.396 | 0.355 | 1.396 | 1.203 | 0.890 | 1.620 | 2.611 | 2.570 |
| | | | 32 | 0.294 | 0.289 | 0.242 | 1.057 | 1.025 | 0.839 | 0.970 | 0.970 | 1.211 |
| | | | 64 | 0.231 | 0.231 | 0.169 | 0.864 | 0.864 | 0.717 | 0.553 | 0.553 | 0.417 |
| | | | 128 | 0.160 | 0.160 | 0.115 | 0.627 | 0.627 | 0.603 | 0.285 | 0.285 | 0.220 |
| | 1.0 | 0.00025 | 16 | 0.401 | 0.369 | 0.366 | 1.243 | 0.896 | 0.899 | 13.097 | 67.179 | 3.492 |
| | | | 32 | 0.320 | 0.309 | 0.259 | 1.191 | 1.125 | 0.833 | 4.406 | 19.769 | 3.220 |
| | | | 64 | 0.248 | 0.245 | 0.183 | 0.947 | 0.898 | 0.788 | 2.164 | 4.136 | 2.417 |
| | | | 128 | 0.166 | 0.153 | 0.126 | 0.709 | 0.690 | 0.772 | 1.231 | 1.405 | 1.623 |
| | | 0.00100 | 16 | 0.420 | 0.366 | 0.363 | 1.335 | 1.077 | 0.883 | 3.569 | 13.571 | 3.409 |
| | | | 32 | 0.316 | 0.279 | 0.258 | 1.178 | 1.043 | 0.821 | 2.126 | 4.547 | 2.642 |
| | | | 64 | 0.221 | 0.217 | 0.174 | 0.850 | 0.839 | 0.788 | 1.180 | 1.264 | 1.461 |
| | | | 128 | 0.167 | 0.167 | 0.120 | 0.642 | 0.642 | 0.682 | 0.675 | 0.672 | 0.512 |
| | | 0.00400 | 16 | 0.400 | 0.381 | 0.353 | 1.208 | 1.131 | 0.884 | 1.877 | 3.567 | 2.735 |
| | | | 32 | 0.317 | 0.324 | 0.246 | 1.135 | 1.147 | 0.826 | 1.115 | 1.205 | 1.774 |
| | | | 64 | 0.232 | 0.230 | 0.167 | 0.825 | 0.818 | 0.731 | 0.617 | 0.613 | 0.579 |
| | | | 128 | 0.156 | 0.156 | 0.110 | 0.542 | 0.542 | 0.524 | 0.358 | 0.358 | 0.250 |
| 0.5 | 0.5 | 0.00025 | 16 | 0.443 | 0.461 | 0.376 | 0.876 | 0.738 | 0.885 | 4.969 | 15.509 | 3.374 |
| | | | 32 | 0.345 | 0.344 | 0.275 | 0.772 | 0.750 | 0.807 | 2.470 | 5.464 | 2.631 |
| | | | 64 | 0.258 | 0.248 | 0.207 | 0.662 | 0.649 | 0.697 | 1.428 | 1.733 | 1.696 |
| | | | 128 | 0.197 | 0.198 | 0.149 | 0.577 | 0.578 | 0.612 | 0.743 | 0.736 | 0.625 |
| | | 0.00100 | 16 | 0.422 | 0.405 | 0.369 | 0.819 | 0.726 | 0.899 | 2.082 | 5.229 | 2.976 |
| | | | 32 | 0.347 | 0.324 | 0.277 | 0.792 | 0.737 | 0.788 | 1.445 | 1.799 | 1.853 |
| | | | 64 | 0.252 | 0.251 | 0.199 | 0.638 | 0.635 | 0.659 | 0.776 | 0.767 | 0.637 |
| | | | 128 | 0.183 | 0.183 | 0.141 | 0.515 | 0.515 | 0.523 | 0.385 | 0.385 | 0.291 |
| | | 0.00400 | 16 | 0.415 | 0.400 | 0.359 | 0.849 | 0.803 | 0.857 | 1.497 | 1.736 | 1.665 |
| | | | 32 | 0.351 | 0.352 | 0.262 | 0.799 | 0.800 | 0.731 | 0.726 | 0.723 | 0.650 |
| | | | 64 | 0.254 | 0.254 | 0.181 | 0.627 | 0.627 | 0.586 | 0.414 | 0.414 | 0.311 |
| | | | 128 | 0.163 | 0.163 | 0.122 | 0.423 | 0.423 | 0.383 | 0.234 | 0.234 | 0.165 |
| | 0.75 | 0.00025 | 16 | 0.424 | 0.485 | 0.380 | 0.801 | 0.851 | 0.881 | 8.590 | 36.604 | 3.404 |
| | | | 32 | 0.335 | 0.336 | 0.275 | 0.756 | 0.709 | 0.794 | 3.312 | 8.892 | 2.885 |
| | | | 64 | 0.245 | 0.247 | 0.199 | 0.650 | 0.655 | 0.723 | 1.620 | 2.589 | 2.241 |
| | | | 128 | 0.194 | 0.184 | 0.146 | 0.596 | 0.578 | 0.624 | 0.963 | 0.964 | 1.065 |
| | | 0.00100 | 16 | 0.441 | 0.492 | 0.382 | 0.863 | 0.963 | 0.865 | 2.884 | 9.093 | 3.190 |
| | | | 32 | 0.335 | 0.362 | 0.270 | 0.757 | 0.801 | 0.789 | 1.580 | 2.624 | 2.439 |
| | | | 64 | 0.244 | 0.244 | 0.191 | 0.639 | 0.638 | 0.689 | 0.958 | 0.958 | 0.811 |
| | | | 128 | 0.182 | 0.182 | 0.137 | 0.539 | 0.539 | 0.551 | 0.516 | 0.516 | 0.385 |
| | | 0.00400 | 16 | 0.429 | 0.411 | 0.370 | 0.847 | 0.780 | 0.855 | 1.626 | 2.341 | 2.346 |
| | | | 32 | 0.331 | 0.329 | 0.256 | 0.766 | 0.759 | 0.752 | 0.989 | 0.991 | 0.982 |
| | | | 64 | 0.230 | 0.230 | 0.181 | 0.599 | 0.599 | 0.600 | 0.529 | 0.529 | 0.404 |
| | | | 128 | 0.153 | 0.153 | 0.122 | 0.435 | 0.435 | 0.414 | 0.271 | 0.271 | 0.203 |
| | 1.0 | 0.00025 | 16 | 0.474 | 0.461 | 0.381 | 0.864 | 0.581 | 0.864 | 10.050 | 33.492 | 3.517 |
| | | | 32 | 0.329 | 0.328 | 0.276 | 0.742 | 0.736 | 0.788 | 3.828 | 15.704 | 3.169 |
| | | | 64 | 0.253 | 0.240 | 0.197 | 0.659 | 0.631 | 0.720 | 1.956 | 3.662 | 2.401 |
| | | | 128 | 0.181 | 0.181 | 0.141 | 0.565 | 0.557 | 0.617 | 1.147 | 1.200 | 1.246 |
| | | 0.00100 | 16 | 0.437 | 0.424 | 0.377 | 0.788 | 0.647 | 0.863 | 3.551 | 11.807 | 3.268 |
| | | | 32 | 0.366 | 0.360 | 0.272 | 0.814 | 0.778 | 0.767 | 1.789 | 3.417 | 2.638 |
| | | | 64 | 0.249 | 0.249 | 0.191 | 0.645 | 0.636 | 0.664 | 1.155 | 1.236 | 1.455 |
| | | | 128 | 0.170 | 0.170 | 0.130 | 0.531 | 0.531 | 0.536 | 0.640 | 0.640 | 0.445 |
| | | 0.00400 | 16 | 0.435 | 0.426 | 0.378 | 0.850 | 0.805 | 0.847 | 1.728 | 3.345 | 2.844 |
| | | | 32 | 0.322 | 0.307 | 0.257 | 0.725 | 0.695 | 0.720 | 1.072 | 1.100 | 1.330 |
| | | | 64 | 0.224 | 0.224 | 0.172 | 0.558 | 0.558 | 0.543 | 0.575 | 0.575 | 0.460 |
| | | | 128 | 0.172 | 0.172 | 0.117 | 0.435 | 0.435 | 0.356 | 0.329 | 0.329 | 0.235 |
